# Quantifying Cross-Modal Shared Information Between Histomorphology and Spatial Transcriptomics via Spatiotemporal Trajectory Correlation

**DOI:** 10.64898/2026.07.13.738201

**Authors:** Xuanxuan He, Ming Feng, Anqi Wang, Xuan Huang, Xiaobang Luo, Xinwang Liu, Tao Sun, Lifei Wang, Kele Xu

**Affiliations:** College of Computer Science and Technology, National University of Defense Technology, Changsha, China; National Key Laboratory of Parallel and Distributed Computing, National University of Defense Technology, Changsha, China; Key Laboratory of Artificial Organs and Computational Medicine of Zhejiang Province, Shulan (Hangzhou) Hospital, Shulan International Medical College, Zhejiang Shuren University, Hangzhou, China; Research Center for Blood Engineering and Manufacturing, Cyrus Tang Medical Institute, Socchow University, Suzhou, China

**Keywords:** Dual-modality Analysis of Spatial Transcriptomics and Histopathological Images, Trajectory Reconstruction and Trajectory Enriched Genes, Pathology Foundation Models, Cross-modal Information Sharing

## Abstract

Histopathological imaging and spatial transcriptomics (ST) provide synergistic morphological and molecular insights into tissue architecture. While conventional downstream analyses predominantly adopt a discrete paradigm, such as segmenting histological images or identifying spatial domains, emerging trajectory reconstruction methods offer a continuous perspective for analyzing these modalities. However, most current studies are confined to single-modality or single-organ analyses, lacking systematic integration across modalities and multiple organs. To address this limitation, we performed trajectory reconstruction on ST-derived gene expression data and on histopathological morphological features extracted using ten widely used pathology pretrained models across multiple cancer samples from six organs. The results demonstrate that trajectory reconstruction, as a continuous analytical framework, effectively bridges spatial transcriptomics and histopathological imaging. More importantly, we propose an innovative framework that uses trajectory pseudotime as a mediating variable to quantify the extent of information sharing between molecular and morphological features. This framework not only provides a new perspective for understanding the intrinsic links between modalities but also establishes a solid theoretical and methodological foundation for future cross-modal translation studies.

## Introduction

Histopathological image analysis remains the cornerstone for cancer diagnosis, subtyping, and prognostic stratification, playing an indispensable role in routine clinical practice. Recent advances in artificial intelligence (AI), widespread digitalization of whole-slide images (WSIs), and the availability of large-scale annotated datasets have enabled the development of specialized deep neural network models tailored to clinical applications, including cancer subtype classification, staging, diagnostic prediction, and prognostic evaluation [1–3]. Furthermore, the emergence of self-supervised learning has catalyzed the pre-training of pathology foundation models, which learn robust representations from vast amounts of unlabeled tissue data [4]. These models typically follow a workflow where large images are partitioned into patches and converted into digitized vector representations that encapsulate complex visual information. In contrast to traditional handcrafted features based on texture and color, features extracted by pre-trained self-supervised foundation models exhibit superior generalization and representational capacity. These digitized vector representations of histopathological image patches have been widely adopted across numerous downstream tasks [5–9]. For instance, histopathological image segmentation can be performed by leveraging similarities among patch-level feature vectors to identify tissue architectures [10]. Furthermore, clustering of patch representations from multiple slides of a given cancer subtype can reveal shared histomorphological phenotypes characteristic of that cancer type [11].

Complementary to the morphological information captured in histopathological images, spatial transcriptomics (ST) provides an orthogonal molecular modality. By integrating high-throughput sequencing with spatial coordinate systems, ST technologies simultaneously capture genome-wide gene expression profiles and spatial positional information in intact tissue sections [12, 13]. The widely used 10x Genomics Visium platform, for example, uses spatially barcoded oligonucleotide arrays to partition tissue sections into capture spots (55 μm diameter, 100 μm center-to-center spacing). Each spot simultaneously measures expression levels of thousands of genes, with barcode sequences mapping expression data back to their original spatial locations [14]. This and related technologies have empowered researchers to resolve molecular heterogeneity in situ, identify gene expression signatures of functionally distinct tissue domains, and even dissect intercellular signaling networks and cellular crosstalk [15–18]. For example, spatially variable gene detection identifies genes whose expression exhibits non-random spatial distributions, thereby revealing tissue spatial heterogeneity and defining functional domains [19]. Spatial domain identification, by integrating gene expression and spatial location, partitions tissues into biologically meaningful spatially coherent regions [20].

Histopathological images and spatial transcriptomic data are inherently paired. In standard ST workflows, tissue sections are first processed for H&E staining and imaging before permeabilization, mRNA capture, and sequencing, enabling precise alignment between each ST spot and its corresponding histopathological image patch. This intrinsic pairing provides a solid foundation for multimodal integration and cross-modality translation. Given the abundance and low cost of archived histopathological data, coupled with the rich molecular detail provided by spatial transcriptomics (ST), morphology translation (predicting gene expression from histology images) has emerged as a vibrant area of research. Methods such as ST-Net, Hist2ST, iStar, and GHIST leverage Transformers, graph neural networks (GNNs), and multi-scale feature extraction to map histomorphological features onto virtual ST profiles, enabling local or whole-slide prediction of gene expression [21–24]. However, morphology translation critically depends on the design of morphological feature extractors and the shared information between extracted features and gene expression. To date, systematic comparisons of shared cross-modal information between diverse morphological feature extraction strategies and spatial transcriptomic modalities remain limited [25].

Single-cell trajectory analysis reconstructs continuous dynamic trajectories from discrete cellular snapshots using pseudotime inference algorithms (e.g., Monocle, Slingshot, PAGA) applied to single-cell RNA sequencing (scRNA-seq) or multi-omic profiles [26–28]. In this framework, each cell is mapped to a position in a low-dimensional latent space and assigned a pseudotime value based on progressive changes in its gene expression program. Widely applied in developmental biology, this approach has successfully recapitulated cellular differentiation from pluripotent stem cells to specialized lineages and has been used to dissect tumor heterogeneity and evolutionary trajectories [29]. Recently, trajectory analysis has been extended to H&E histopathological imaging and spatial transcriptomics [30]. In a prostate cancer study, ST spots replaced single cells as the analytical unit; trajectory algorithms organized spots into a linear, quantifiable ‘malignant progression trajectory’ based on expression similarity, enabling systematic identification of genes associated with core disease progression programs [31]. Similarly, in a lung adenocarcinoma study, whole H&E slides were partitioned into patches, and feature vectors extracted by a fine-tuned pathological foundation model (Phikon) were used as surrogates for transcriptomic profiles to infer pseudotime, enabling spatially resolved exploration of tumor differentiation trajectories [32].

However, existing trajectory analyses remain largely restricted to single organs and single modalities, lacking large-scale multi-organ, multi-modal investigations. To fill this gap, this study conducted dual-trajectory analyses based on ST and morphology on various cancer samples from colorectal, prostate, breast, lymph node, skin, and brain tissues [33]. ST-based trajectories successfully enriched genes specific to peritumoral and tumor regions, and revealed a transformation trajectory from stroma to malignant epithelium in epithelial adenoma samples. For histomorphology-based analysis, we extracted features using ten mainstream pathology and general visual models [5, 6, 8, 9, 34–37], then systematically quantified correlations between ST-derived trajectories and those generated from each model’s morphological features. Despite not being explicitly designed for morphology-to-ST translation, we observed that larger, more advanced models trained on larger pre-training datasets generated morphological trajectories highly correlated with ST-derived trajectories, indicating that these models capture greater shared information between morphology and gene expression. These findings were supported by trajectory-enriched gene analysis and morphology-to-transcriptome translation performance.

## Results

### Overview of the Analytical Pipeline

Following previously established methods [30–32], we computed pseudotime for each ST sample and its matched H&E histopathological image. Briefly, ST pseudotime was inferred from spot-level gene expression profiles. For H&E images, we adopted two patch extraction strategies: fixed-resolution patches and spot-aligned patches. These patches were subsequently transformed into image feature vectors using ten mainstream pathology and general visual models. Pseudotime trajectories were independently inferred from these image feature vectors (model-extracted histomorphological feature vectors) for each sample. For root node selection, we first identified the starting cluster by minimizing the mean tumor cell proportion and maximizing the spot scale. The root spot was then defined as the one with the minimal tumor proportion within this cluster, using the first diffusion map component as the final tie-breaking metric to ensure the root node was derived, as much as possible, from non-tumor tissue. After root determination, a k-nearest neighbor (KNN) graph was constructed from feature vectors (gene expression for ST; model-extracted histomorphological feature for H&E). Using a diffusion map algorithm initiated from the root node, we computed diffusion components for each spot, generating gene-based and image-based pseudotime orderings. We then calculated correlations between ST-derived pseudotime trajectories and those generated from each pathological foundation model for every sample (Methods) (Fig. 1a).

**Fig. 1.**
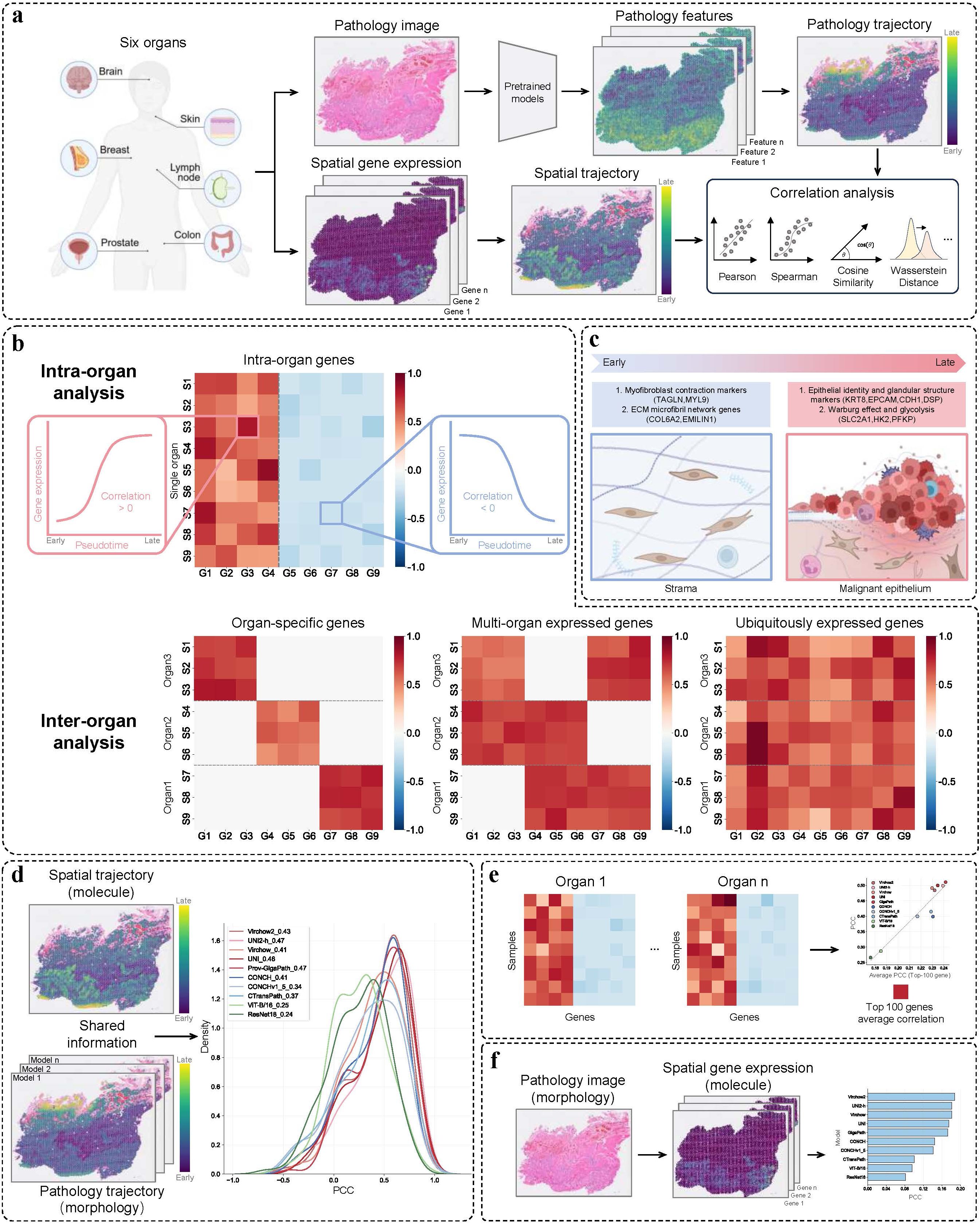
| Integrated framework for dual-trajectory analysis of morphology and spatial transcriptomics. a. Pathological foundation models were used to extract morphological representations from histopathological images and reconstruct morphological trajectories, while trajectories were independently constructed from spatial transcriptomic data. Correlations between morphology-derived trajectories and molecular trajectories were then quantified. b. Gene enrichment analysis in early and late pseudotime along spatial transcriptome-derived trajectories, followed by cross-tissue comparisons to identify shared and tissue-specific gene signatures. c. Early trajectory stages were significantly enriched for stromal cell-related gene sets, whereas late stages were enriched for malignant epithelial cell-related gene sets, demonstrating that trajectory analysis recapitulates the spatiotemporal pattern from stroma to malignant regions in epithelial adenocarcinomas. Created with BioRender.com. d. Systematic evaluation of correlation distributions between morphological trajectories derived from ten models and spatial transcriptome-based molecular trajectories across multiple samples from six organs. This correlation is used to quantify the degree of information sharing between molecular and morphological features. e. Model-generated trajectories exhibit a stronger correlation with spatial transcriptomics (ST)-derived trajectories as the monotonicity of enriched gene expression dynamics increases along the pseudotime axis. f. The morphology-to-transcriptome cross-modal prediction task shows that higher correlation between morphological and molecular trajectories corresponds to improved accuracy in reconstructing spatial gene expression patterns.

For each gene, spearman correlation coefficients were computed between its expression vector and the pseudotime vector for each sample [31]. A correlation coefficient approaching −1 indicated early enrichment and progressive downregulation, suggestive of normal tissue association. Conversely, a coefficient approaching 1 indicated low early expression and progressive upregulation, suggestive of tumor association. We designated these two classes as early- and late-trajectory enriched genes, respectively. For all samples from the same organ, we calculated the top 100 or top 50 organ-enriched early- and late-trajectory genes, then filtered out samples whose correlation coefficients were inconsistent with most samples. Following sample filtering, further analysis of early- and late-trajectory gene sets enabled the identification of pan-tissue and tissue-specific genes (Fig. 1b). Functional characterization revealed that late-trajectory genes were associated with malignant programs, while early-trajectory genes in adenocarcinomas were enriched for stromal signatures, illustrating a stromal-to-malignant epithelial transition and validating the effectiveness of trajectory analysis for ST data (Fig. 1c).

We next quantified the distribution of correlations between ST-based trajectories and morphology-based trajectories across multiple organs and samples. Through this dual trajectory construction and correlation analysis, we aimed to characterize the extent of shared information between histomorphological features and transcriptomic profiles at the level of trajectory inference (Fig. 1d). These findings were corroborated by trajectory-enriched gene analysis and morphology-to-transcriptome translation performance (Fig. 1e, f).

### ST-Based Trajectory Inference and Identification of Early/Late Trajectory-Enriched Genes

We conducted large-scale pseudotime trajectory analysis across multiple cancer types derived from colorectal, prostate, breast, lymph node, skin, and brain tissues, utilizing ST expression vectors. By quantifying dynamic correlations between gene expression and pseudotime, we systematically identified gene modules enriched in early and late trajectory phases for each sample.

In tissue-specific analyses, late pseudotime phases were universally hallmarked by core malignant programs. In colorectal cancer, late stages were strongly enriched for diagnostic markers (e.g., *CEACAM5*, *EPCAM*) and genes governing metastatic invasion, stemness, the Warburg effect, proliferation, antioxidant defense, and chemoresistance. By contrast, early stages were dominated by extracellular matrix (ECM) genes (e.g., *COLs*) and myofibroblast markers (e.g., *ACTA2*). In prostate cancer, late pseudotime was characterized by lineage-defining markers (e.g., *KLK3*, *EPCAM*) and genes involved in androgen signaling, lipid metabolism, protein homeostasis, and therapy resistance. Early stages preferentially expressed stromal and smooth muscle regulators (*ACTA2*, *MYH11*, *DES*, *TPM1/2*, *CALD1*) that mediate ECM assembly, contractility, and mechanotransduction, consistent with previous reports [31]. In breast cancer, late stages were enriched for luminal markers (*ESR1*, *GATA3*) and genes driving proliferation, epithelial differentiation, metabolic reprogramming, and stress tolerance, whereas early stages were marked by HLA molecules, immune genes, and vascular network factors supporting immune surveillance and metabolic homeostasis. In lymph node metastases of renal cell carcinoma, late stages featured renal tubular injury markers (e.g., *LRP2*) [38], hypoxia adaptation, epithelial-mesenchymal transition (EMT), and invasive programs, while early stages were defined by vascular patterning (*CLDN5*, *PECAM1*), dense ECM deposition, and lymphocyte homing (Fig. 2a; Figs. S1-S4). Additionally, late-stage squamous cell carcinoma was enriched for hyperproliferative keratins (e.g., *KRT6A/B/C*), pro-inflammatory *S100* family members, and gap junction components; late-stage melanoma for *SOX10*-driven melanogenesis (*TYR*, *PMEL*, etc.) and lipid metabolism genes; and late-stage brain cancer samples for hypoxia inducers and *VEGFA*-mediated aberrant angiogenesis (Fig. 2b; Fig. S5). Collectively, these results establish that ST pseudotime analysis robustly identifies key genes linked to cancer progression and stromal identity across diverse tissue contexts.

**Fig. 2.**
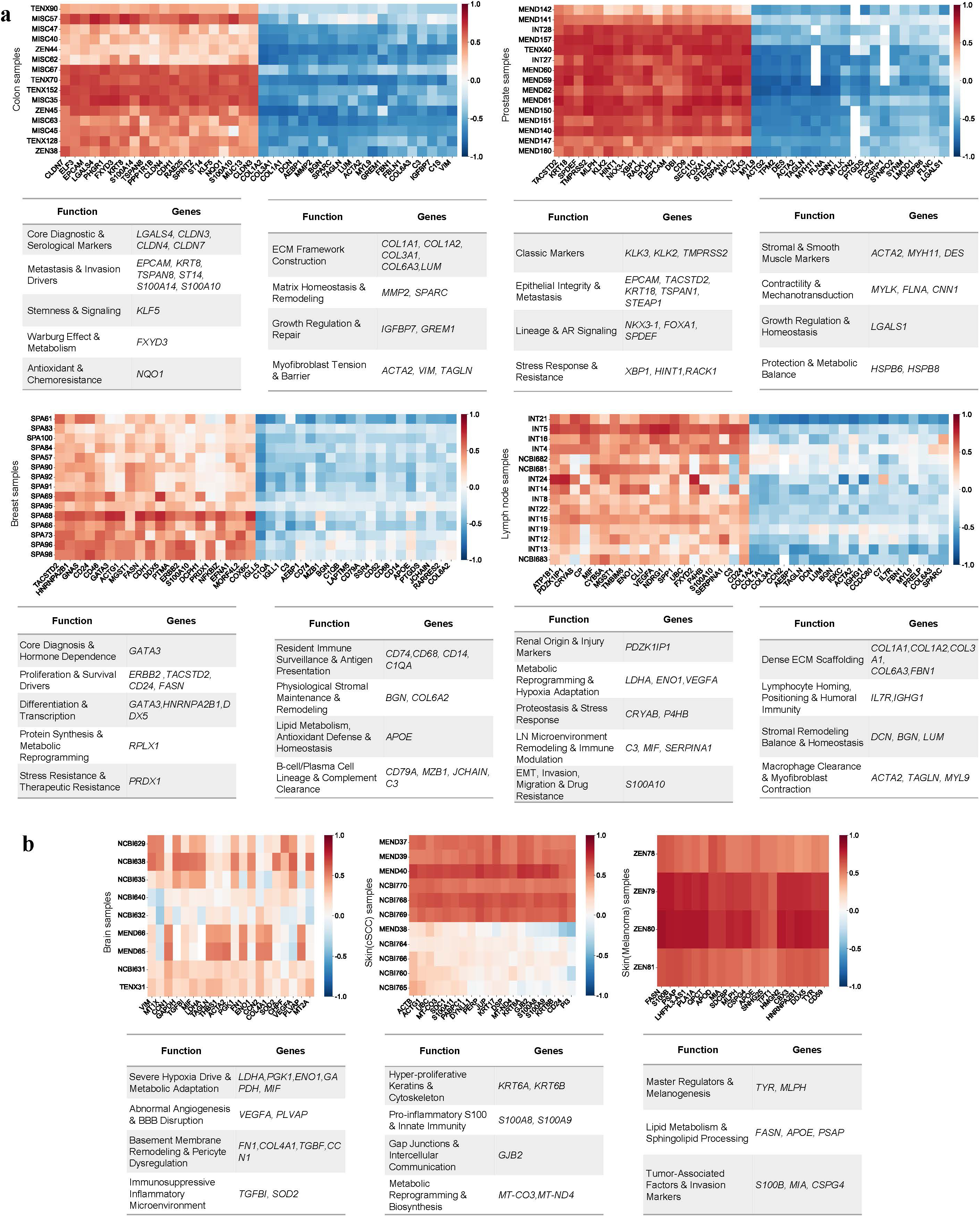
| Spatial transcriptome-derived trajectories enrich adjacent normal- and tumor-associated genes in early and late pseudotime, respectively. a. Top 20 genes most strongly positively (late trajectory stages) or negatively (early trajectory stages) correlated with pseudotime and their functional annotations enriched along spatial transcriptome-derived trajectories in colorectal, prostate, breast, and lymph node samples. The heatmap illustrates correlation strength: deep red signifies strong positive correlation, deep blue signifies strong negative correlation, and white indicates no correlation. b. Top 20 genes most strongly positively (late trajectory stages) correlated with pseudotime and their functional annotations enriched along spatial transcriptome-derived trajectories in brain and skin (melanoma, squamous cell carcinoma) samples. The staining logic is depicted in a.

Pan-cancer integration revealed that recovered gene signatures comprised both pan-tumor and tissue-restricted markers, with epithelial adenocarcinomas (colorectal, prostate, breast, and renal lymph node metastases) displaying marked molecular convergence. Among late-enriched genes, canonical epithelial identity and glandular structural markers (*KRT8*, *KRT19*, *EPCAM*, *DSP*) as well as tight junction stability factors (*CLDN3*, *CLDN4*, *CLDN7*) were predominantly detected in epithelial malignancies. Concomitantly, genes driving proliferative signaling, stemness, and protease activity (*TMPRSS2*, *ERBB3*, *TPD52*, *CD24*, *SPINT2*) were highly expressed in these samples. By contrast, metabolic and stress-associated genes governing the Warburg effect, glycolysis (*SLC2A1*, *HK2*, *PFKP*), angiogenesis (*VEGFA*), ER stress, and autophagy (*CANX*, *VMP1*, *YWHAZ*, etc.) were broadly enriched across most organs, extending beyond epithelial tumors (Fig. 3a). Tissue-specific genes exhibited strict lineage restriction, encompassing intestinal epithelial markers (*LGALS4*, *CEACAM5*) [39], prostate antigens (*KLK3*, *NKX3-1*) [40], breast hormone receptors (*ESR1*, *GATA3*) [41], epidermal keratinization genes (*KRT6A*, *IVL*) [42], and melanocyte lineage factors (*TYR*, *SOX10*) [43] (Fig. 3b). In early pseudotime phases, epithelial adenocarcinomas similarly displayed a combination of shared and tissue-specific gene expression: most samples converged on myofibroblast contractility markers (*TAGLN*, *MYL9*) and ECM microfibril network genes (*COL6A2*, *EMILIN1*), whereas other transcripts exhibited organ-restricted patterns. For example, *DES* and *MYH11* were prominent in colorectal and prostate cancers; *GREM1*, *SLIT3*, and *SSC5D* were specific to colorectal tissue [44]; and *MAOB* and *SMTN* were prostate-enriched (Fig. 3c).

**Fig. 3.**
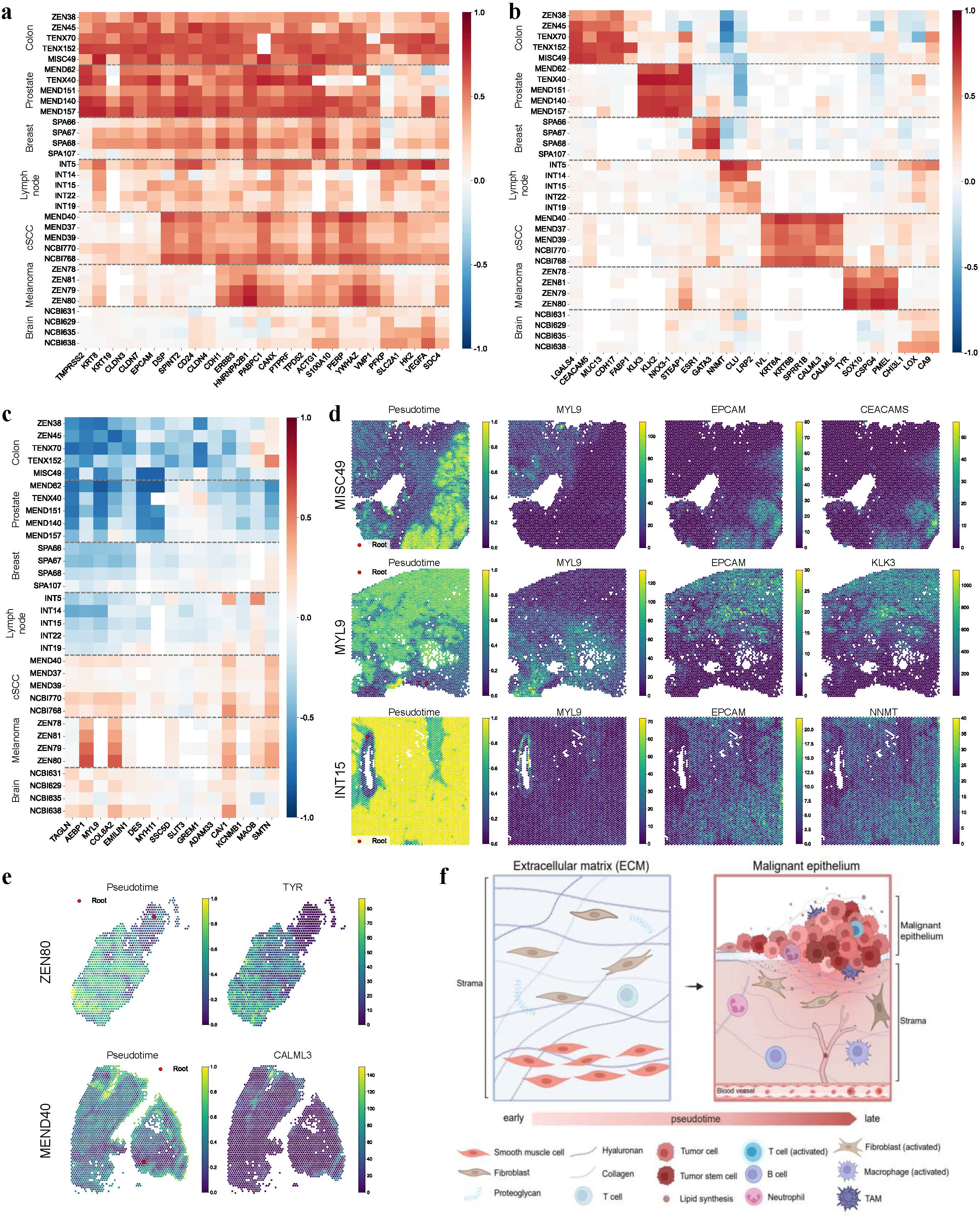
| Cross-tissue analysis of trajectory-enriched genes reveals an ‘early stromal, late malignant epithelial’ spatiotemporal pattern in epithelial adenocarcinomas. a. Integrated analysis of commonly or partially shared genes enriched in late trajectory stages across colorectal, prostate, breast, lymph node, skin (melanoma and cutaneous squamous cell carcinoma), and brain samples. b. Analysis of tissue-specific genes enriched in late trajectory stages across the above samples. c. Integrated analysis of commonly or tissue-specifically expressed genes enriched in early trajectory stages in colorectal, prostate, breast, and lymph node samples. d. Spatial in situ mapping of trajectory-enriched gene expression and pseudotime onto histopathological images of colorectal (MISC49), prostate (TENX40), and lymph node (INT15) tissues. e. Spatial in situ mapping of trajectory-enriched gene expression and pseudotime onto histopathological images of skin tissues. f. Schematic illustrating the biological interpretation of pseudotime progression from early to late in epithelial adenocarcinoma samples. Created with BioRender.com.

Spatial in situ mapping of gene expression and pseudotime onto histopathological images revealed that in epithelial adenocarcinomas (including samples from colon, prostate, and lymph node) early pseudotime values strongly colocalized with stromal gene expression, while late pseudotime values precisely overlapped with malignant epithelial gene signatures (Fig. 3d), visually validating the spatiotemporal trajectory of ‘early stromal, late malignant epithelial’ progression. In skin and brain tumors, late pseudotime also mapped accurately to malignant cell compartments (Fig. 3e). Together, these observations indicate that pseudotime captures a fundamental biological axis: in epithelial adenocarcinomas, progression from early to late pseudotime reflects a continuum from stromal dominance to malignant epithelial proliferation and invasion. Early phases are governed by genes that construct the ECM scaffold, maintain tissue mechanical tension, and support immune surveillance and physiological repair, which are defining features of stromal identity. Late phases, by contrast, switch to epithelial oncogenic programs driving rapid proliferation, metabolic reprogramming, stress survival, stemness, and metastatic dissemination. In summary, trajectory-based pseudotime inference represents a powerful analytical framework for dissecting spatial transcriptomic data, particularly in ordering the transition from stromal to malignant regions in epithelial adenocarcinoma samples (Fig. 3f).

### Histomorphology Model-Based Trajectory Inference and Correlation with ST Molecular Trajectories

Adopting the analytical framework established for ST, we next performed large-scale histomorphological trajectory inference using matched H&E images. For each ST spot, we employed two patch extraction strategies: fixed-resolution patches (224×224 pixels, 0.5 µm/pixel), which typically cover a larger area than individual spots, capture global tissue architecture, and align with standard histopathology model input requirements (large region); and spot-aligned patches, which precisely match the physical dimensions of ST spots, focusing on spot-localized cellular communities and microenvironmental features to enable direct one-to-one correspondence with gene expression profiles (small region). We systematically evaluated ten distinct models for extracting histomorphological feature vectors across both large and small regions: Virchow, Virchow2, UNI2-h, UNI, Prov-GigaPath, ResNet18 (RN18), Vision Transformer-Base/16 (ViT-B/16), CONCH, CONCHv1_5, and CTransPath. For each sample, 20 morphological pseudotime trajectories (10 for the large region and 10 for the small region) were constructed based on these model-extracted histomorphological feature vectors, and their concordance with corresponding spatial transcriptomics (ST)-derived molecular pseudotime was quantified (Fig. 4a, b; Figs. S6-S12).

**Fig. 4.**
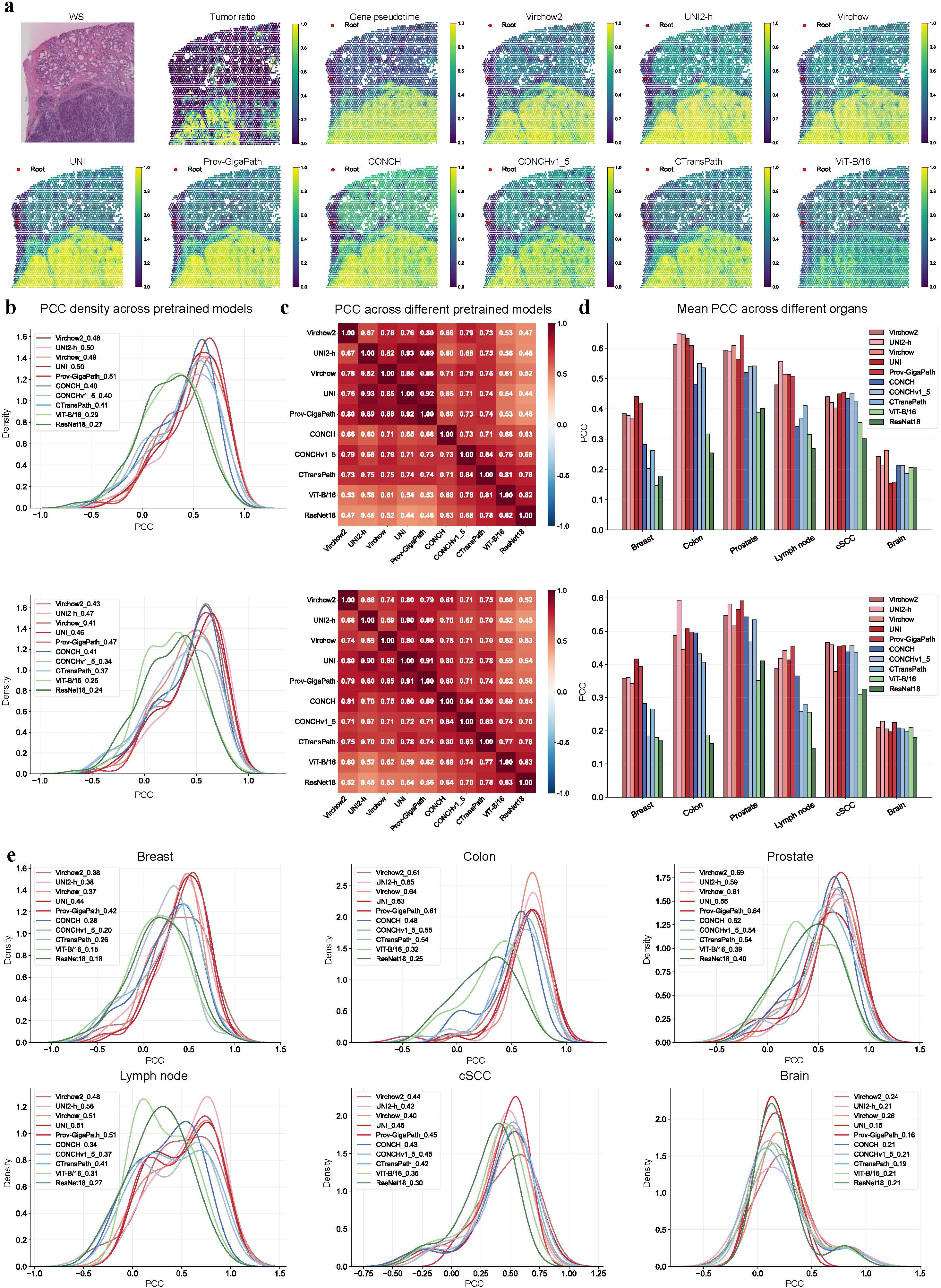
| Evaluation of correlations between morphological trajectories inferred by histopathology foundation models and spatial transcriptome-derived molecular trajectories. a. Original histopathological images and spatial mapping of cancer cell fraction, spatial transcriptome molecular pseudotime, and morphological trajectory pseudotime from nine histopathology foundation models on tissue sections (MEND62, large region). b. Schematic of feature extraction strategies (large and small regions) and distribution plots of correlations between morphological trajectories from various foundation models and spatial transcriptome molecular trajectories across multiple samples from six organs. c. Pairwise correlations among morphological trajectories generated by the ten histopathology foundation models (top: large region; bottom: small regions). d. Mean values of correlations between morphological trajectories from each model and spatial transcriptome molecular trajectories across organs (large and small regions). e. Distribution of correlations between morphological trajectories from each model and spatial transcriptome molecular trajectories across different organs (large region).

Correlation analysis (Pearson and Spearman correlation coefficients, Kendall’s tau coefficient, Wasserstein distance, cosine similarity) revealed a clear performance hierarchy (Fig. 4b, c; Figs. S13-S14; Tables. S1-S5). Under both patch extraction strategies, vision Transformer models pre-trained on large-scale histopathology data (Virchow, Virchow2, UNI2-h, UNI, Prov-GigaPath) exhibited the strongest correlations with ST molecular pseudotime; CONCH-series and CTransPath models performed moderately; and baseline models trained on natural images (ResNet18, ViT-B/16) showed the weakest concordance. Notably, the fixed-resolution (large-region) patch strategy outperformed spot-aligned patches (small-region) across almost all models (Fig. 4b, c; Figs. S13-S14; Tables. S1-S5). This stratification was primarily governed by three factors: domain specificity of training data, model architecture, and parameter scale. Specifically, models pre-trained on massive histopathology slides significantly outperformed those trained on general image corpora, and large-scale vision Transformer architectures (e.g., ViT-H/G14) exhibited vastly superior representation learning compared to smaller ResNet18 or ViT-B/16 models. We also observed pronounced tissue-specific variability: colon and prostate samples showed the highest morphology-molecule temporal concordance, while brain samples showed the lowest, likely reflecting inherent differences in morphological complexity, cellular heterogeneity, and spatial organization of gene expression across organ systems (Fig. 4d, e).

Spatial in situ mapping of trajectories for representative samples further validated these quantitative findings. In displayed samples, trajectories generated by high-correlation models (notably UNI2-h and Prov-GigaPath) exhibited near-identical pseudotime distributions to ST-derived trajectories; in contrast, trajectories from low-correlation models (ResNet18, ViT-B/16) lacked substantial alignment (Fig. 5a, b). To quantify this at the level of functional gene modules, we calculated mean correlations between gene expression trends (encompassing both early stromal and late malignant epithelial trajectory-enriched genes) and their respective pseudotime values across four epithelial adenocarcinomas. While specific gene sets varied across models, UNI2-h and Prov-GigaPath exhibited significantly stronger correlations between gene expression dynamics and pseudotime than ResNet18 and ViT-B/16, closely approximating ST benchmark performance (Fig. 5c; Figs. S15-S30). Gene set overlap analysis confirmed that top 100 key genes enriched by high-performance models showed far greater intersection with ST-enriched genes than low-performance models (Table S6). Scatterplot visualization further revealed a strong linear association: the similarity between model-generated and ST trajectories correlated positively with the monotonicity of enriched gene expression dynamics along pseudotime. In other words, morphological models that accurately capture stromal features early and malignant epithelial features late reconstruct pseudotime trajectories that closely reflect the biological progression revealed by ST (Fig. 5d).

**Fig. 5.**
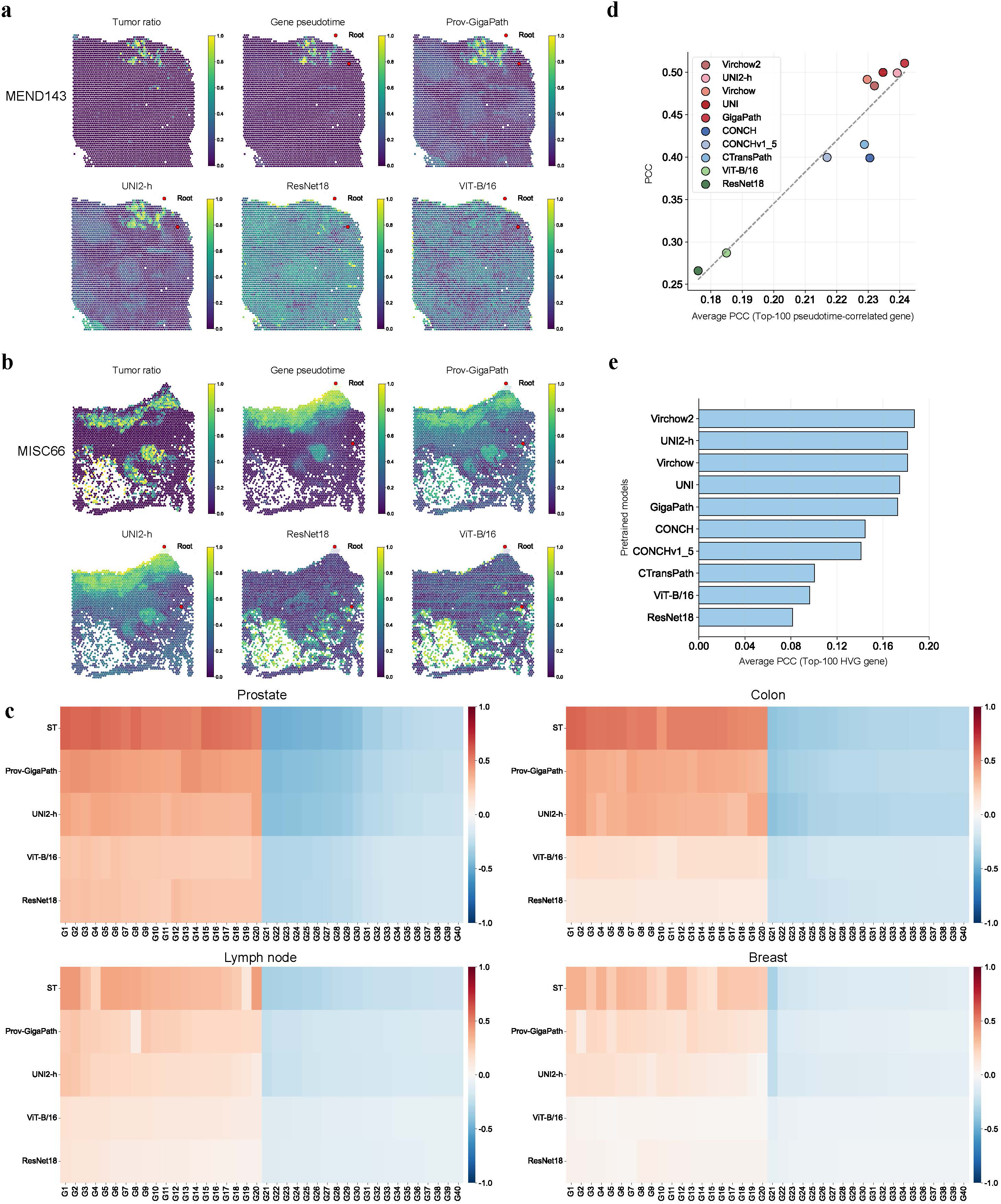
| Genomic dissection of the molecular mechanisms underlying similarity between model-derived morphological trajectories and spatial transcriptome molecular trajectories. a. Comparison of trajectories derived from morphological features (extracted by the top-performing and worst-performing models) with spatial transcriptome-based molecular trajectories in prostate cancer samples. (MEND143, large region). b. Comparison of trajectories derived from morphological features (extracted by the top-performing and worst-performing models) with spatial transcriptome-based molecular trajectories in colorectal cancer samples (MISC66, large region). c. Average correlations between pseudotime and expression trends of early- and late-stage enriched genes from trajectories generated by top- and worst-performing models in colorectal, prostate, breast, and lymph node samples. d. Similarity between model-derived trajectories and spatial transcriptome (ST) trajectories is significantly positively correlated with the monotonicity of dynamic gene expression changes along pseudotime. e. Results of the cross-modal translation task from morphological features to spatial transcriptomic profiles for each model.

Collectively, despite these pathological foundation models not being directly trained on ST data, our results demonstrate a striking convergence between extracted morphological features and gene expression profiles as training data evolves from general images to domain-specific histopathology slides, model architectures advance from conventional CNNs to large-scale vision Transformers, and parameter scales expand. This “morphology-molecule” correlation was further validated by our morphology-to-ST translation task: the model’s cross-modal translation capability is highly consistent with its trajectory inference performance. (Fig. 5e).

These findings establish trajectory inference as a viable metric for shared information between histomorphological and ST expression features. Morphological features extracted by different pathological models can be evaluated for shared information with transcriptomic profiles via dual trajectory analysis and inter-trajectory correlation.

### Analysis of Samples with Severe Modality-Specific Trajectory Divergence

While trajectories derived from most pathology and general visual models showed strong positive correlations with ST trajectories, a small subset of samples exhibited near-zero or even negative correlations across all models. Case inspection revealed striking divergence in sample INT4 (Fig. 6a). In INT4, the ST trajectory was characterized by early enrichment of plasma cell-specific diagnostic markers (*IGHG*, *IGKC*, *MZB1*, *JCHAIN*), genes driving robust antibody synthesis and secretion (*DERL3*), B-cell lineage maintenance (*PAX5, CD79A*), ECM remodeling and fibrosis (*COL10A1, COMP, LUM, DCN, CCN2, MGP*), and smooth muscle/myofibroblast contractility (*ACTG2, TAGLN*). In contrast, the late ST trajectory was enriched for tissue-specific markers (renal-associated *FXYD2, SPP1*), metastatic invasion (*FN1, LOX*), stemness, Warburg effect metabolism (*LDHA, SLC16A3, PKM*), and rapid proliferation (high expression of ribosomal RPL/RPS genes and mitochondrial oxidative phosphorylation genes to support energy and biosynthesis). Genes involved in chaperone and stress response (*SOD2, HSP90B1, CALR, P4HB*), chemoresistance (*BIRC3, DDIT4*), and MHC class I-mediated immune crosstalk were also upregulated. These ST-enriched genes recapitulate the previously defined trajectory from stromal toward malignant epithelial regions. (Fig. 6b, c).

**Fig. 6.**
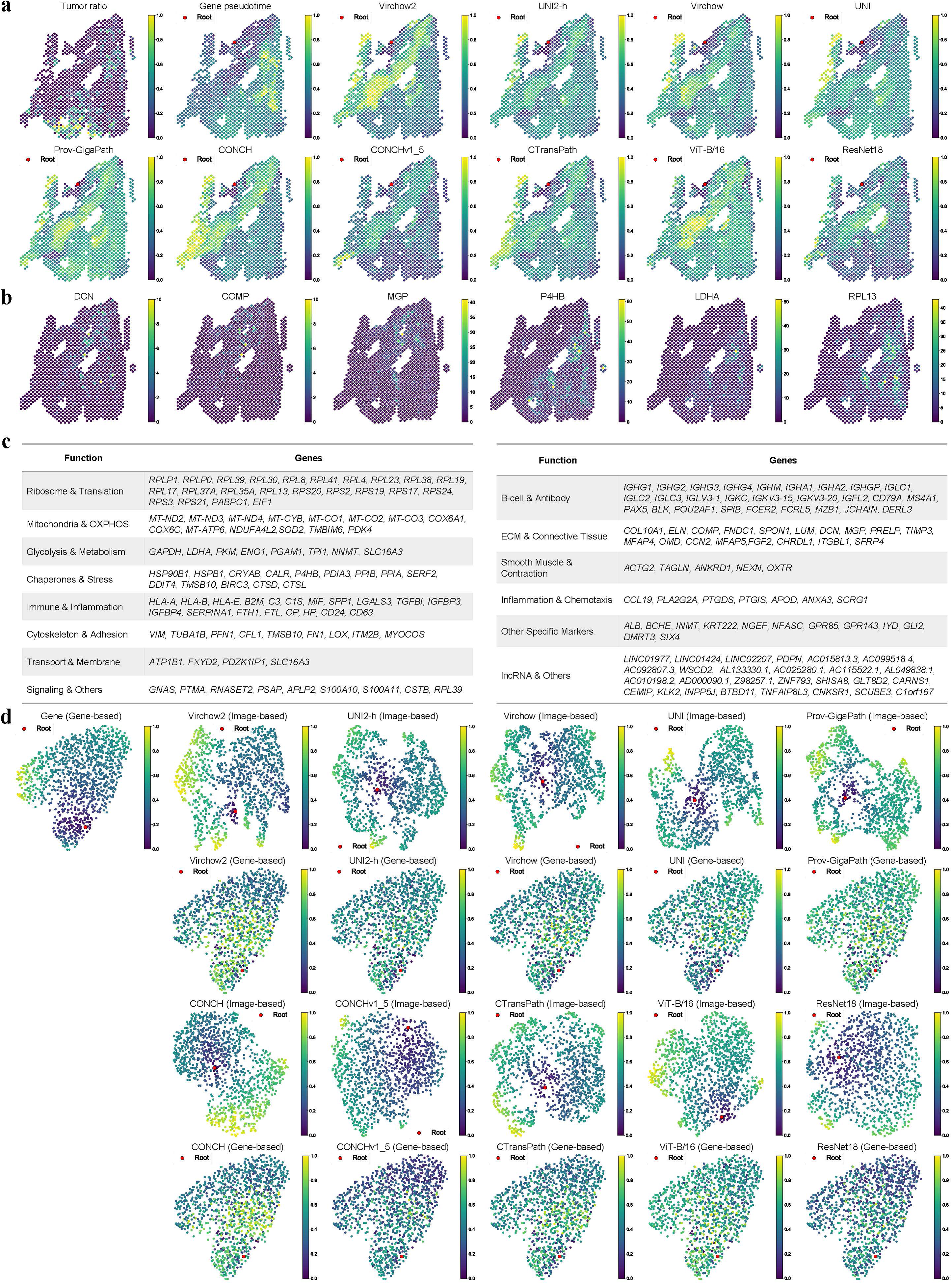
| In-depth characterization of samples with low morphology-molecule trajectory correlation. a. Spatial mapping of cancer cell fraction, spatial transcriptome molecular pseudotime, and morphological trajectories from ten pathology and general visual models onto histopathological images of sample INT4 (large region). b. Spatial expression patterns of stroma-related and malignant tumor-related genes in sample INT4. c. Gene sets and functional annotations enriched in early and late stages of the spatial transcriptome trajectory in sample INT4. d. UMAP visualization of dimensionality-reduced spatial transcriptome gene expression vectors and model-extracted histomorphological feature vectors, colored by molecular or morphological pseudotime.

Conversely, trajectories derived from most pathology and general visual models erroneously enriched genes related to ribosomal translation, metabolic reprogramming, and stress/chaperone responses (characteristics of malignant epithelial cells) in the early stage, while enriching ECM structure and remodeling genes (e.g., *COMP, DCN, MFAP5, COL14A1, COL11A1*) in the late stage., indicating failure to accurately capture transition from stromal to malignant regions (Table S7).

We further visualized these pronounced differences using UMAP embeddings of ST gene expression vectors and model-extracted histomorphological feature vectors, colored and rooted by corresponding pseudotime. UMAP structures derived from model-extracted histomorphological features diverged substantially from ST-based structures. Moreover, their origin points differed: ST-based trajectories initiated at the UMAP periphery, whereas most model-extracted histomorphological trajectories began in the central region. Coloring ST UMAP plots with model-generated pseudotime yielded patterns that were strikingly distinct from those obtained with ST pseudotime. Moreover, assignments based on these model-generated pseudotime values exhibited a disordered mixing of early and late points, in sharp contrast to the ordered progression observed with ST-derived pseudotime (Fig. 6d).

In contrast, sample MEND60 exhibited strong correlations between all model-extracted histomorphological pseudotime and ST-derived pseudotime. In this sample, UMAP structures from model-extracted histomorphological features closely resembled those from ST spot gene expression, with consistent origin positions and matching pseudotime coloring patterns, markedly contrasting observations in INT4 (Fig. S31).

Collectively, these findings demonstrate that the pronounced discrepancies between model-derived and ST trajectories in INT4 arise from extensive divergence between model-extracted histomorphological feature vectors and ST spot gene expression vectors.

## Discussion

In this study, we performed large-scale trajectory analysis across multiple cancer types, using spatial transcriptomic data and histopathological images processed with model-extracted histomorphological features.

Trajectory analysis of ST data revealed distinct gene enrichment patterns in tumor versus adjacent non-tumor regions. Notably, in malignant epithelial adenocarcinomas (from colon, prostate, breast, and lymph node), early pseudotime was enriched for genes involved in ECM scaffold formation (e.g., collagens), tissue mechanical tension (smooth muscle markers), immune surveillance, and physiological tissue repair. Late pseudotime, by contrast, was enriched for epithelial oncogenes driving rapid proliferation, metabolic reprogramming (glycolysis, lipid synthesis), stress survival, stemness, and invasive metastasis. This early-to-late pseudotime progression delineates a biological trajectory from a stromal-like state to a malignant epithelial phenotype.

Trajectory analysis of histopathological images using model-extracted histomorphological features revealed that, despite these models not being trained on ST data, correlations between model-derived histomorphological trajectories and ST trajectories progressively increased as training data evolved from general images to domain-specific histopathology slides, model architectures advanced from early CNNs to modern vision Transformers, and parameter scales expanded. This trend indicates that representations learned by foundation models are increasingly aligned with ST profiles. Analogous to ST-based gene enrichment, model-derived histomorphological trajectories also enriched stroma-related genes early and malignancy-associated genes late. Furthermore, the stronger the correlation with ST trajectories, the more similar the enriched gene sets. Specifically, the extent to which foundation model trajectories reflected the stromal-to-malignant epithelial transition correlated positively with their concordance with ST trajectories. These findings further confirm that, across analyzed samples, progressive advances in foundation models yield histomorphological features increasingly consistent with spatial transcriptomic profiles.

Subsequent experiments predicting ST profiles from histopathological images validated this trend: models with more advanced architectures and larger parameter scales extracted features that more accurately predicted ST gene expression patterns. However, despite this overall trend, significant discrepancies persisted in a small number of specific contexts.

Spatial domain identification and histopathological image segmentation, common downstream tasks in ST and histomorphological analysis, are inherently biased toward discrete classification, assigning clear categorical labels to spatial spots or image patches [10, 11, 19, 20]. Yet ST and histomorphological data inherently exhibit strong spatiotemporal continuity [45]. Guided by previous methodological frameworks [30–32, 46, 47], we reconstructed trajectories in multiple cancer samples using both ST expression profiles and H&E histomorphological features. Both modalities yielded consistent biological trajectories: stroma-related genes enriched at trajectory onset, followed by gradual transition toward malignancy-associated genes. This result clearly delineates the spatiotemporal organization of stromal and malignant epithelial compartments, strongly validating the utility of continuous analytical frameworks in spatial transcriptomics and histopathological imaging research. We conclude that trajectory reconstruction emphasizing continuity should be valued alongside discrete spatial domain identification (or histopathological segmentation) as complementary, indispensable analytical tools for ST and histopathological image research.

H&E-stained histopathological sections and spatial transcriptomics represent fundamentally distinct data modalities. H&E slides offer ease of preparation, low cost, and extensive archived datasets. By contrast, although ST remains relatively costly and data-limited, it provides precise molecular expression measurements that serve as powerful biomarkers for targeted clinical diagnosis and therapy, supporting refined downstream applications such as survival prediction and immunotherapy guidance. Consequently, numerous studies have aimed to predict ST gene expression from H&E-derived morphological features. However, systematic investigations into shared information between these two modalities, especially regarding how to design or select morphological features to maximize such shared information, remain scarce [25].

In this study, we systematically evaluated ten distinct morphological feature extraction approaches. For each method, starting from consistent root points, we computed trajectories derived from model-extracted morphological features and directly from ST expression profiles, then quantified inter-trajectory correlations. We propose that this correlation serves as a metric for shared information between morphological and ST modalities. This metric is supported by correlations in trajectory-enriched gene sets and validated by morphology-to-transcriptome translation performance.

Using this metric, we find that among current models, those with more modern architectures, larger parameter counts, and pre-training on H&E histopathology capture morphological features more consistent with transcriptomic profiles-despite no explicit training on transcriptomic data. Furthermore, our analyses indicate that epithelial adenocarcinoma samples are particularly amenable to trajectory-based analysis. We anticipate that the analytical framework and findings presented herein will provide valuable insights for integrating morphological and spatial transcriptomic data and offer guidance for addressing key challenges in this rapidly evolving field.

## Methods

### Spatial Transcriptomics and H&E Histopathology Data

This study utilized the HEST-1k dataset as the primary source [33]. HEST-1k is a large-scale dataset pairing spatial transcriptomics with histology images, comprising 1,229 spatial transcriptomics samples, each matched with corresponding high-resolution hematoxylin and eosin (H&E)-stained whole-slide images (WSIs) and rich metadata. The dataset integrates 153 public and in-house cohorts, covering 26 organs across two species-*Homo sapiens* and *Mus musculus*-and includes 367 tumor samples representing 25 cancer types, yielding approximately 2.1 million expression-morphology pairs and encompassing more than 76 million nuclear instances. We selected cancer-associated data from the colorectum, prostate, breast, lymph nodes, skin, and brain. Guided by spatial transcriptomic trajectory analysis, we further filtered samples to retain those with superior enrichment, resulting in a final cohort of 146 samples: 32 colorectal, 32 prostate, 41 breast, 16 lymph node, 15 skin, and 10 brain samples (Table S8).

### Morphological Feature Extraction

We implemented and systematically compared ten representative morphological feature extractors (vision models), spanning paradigms pre-trained on natural images, histopathology images, and multi-modal data, with a focus on contrasting their modalities, training data scale, learning strategies, architectural designs, and representational capabilities (Table 1). **Data modality:** Models fell into three categories: natural image (NI), pathology image (PI), and multi-modal pathology image-text (PI+T). Baseline models ResNet18 and ViT-B/16 were trained on natural images. CTransPath, UNI, and the Virchow series were specialized for pre-training on large-scale pathology images, whereas multi-modal approaches including CONCH and Prov-GigaPath incorporated textual supervision to enable cross-modal alignment.

**Table 1.**
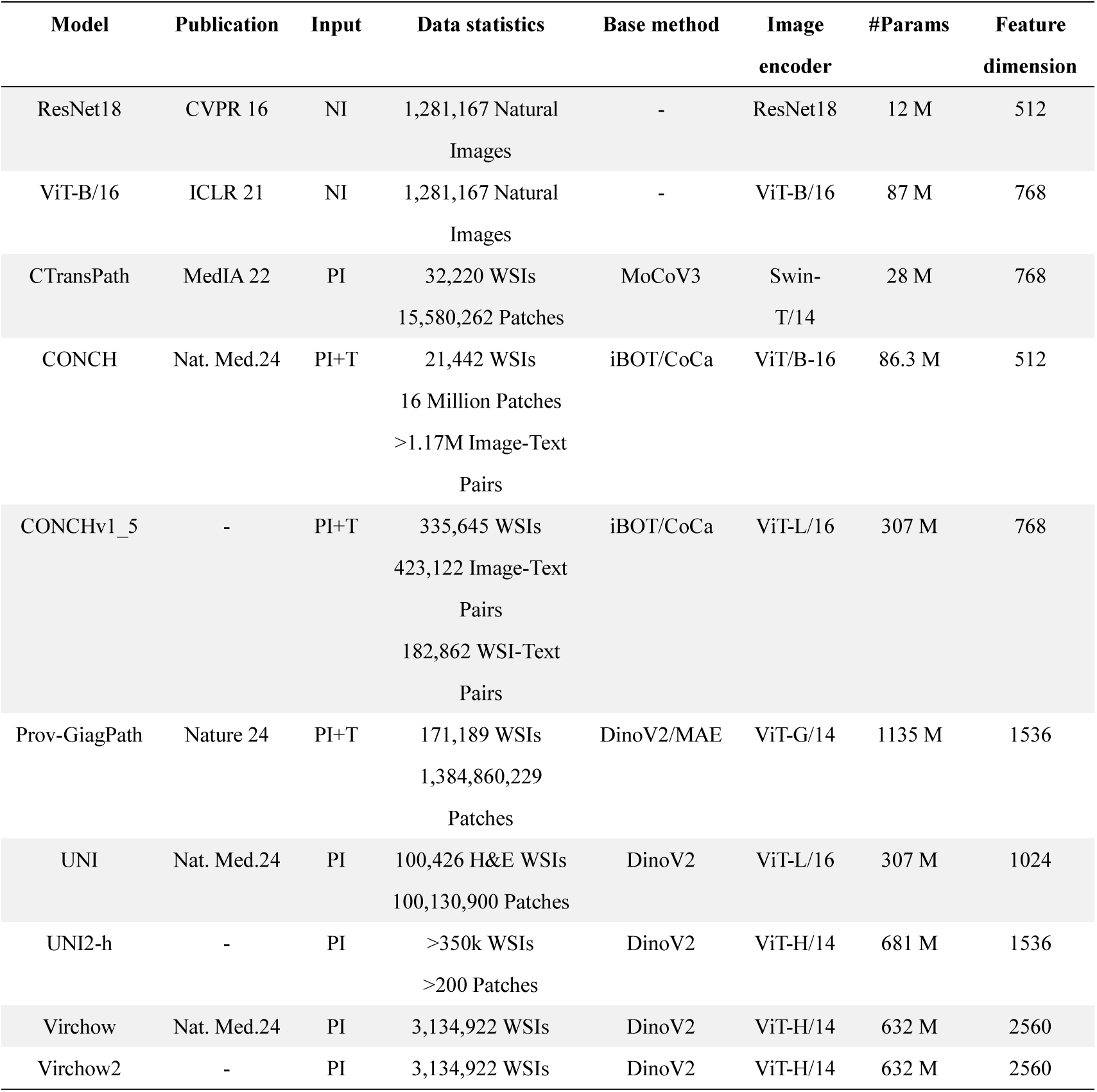
| Comparison of pretrained models. Abbreviations: NI (Natural Images), PI (Pathological Images), and T (Text).

#### Pre-training data scale

ResNet18 and ViT-B/16 relied primarily on natural image datasets (e.g., ImageNet, ∼1.28 million images). All other models were pathology-image-based. Early methods such as CTransPath were trained on ∼30,000 WSIs, while UNI scaled this to ∼100,000 WSIs. More recent approaches including Prov-GigaPath were trained on ∼170,000 WSIs, and the Virchow series was pre-trained on over 3 million WSIs, representing the current upper bound in data scale.

#### Pre-training strategy

Parameters of baseline models (ResNet18 and ViT-B/16) were learned via supervised learning. All other methods adopted self-supervised learning as the core pre-training paradigm.

#### Image encoder architecture

ResNet18 utilized conventional convolutional structures for feature extraction, whereas all other models were Transformer-based. Except for CTransPath, which employed a hierarchical Swin Transformer, all other models were built on the vision transformer (ViT) architecture. These models varied substantially in parameter count: from ViT-B to ViT-G, model size scaled superlinearly with increasing embedding dimension (width) and network depth, approximating an exponential expansion.

### Sample Pseudo-Time Calculation and Comparison of Two Modal Trajectories

#### In-Sample Spot Filtering

For each spot, we quantified the number of detected genes and retained only spots with at least 100 detected genes. For each gene, we computed its detection frequency across all spots and retained genes expressed in at least three spots. Spots with mitochondrial gene expression proportions exceeding 20% were excluded. We then applied tissue contour filtering: each spot was approximated as a circular region, and a spot was classified as intratumoral/intratissue if its centroid lay within the tissue boundary or if its overlapping area with the tissue region exceeded 70%. Finally, we performed cell-count filtering based on cellular geometric information: the overlapping area between each cell and neighboring spots was calculated, and each cell was uniquely assigned to the spot with which it shared the largest overlap to avoid double-counting. We then enumerated the number of cells per spot and discarded spots with zero cells, ensuring subsequent analyses were based on spatial units with valid cellular information.

#### Sample Filtering

At the sample level, only samples with ≥100 valid spots were retained. Additionally, only samples annotated as ‘Cancer’ were included according to disease status metadata, focusing analyses on tumor-associated spatial transcriptomic profiles. For all samples from the same organ, after calculating the top 100 or top 50 organ-enriched early- and late-trajectory genes, we filtered out samples whose correlation coefficients were inconsistent with most samples, and further retained those with superior enrichment under the guidance of spatial transcriptomic trajectory analysis.

#### Feature Extraction and Dimensionality Reduction

In the feature construction stage, both gene expression and imaging features were extracted for downstream analysis. For gene features, highly variable genes (HVGs) were selected from normalized, log-transformed, and standardized gene expression matrices to represent molecular profiles. For image features, pre-trained models were used to extract patch representations from tissue images corresponding to each spot via two strategies: (1) extracting fixed physical-size image patches (0.5 μm per pixel) centered on each spot to match the input dimensions of common pre-trained foundation models (PFMs), thereby leveraging representations learned from large-scale data; and (2) extracting image patches matching the exact spatial dimensions of the corresponding spot to ensure strict alignment between imaging information and spatial transcriptomic measurement regions, enabling more accurate characterization of the local tissue microenvironment. Image features were then matched to their respective spots via barcode information and standardized to ensure comparability across modalities.

#### Pseudo-Time Calculation

For each sample, principal component analysis (PCA) was first applied to reduce the dimensionality of gene expression features, and a k-nearest neighbor (KNN) adjacency graph was constructed. Diffusion maps were then used to refine adjacency relationships for better characterization of the data manifold. Leiden clustering was subsequently performed to assign cluster labels to all spots. Pseudo-time initiation was determined via a two-step procedure: first, the average tumor cell proportion of each cluster was computed using cellular composition data, and the cluster with the lowest tumor proportion was prioritized as the root cluster; in cases of multiple candidates, the cluster containing the largest number of spots was chosen. Second, a root spot was selected within the root cluster, prioritizing spots with a tumor cell proportion of 0 (or minimum); among multiple candidates, the spot with the smallest value along the first diffusion component of the diffusion map was designated the final root. Before pseudo-time computation, we verified that KNN graphs constructed from both gene and image features formed single connected components (connected component count = 1) to ensure graph integrity. Diffusion processes were then modeled using diffusion maps, and diffusion pseudotime (DPT) was employed to calculate the diffusion distance of each spot from the root node, yielding pseudo-time distributions derived separately from gene and image features. Results were validated to exclude NaN or infinite (INF) pseudo-time values; samples containing such invalid values were discarded.

#### Pseudo-Time Similarity Quantification

To assess consistency between pseudo-time estimates derived from gene and image features, multiple metrics were used for quantitative evaluation: Pearson and Spearman correlation coefficients to measure linear and monotonic associations, Kendall’s tau coefficient to evaluate rank consistency, Wasserstein distance to compare distributional differences, and cosine similarity to characterize overall trend similarity.

#### Trajectory-Enriched Gene Identification

For each sample, in-sample spot filtering was performed as described above. We then quantified gene expression across all tissue samples and retained genes expressed in more than half of the samples, with Ensembl IDs uniformly mapped to gene symbols. For each retained gene, Spearman correlation coefficients were computed between its expression vector and the pseudo-time vector for each sample. Trajectory-associated genes were enriched using two strategies: (1) adaptive thresholding based on sample count: starting from a correlation threshold of 0.5 and decreasing in steps of 0.01, we counted the number of samples in which each gene exhibited a correlation coefficient exceeding the current threshold, and selected the top 50 or top 100 positively and negatively correlated genes by sample count; (2) mean-rank-based selection: average correlation coefficients across all tissue samples were computed for each gene, and the top 50 or top 100 positively and negatively correlated genes were selected directly by mean value.

### Pathology-to-Spatial-Transcriptomics Prediction Paradigm

In this study, we established a task for predicting spatial transcriptomic expression from H&E histopathology images using the HEST-1k dataset. Experiments were conducted on a subset of 96 human tissue samples from the colon and prostate within HEST-1k, partitioned into training, validation, and test sets at a ratio of 70%:15%:15%. Gene expression values were normalized via log1p transformation, and the top 250 highly variable genes (HVGs) per sample were selected as prediction targets. The pathology-spatial-transcriptomics association model was implemented using the MISO framework, which models relationships among spatial positions (spots) across tissue sections via multiple instance learning (MIL) and incorporates Transformer architectures to capture local and cross-region tissue morphological features. Model performance was evaluated using the Pearson correlation coefficient (PCC). Training and evaluation were implemented using the PyTorch Lightning framework and performed on a single NVIDIA RTX 4090 GPU.

## Author contributions

K.X., L.W., and M.F. envisioned the project. L.W., K.X. and M.F. conceptualized the study and designed the experiments. X.H. and M.F. implemented the code. L.W., A.W., K.X., M.F. and X.H. performed the analysis and wrote the paper. X.H., X.Luo, X.Liu and T.S. provided assistance in writing and analysis.

## Funding

Not applicable.

## Availability of data and materials

The HEST-1k dataset are publicly available at https://huggingface.co/datasets/MahmoodLab/hest. HEST-1k comprises matched pathology slides and spatial gene expression pro files, enabling cross-modal analysis between tissue morphology and molecular signals. The pathology foundation models used in this study were obtained from publicly accessible repositories. Specifically, ResNet-18 and ViT-B/16 are available via the timm library (https://huggingface.co/timm). CTransPath can be accessed at https://github.com/Xiyue-Wang/TransPath. CONCH and CONCHv1_5 are available at https://huggingface.co/MahmoodLab/CONCH/tree/main and https://huggingface.co/MahmoodLab/conchv1_5, respectively. Prov-GigaPath is available at https://huggingface.co/prov-gigapath/prov-gigapath. UNI and UNI2-h are available at https://huggingface.co/MahmoodLab/UNI and https://huggingface.co/MahmoodLab/UNI2-h, respectively. Virchow and Virchow2 can be accessed at https://huggingface.co/paige-ai/Virchow and https://huggingface.co/paige-ai/Virchow2. All analysis code used in this study is publicly available at https://github.com/AndrewTal/CrossModalTrajCorr.

## Declarations

Not applicable.

## Ethics approval and consent to participate

Not applicable.

## Consent for publication

Not applicable.

## Competing interests

The authors declare that they have no competing interests. Each of the funding bodies took part in the design of the study and collection, analysis, and interpretation of data, and the writing of the manuscript.

## Notes

https://huggingface.co/datasets/MahmoodLab/hest

### Competing Interest Statement

The authors have declared no competing interest.

## Reference

1. Srinidhi, C.L., O. Ciga, and A.L. Martel, Deep neural network models for computational histopathology: A survey. Med Image Anal, 2021. 67: p. 101813.

2. van der Laak, J., G. Litjens, and F. Ciompi, Deep learning in histopathology: the path to the clinic. Nat Med, 2021. 27(5): p. 775–784.

3. Shmatko, A., et al., Artificial intelligence in histopathology: enhancing cancer research and clinical oncology. Nat Cancer, 2022. 3(9): p. 1026–1038.

4. Krishnan, R., P. Rajpurkar, and E.J. Topol, Self-supervised learning in medicine and healthcare. Nat Biomed Eng, 2022. 6(12): p. 1346–1352.

5. Vorontsov, E., et al., A foundation model for clinical-grade computational pathology and rare cancers detection. Nat Med, 2024. 30(10): p. 2924–2935.

6. Chen, R.J., et al., Towards a general-purpose foundation model for computational pathology. Nat Med, 2024. 30(3): p. 850–862.

7. Wang, X., et al., A pathology foundation model for cancer diagnosis and prognosis prediction. Nature, 2024. 634(8035): p. 970–978.

8. Lu, M.Y., et al., A visual-language foundation model for computational pathology. Nat Med, 2024. 30(3): p. 863–874.

9. Xu, H., et al., A whole-slide foundation model for digital pathology from realworld data. Nature, 2024. 630(8015): p. 181–188.

10. Cisternino, F., et al., Self-supervised learning for characterising histomorphological diversity and spatial RNA expression prediction across 23 human tissue types. Nat Commun, 2024. 15(1): p. 5906.

11. Claudio Quiros, A., et al., Mapping the landscape of histomorphological cancer phenotypes using self-supervised learning on unannotated pathology slides. Nat Commun, 2024. 15(1): p. 4596.

12. Marx, V., Method of the Year: spatially resolved transcriptomics. Nat Methods, 2021. 18(1): p. 9–14.

13. Palla, G., et al., Spatial components of molecular tissue biology. Nat Biotechnol, 2022. 40(3): p. 308–318.

14. 10x Visium Genomics Visium Spatial Gene Expression.

15. Fischer, D.S., A.C. Schaar, and F.J. Theis, Modeling intercellular communication in tissues using spatial graphs of cells. Nat Biotechnol, 2023. 41(3): p. 332–336.

16. Tejada-Lapuerta, A., et al., Nicheformer: a foundation model for single-cell and spatial omics. Nat Methods, 2025. 22(12): p. 2525–2538.

17. Stickels, R.R., et al., Highly sensitive spatial transcriptomics at near-cellular resolution with Slide-seqV2. Nat Biotechnol, 2021. 39(3): p. 313–319.

18. Liu, Y., et al., High-Spatial-Resolution Multi-Omics Sequencing via Deterministic Barcoding in Tissue. Cell, 2020. 183(6): p. 1665–1681 e18.

19. Yan, G., S.H. Hua, and J.J. Li, Categorization of 34 computational methods to detect spatially variable genes from spatially resolved transcriptomics data. Nat Commun, 2025. 16(1): p. 1141.

20. Kang, L., et al., Benchmarking computational methods for detecting spatial domains and domain-specific spatially variable genes from spatial transcriptomics data. Nucleic Acids Res, 2025. 53(7).

21. Fu, X., et al., Spatial gene expression at single-cell resolution from histology using deep learning with GHIST. Nat Methods, 2025. 22(9): p. 1900–1910.

22. Zhang, D., et al., Inferring super-resolution tissue architecture by integrating spatial transcriptomics with histology. Nat Biotechnol, 2024. 42(9): p. 1372–1377.

23. Zeng, Y., et al., Spatial transcriptomics prediction from histology jointly through Transformer and graph neural networks. Brief Bioinform, 2022. 23(5).

24. He, B., et al., Integrating spatial gene expression and breast tumour morphology via deep learning. Nat Biomed Eng, 2020. 4(8): p. 827–834.

25. Chelebian, E., C. Avenel, and C. Wahlby, Combining spatial transcriptomics with tissue morphology. Nat Commun, 2025. 16(1): p. 4452.

26. Wolf, F.A., et al., PAGA: graph abstraction reconciles clustering with trajectory inference through a topology preserving map of single cells. Genome Biol, 2019. 20(1): p. 59.

27. Qiu, X., et al., Reversed graph embedding resolves complex single-cell trajectories. Nat Methods, 2017. 14(10): p. 979–982.

28. Street, K., et al., Slingshot: cell lineage and pseudotime inference for single-cell transcriptomics. BMC Genomics, 2018. 19(1): p. 477.

29. Wang, M., et al., Unraveling temporal and spatial biomarkers of epithelialmesenchymal transition in colorectal cancer: insights into the crucial role of immunosuppressive cells. J Transl Med, 2023. 21(1): p. 794.

30. Pham, D., et al., Robust mapping of spatiotemporal trajectories and cell-cell interactions in healthy and diseased tissues. Nat Commun, 2023. 14(1): p. 7739.

31. Smelik, M., et al., Combining Spatial Transcriptomics, Pseudotime, and Machine Learning Enables Discovery of Biomarkers for Prostate Cancer. Cancer Res, 2025. 85(13): p. 2514–2526.

32. Liu, Y., et al., Image-based inference of tumor cell trajectories enables large-scale cancer progression analysis. Sci Adv, 2025. 11(29): p. eadv9466.

33. Jaume, G., Doucet, P., Song, A. H., Lu, M. Y., Almagro-Pérez, C., Wagner, S. J., Vaidya, A. J., Chen, R. J., Williamson, D. F. K., Kim, A., & Mahmood, F., HEST-1k: A Dataset for Spatial Transcriptomics and Histology Image Analysis. Advances in Neural Information Processing Systems, 2024. 37: p. 53798–53833.

34. He, K., Zhang, X., Ren, S., & Sun, J., Deep Residual Learning for Image Recognition. 2015.

35. Dosovitskiy, A., Beyer, L., Kolesnikov, A., Weissenborn, D., Zhai, X., Unterthiner, T., Dehghani, M., Minderer, M., Heigold, G., Gelly, S., Uszkoreit, J., & Houlsby, N., An Image is Worth 16x16 Words: Transformers for Image Recognition at Scale. 2020.

36. Zimmermann, E., Vorontsov, E., Viret, J., Casson, A., Zelechowski, M., Shaikovski, G., Tenenholtz, N., Hall, J., Klimstra, D., Yousfi, R., Fuchs, T., Fusi, N., Liu, S., & Severson, K., Virchow2: Scaling Self-Supervised Mixed Magnification Models in Pathology. 2024.

37. Wang, X., et al., Transformer-based unsupervised contrastive learning for histopathological image classification. Med Image Anal, 2022. 81: p. 102559.

38. Yu, Z., et al., Integrative Single-Cell Analysis Reveals Transcriptional and Epigenetic Regulatory Features of Clear Cell Renal Cell Carcinoma. Cancer Res, 2023. 83(5): p. 700–719.

39. Liscu, H.D., et al., Biomarkers in Colorectal Cancer: Actual and Future Perspectives. Int J Mol Sci, 2024. 25(21).

40. Cancer Genome Atlas Research, N., The Molecular Taxonomy of Primary Prostate Cancer. Cell, 2015. 163(4): p. 1011–25.

41. Martin, E.M., et al., The estrogen receptor/GATA3/FOXA1 transcriptional network: lessons learned from breast cancer. Curr Opin Struct Biol, 2021. 71: p. 65–70.

42. Pandey, S., et al., Combined loss of expression of involucrin and cytokeratin 13 is associated with poor prognosis in squamous cell carcinoma of mobile tongue. Head Neck, 2021. 43(11): p. 3374–3385.

43. Wang, L., et al., The SOX10-ACAT2-Cholesterol Synthesis Axis Is Required for Melanoma Proliferation. Int J Biol Sci, 2026. 22(4): p. 1717–1732.

44. Huang, T., et al., The emerging role of Slit-Robo pathway in gastric and other gastro intestinal cancers. BMC Cancer, 2015. 15: p. 950.

45. Smith, E.A. and H.C. Hodges, The Spatial and Genomic Hierarchy of Tumor Ecosystems Revealed by Single-Cell Technologies. Trends Cancer, 2019. 5(7): p. 411–425.

46. Chitra, U., et al., Mapping the topography of spatial gene expression with interpretable deep learning. Nat Methods, 2025. 22(2): p. 298–309.

47. Liang, Q., et al., LSGI: interpretable spatial gradient analysis for spatial transcriptomics data. Genome Biol, 2025. 26(1): p. 238.

