## Supplemental Figure S1-S31,Supplemental Table S1-S8 for "Quantifying Cross-Modal Shared Information Between Histomorphology and Spatial Transcriptomics via Spatiotemporal Trajectory Correlation"

Supplementary Figures Legend

Fig. S1 Top 100 genes most strongly positively or negatively correlated with pseudotime, and their functional annotations, enriched along spatial transcriptome-derived trajectories in colorectal samples.

Fig. S2 Top 100 genes most strongly positively or negatively correlated with pseudotime, and their functional annotations, enriched along spatial transcriptome-derived trajectories in prostate samples.

Fig. S3 Top 100 genes most strongly positively or negatively correlated with pseudotime, and their functional annotations, enriched along spatial transcriptome-derived trajectories in breast samples.

Fig. S4 Top 100 genes most strongly positively or negatively correlated with pseudotime, and their functional annotations, enriched along spatial transcriptome-derived trajectories in lymph node samples.

Fig. S5 Top 100 genes most strongly positively correlated with pseudotime, and their functional annotations, enriched along spatial transcriptome-derived trajectories in brain and skin (melanoma, squamous cell carcinoma) samples.

Fig. S6 Original histopathological images and spatial mapping of cancer cell fraction, spatial transcriptome molecular pseudotime, and morphological trajectory pseudotime from nine histopathology foundation models on tissue sections (MEND62, small region).

Fig. S7 Original histopathological images and spatial mapping of cancer cell fraction, spatial transcriptome molecular pseudotime, and morphological trajectory pseudotime from ten models on samples INT10 and MEND160 (large region).

Fig. S8 Original histopathological images and spatial mapping of cancer cell fraction, spatial transcriptome molecular pseudotime, and morphological trajectory pseudotime from ten models on samples MISC73 and INT2 (large region).

Fig. S9 Original histopathological images and spatial mapping of cancer cell fraction, spatial transcriptome molecular pseudotime, and morphological trajectory pseudotime from ten models on samples MEND153 and NCBI637 (large region).

Fig. S10 Original histopathological images and spatial mapping of cancer cell fraction, spatial transcriptome molecular pseudotime, and morphological trajectory pseudotime from ten models on samples INT10 and MEND160 (small region).

Fig. S11 Original histopathological images and spatial mapping of cancer cell fraction, spatial transcriptome molecular pseudotime, and morphological trajectory pseudotime from ten models on samples MISC73 and INT2 (small region).

Fig. S12 Original histopathological images and spatial mapping of cancer cell fraction, spatial transcriptome molecular pseudotime, and morphological trajectory pseudotime from ten models on samples MEND153 and NCBI637 (small region).

Fig. S13 Distributions of correlation coefficients (left), mean and median values (middle), and pairwise correlations (right) between morphological trajectories (large-region features) from each histopathology foundation model and spatial transcriptome molecular trajectories across multiple samples from six organs.

Fig. S14 Distributions of correlation coefficients (left), mean and median values (middle), and pairwise correlations (right) between morphological trajectories (small-region features) from each histopathology foundation model and spatial transcriptome molecular trajectories across multiple samples from six organs.

Fig. S15 Top 100 genes most strongly positively or negatively correlated with pseudotime enriched along morphological trajectories derived from large-region Prov-GigaPath features in colorectal samples.

Fig. S16 Top 100 genes most strongly positively or negatively correlated with pseudotime enriched along morphological trajectories derived from large-region Prov-GigaPath features in prostate samples.

Fig. S17 Top 100 genes most strongly positively or negatively correlated with pseudotime enriched along morphological trajectories derived from large-region Prov-GigaPath features in breast samples.

Fig. S18 Top 100 genes most strongly positively or negatively correlated with pseudotime enriched along morphological trajectories derived from large-region Prov-GigaPath features in lymph node (metastatic renal cell carcinoma) samples.

Fig. S19 Top 100 genes most strongly positively or negatively correlated with pseudotime enriched along morphological trajectories derived from large-region UNI2-h features in colorectal samples.

Fig. S20 Top 100 genes most strongly positively or negatively correlated with pseudotime enriched along morphological trajectories derived from large-region UNI2-h features in prostate samples.

Fig. S21 Top 100 genes most strongly positively or negatively correlated with pseudotime enriched along morphological trajectories derived from large-region UNI2-h features in breast samples.

Fig. S22 Top 100 genes most strongly positively or negatively correlated with pseudotime enriched along morphological trajectories derived from large-region UNI2-h features in lymph node (metastatic renal cell carcinoma) samples.

Fig. S23 Top 100 genes most strongly positively or negatively correlated with pseudotime enriched along morphological trajectories derived from large-region ResNet18 features in colorectal samples.

Fig. S24 Top 100 genes most strongly positively or negatively correlated with pseudotime enriched along morphological trajectories derived from large-region ResNet18 features in prostate samples.

Fig. S25 Top 100 genes most strongly positively or negatively correlated with pseudotime enriched along morphological trajectories derived from large-region ResNet18 features in breast samples.

Fig. S26 Top 100 genes most strongly positively or negatively correlated with pseudotime enriched along morphological trajectories derived from large-region ResNet18 features in lymph node (metastatic renal cell carcinoma) samples.

Fig. S27 Top 100 genes most strongly positively or negatively correlated with pseudotime enriched along morphological trajectories derived from large-region ViT-B/16 features in colorectal samples.

Fig. S28 Top 100 genes most strongly positively or negatively correlated with pseudotime enriched along morphological trajectories derived from large-region ViT-B/16 features in prostate samples.

Fig. S29 Top 100 genes most strongly positively or negatively correlated with pseudotime enriched along morphological trajectories derived from large-region ViT-B/16 features in breast samples.

Fig. S30 Top 100 genes most strongly positively or negatively correlated with pseudotime enriched along morphological trajectories derived from large-region ViT-B/16 features in lymph node (metastatic renal cell carcinoma) samples.

Fig. S31 UMAP visualization of dimensionality-reduced spatial transcriptome gene expression vectors and histopathological image feature vectors extracted by each foundation model for sample MEND62, colored by molecular or morphological pseudotime.

Fig.S1

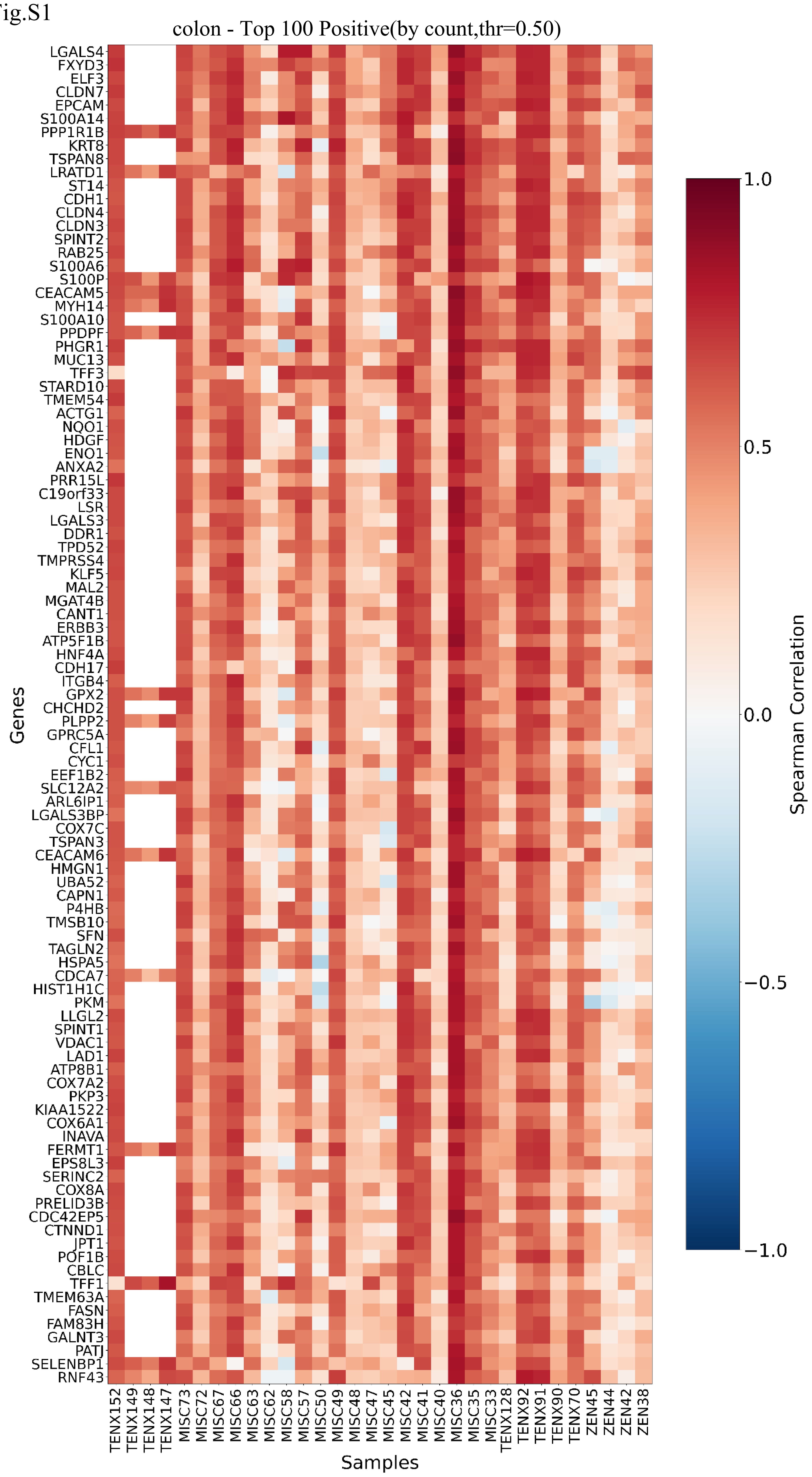


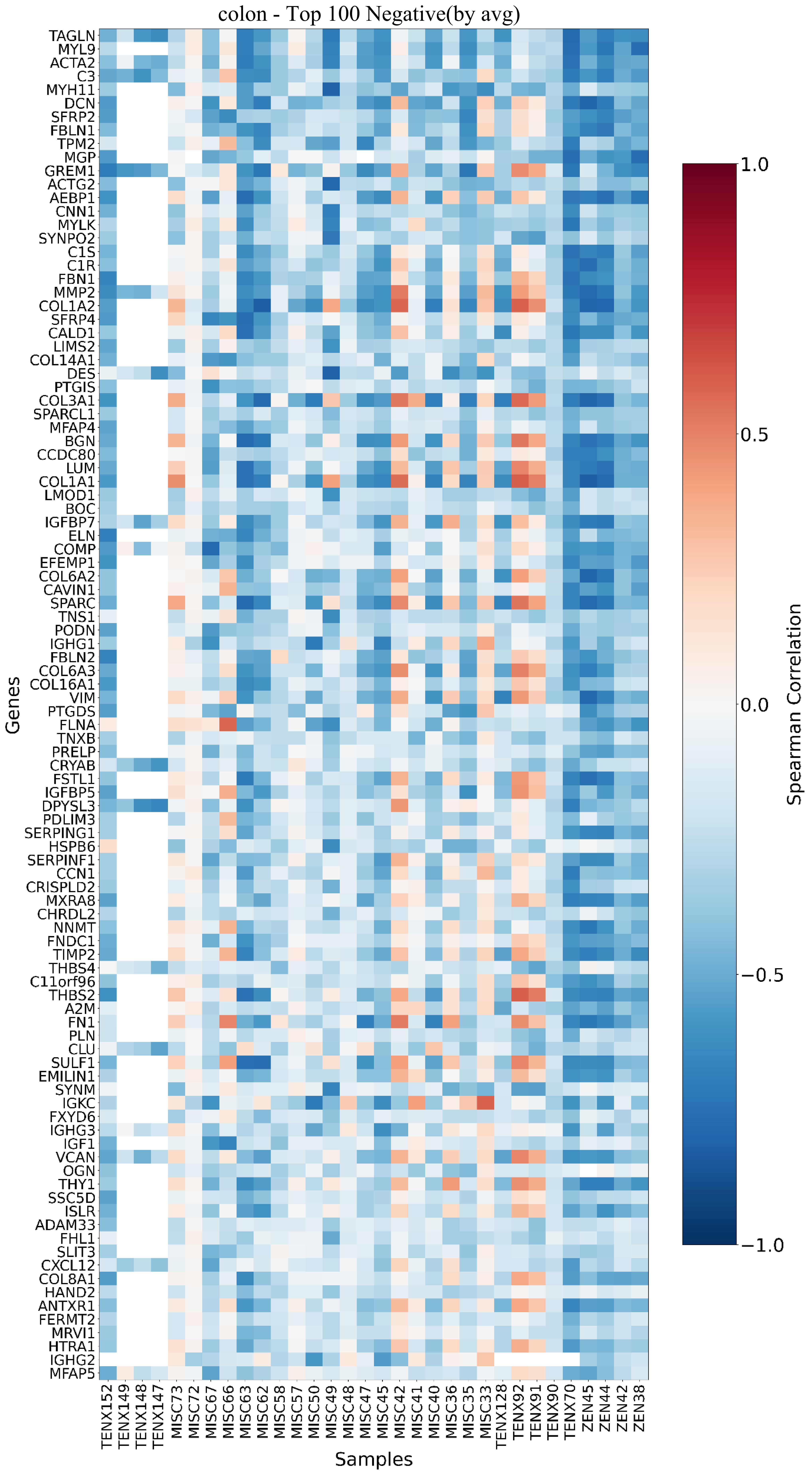


| Function | Genes |
|---|---|
| Core Diagnostic & Serological Markers | *CEACAM5 (CEA), CEACAM6, LGALS4, CLDN3, CLDN4, CLDN7* |
| Metastasis & Invasion Drivers | *EPCAM, KRT8, TSPAN8, ANXA2, ST14, DDR1, S100A14, S100A10 (P), S100A6* |
| Stemness & Signaling | *SOX9, KLF5, RNF43, ERBB3, TFF1, TFF3, HNF4A* |
| Warburg Effect & Metabolism | *PKM, ENO1, FXYD3, ATP5F1B, ATP5IF1, COX7C, COX8A, ASS1* |
| Antioxidant & Chemoresistance | *GPX2, NQO1, PRDX2, HSPA5, SFN* |

| Function | Genes |
|---|---|
| ECM Framework Construction | *COL1A1, COL1A2, COL3A1, COL6A2, COL6A3, FN1, ELN, LUM* |
| Matrix Homeostasis & Remodeling | *MMP2, TIMP2, THBS2, THBS4, SPARC* |
| Growth Regulation & Repair | *IGF1, IGFBP5, IGFBP7, GREM1, SFRP2, SFRP4, CXCL12* |
| Myofibroblast Tension & Barrier | *ACTA2, VIM, TAGLN, MYH11, DES* |

Fig.S2

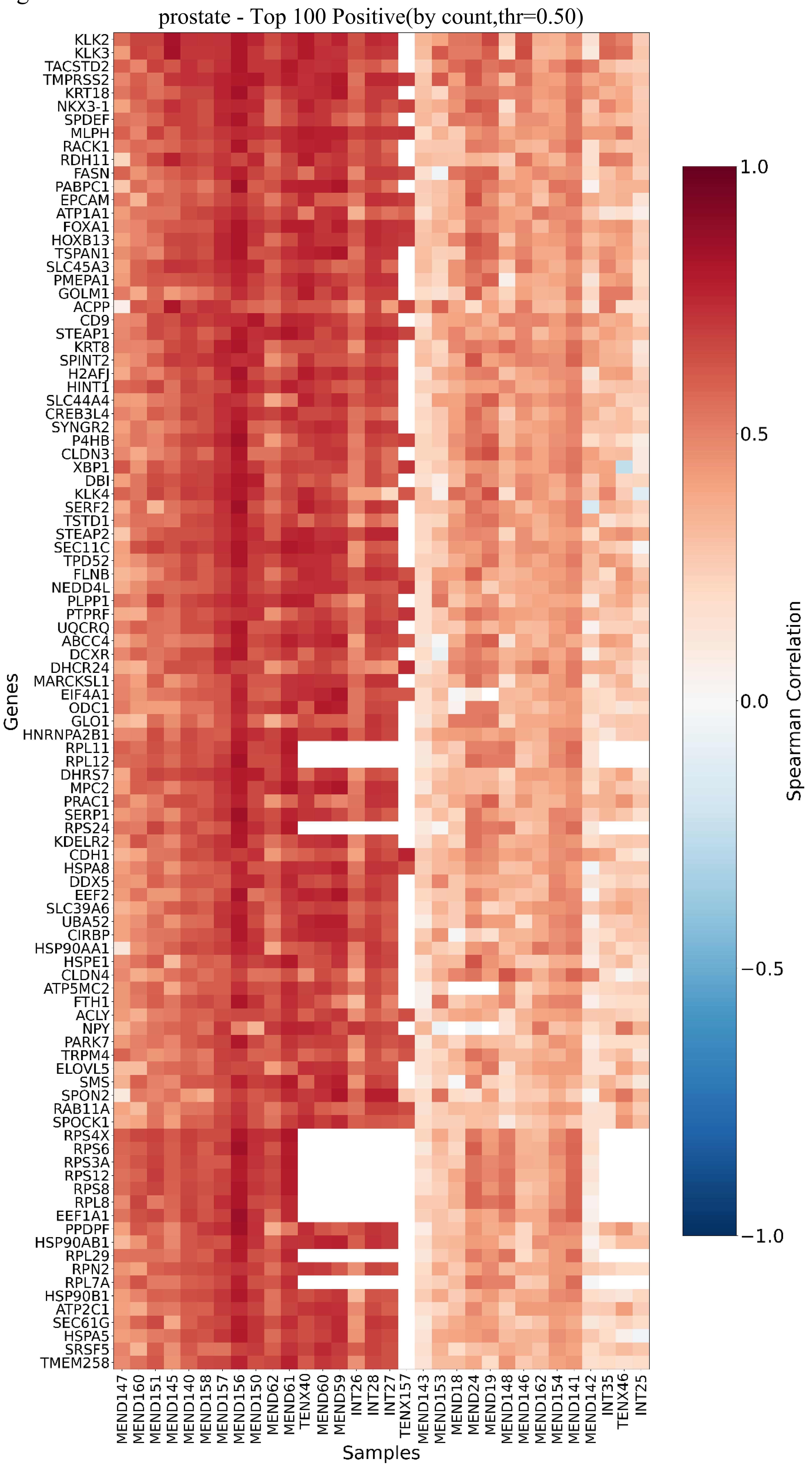


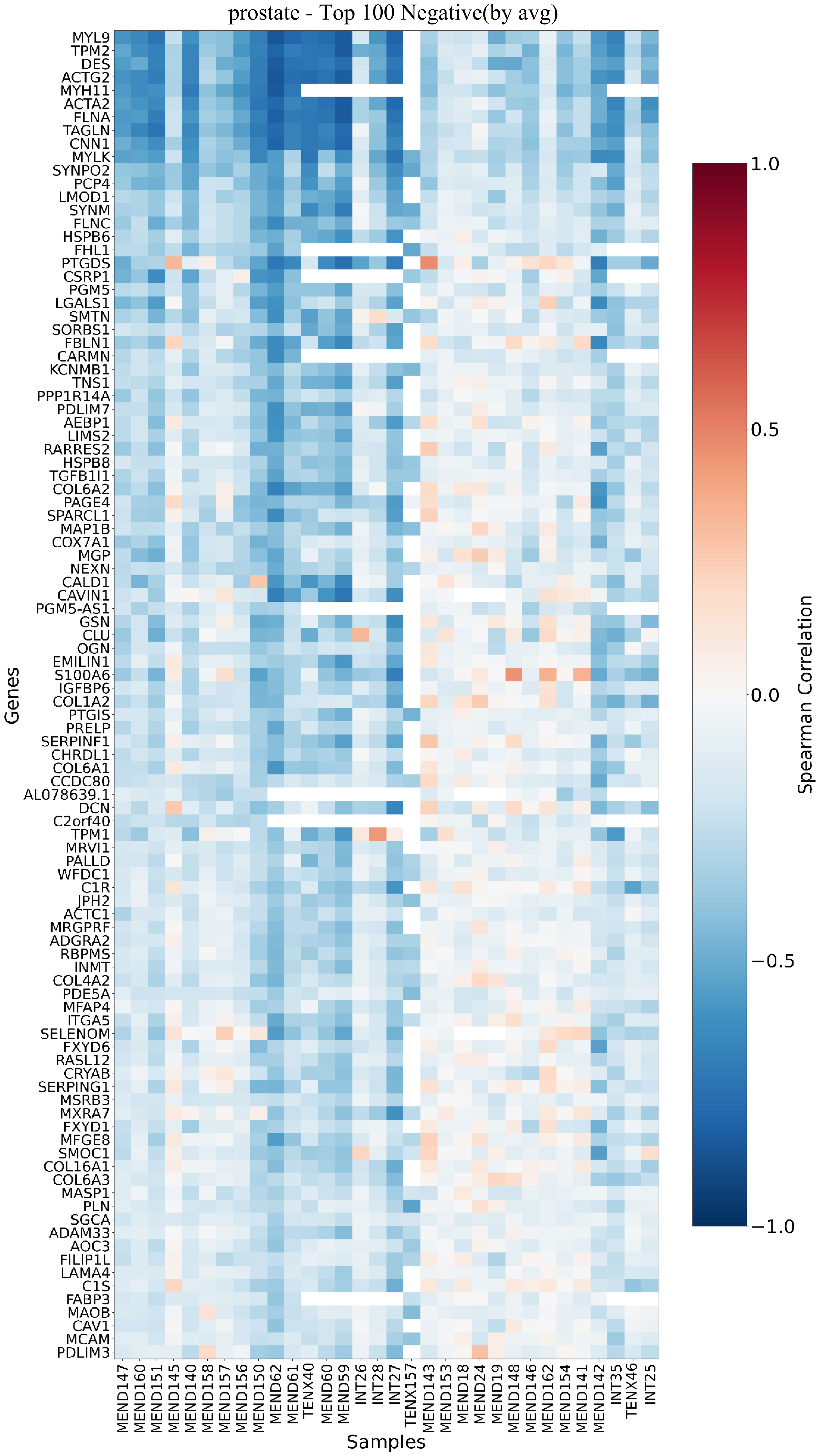


| Function | Genes |
|---|---|
| Classic Markers | *KLK3 (PSA), KLK2, KLK4, ACPP (PAP), TMPRSS2* |
| Epithelial Integrity & Metastasis | *EPCAM, TACSTD2, KRT8, KRT18, CLDN3, CLDN4, TSPAN1, STEAP1, STEAP2* |
| Lineage & AR Signaling | *NKX3-1, HOXB13, FOXA1, SPDEF* |
| Lipid Metabolism Reprogramming | *FASN, ACLY, DHCR24, ODC1, ATP1A1* |
| Stress Response & Resistance | *HSPA8, HSP90AA1, PARK7, XBP1, GLO1, P4HB* |

| Function | Genes |
|---|---|
| Stromal & Smooth Muscle Markers | *ACTA2 (α-SMA), MYH11, DES (Desmin), TPM1/2, CALD1* |
| ECM Architecture & Support | *COL1A2, COL6A1/2/3, LAMA4, DCN (Decorin), SPARC* |
| Contractility & Mechanotransduction | *MYLK, FLNA/B, TAGLN (SM22α), CNN1, LIMS2, PDLIM3/7* |
| Growth Regulation & Homeostasis | *TGFB1I1, IGFBP6, LGALS1, CLU (Clusterin), S100A6, ADAM33* |
| Protection & Metabolic Balance | *CRYAB, HSPB6/8, KCNMB1, FXYD1/6, CAV1* |

Fig.S3

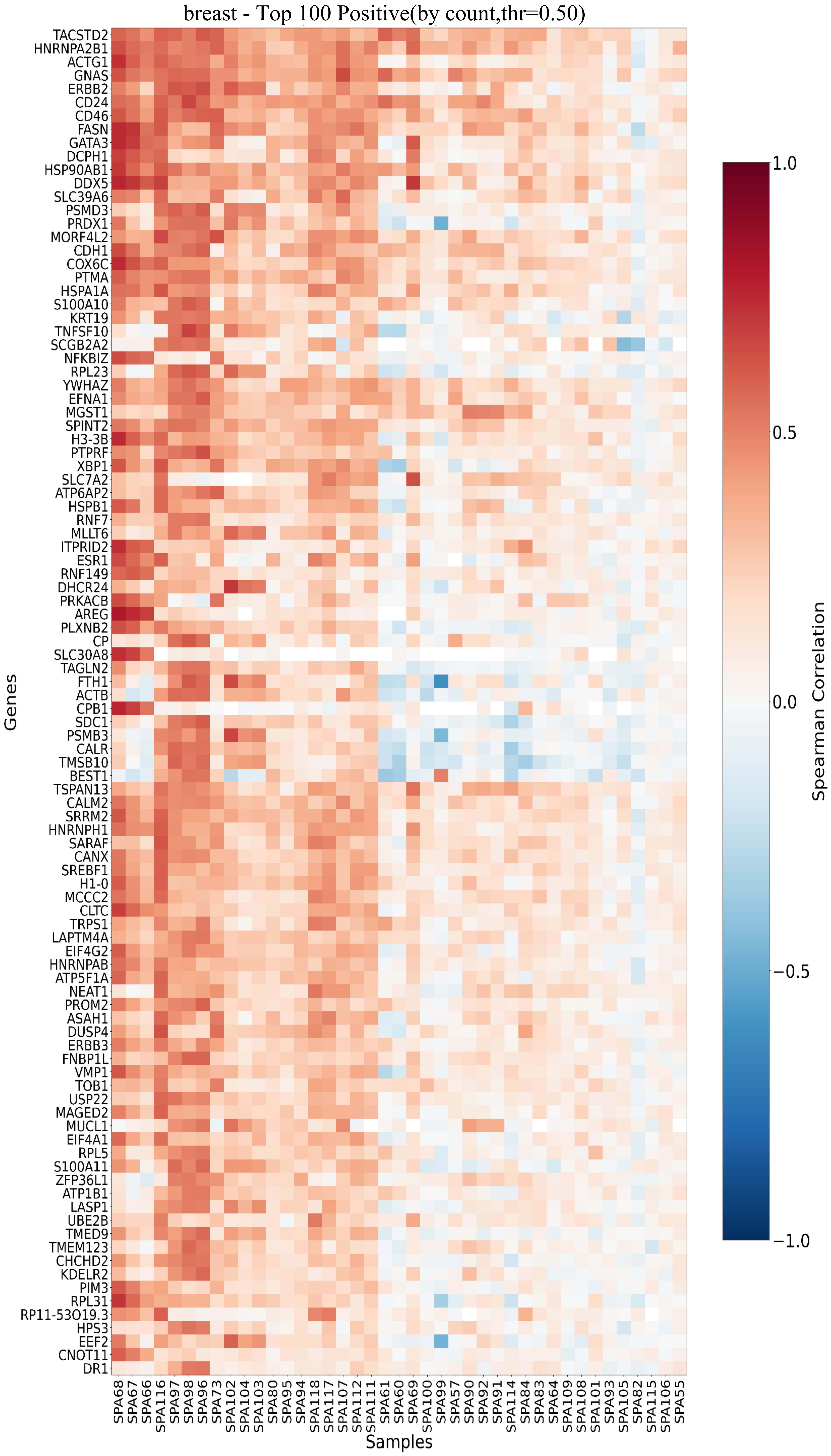


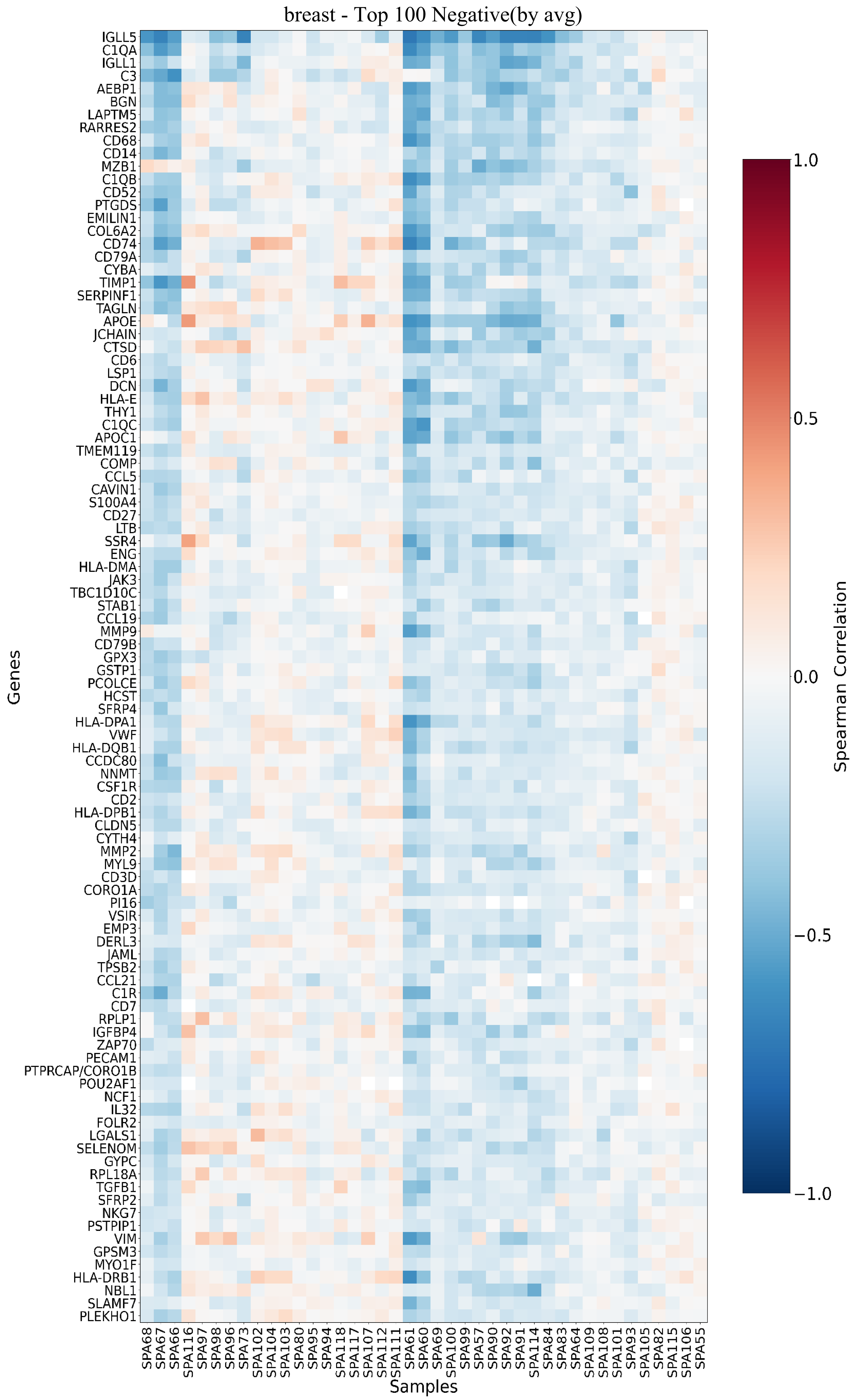


| Function | Genes |
|---|---|
| Core Diagnosis & Hormone Dependence | *ESR1, GATA3, TRPS1, KRT19, SCGB2A2* |
| Proliferation & Survival Drivers | *ERBB2 (HER2), ERBB3, TACSTD2 (TROP2), CD24, FASN* |
| Differentiation & Transcription | *GATA3, XBP1, HNRNPA2B1/H1, DDX5, ZFP36L1* |
| Protein Synthesis & Metabolic Reprogramming | *RPL23, RPL5, RPL31, RPLP0,IF4A1, EIF4G2 EEF2, SREBF1, ATP5F1A, ATP6AP2, ATP1B1, SLC39A6, SLC7A2, SLC30A8* |
| Stress Resistance & Therapeutic Resistance | *HSP90AB1, HSPA1A, HSPB1, CALR, PRDX1, FTH1* |

| Function | Genes |
|---|---|
| Resident Immune Surveillance & Antigen Presentation | *CD74, HLA-DRB1/DPB1/DQB1/DPA1 CD68, CD14, C1QA/B/C* |
| Normal Microvascular Network | *PECAM1 (CD31), VWF, ENG (CD105), CLDN5* |
| Tissue-Specific Immune Defense & Inflammation Regulation | *CCL19, CCL21, CCL5, IL32, JAK3, ZAP70, CD3D, CD2, NKG7* |
| Physiological Stromal Maintenance & Remodeling | *VIM, TAGLN, MYL9, DCN, BGN, COL6A2, EMILIN1, COMP, SFRP2/4, TGFB1, MMP2/9, TIMP1, AEBP1* |
| Lipid Metabolism, Antioxidant Defense & Homeostasis | *APOE, APOC1, GPX3, GSTP1, SELENOM, NNMT, FOLR2* |
| B-cell/Plasma Cell Lineage & Complement Clearance | *CD79A/B, MZB1, JCHAIN, POU2AF1, C3, C1R, VSIR, LTB* |

Fig.S4

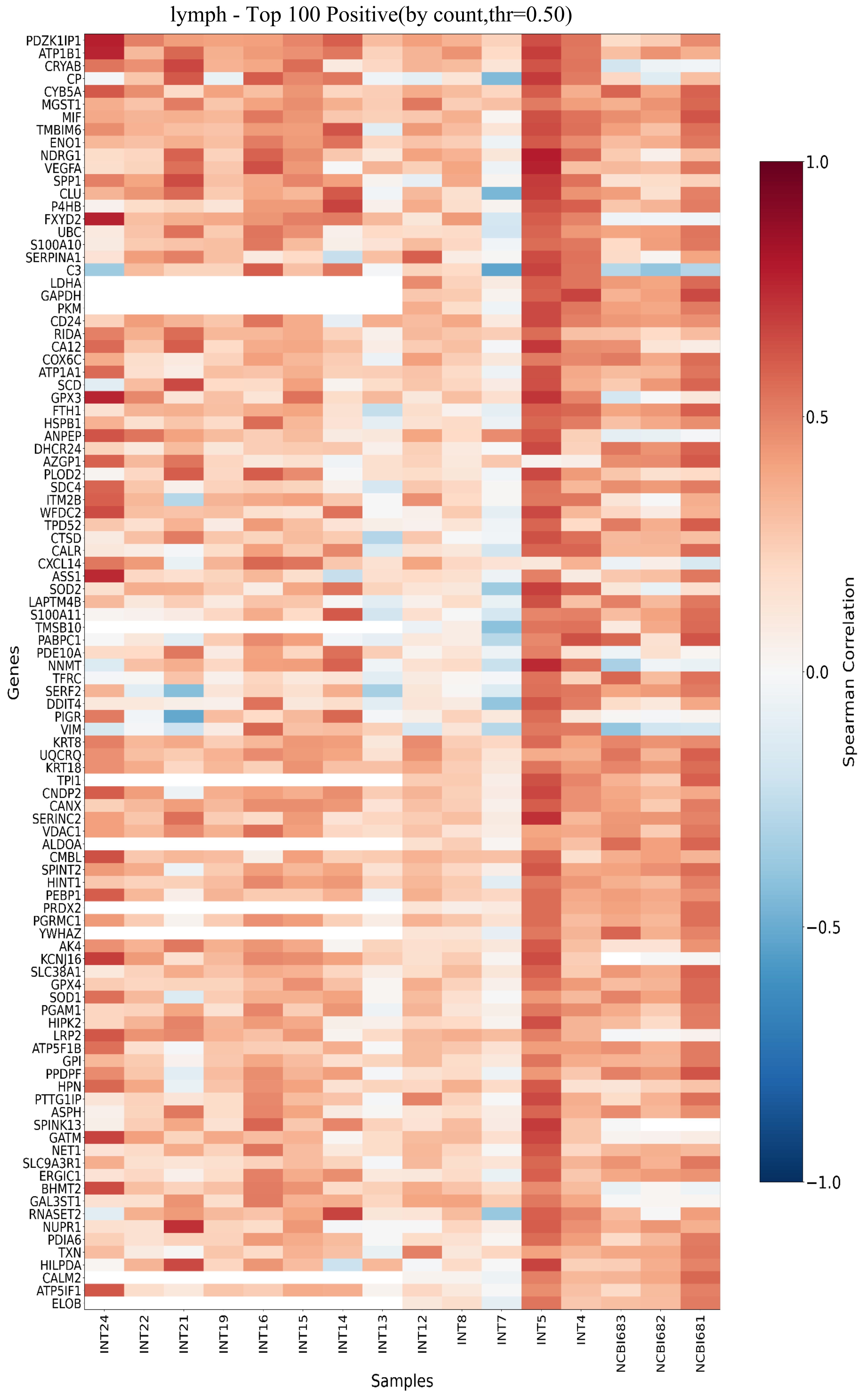


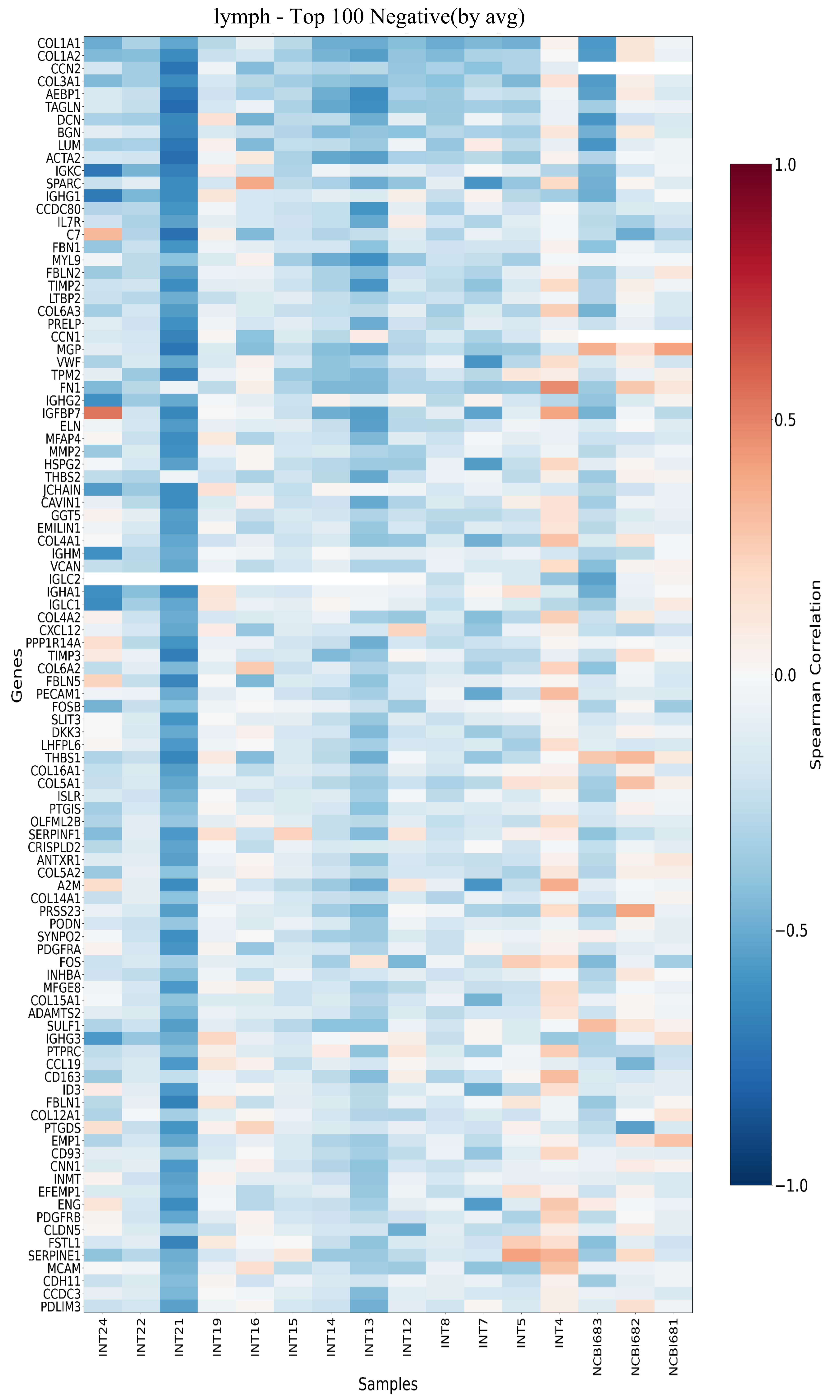


| Function | Genes |
|---|---|
| Renal Origin & Injury Markers | *LRP2 (Megalin), TFCP2L1, PDZK1IP1, SLC4A4, WFDC2* |
| Metabolic Reprogramming & Hypoxia Adaptation | *HK2, LDHA, GAPDH, ENO1, PKM, NDRG1, DDIT4, EGLN3, VEGFA* |
| Proteostasis & Stress Response | *CRYAB, HSPB1, P4HB, SOD2, GPX3, MT1F/G, TXN* |
| LN Microenvironment Remodeling & Immune Modulation | *SPP1 (OPN), C3, MIF, CXCL14, ANGPTL4, ITGB8, SERPINA1* |
| EMT, Invasion, Migration & Drug Resistance | *VIM, EPCAM, MUC1, MET, DDR1,S100A10/A11, ABCC3* |

| Function | Genes |
|---|---|
| HEV & Vascular Network Architecture | *CLDN5, PECAM1 (CD31), VWF, ENG, (CD105), CD93* |
| Dense ECM Scaffolding | *COL1A1/2, COL3A1, COL4A1/2, COL5A1/2, COL6A2/3, ELN, FBN1* |
| Lymphocyte Homing, Positioning & Humoral Immunity | *CCL19, CXCL12, IL7R, PTPRC (CD45) JCHAIN, IGHG1/2/3, IGHA1, IGHM, IGKC, IGLC1/2* |
| Stromal Remodeling Balance & Homeostasis | *MMP2, TIMP2/3, SPARC, DCN, BGN, LUM, VCAN* |
| Macrophage Clearance & Myofibroblast Contraction | *CD163, MFGE8, ACTA2 (α-SMA), TAGLN, MYL9, TPM2* |

Fig.S5

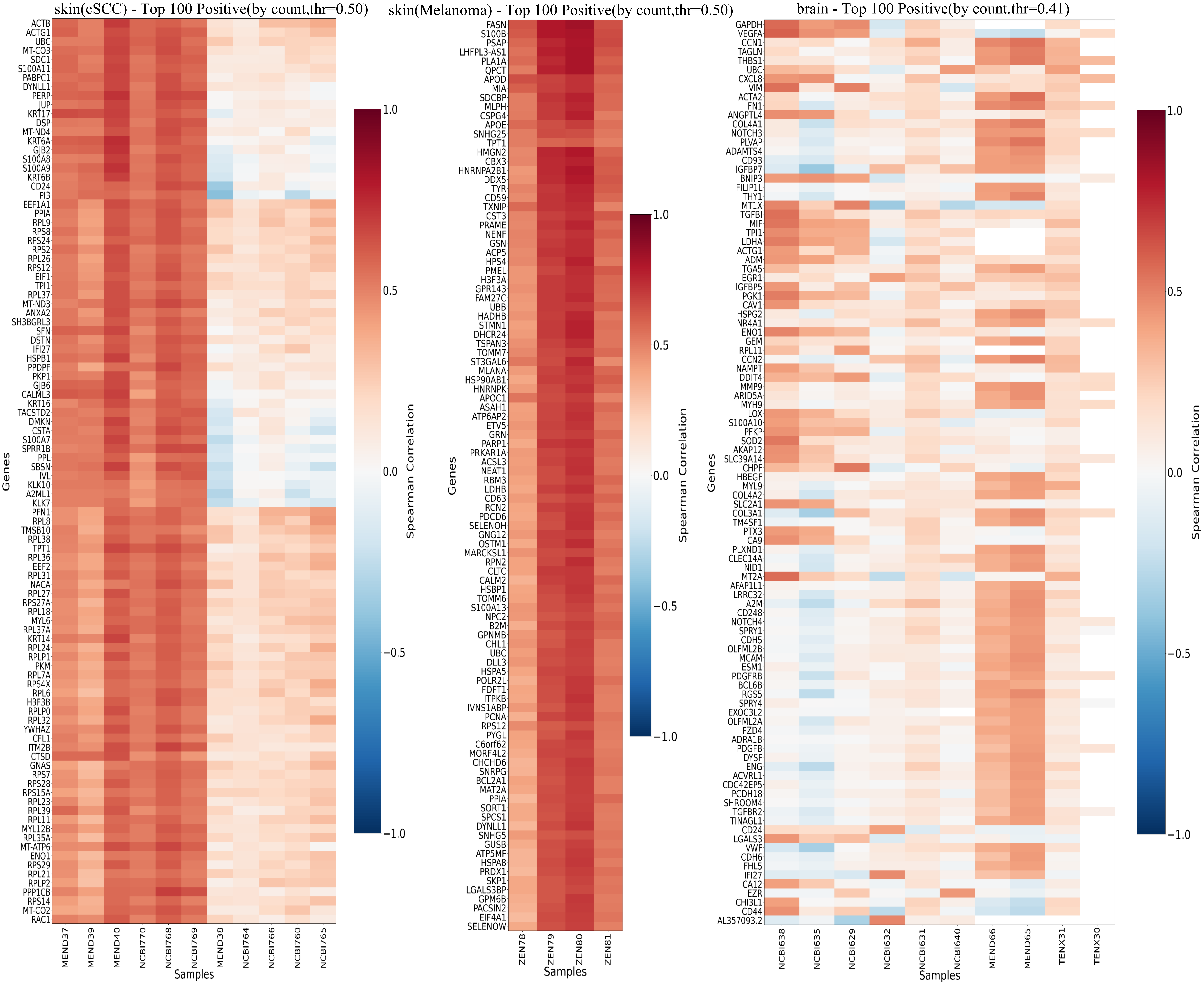


| Function | Genes |
|---|---|
| Hyper-proliferative Keratins & Cytoskeleton | *KRT6A, KRT6B, KRT6C KRT16, KRT17* |
| Pro-inflammatory S100 & Innate Immunity | *S100A8, S100A9, S100A7 (Psoriasin)* |
| Gap Junctions & Intercellular Communication | *GJB2 (Connexin 26), GJB6 (Connexin 30)* |
| Metabolic Reprogramming & Biosynthesis | *RPL/RPS系列, MT-ND/CO/CYB系列* |

| Function | Genes |
|---|---|
| Master Regulators & Melanogenesis | *SOX10 , TYR, PMEL, MLANA (MART-1), GPR143* |
| Lipid Metabolism & Sphingolipid Processing | *FASN, APOE, ASAH1, PSAP* |
| Tumor-Associated Factors & Invasion Markers | *S100B, MIA (Melanoma Inhibitory Activity), CSPG4 (NG2)* |

| Function | Genes |
|---|---|
| Severe Hypoxia Drive & Metabolic Adaptation | *CA9, BNIP3, SLC2A1 (GLUT1)* |
| Abnormal Angiogenesis & BBB Disruption | *VEGFA, PLVAP (Plasmalemma Vesicle Associated Protein)* |
| Basement Membrane Remodeling & Pericyte Dysregulation | *MMP9, ADAMTS4 PDGFB, RGS5* |
| Immunosuppressive Inflammatory Microenvironment | *CXCL8 (IL-8), TGFBR2, CHI3L1 (YKL-40)* |

Fig.S6

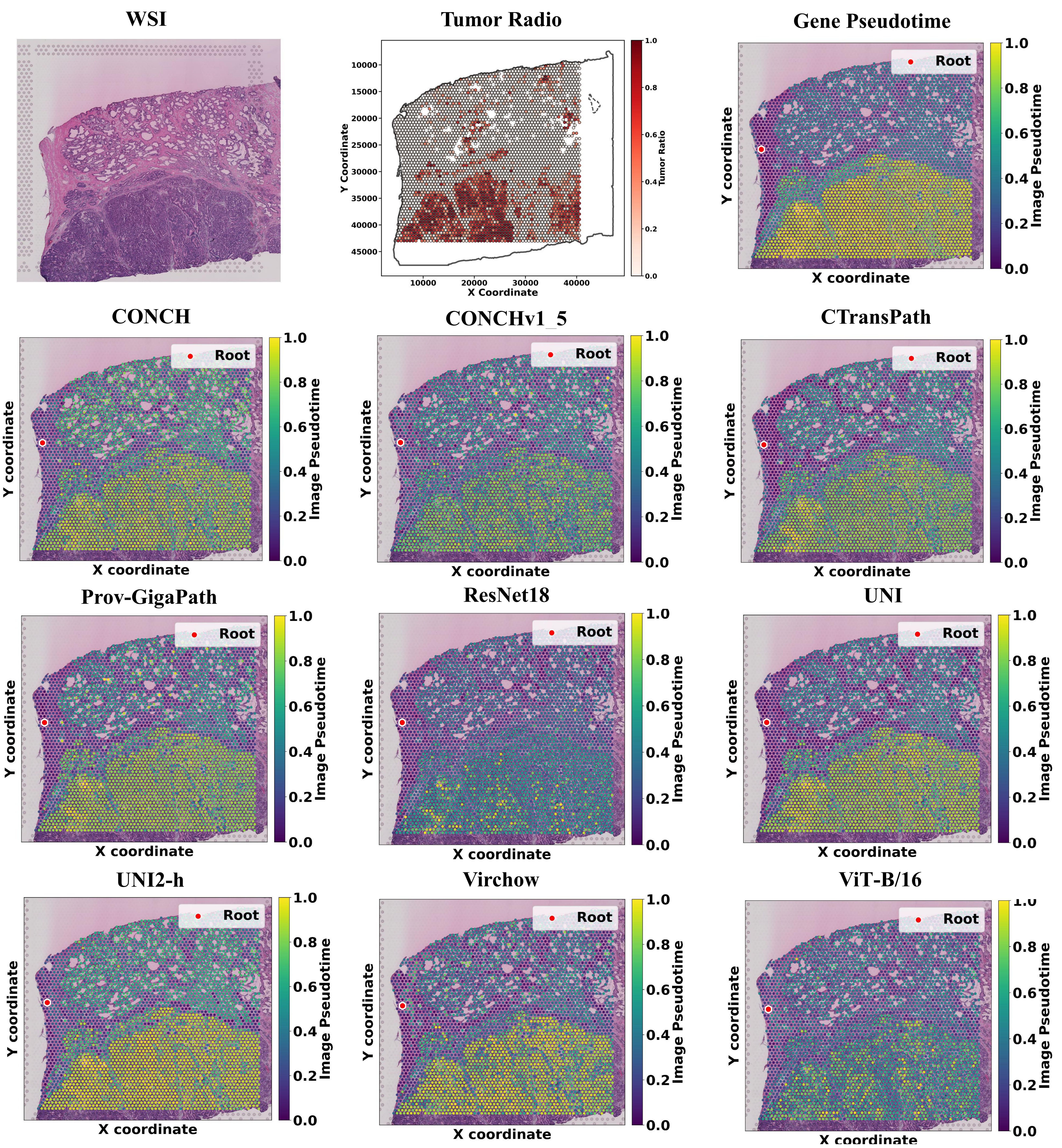


MEND62

Fig.S7

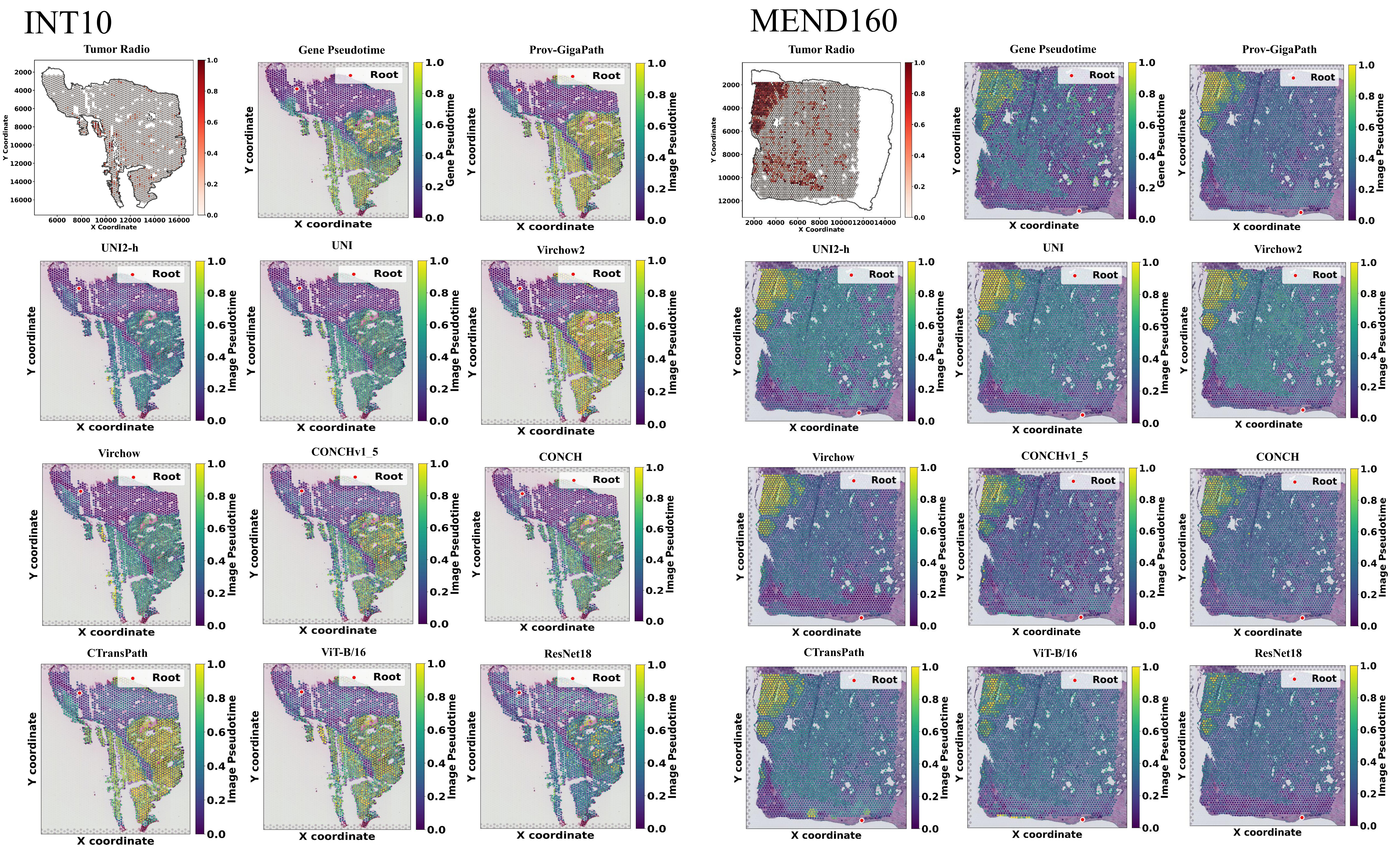
INT10
MEND160
Tumor Radio
Gene Pseudotime
Prov-GigaPath
UNI2-h
UNI
Virchow2
Virchow
CONCHv1_5
CONCH
CTransPath
ViT-B/16
ResNet18
Root
X coordinate
Y coordinate
X Coordinate
Y Coordinate
Gene Pseudotime
Image Pseudotime

Fig.S8

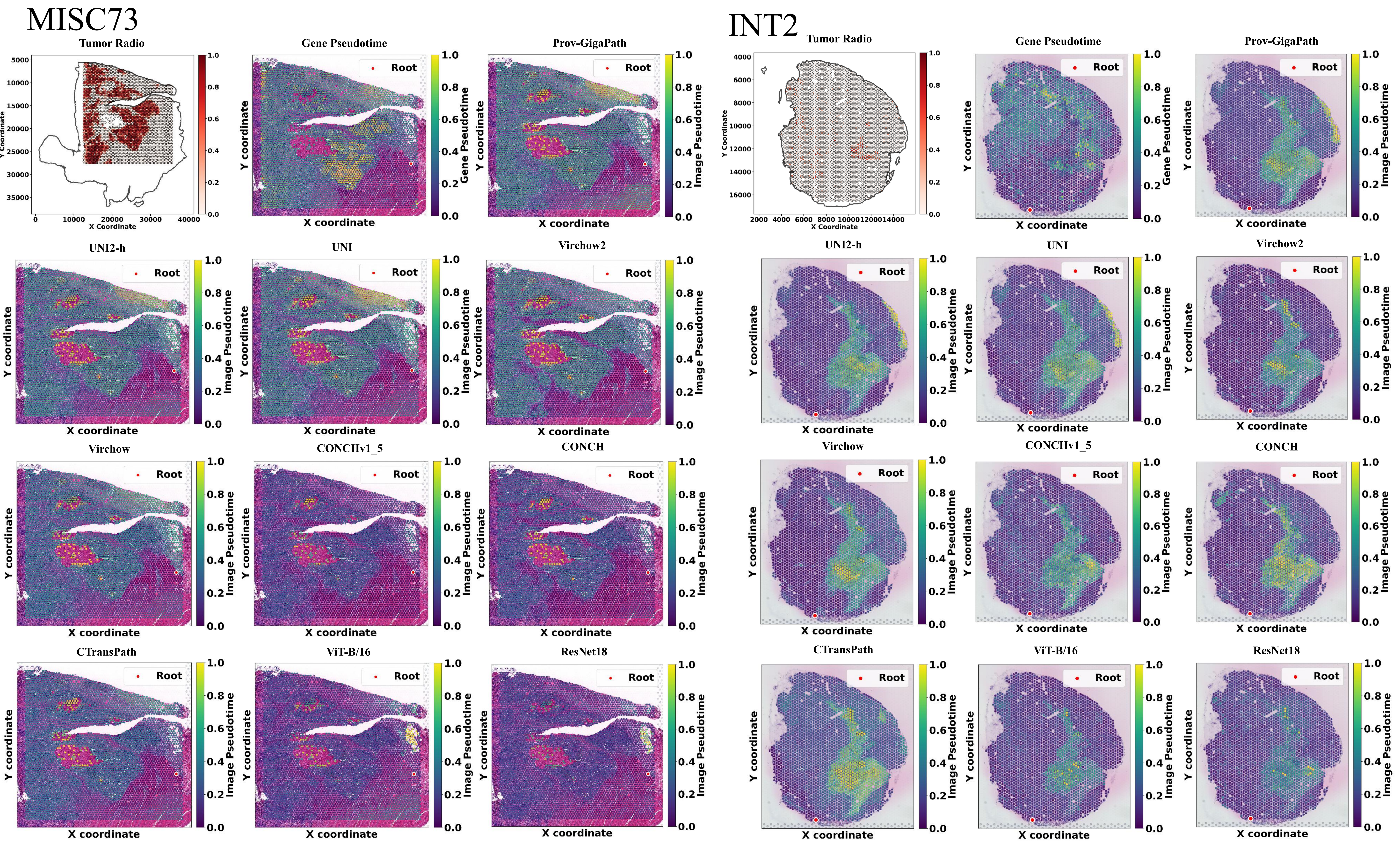
MISC73
INT2
Tumor Radio
Gene Pseudotime
Prov-GigaPath
UNI2-h
UNI
Virchow2
Virchow
CONCHv1_5
CONCH
CTransPath
ViT-B/16
ResNet18
Root
X coordinate
Y coordinate
X Coordinate
Y Coordinate
Gene Pseudotime
Image Pseudotime

Fig.S9

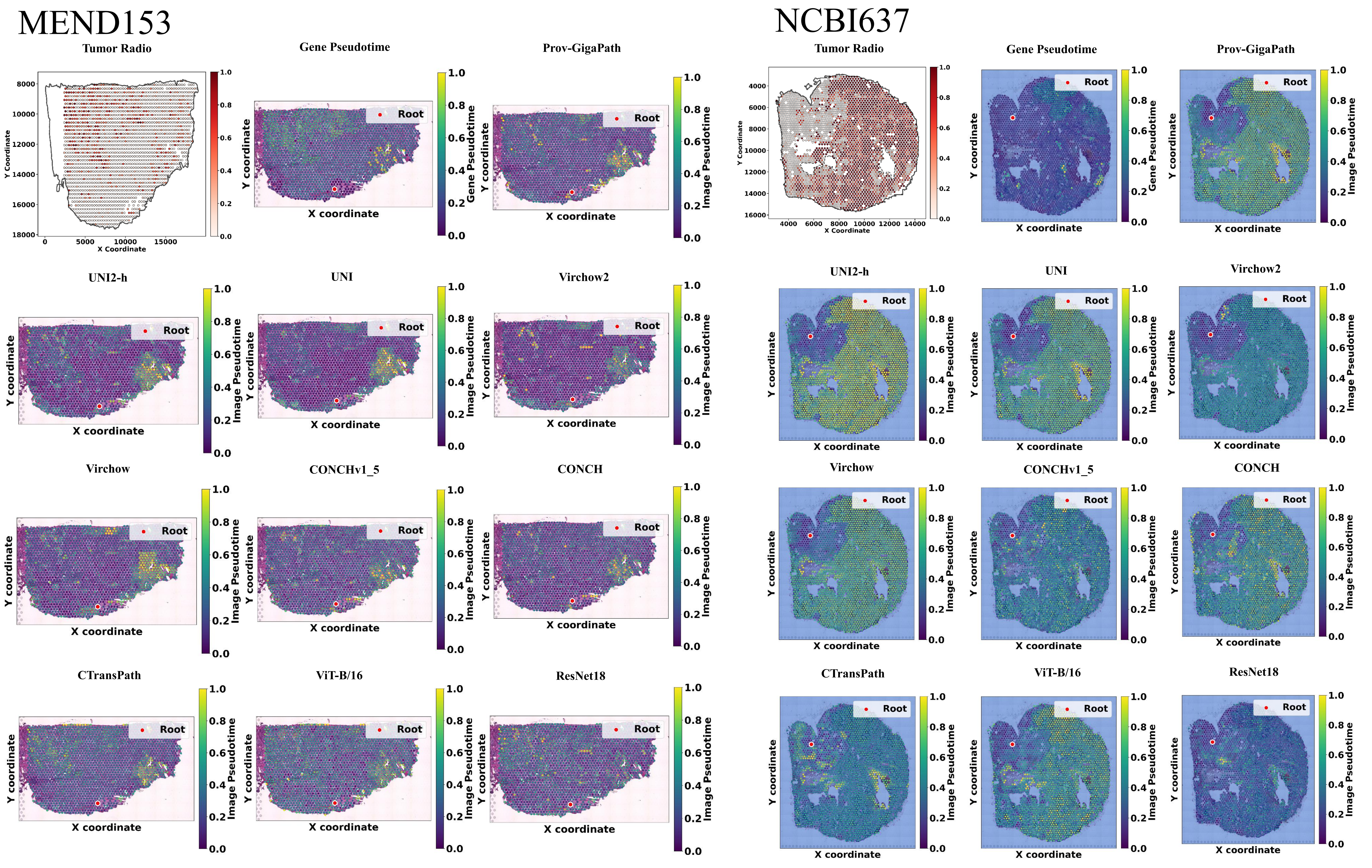
MEND153
NCBI637
Tumor Radio
Gene Pseudotime
Prov-GigaPath
UNI2-h
UNI
Virchow2
Virchow
CONCHv1_5
CONCH
CTransPath
ViT-B/16
ResNet18
Root
X coordinate
Y coordinate
X Coordinate
Y Coordinate
Gene Pseudotime
Image Pseudotime

Fig.S10

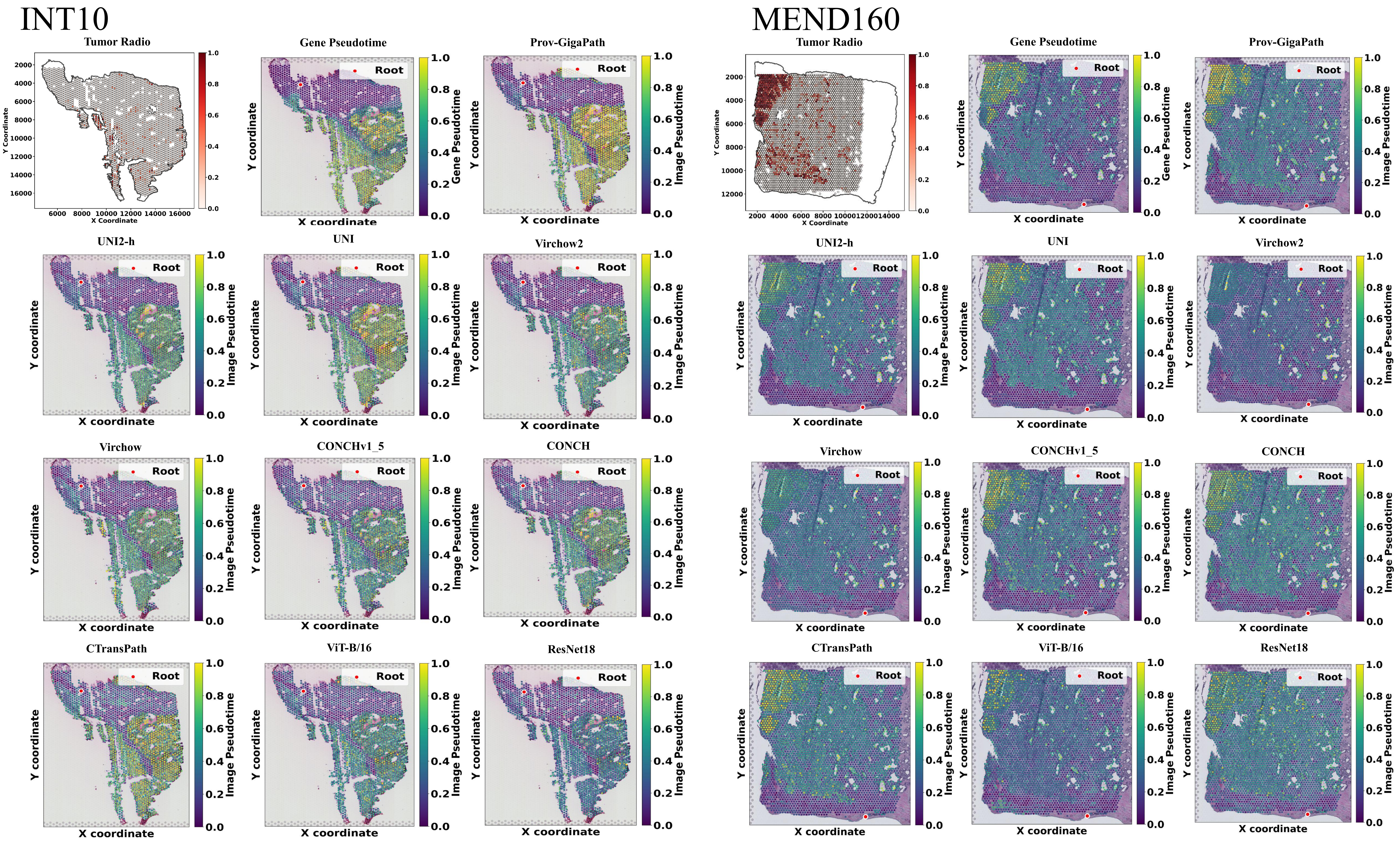
INT10
MEND160
Tumor Radio
Gene Pseudotime
Prov-GigaPath
UNI2-h
UNI
Virchow2
Virchow
CONCHv1_5
CONCH
CTransPath
ViT-B/16
ResNet18
Root
X coordinate
Y coordinate
X Coordinate
Y Coordinate
Gene Pseudotime
Image Pseudotime

Fig.S11

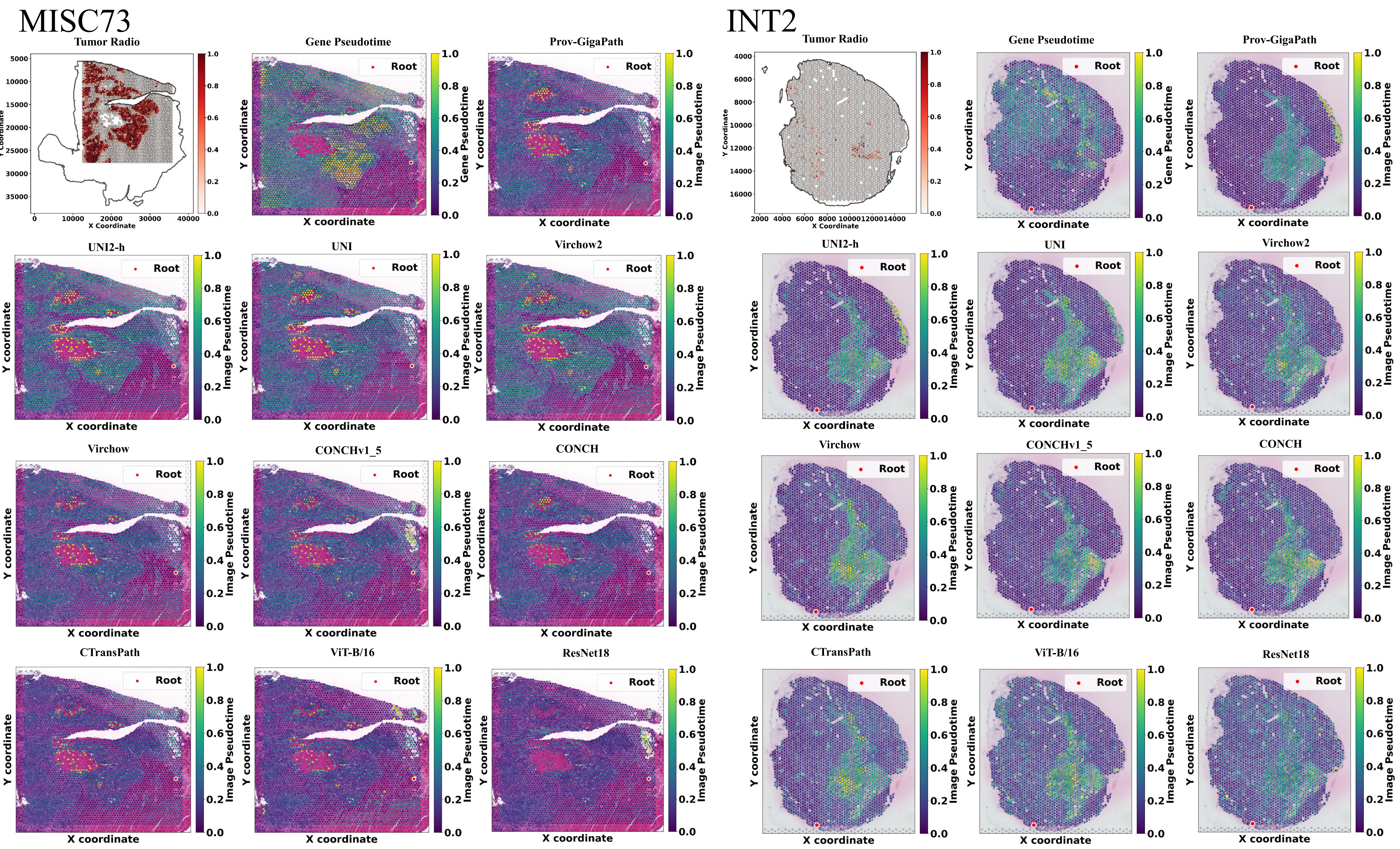
MISC73
Tumor Radio
X Coordinate
Y Coordinate
Gene Pseudotime
Prov-GigaPath
UNI2-h
UNI
Virchow2
Virchow
CONCHv1_5
CONCH
CTransPath
ViT-B/16
ResNet18
Root
X coordinate
Y coordinate
Image Pseudotime
INT2

Fig.S12

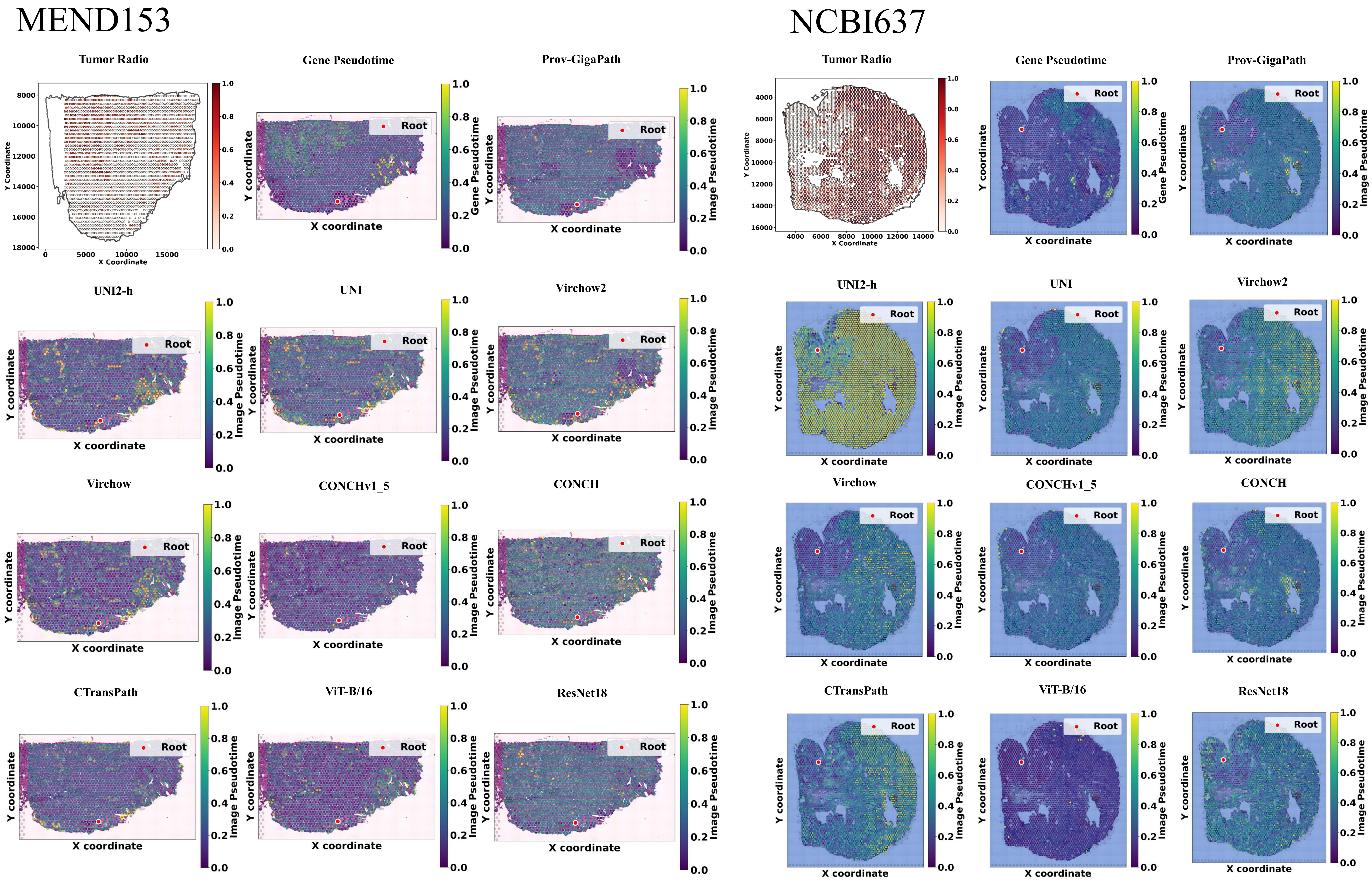
MEND153
NCBI637
Tumor Radio
Gene Pseudotime
Prov-GigaPath
UNI2-h
UNI
Virchow2
Virchow
CONCHv1_5
CONCH
CTransPath
ViT-B/16
ResNet18
Root
X coordinate
Y coordinate
X Coordinate
Y Coordinate
Gene Pseudotime
Image Pseudotime

### Fig.S13

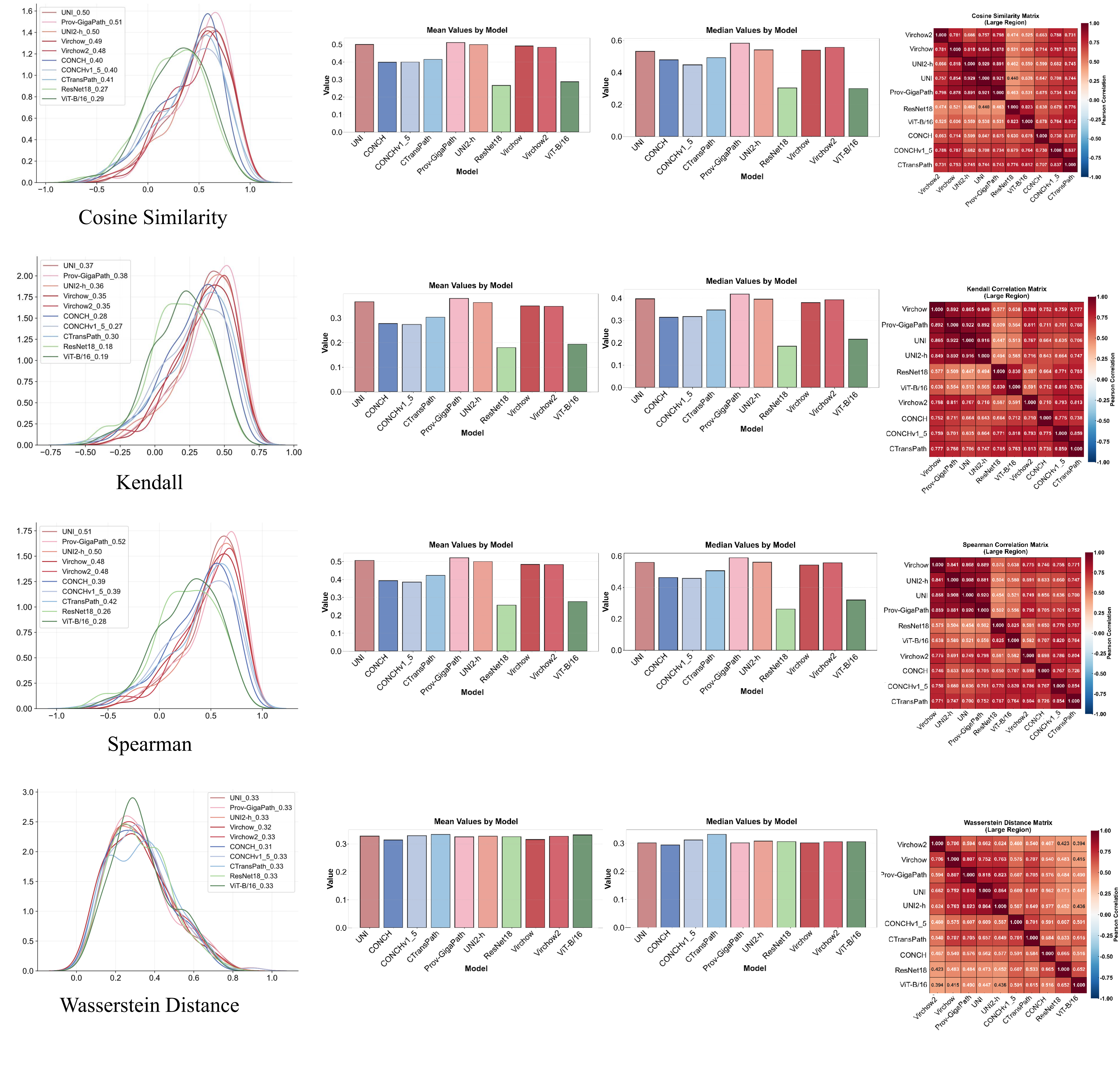
UNI_0.50
Prov-GigaPath_0.51
UNI2-h_0.50
Virchow_0.49
Virchow2_0.48
CONCH_0.40
CONCHv1_5_0.40
CTransPath_0.41
ResNet18_0.27
ViT-B/16_0.29
Cosine Similarity
Mean Values by Model
Median Values by Model
Value
Model
UNI
CONCH
CONCHv1_5
CTransPath
Prov-GigaPath
UNI2-h
ResNet18
Virchow
Virchow2
ViT-B/16
Cosine Similarity Matrix
(Large Region)
Pearson Correlation
UNI_0.37
Prov-GigaPath_0.38
UNI2-h_0.36
Virchow_0.35
Virchow2_0.35
CONCH_0.28
CONCHv1_5_0.27
CTransPath_0.30
ResNet18_0.18
ViT-B/16_0.19
Kendall
Kendall Correlation Matrix
(Large Region)
UNI_0.51
Prov-GigaPath_0.52
UNI2-h_0.50
Virchow_0.48
Virchow2_0.48
CONCH_0.39
CONCHv1_5_0.39
CTransPath_0.42
ResNet18_0.26
ViT-B/16_0.28
Spearman
Spearman Correlation Matrix
(Large Region)
UNI_0.33
Prov-GigaPath_0.33
UNI2-h_0.33
Virchow_0.32
Virchow2_0.33
CONCH_0.31
CONCHv1_5_0.33
CTransPath_0.33
ResNet18_0.33
ViT-B/16_0.33
Wasserstein Distance
Wasserstein Distance Matrix
(Large Region)

### Fig.S14

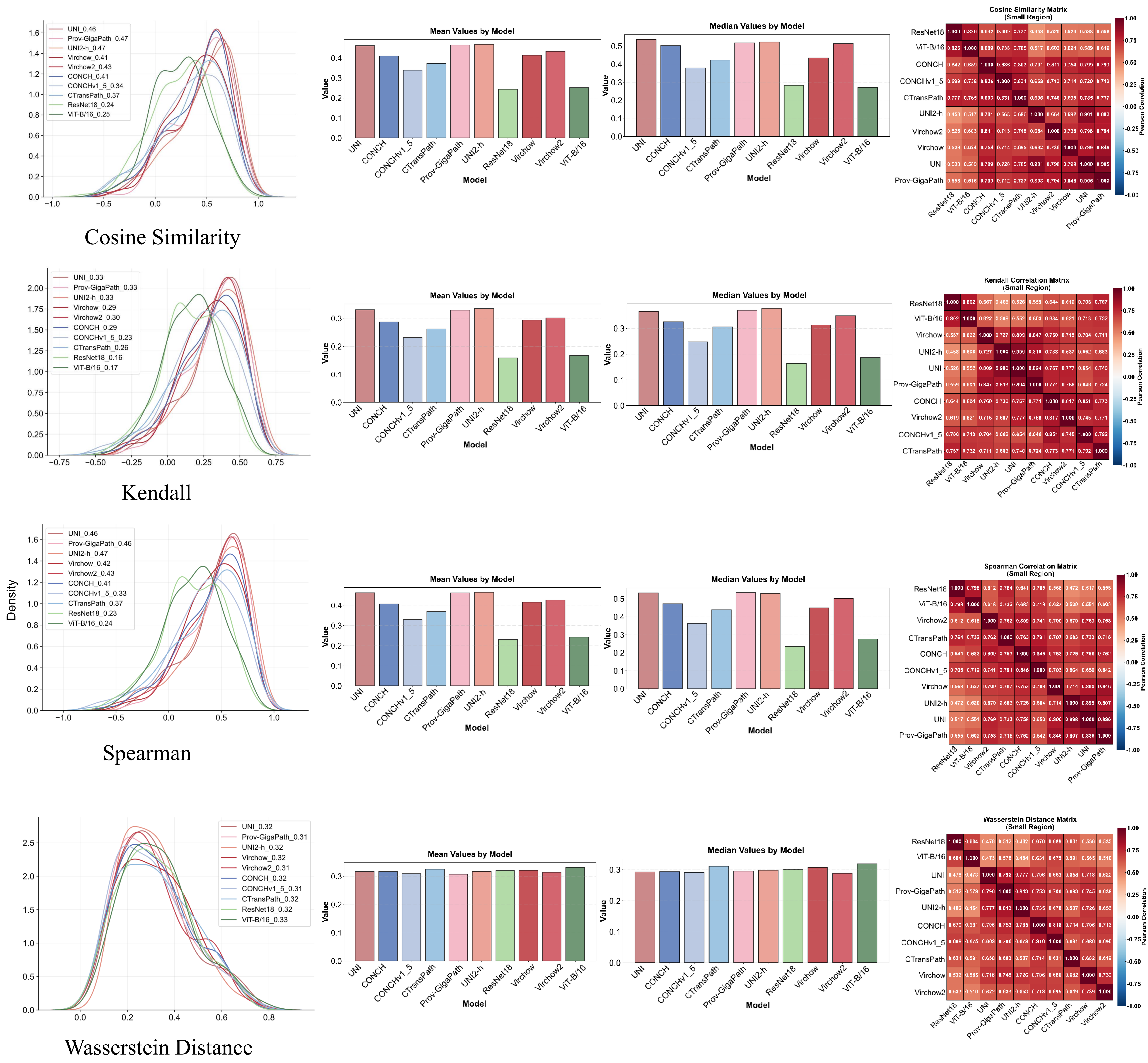
Cosine Similarity
UNI_0.46
Prov-GigaPath_0.47
UNI2-h_0.47
Virchow_0.41
Virchow2_0.43
CONCH_0.41
CONCHv1_5_0.34
CTransPath_0.37
ResNet18_0.24
ViT-B/16_0.25
Mean Values by Model
Median Values by Model
Value
Model
Cosine Similarity Matrix
(Small Region)
Kendall
UNI_0.33
Prov-GigaPath_0.33
UNI2-h_0.33
Virchow_0.29
Virchow2_0.30
CONCH_0.29
CONCHv1_5_0.23
CTransPath_0.26
ResNet18_0.16
ViT-B/16_0.17
Kendall Correlation Matrix
(Small Region)
Spearman
Density
UNI_0.46
Prov-GigaPath_0.46
UNI2-h_0.47
Virchow_0.42
Virchow2_0.43
CONCH_0.41
CONCHv1_5_0.33
CTransPath_0.37
ResNet18_0.23
ViT-B/16_0.24
Spearman Correlation Matrix
(Small Region)
Wasserstein Distance
UNI_0.32
Prov-GigaPath_0.31
UNI2-h_0.32
Virchow_0.32
Virchow2_0.31
CONCH_0.32
CONCHv1_5_0.31
CTransPath_0.32
ResNet18_0.32
ViT-B/16_0.33
Wasserstein Distance Matrix
(Small Region)
Pearson Correlation

### Fig.S15

#### Prov-GigaPath

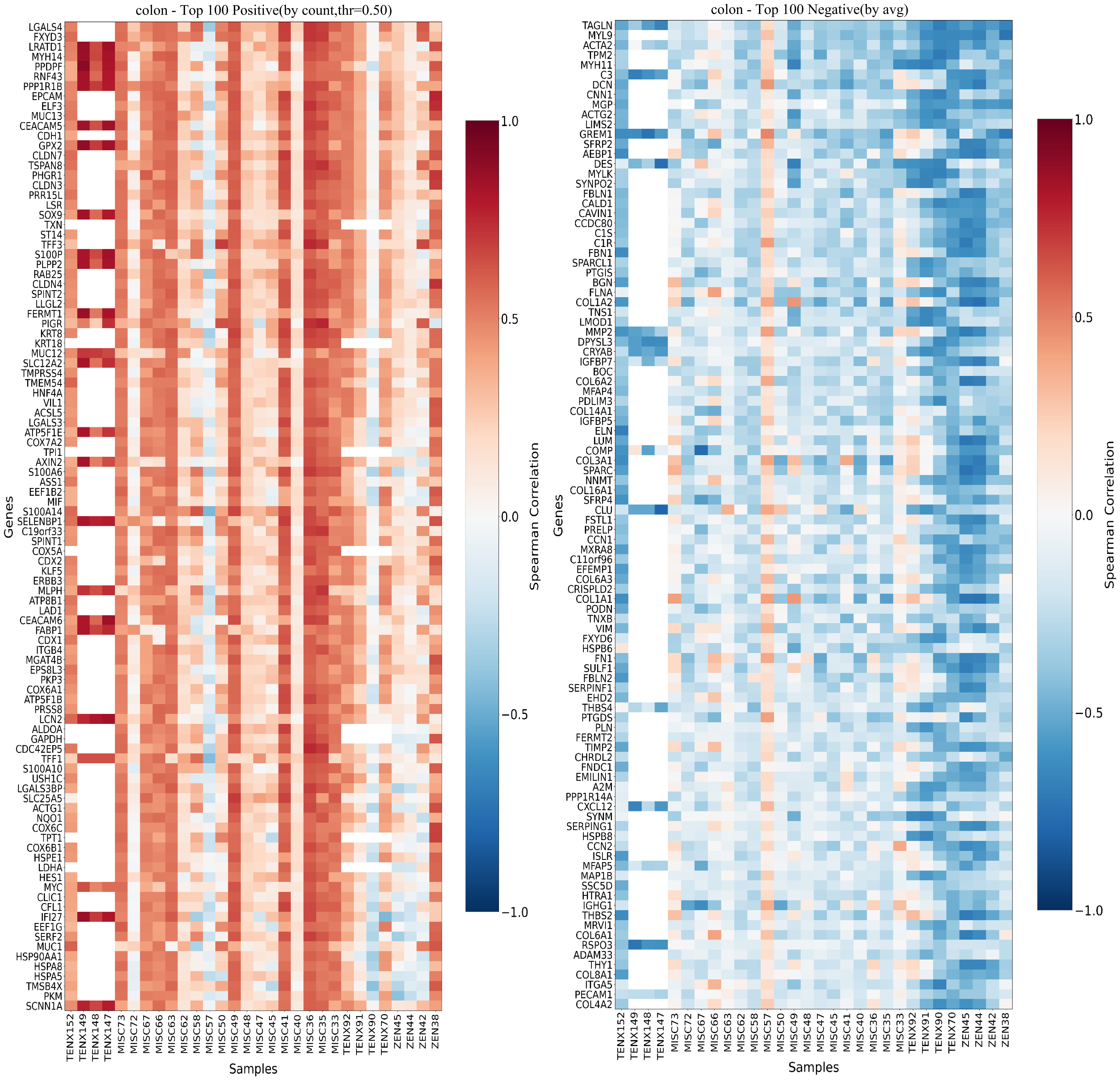

### Fig.S16

#### Prov-GigaPath

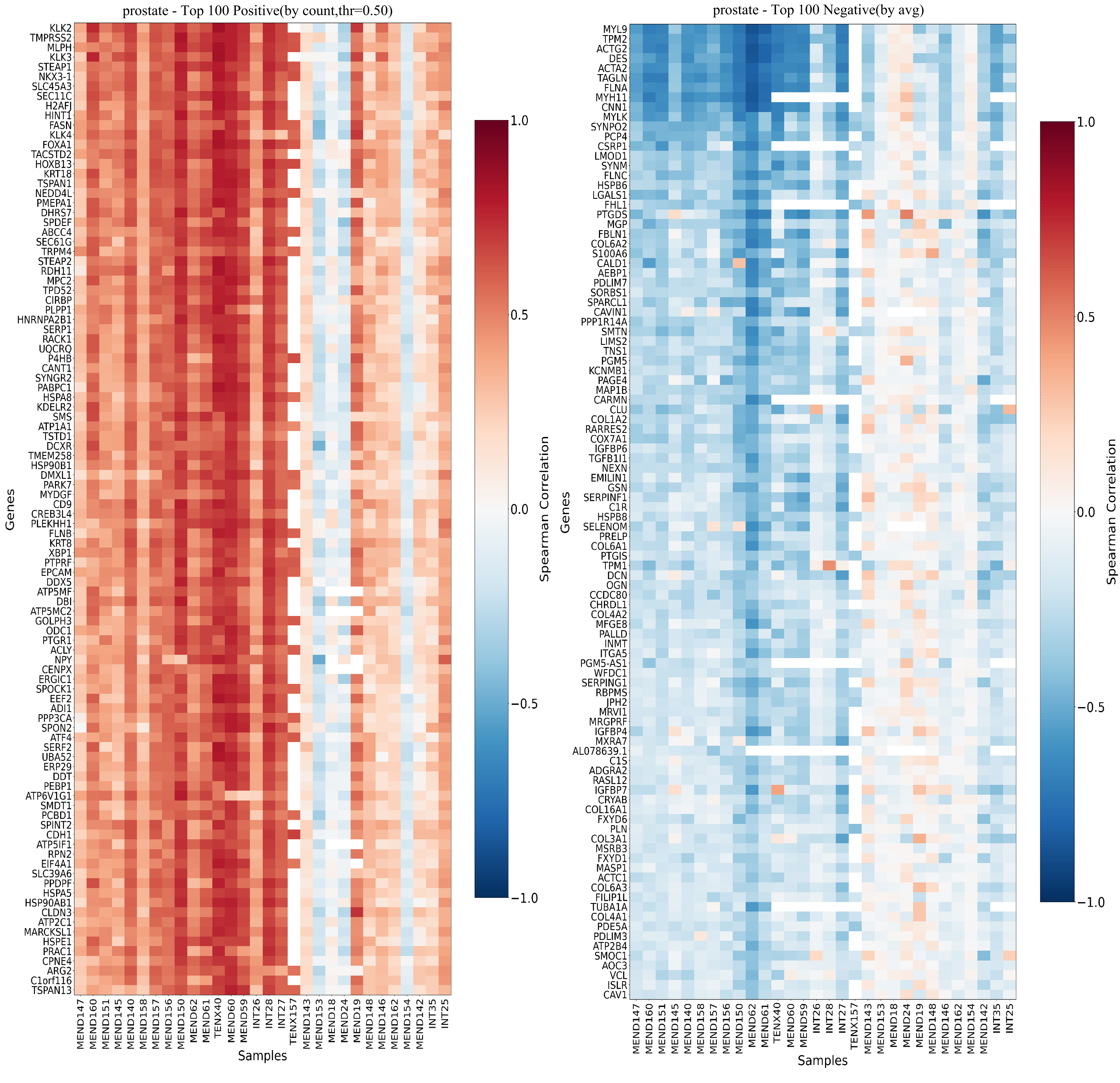

### Fig.S17

#### Prov-GigaPath

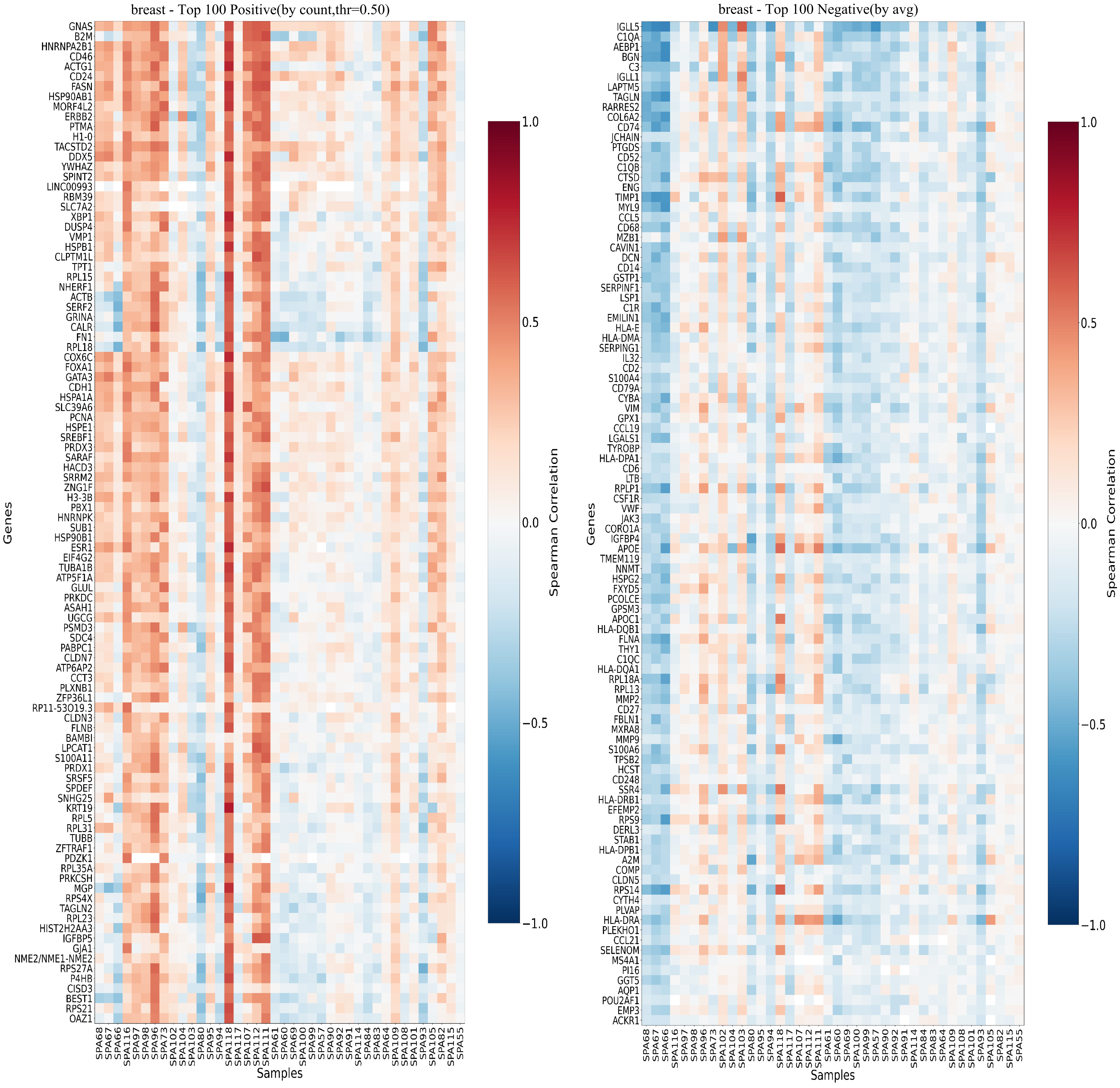

### Fig.S18

#### Prov-GigaPath

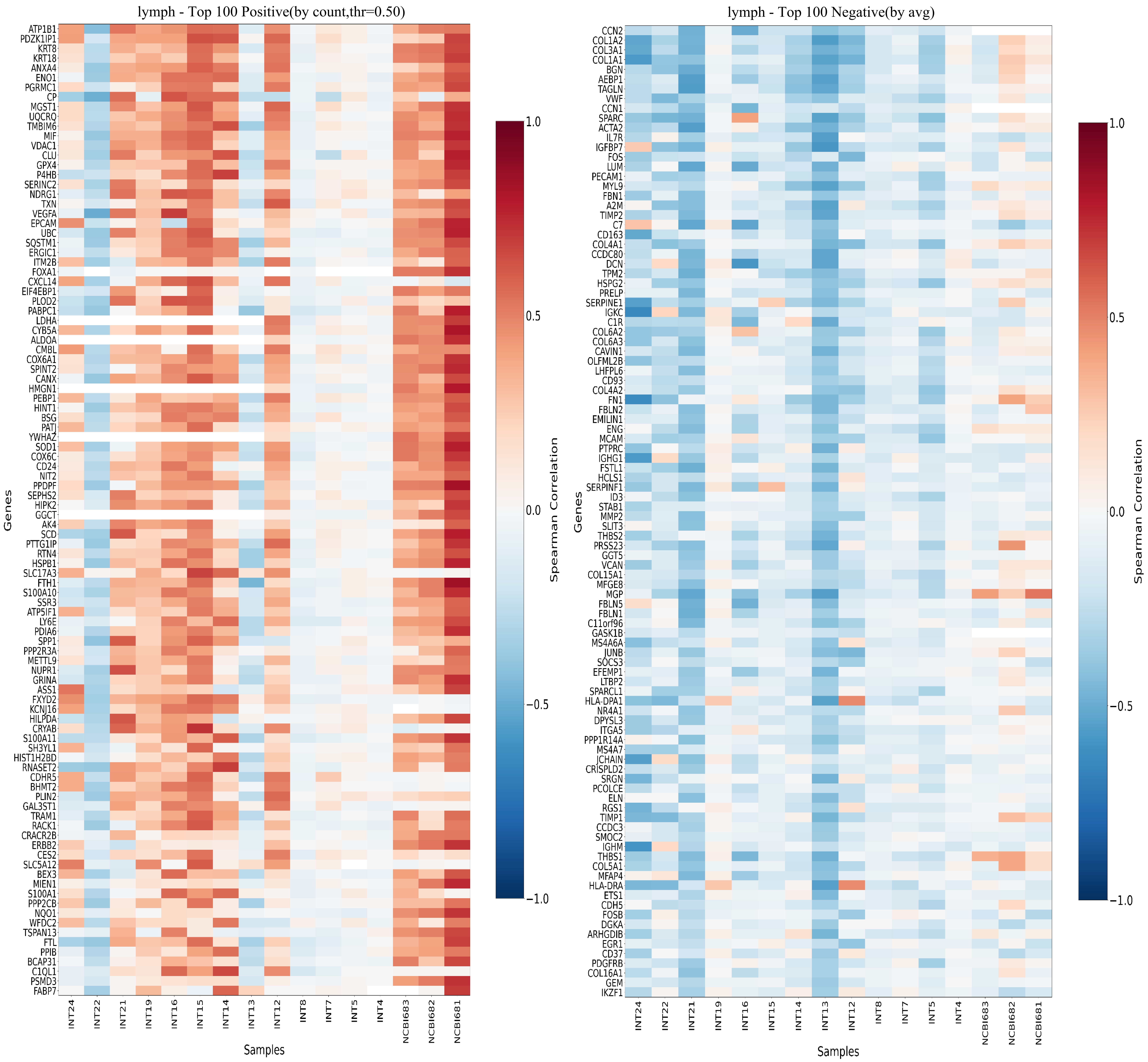

Fig.S19

UNI2-h

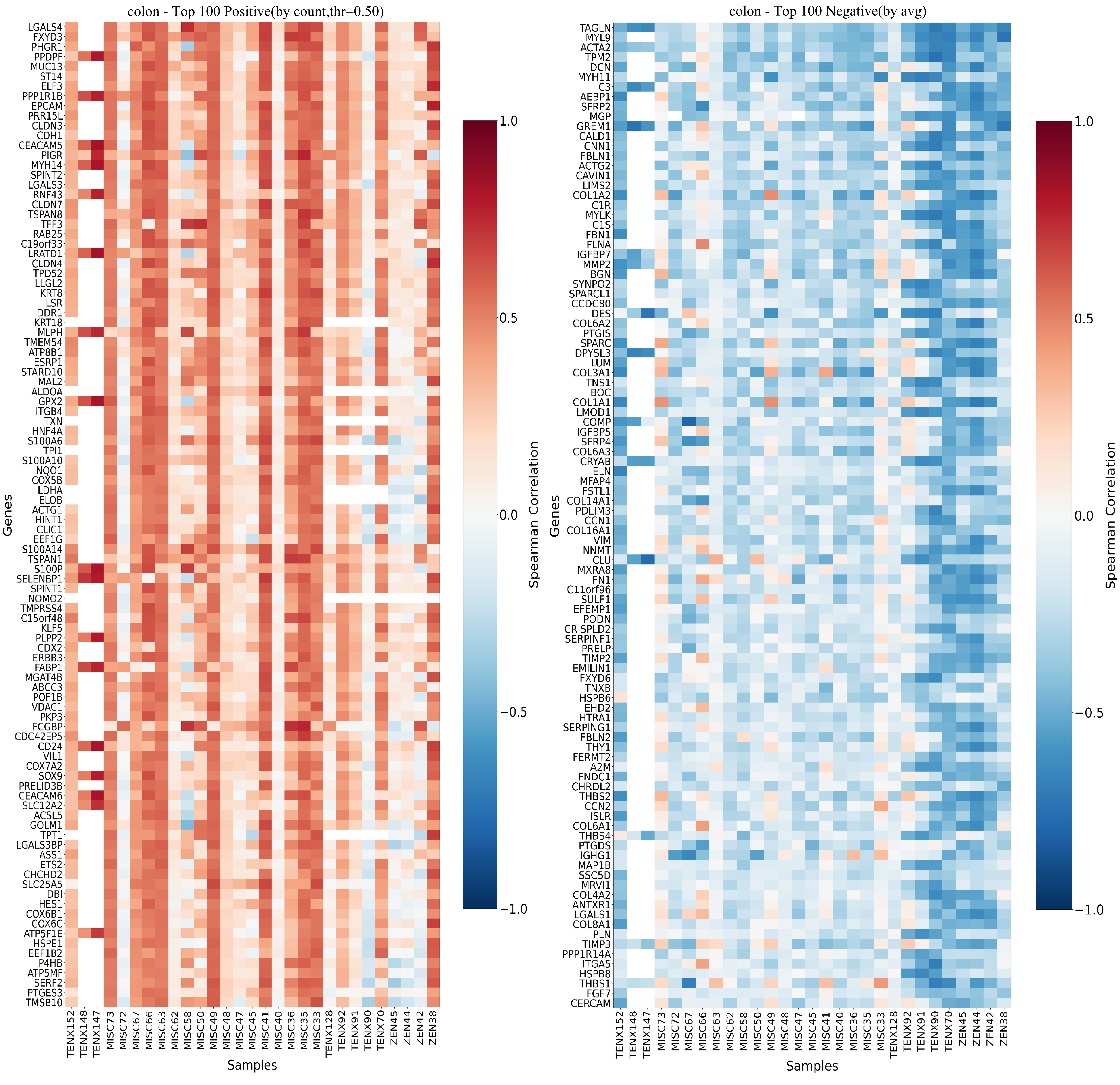

### Fig.S20

#### UNI2-h

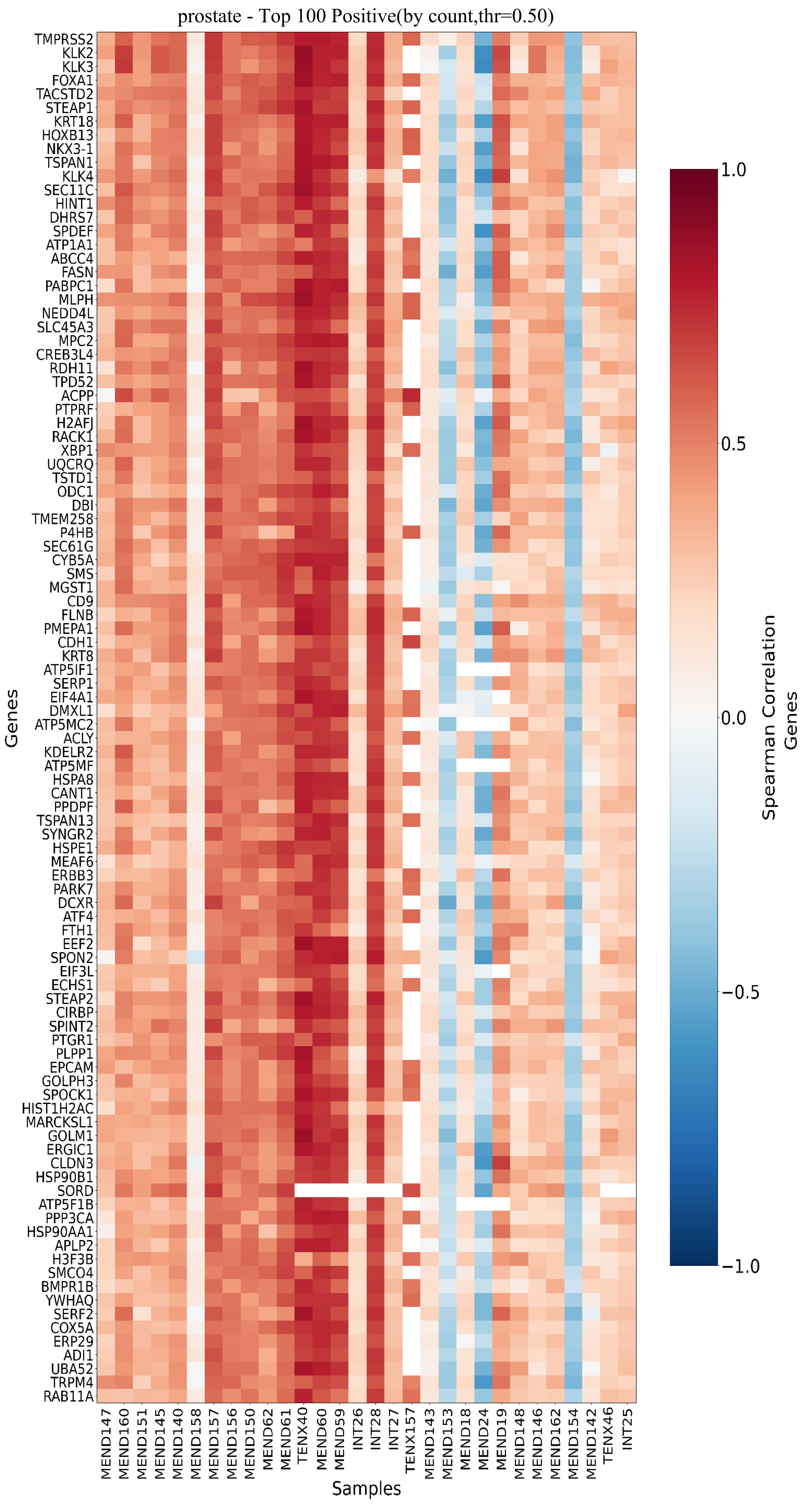


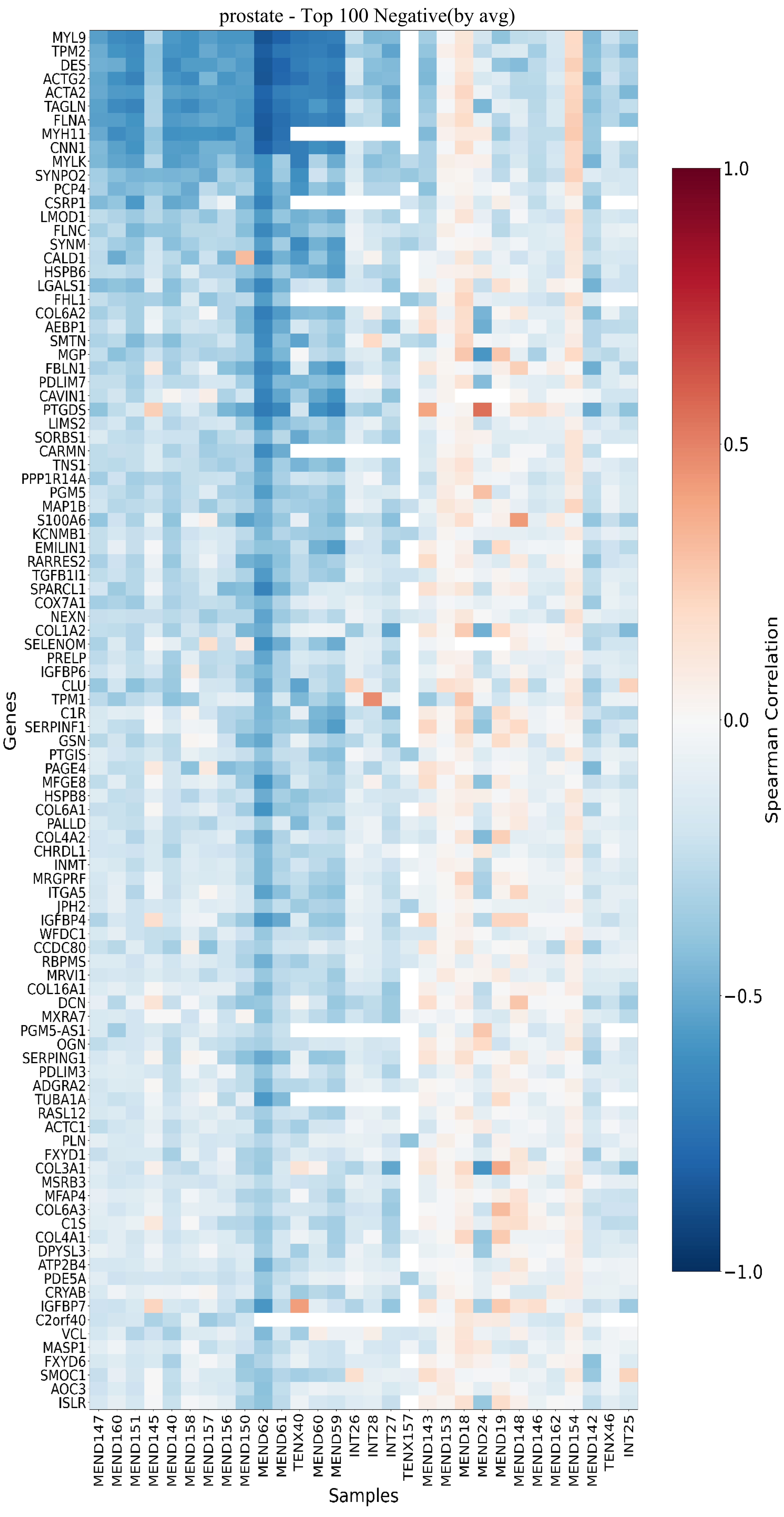

### Fig.S21

#### UNI2-h

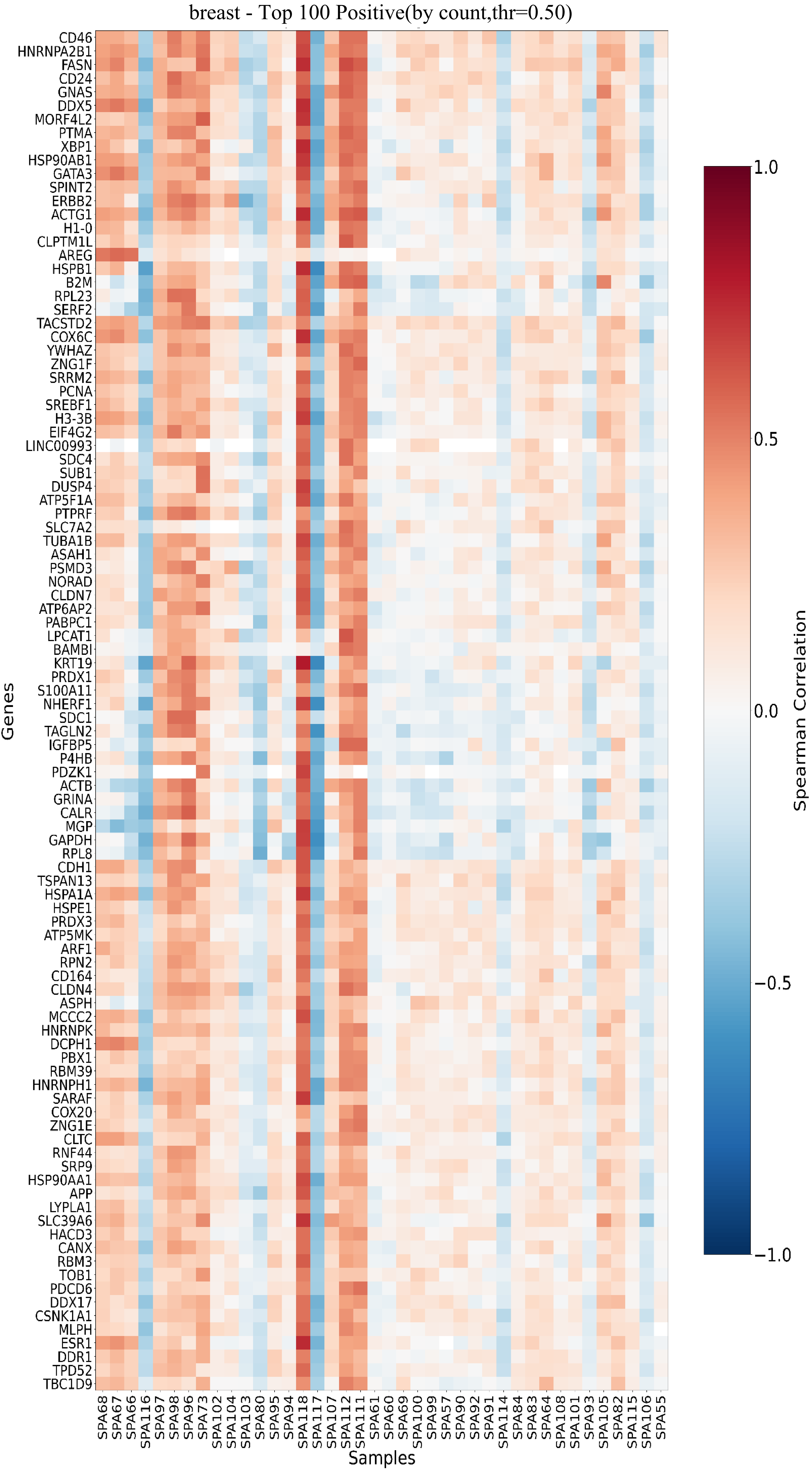


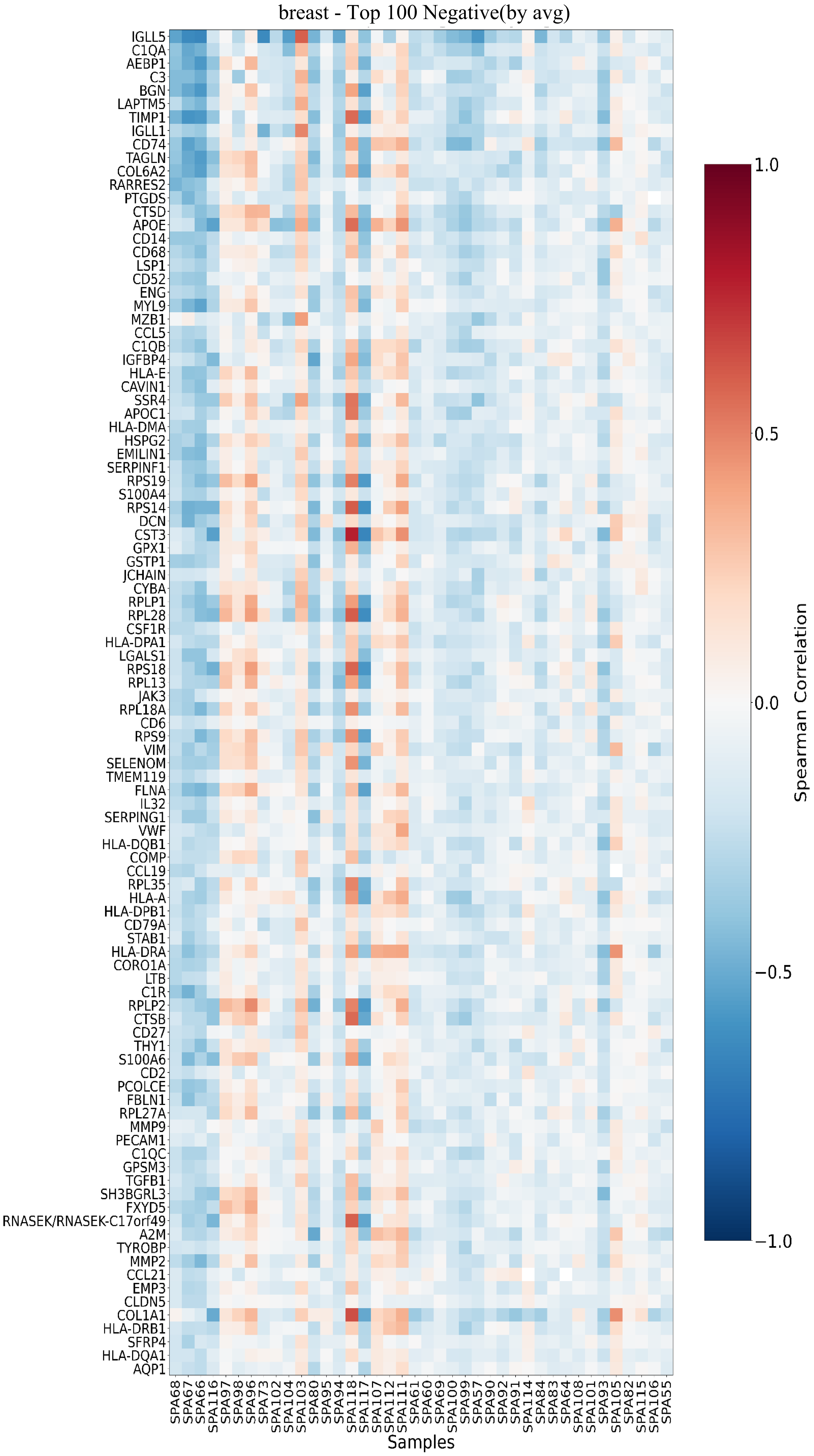

### Fig.S22

#### UNI2-h

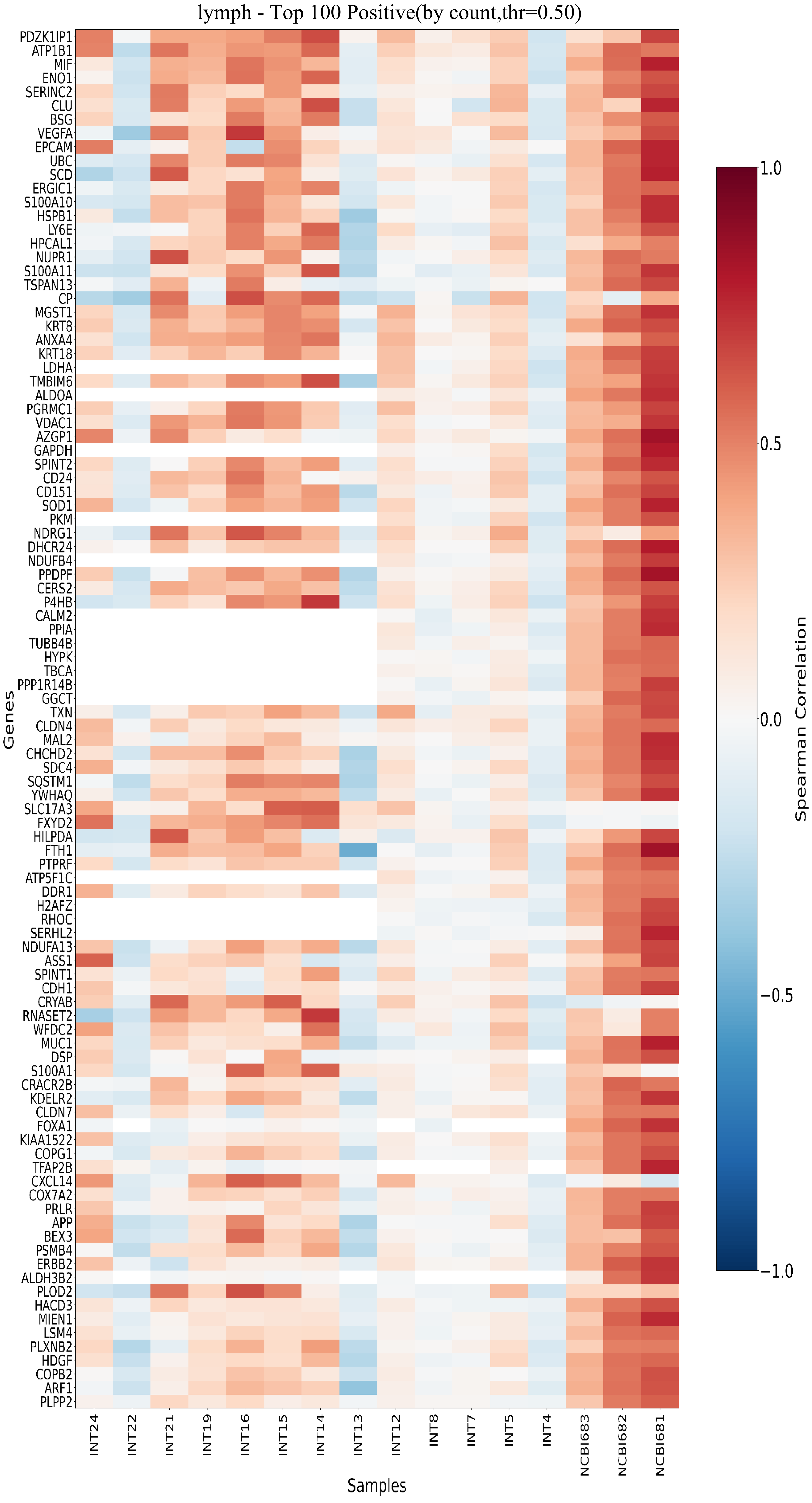


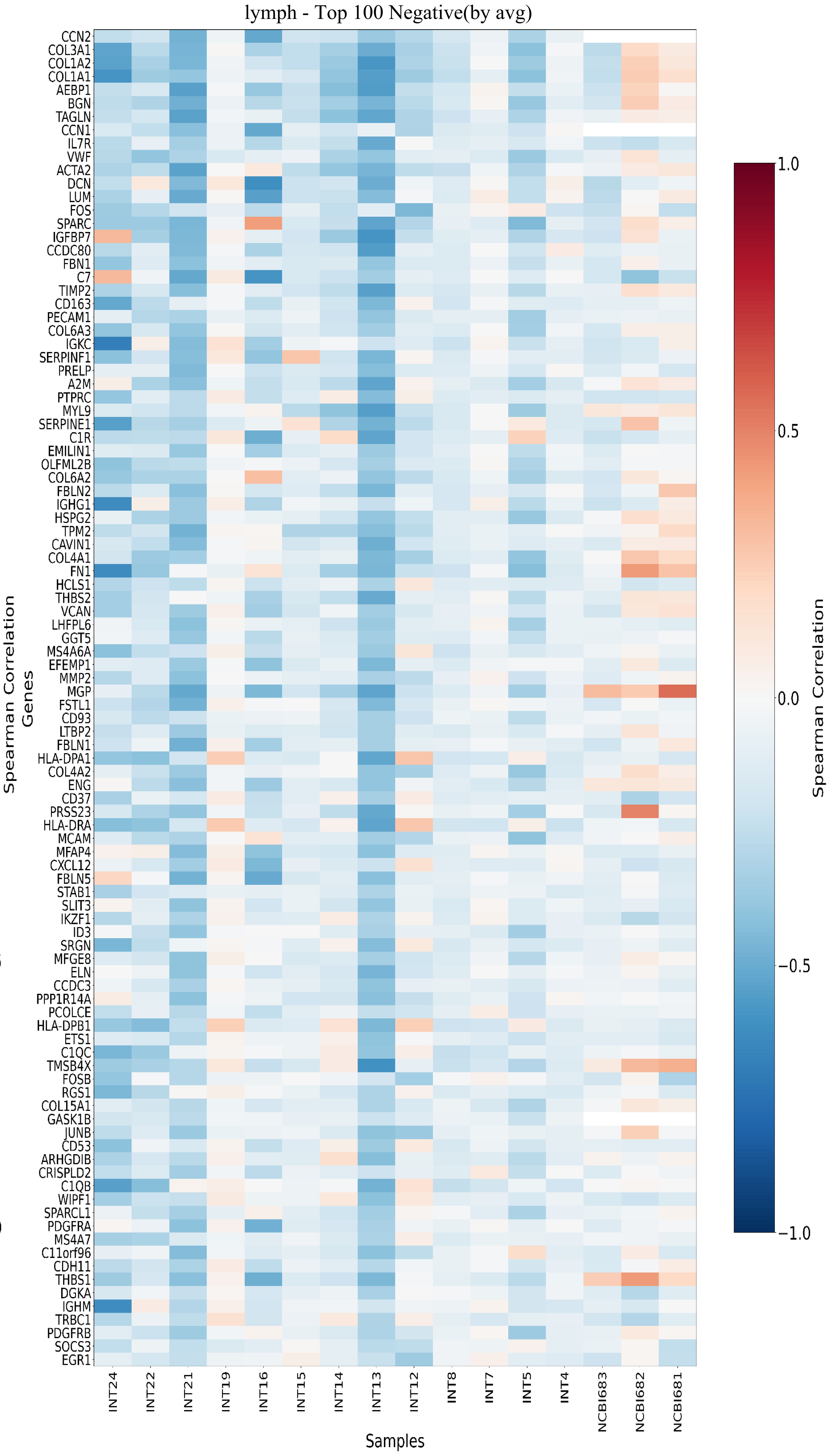

### Fig.S23

#### ResNet18

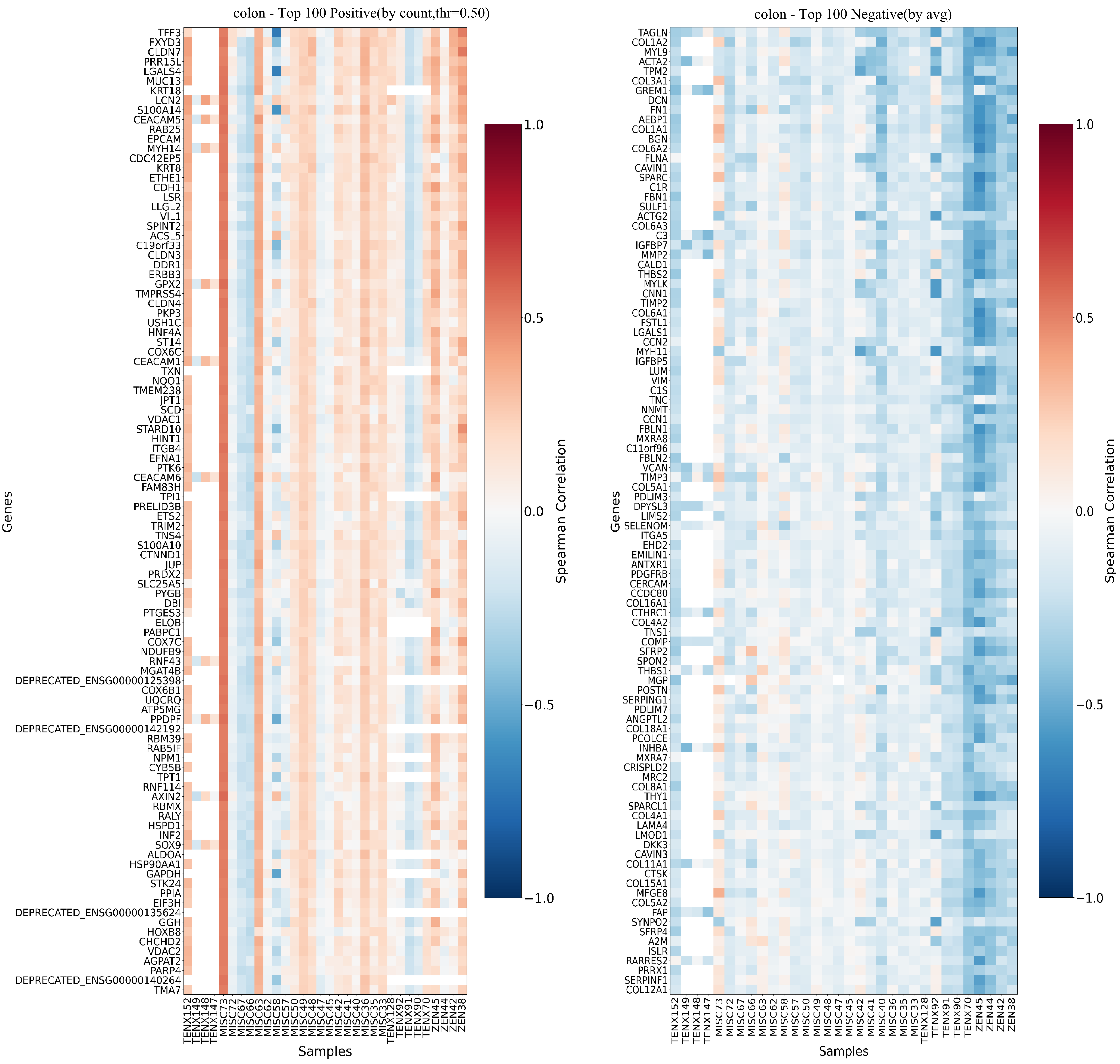

### Fig.S24

#### ResNet18

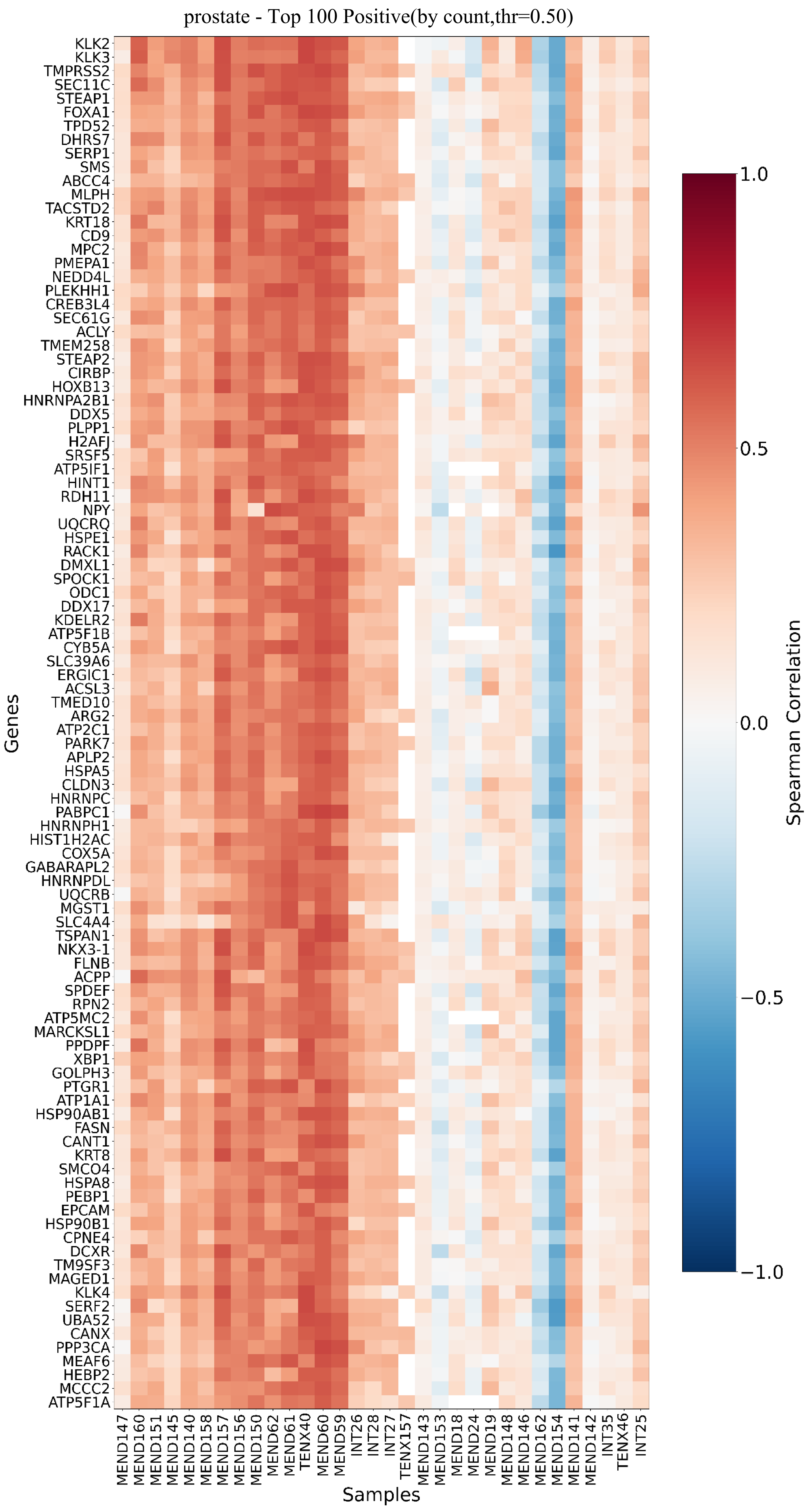


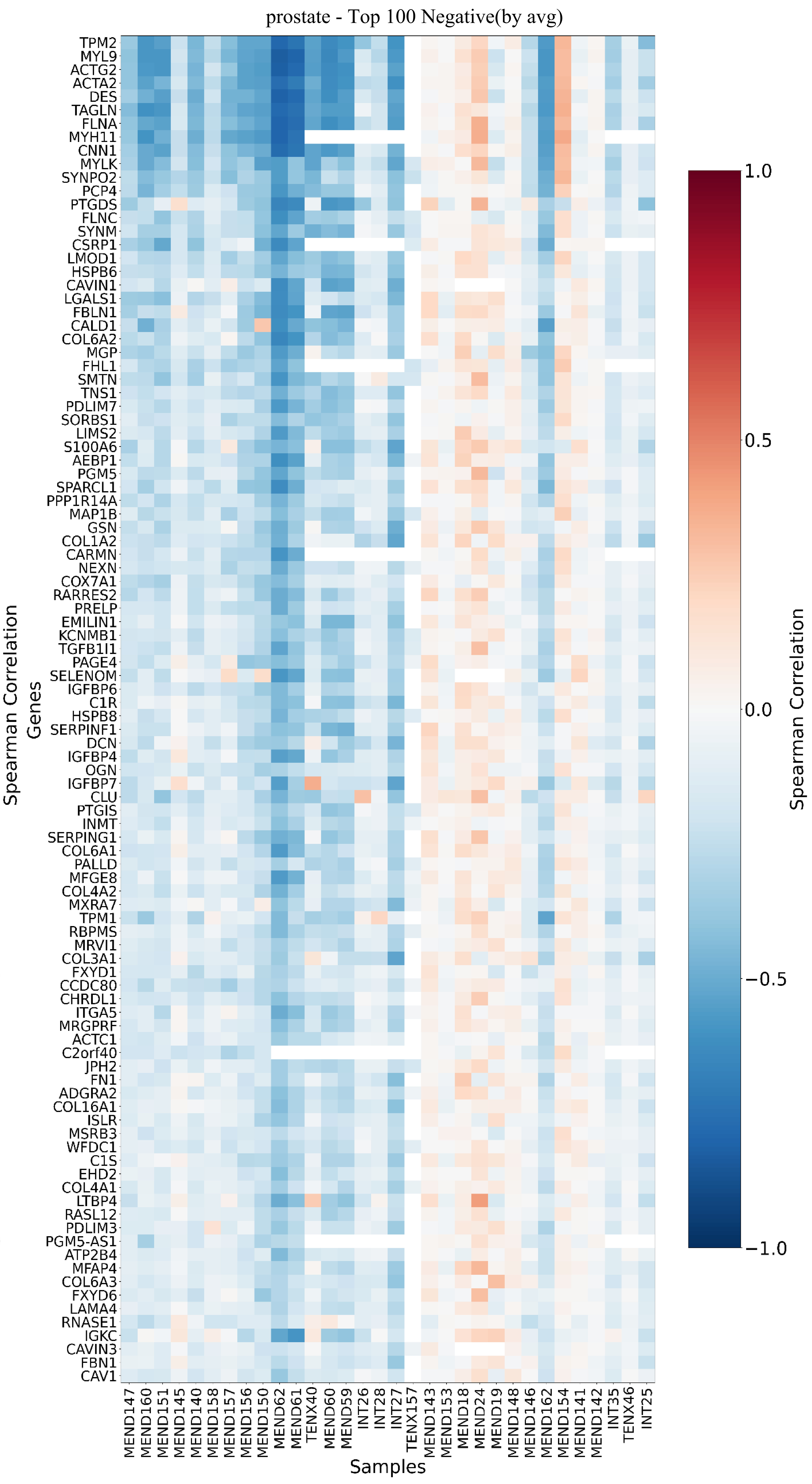

### Fig.S25

#### ResNet18

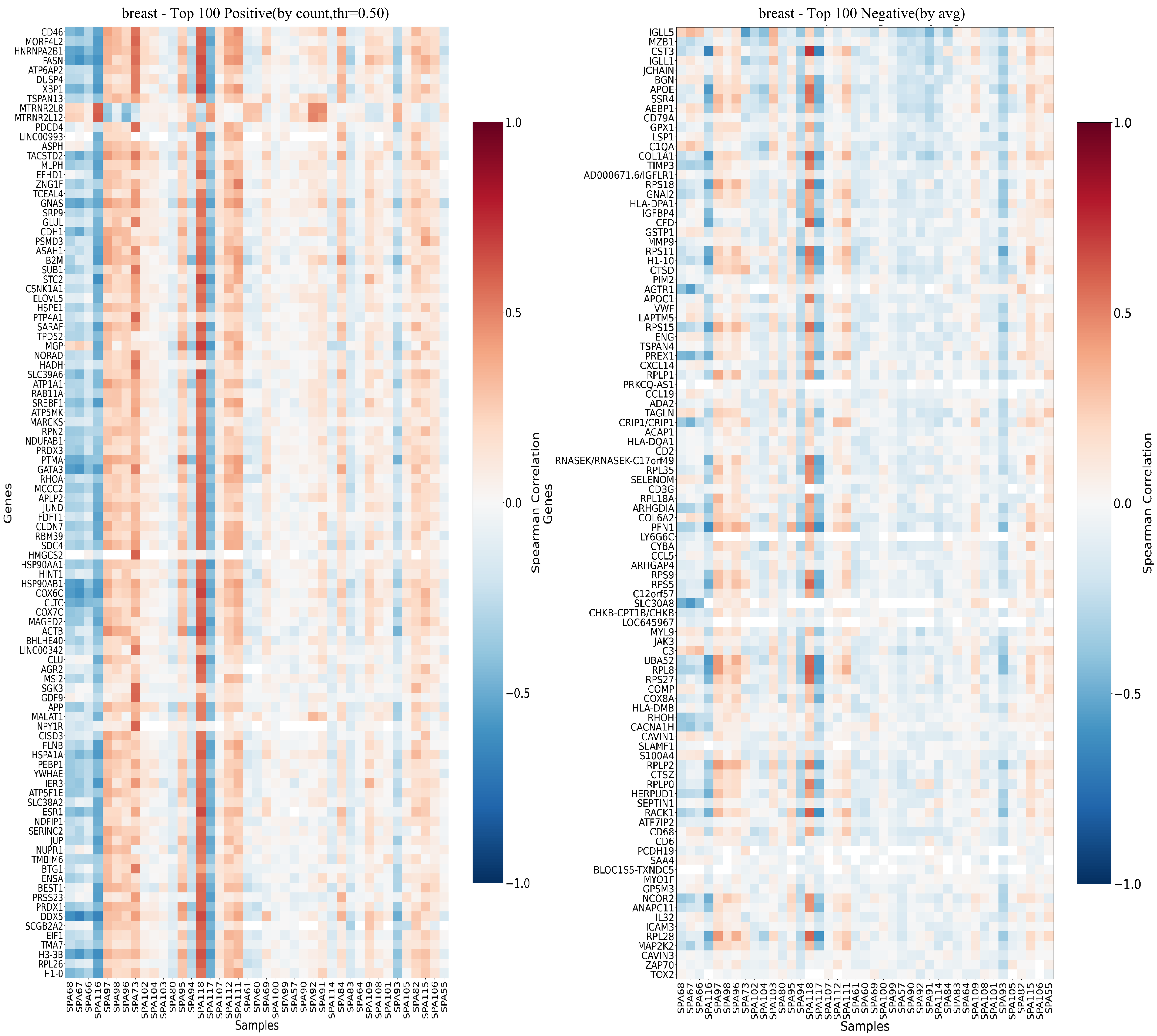

### Fig.S26

#### ResNet18

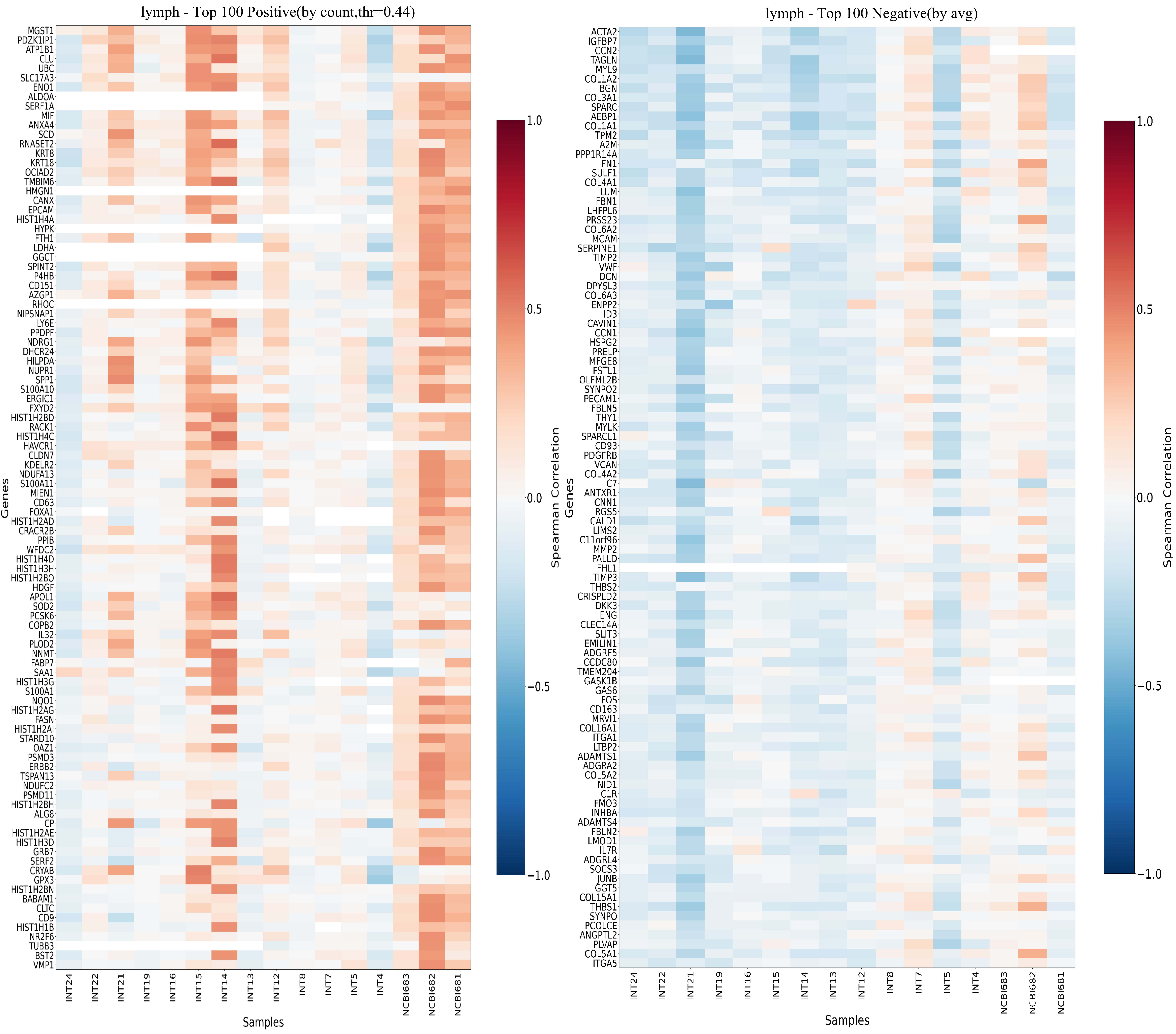

### Fig.S27

#### ViT-B/16

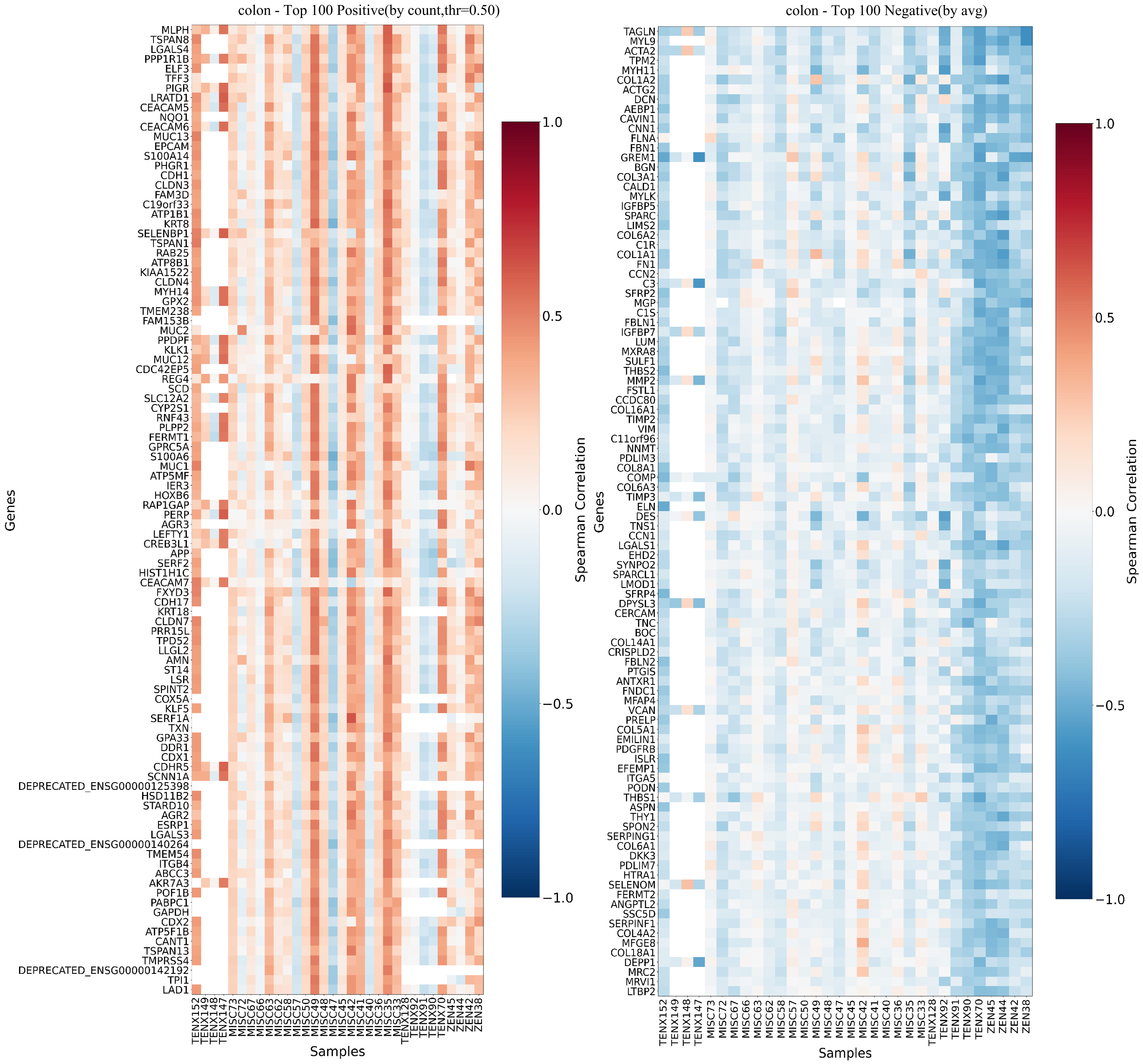

### Fig.S28

#### ViT-B/16

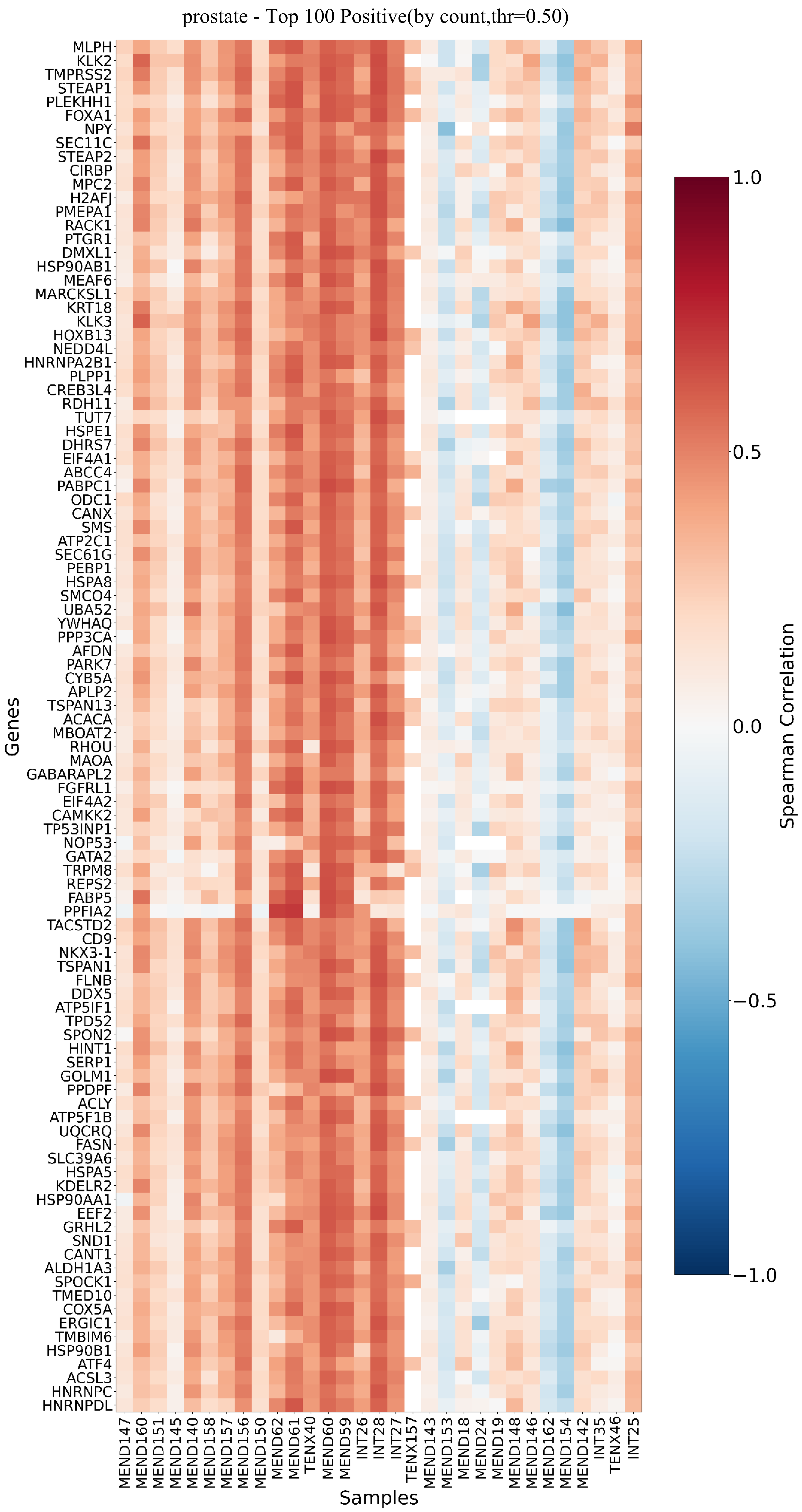


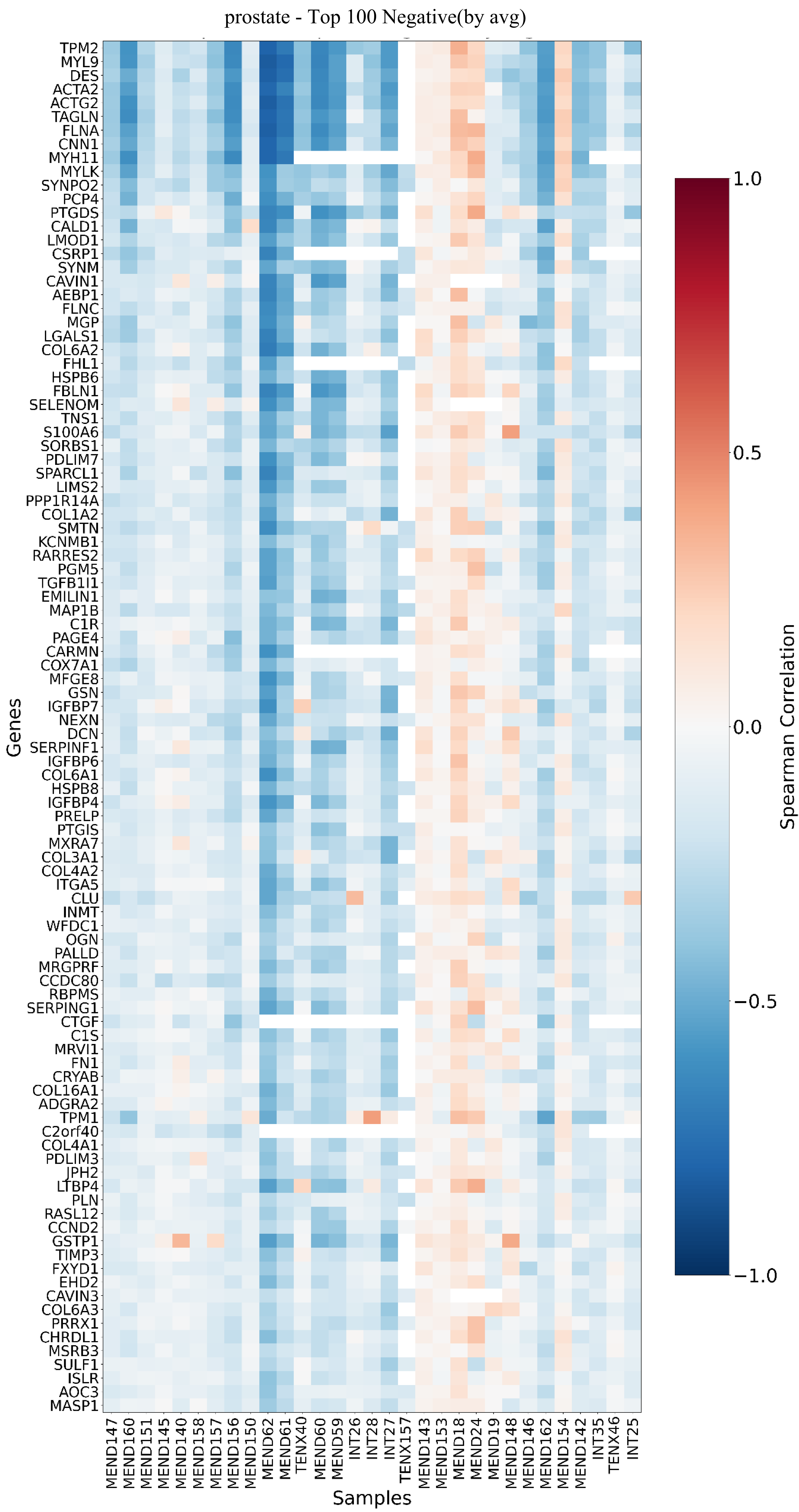

### Fig.S29

#### ViT-B/16

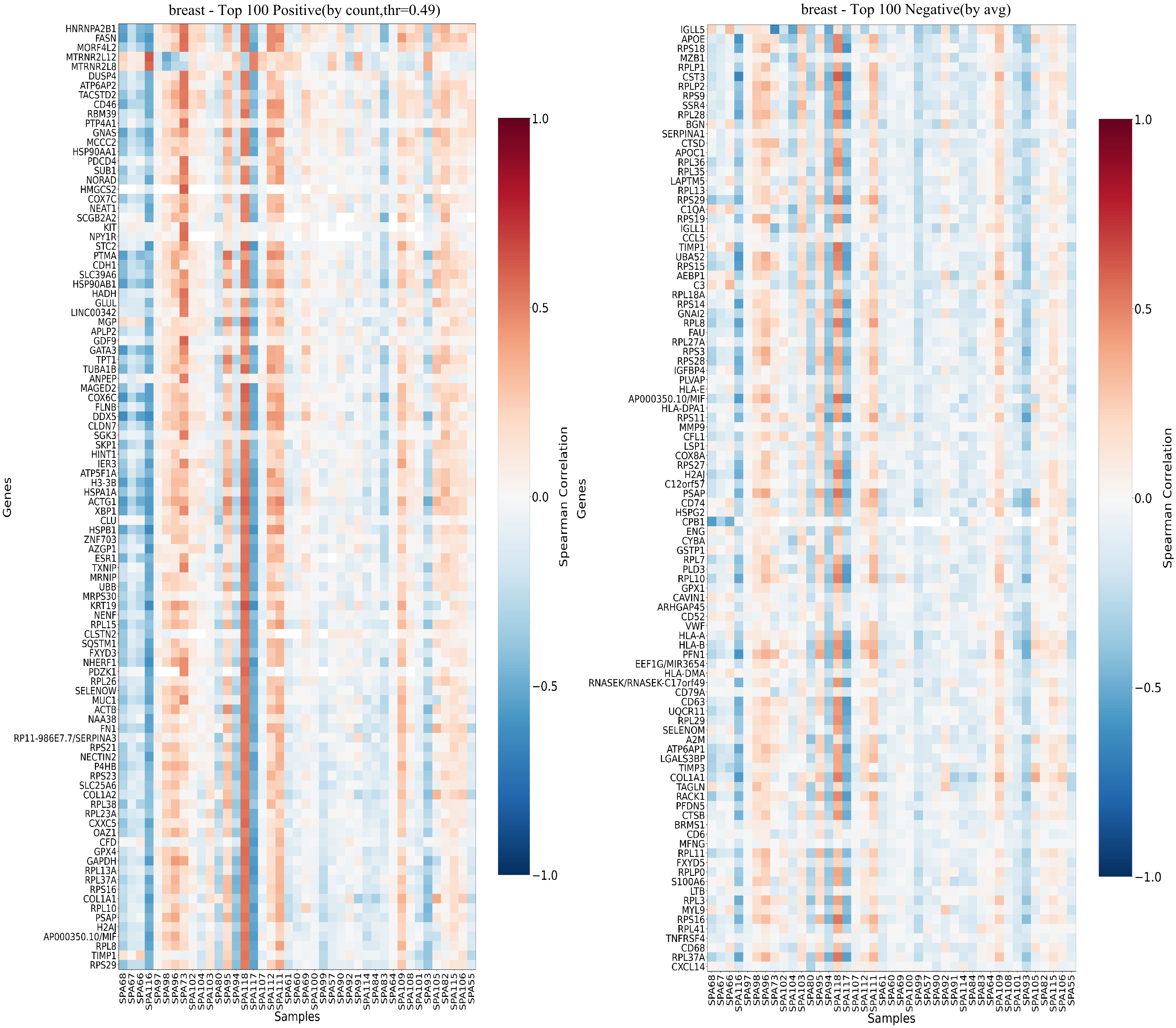

### Fig.S30

#### ViT-B/16

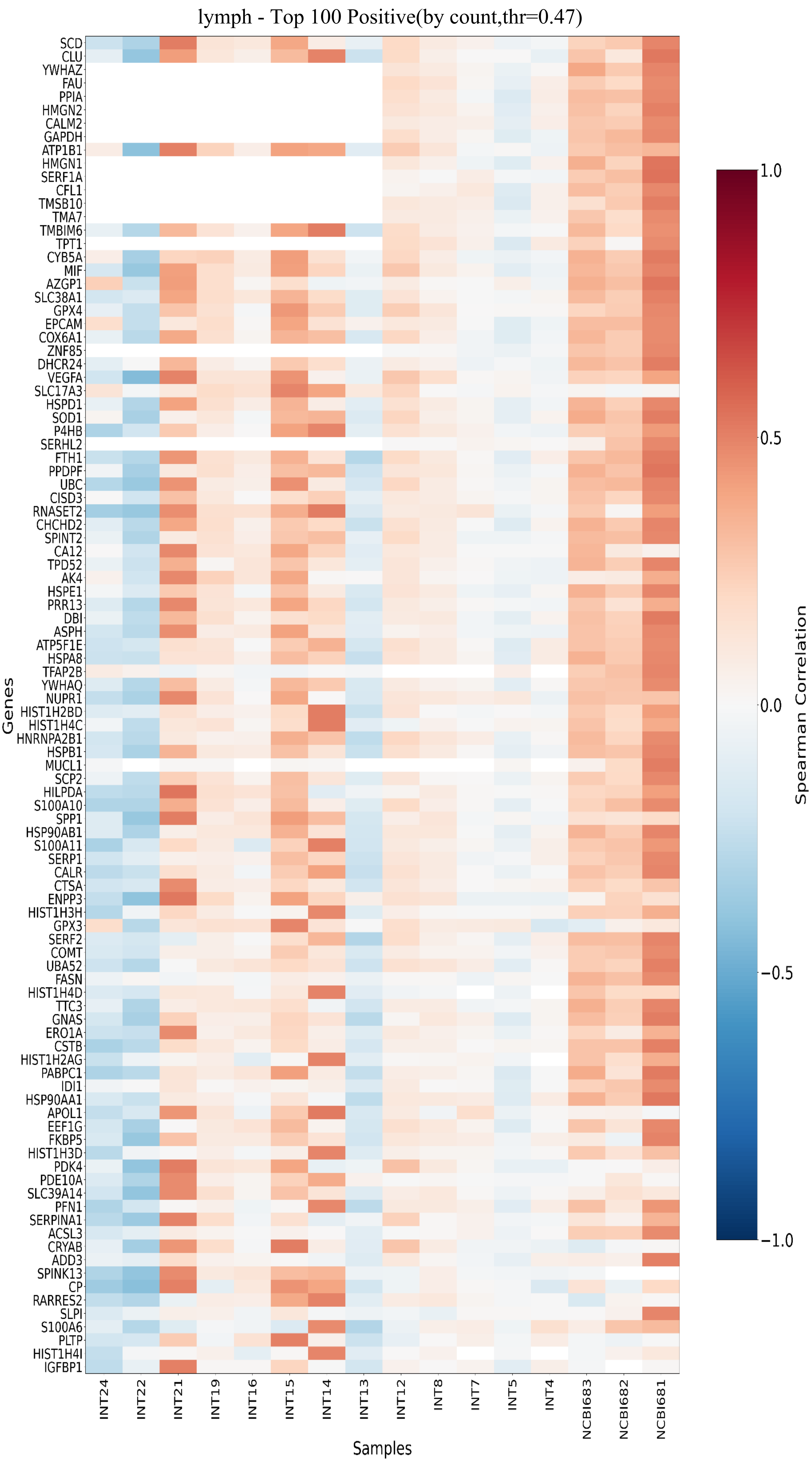


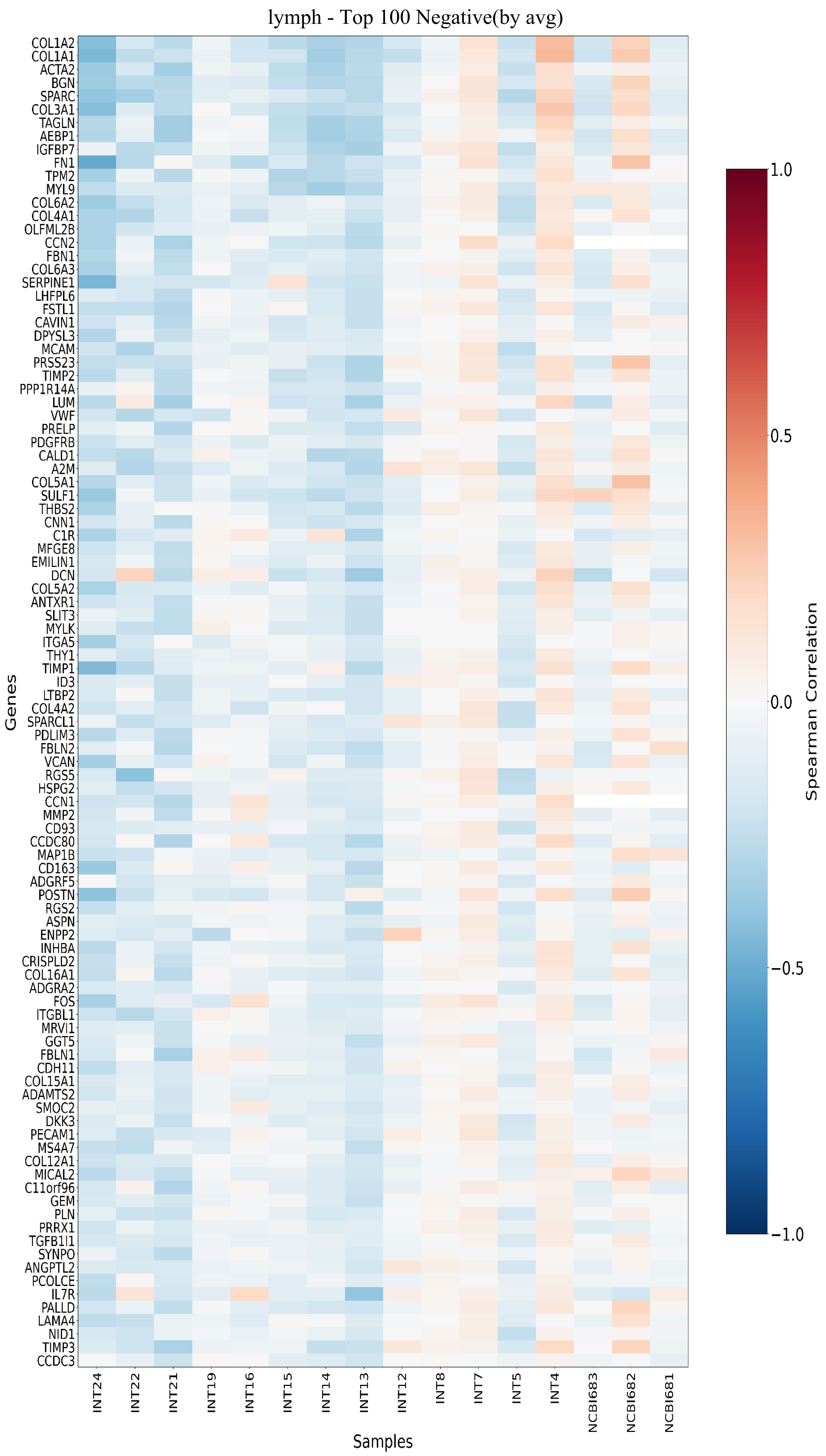

### Fig.S31

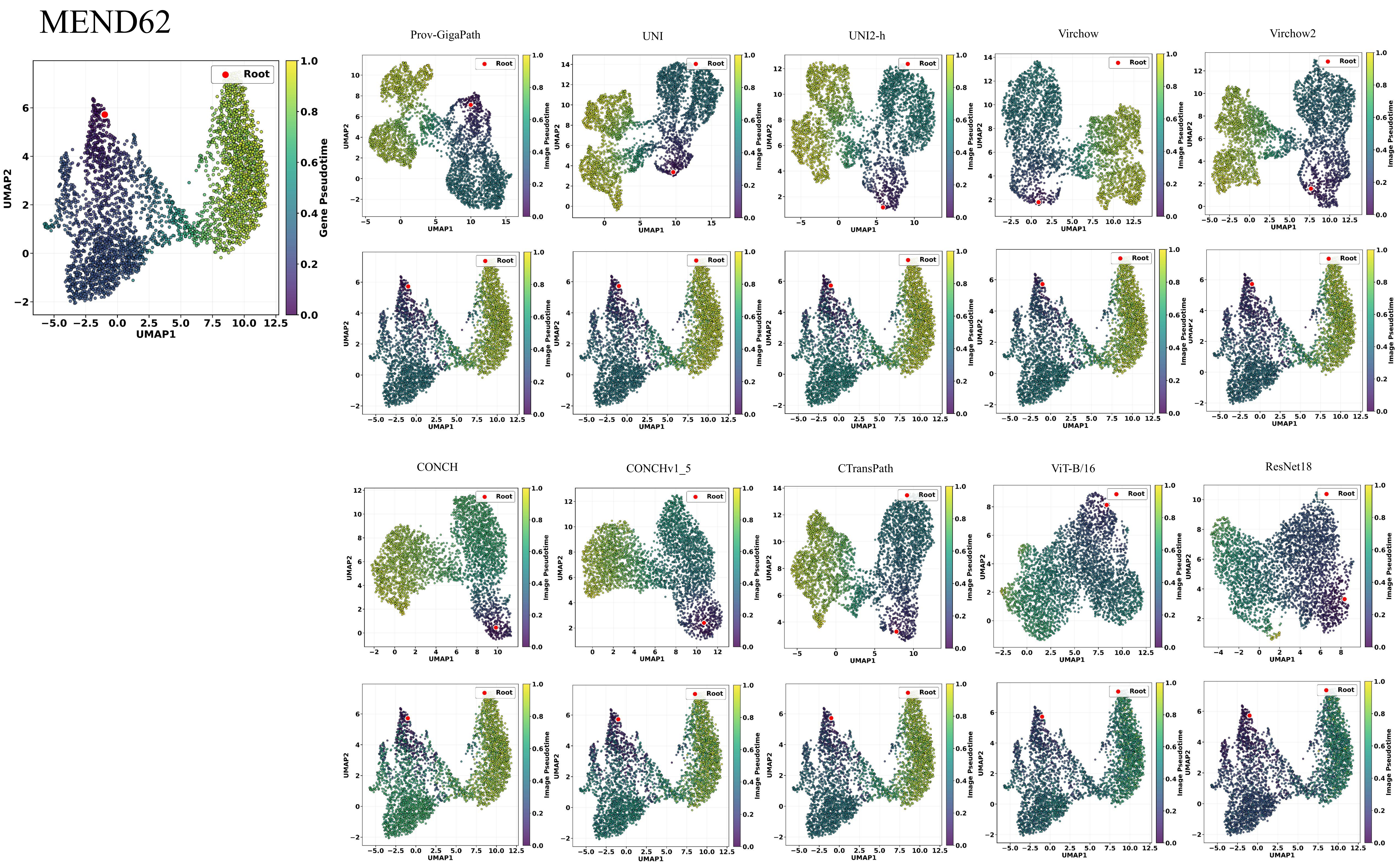
MEND62
Root
Gene Pseudotime
UMAP1
UMAP2
Prov-GigaPath
UNI
UNI2-h
Virchow
Virchow2
CONCH
CONCHv1_5
CTransPath
ViT-B/16
ResNet18
Image Pseudotime

Supplementary Tables Legend

Table S1. Correlations (Pearson) between model-derived histomorphological trajectories and spatial transcriptomic trajectories across all samples.

Table S2. Correlations (Kendall_tau) between model-derived histomorphological trajectories and spatial transcriptomic trajectories across all samples.

Table S3. Correlations (Spearman) between model-derived histomorphological trajectories and spatial transcriptomic trajectories across all samples.

Table S4. Correlations (Wasserstein_distance) between model-derived histomorphological trajectories and spatial transcriptomic trajectories across all samples.

Table S5. Correlations (Cosine_similarity) between model-derived histomorphological trajectories and spatial transcriptomic trajectories across all samples.

Table S6. The overlap of key genes enriched by different models (high-performance vs. low-performance) with spatial transcriptomics (ST)-enriched genes.

Table S7. The early and late trajectory genes enriched by trajectories derived from different models on INT4 samples.

Table S8. The samples used in this study.

### Table.S1

| pearson | UNI_big_region | UNI_small_region | CONCH_big_region | CONCH_small_region | CONCH15_big_region | CONCH15_small_region | CTransPath_big_region | CTransPath_small_region | GigaPath_big_region | GigaPath_small_region | UNI2h_big_region | UNI2h_small_region | RN18_big_region | RN18_small_region | Virchow_big_region | Virchow_small_region | Virchow2_big_region | Virchow2_small_region | ViTB16_big_region | ViTB16_small_region |
|---|---|---|---|---|---|---|---|---|---|---|---|---|---|---|---|---|---|---|---|---|
| INT13 | 0.4905 | 0.351 | 0.4385 | 0.4961 | 0.0748 | 0.0249 | 0.2625 | 0.431 | 0.4898 | 0.4599 | 0.5968 | 0.4447 | 0.216 | 0.2589 | 0.4617 | 0.3435 | 0.3771 | 0.527 | 0.302 | 0.3778 |
| INT14 | 0.8364 | 0.8672 | 0.7575 | 0.7575 | 0.8028 | 0.7311 | 0.8618 | 0.8176 | 0.8672 | 0.8448 | 0.8565 | 0.8372 | 0.7836 | 0.7275 | 0.8875 | 0.8827 | | 0.806 | 0.8206 | 0.7397 |
| INT15 | 0.8332 | 0.8381 | 0.6913 | 0.6913 | 0.7346 | 0.7083 | 0.8225 | 0.7 | 0.855 | 0.8451 | 0.8347 | 0.8311 | 0.5898 | 0.3013 | 0.8401 | 0.834 | 0.8187 | 0.8019 | 0.6305 | 0.5417 |
| INT16 | 0.7746 | 0.699 | 0.625 | 0.625 | 0.6576 | 0.5737 | 0.6826 | 0.3355 | 0.7558 | 0.6444 | 0.7817 | 0.6138 | 0.2577 | 0.248 | 0.8063 | 0.7489 | 0.7409 | 0.6131 | 0.2996 | 0.3752 |
| INT19 | 0.3613 | 0.3591 | 0.312 | 0.379 | 0.3625 | 0.2055 | 0.3189 | 0.3194 | 0.3487 | 0.3436 | 0.4102 | 0.3884 | 0.349 | 0.3474 | 0.373 | 0.3872 | 0.3719 | 0.3776 | 0.3463 | 0.3382 |
| INT21 | 0.8539 | 0.6319 | 0.6008 | 0.6008 | 0.813 | 0.6423 | 0.7978 | 0.2247 | 0.8289 | 0.7259 | 0.8405 | 0.7666 | 0.6003 | -0.029 | 0.8219 | 0.7411 | 0.8419 | 0.6105 | 0.6277 | 0.5092 |
| INT22 | 0.2314 | 0.3386 | 0.3356 | 0.3356 | 0.0181 | 0.228 | 0.0541 | 0.1921 | 0.1745 | 0.1332 | 0.3807 | 0.2323 | 0.3038 | 0.2017 | 0.2925 | 0.337 | 0.3103 | 0.023 | 0.0602 | 0.152 |
| INT24 | 0.8311 | 0.4413 | 0.0626 | 0.0625 | 0.6506 | 0.2459 | 0.463 | -0.3187 | 0.7763 | 0.4758 | 0.8314 | 0.7935 | -0.1728 | -0.2066 | 0.7208 | 0.6141 | 0.8399 | 0.4213 | 0.1307 | -0.1079 |
| INT25 | 0.5191 | 0.3761 | 0.3839 | 0.322 | 0.3755 | 0.2768 | 0.4209 | 0.3758 | 0.5734 | 0.4111 | 0.6251 | 0.4343 | 0.4251 | 0.3111 | 0.5668 | 0.2939 | 0.5474 | 0.3967 | 0.3946 | 0.4074 |
| INT26 | 0.4776 | 0.4804 | 0.4865 | 0.4869 | 0.5358 | 0.4279 | 0.6032 | 0.6127 | 0.6054 | 0.5448 | 0.4434 | 0.4731 | 0.5944 | 0.515 | 0.6326 | 0.326 | 0.5614 | 0.5082 | 0.5483 | 0.3213 |
| INT27 | 0.5501 | 0.7302 | 0.7374 | 0.7374 | 0.8026 | 0.7908 | 0.7433 | 0.7235 | 0.8131 | 0.8376 | 0.5208 | 0.7606 | 0.5952 | 0.5163 | 0.9091 | 0.6566 | 0.8333 | | 0.7153 | 0.7769 |
| INT28 | 0.7627 | 0.7617 | 0.7044 | 0.7044 | 0.7783 | 0.787 | 0.8536 | 0.7964 | 0.8562 | 0.8093 | 0.7747 | 0.6825 | 0.4916 | 0.5009 | 0.8598 | 0.7313 | 0.8036 | 0.7248 | 0.7236 | 0.6934 |
| INT35 | 0.47 | 0.6115 | 0.6047 | 0.6047 | 0.594 | 0.6496 | 0.4907 | 0.5139 | 0.5195 | 0.6982 | | 0.6486 | 0.4114 | 0.4784 | 0.5989 | 0.413 | 0.5988 | 0.641 | 0.4284 | 0.3892 |
| INT4 | 0.1723 | -0.2142 | -0.1356 | -0.1356 | -0.3005 | -0.3216 | -0.1145 | -0.0099 | 0.1473 | 0.0008 | -0.0259 | -0.2641 | -0.2797 | -0.2786 | -0.0772 | 0.0181 | -0.3052 | -0.1267 | -0.0377 | -0.1733 |
| INT5 | 0.0935 | -0.2235 | -0.1663 | -0.1663 | -0.0452 | -0.3149 | -0.1134 | -0.1144 | 0.0996 | 0.1013 | 0.283 | -0.1941 | 0.1744 | -0.2211 | 0.0796 | -0.0614 | 0.3156 | 0.0617 | 0.0646 | -0.0338 |
| INT7 | 0.214 | 0.1807 | 0.0269 | 0.0269 | 0.081 | -0.0473 | 0.1793 | 0.2061 | 0.1351 | 0.1618 | 0.1939 | 0.1423 | -0.0187 | -0.0835 | 0.1568 | 0.183 | 0.2346 | 0.2511 | 0.0833 | -0.0155 |
| INT8 | 0.0547 | -0.0209 | 0.0714 | 0.0714 | 0.0445 | 0.0735 | 0.1003 | 0.0148 | 0.067 | 0.0522 | 0.1202 | 0.0387 | 0.1018 | 0.0232 | 0.119 | 0.0792 | 0.0942 | 0.0144 | 0.0605 | 0.0336 |
| MEND140 | 0.4502 | 0.6763 | 0.5405 | 0.5405 | 0.3616 | 0.2261 | 0.2749 | 0.5448 | 0.5824 | 0.634 | 0.4512 | 0.6906 | 0.4156 | 0.5373 | 0.5727 | 0.6864 | 0.6275 | 0.6899 | 0.2601 | 0.0089 |
| MEND142 | 0.7103 | 0.7527 | 0.8106 | 0.8106 | 0.7773 | 0.7587 | 0.4151 | 0.6692 | 0.7066 | 0.788 | 0.7464 | 0.8165 | 0.1371 | 0.326 | 0.6467 | 0.6877 | 0.6367 | 0.7592 | 0.6259 | 0.4186 |
| MEND143 | 0.8232 | 0.8207 | 0.7017 | 0.7017 | 0.7403 | 0.5158 | 0.6765 | 0.8265 | 0.827 | 0.7669 | 0.7385 | 0.7986 | 0.1106 | 0.124 | 0.8067 | 0.7306 | 0.8089 | 0.737 | 0.216 | 0.0381 |
| MEND145 | 0.7568 | 0.6558 | 0.6842 | 0.6842 | 0.7676 | 0.4582 | 0.5078 | 0.6597 | 0.8105 | 0.6039 | 0.7592 | 0.6211 | 0.3868 | 0.2806 | 0.7641 | 0.5681 | 0.763 | 0.6172 | 0.2498 | 0.1895 |
| MEND146 | 0.5008 | 0.4758 | 0.2822 | 0.2822 | 0.5187 | 0.1866 | 0.2487 | 0.3988 | 0.5952 | 0.5143 | 0.567 | 0.5785 | 0.2506 | 0.1826 | 0.4881 | 0.3856 | 0.4544 | 0.499 | 0.266 | 0.2907 |
| MEND147 | 0.5438 | 0.4645 | 0.4311 | 0.4311 | 0.5377 | 0.4321 | 0.4932 | 0.4674 | 0.6785 | 0.5729 | 0.5306 | 0.391 | 0.2539 | 0.285 | 0.534 | 0.3974 | 0.5487 | 0.432 | 0.3164 | 0.179 |
| MEND150 | 0.6985 | 0.8634 | 0.8557 | 0.8557 | 0.5309 | 0.8463 | 0.6449 | 0.8301 | 0.8549 | 0.8727 | 0.6764 | 0.8607 | 0.5897 | 0.6266 | 0.8237 | 0.7905 | 0.7813 | 0.8663 | 0.2394 | 0.7173 |
| MEND151 | 0.4458 | 0.675 | 0.7187 | 0.6998 | 0.5609 | 0.5417 | 0.5123 | | 0.5878 | 0.6539 | 0.4875 | 0.6911 | 0.5311 | 0.5522 | 0.4152 | 0.5582 | 0.585 | 0.7058 | 0.2405 | 0.3622 |
| MEND153 | 0.2038 | 0.0041 | -0.039 | -0.039 | 0.0298 | -0.3557 | 0.5414 | 0.0516 | 0.3626 | 0.002 | 0.3166 | -0.0133 | 0.264 | 0.131 | 0.1258 | -0.1193 | -0.0166 | -0.1849 | 0.2176 | 0.1137 |
| MEND154 | -0.0612 | 0.3486 | 0.7218 | 0.4888 | -0.1209 | 0.3608 | 0.0719 | 0.4778 | 0.1668 | 0.5109 | 0.0862 | 0.2299 | -0.1591 | 0.3477 | 0.25 | 0.5003 | 0.0053 | 0.5071 | -0.0413 | 0.3414 |
| MEND157 | 0.6906 | 0.7341 | 0.6241 | 0.7479 | 0.6804 | 0.697 | 0.6241 | 0.6377 | 0.7261 | 0.7477 | 0.6872 | 0.7439 | 0.6475 | 0.6068 | 0.733 | 0.7117 | 0.6913 | 0.6144 | 0.4984 | 0.6744 |
| MEND158 | 0.4973 | 0.6981 | -0.1932 | 0.6468 | 0.4359 | 0.3707 | 0.5325 | 0.6647 | 0.7003 | 0.6957 | 0.3975 | 0.6795 | 0.3085 | 0.4014 | 0.3682 | 0.518 | 0.6989 | 0.5593 | 0.0466 | 0.3979 |
| MEND160 | 0.734 | 0.8212 | 0.7804 | 0.7718 | 0.7555 | 0.6614 | 0.7472 | 0.8012 | 0.778 | 0.8085 | 0.7146 | 0.7684 | 0.7169 | 0.6213 | 0.7971 | 0.7706 | 0.7142 | 0.726 | 0.6781 | 0.6534 |
| MEND162 | 0.6268 | 0.3568 | 0.2422 | 0.2422 | 0.4716 | 0.2215 | 0.3857 | 0.2825 | 0.439 | 0.1868 | 0.6588 | 0.5189 | 0.0974 | 0.2985 | 0.5053 | 0.2185 | 0.3566 | 0.2783 | 0.1369 | 0.1559 |
| MEND18 | -0.0061 | 0.0996 | 0.151 | 0.151 | -0.1361 | -0.0917 | -0.0125 | 0.011 | 0.1576 | 0.1943 | 0.2264 | 0.2463 | 0.013 | 0.1471 | 0.0516 | 0.1232 | 0.1948 | 0.2433 | -0.0499 | 0.0396 |
| MEND19 | 0.7453 | 0.732 | 0.6828 | 0.6828 | 0.6864 | 0.6713 | 0.6884 | 0.7116 | 0.7507 | 0.7366 | 0.6941 | 0.6818 | 0.374 | 0.4608 | 0.6422 | 0.6 | 0.6508 | 0.6881 | 0.4116 | 0.459 |
| MEND24 | -0.2302 | -0.2012 | 0.1306 | 0.2086 | -0.3479 | -0.2443 | -0.1453 | -0.1823 | -0.1578 | -0.0885 | -0.343 | -0.2838 | -0.1149 | 0.1283 | 0.0118 | 0.0726 | 0.1919 | 0.2655 | -0.1891 | -0.1671 |
| MEND37 | 0.6194 | 0.6575 | 0.595 | 0.595 | 0.5717 | 0.5709 | 0.5114 | 0.572 | 0.5665 | 0.7239 | 0.52 | 0.7531 | 0.2967 | 0.3753 | 0.4912 | 0.4512 | 0.6024 | 0.6305 | 0.4701 | 0.3902 |
| MEND38 | 0.569 | 0.5947 | 0.4805 | 0.4805 | 0.4885 | 0.4019 | 0.5082 | 0.5485 | 0.4194 | 0.4828 | 0.4079 | 0.4969 | 0.1022 | 0.2703 | 0.3464 | 0.3756 | 0.5142 | 0.5946 | 0.2944 | 0.1859 |
| MEND39 | 0.7244 | 0.6918 | 0.6092 | 0.653 | 0.7032 | 0.6633 | 0.6014 | 0.6404 | 0.6198 | 0.6965 | 0.6632 | 0.691 | 0.402 | 0.3222 | 0.5955 | 0.5362 | 0.6867 | 0.754 | 0.5099 | 0.3176 |
| MEND40 | 0.6236 | 0.6458 | 0.5509 | 0.5508 | 0.6208 | 0.5739 | 0.5462 | 0.6073 | 0.5844 | 0.5997 | 0.5408 | 0.6913 | 0.3224 | 0.407 | 0.5953 | 0.394 | 0.6349 | 0.6475 | 0.5866 | 0.4653 |
| MEND59 | 0.8945 | 0.4785 | 0.1604 | 0.1604 | 0.672 | 0.2161 | 0.8455 | 0.3679 | 0.8545 | 0.4507 | 0.91 | 0.34 | 0.7016 | 0.3916 | | 0.4563 | 0.8394 | 0.1552 | 0.6612 | 0.3666 |
| MEND60 | 0.8991 | 0.796 | 0.6509 | 0.6509 | 0.8251 | 0.6801 | 0.8967 | 0.822 | 0.8825 | 0.7782 | 0.9085 | 0.7553 | 0.7557 | 0.7318 | 0.8856 | 0.6484 | 0.8839 | 0.7013 | 0.7371 | 0.3323 |
| MEND61 | 0.9632 | 0.8163 | 0.6022 | 0.6022 | 0.8126 | 0.6324 | 0.9472 | 0.7357 | 0.9592 | 0.6268 | 0.9557 | 0.8014 | 0.814 | 0.6238 | 0.9501 | 0.7772 | 0.8845 | 0.5121 | 0.7853 | 0.511 |
| MEND62 | 0.937 | 0.8775 | 0.7008 | 0.7008 | 0.8452 | 0.7486 | 0.9349 | 0.8407 | 0.9389 | 0.8209 | 0.9083 | 0.7869 | 0.8495 | 0.7299 | 0.9276 | 0.8125 | 0.9334 | 0.8178 | 0.7685 | 0.6992 |
| MEND66 | 0.0174 | 0.0646 | -0.0144 | -0.0144 | 0.0027 | 0.0357 | 0.0061 | 0.1196 | 0.0544 | 0.075 | -0.0157 | 0.0503 | 0.0597 | 0.1067 | 0.1963 | 0.2165 | 0.1066 | 0.1047 | 0.0987 | 0.0837 |
| MISC33 | 0.8118 | | 0.6218 | 0.6527 | 0.4815 | 0.0211 | 0.6638 | 0.4251 | 0.7715 | 0.6298 | 0.8046 | 0.7623 | 0.0765 | -0.4443 | 0.7912 | 0.6399 | 0.8175 | 0.7458 | 0.4994 | -0.3167 |
| MISC35 | 0.8407 | 0.7886 | 0.6194 | 0.6194 | 0.8079 | 0.6025 | 0.8011 | 0.4175 | 0.8402 | 0.7704 | 0.8383 | 0.7964 | 0.1465 | 0.1291 | 0.8434 | 0.6261 | | 0.676 | 0.598 | -0.0084 |
| MISC36 | 0.8608 | 0.7868 | 0.6311 | 0.6311 | 0.8665 | 0.6744 | 0.7249 | 0.6135 | 0.8632 | 0.7518 | 0.8679 | 0.7771 | 0.3579 | 0.334 | 0.8253 | 0.7655 | 0.6201 | 0.2531 | 0.4607 | 0.2815 |
| MISC40 | 0.5126 | 0.1357 | 0.5397 | 0.5397 | 0.5618 | 0.5668 | 0.2555 | 0.3177 | 0.5169 | 0.4806 | 0.4745 | 0.407 | 0.4171 | 0.073 | 0.5279 | 0.3237 | 0.4873 | 0.3685 | 0.0101 | 0.2777 |
| MISC41 | 0.7068 | 0.4552 | 0.5303 | 0.5303 | 0.6204 | 0.5278 | 0.6084 | 0.0856 | 0.7513 | 0.1756 | 0.7362 | 0.6622 | 0.1322 | 0.0736 | 0.77 | 0.1768 | | 0.6577 | 0.2994 | 0.239 |
| MISC45 | 0.4585 | 0.3006 | 0.2458 | 0.2458 | 0.3869 | 0.2568 | 0.3179 | 0.2304 | 0.5278 | 0.3205 | 0.5217 | 0.4771 | -0.0944 | -0.0736 | 0.5736 | 0.3788 | 0.533 | 0.3934 | 0.1069 | 0.0484 |
| MISC47 | 0.5058 | 0.1549 | 0.0421 | 0.0421 | 0.2932 | -0.0334 | 0.4382 | 0.0502 | 0.5875 | 0.1503 | 0.51 | 0.3665 | 0.123 | 0.0718 | 0.54 | 0.2393 | 0.3827 | 0.0832 | 0.1422 | 0.0186 |
| MISC48 | 0.6799 | 0.6761 | 0.5564 | 0.5564 | 0.6052 | 0.4997 | 0.6772 | 0.669 | 0.6926 | 0.6014 | 0.6426 | 0.6368 | 0.5214 | 0.4076 | 0.6 | 0.5491 | 0.6489 | 0.5637 | 0.5291 | 0.4731 |
| MISC49 | 0.6858 | 0.6425 | 0.5361 | 0.5361 | 0.5947 | 0.4962 | 0.6553 | 0.5431 | 0.7307 | 0.5143 | 0.7068 | 0.6744 | 0.0757 | -0.2555 | 0.6684 | 0.5801 | 0.7172 | 0.6693 | 0.3985 | 0.3953 |
| MISC50 | 0.6779 | 0.6273 | 0.4786 | 0.4786 | 0.6201 | 0.5429 | 0.6364 | 0.418 | 0.7028 | 0.5588 | 0.7055 | 0.6856 | 0.3499 | 0.1485 | 0.656 | 0.3361 | 0.6476 | 0.5131 | 0.287 | 0.226 |
| MISC62 | 0.5853 | 0.1097 | 0.0013 | 0.4082 | 0.5104 | 0.377 | 0.4975 | 0.3067 | 0.6476 | 0.3068 | 0.6614 | 0.457 | 0.3335 | 0.2441 | 0.637 | 0.2524 | 0.6455 | 0.3378 | 0.2174 | 0.2684 |
| MISC63 | -0.0631 | -0.0735 | -0.0154 | -0.0153 | -0.0354 | -0.0502 | -0.0518 | -0.069 | 0.0303 | -0.0099 | -0.04 | -0.0234 | -0.0056 | -0.0104 | 0.0549 | 0.0255 | 0.0116 | -0.0025 | -0.0907 | -0.0778 |
| MISC66 | 0.8362 | 0.2812 | 0.157 | 0.1536 | 0.5509 | -0.0359 | 0.2675 | 0.0184 | 0.7342 | 0.0965 | 0.9179 | 0.582 | -0.2701 | -0.1869 | 0.6053 | 0.0401 | 0.8713 | 0.3123 | -0.1398 | -0.0609 |
| MISC67 | 0.8979 | 0.5294 | 0.352 | 0.352 | 0.603 | 0.334 | 0.4484 | 0.3353 | 0.8123 | 0.4227 | 0.8923 | 0.7539 | -0.094 | 0.0491 | 0.651 | 0.5302 | 0.9085 | 0.6141 | 0.201 | -0.0116 |
| MISC72 | 0.6146 | 0.5457 | 0.5162 | 0.5162 | 0.6241 | 0.5325 | 0.6336 | 0.4873 | 0.6288 | 0.5505 | 0.5798 | 0.59 | 0.3354 | 0.1734 | 0.6046 | 0.4798 | 0.5995 | 0.5652 | 0.3301 | 0.2578 |
| MISC73 | 0.7901 | 0.6787 | 0.6833 | 0.6833 | 0.6804 | 0.6166 | 0.7413 | 0.4346 | 0.6948 | 0.5791 | 0.7848 | 0.7602 | 0.3835 | 0.283 | 0.7606 | 0.5903 | 0.7929 | 0.7353 | 0.0737 | 0.1406 |
| NCBI629 | 0.1944 | 0.1758 | 0.1346 | 0.1346 | 0.0683 | 0.1793 | 0.036 | 0.1796 | 0.1445 | 0.1688 | 0.1614 | 0.1355 | 0.0905 | 0.1534 | 0.1973 | 0.1718 | 0.2032 | 0.1861 | 0.2054 | 0.1362 |
| NCBI631 | 0.0919 | -0.0519 | 0.1848 | 0.1518 | 0.0348 | 0.0072 | 0.0243 | -0.0356 | -0.0326 | -0.0563 | -0.009 | 0.0072 | 0.1153 | -0.1243 | 0.0808 | 0.0208 | -0.0391 | -0.0767 | -0.0996 | 0.0152 |
| NCBI632 | 0.385 | 0.3376 | 0.3934 | 0.3934 | 0.4024 | 0.3993 | 0.3629 | 0.4146 | 0.4323 | 0.3867 | 0.4289 | 0.2536 | 0.2511 | 0.2809 | 0.4264 | 0.3108 | 0.4108 | 0.4037 | 0.389 | 0.4076 |
| NCBI635 | 0.3176 | 0.3175 | 0.2589 | 0.2589 | 0.3455 | 0.285 | 0.3303 | 0.3425 | 0.312 | 0.3308 | 0.2964 | 0.3106 | 0.3137 | 0.3937 | 0.3315 | 0.3126 | 0.3217 | 0.3983 | 0.23 | 0.3259 |
| NCBI638 | -0.0882 | -0.0375 | -0.1164 | -0.1164 | 0.0081 | -0.0682 | -0.0702 | -0.1281 | -0.0619 | -0.0397 | -0.158 | -0.0066 | 0.0815 | 0.0186 | -0.0012 | 0.0735 | -0.1425 | -0.0413 | -0.0448 | -0.0232 |
| NCBI640 | 0.2398 | 0.1497 | 0.1353 | 0.1353 | 0.1573 | 0.2001 | 0.0257 | 0.08 | 0.2171 | 0.2009 | 0.348 | 0.3004 | 0.0898 | 0.0615 | 0.2694 | 0.1536 | 0.2917 | 0.1487 | 0.2555 | 0.2502 |
| NCBI681 | 0.79 | 0.7509 | 0.6673 | 0.6673 | 0.6899 | | 0.8028 | 0.5559 | 0.7816 | 0.7708 | 0.7728 | 0.7712 | 0.4493 | 0.2578 | 0.7885 | 0.6461 | 0.8421 | 0.6438 | 0.5709 | 0.4108 |
| NCBI682 | 0.6119 | 0.7103 | 0.4657 | 0.7008 | 0.5802 | 0.4575 | 0.6612 | 0.6603 | 0.7069 | 0.7024 | 0.6601 | 0.6534 | 0.4095 | 0.35 | 0.5793 | 0.5871 | 0.6447 | 0.6406 | 0.486 | 0.5394 |
| NCBI683 | 0.5412 | 0.5436 | 0.5416 | 0.5416 | 0.5635 | 0.5121 | 0.5592 | 0.5064 | 0.5524 | 0.5306 | 0.5417 | 0.5381 | 0.4845 | 0.4285 | 0.5521 | 0.5525 | 0.5802 | 0.5899 | 0.5276 | 0.4844 |
| NCBI760 | 0.3717 | 0.3749 | 0.3651 | 0.3651 | 0.3908 | 0.3893 | 0.3855 | 0.4026 | 0.4158 | 0.415 | 0.3725 | 0.3857 | 0.3851 | 0.3849 | 0.3828 | 0.4047 | 0.3027 | 0.2993 | 0.3265 | 0.2658 |
| NCBI764 | 0.2323 | 0.1949 | 0.1937 | 0.1937 | 0.1469 | 0.1622 | 0.1708 | 0.1562 | 0.2413 | 0.2058 | 0.2247 | 0.206 | 0.1296 | 0.1099 | 0.2004 | 0.186 | 0.2247 | 0.1588 | 0.0434 | -0.0266 |
| NCBI765 | 0.5045 | 0.4914 | 0.5457 | 0.5457 | 0.484 | 0.4948 | 0.552 | 0.556 | 0.5205 | 0.5341 | 0.509 | 0.5224 | 0.451 | 0.4308 | 0.5381 | 0.5323 | 0.4508 | 0.5549 | 0.425 | 0.4778 |
| NCBI766 | 0.4558 | 0.444 | 0.4514 | 0.4514 | 0.448 | 0.4489 | 0.4044 | 0.3588 | 0.3704 | 0.3907 | 0.442 | 0.4107 | 0.4413 | 0.4477 | 0.4641 | 0.4551 | 0.2525 | 0.425 | 0.4677 | 0.4315 |
| NCBI768 | 0.5423 | 0.5439 | 0.6093 | 0.6093 | 0.6552 | 0.6566 | 0.6053 | 0.5685 | 0.5994 | 0.5763 | 0.605 | 0.5675 | 0.5075 | 0.5136 | 0.4975 | 0.5242 | 0.685 | 0.6092 | 0.537 | 0.5521 |
| NCBI769 | 0.5001 | | 0.6326 | 0.6326 | 0.5566 | 0.6822 | 0.5838 | 0.558 | 0.598 | 0.548 | 0.4609 | 0.4686 | 0.5031 | 0.4983 | 0.5782 | 0.5433 | 0.6306 | 0.6184 | 0.4426 | 0.5332 |
| NCBI770 | -0.1996 | -0.0904 | -0.2631 | -0.2631 | -0.0994 | -0.0227 | -0.2183 | -0.1633 | 0.0678 | -0.159 | -0.1167 | -0.1483 | -0.2312 | -0.1781 | -0.2551 | -0.2375 | -0.1477 | -0.1666 | -0.2004 | -0.1857 |
| SPA100 | 0.4046 | 0.5788 | -0.1386 | -0.1386 | 0.2951 | 0.3088 | 0.3931 | 0.3295 | 0.6149 | 0.6731 | 0.5761 | 0.5571 | 0.0277 | -0.0143 | 0.5912 | 0.6114 | 0.5791 | 0.5755 | 0.0827 | 0.1206 |
| SPA101 | 0.4192 | 0.373 | 0.3731 | 0.3731 | 0.0927 | 0.232 | 0.366 | 0.422 | 0.388 | 0.3457 | 0.4524 | 0.4297 | 0.3041 | 0.3426 | 0.4294 | 0.4352 | 0.358 | 0.4416 | 0.2902 | 0.3061 |
| SPA102 | 0.4029 | 0.4161 | 0.3377 | 0.3377 | 0.3687 | 0.3479 | 0.4654 | 0.4571 | 0.0467 | 0.1713 | 0.399 | 0.4424 | 0.35 | 0.3297 | 0.4013 | 0.4199 | 0.3591 | 0.3973 | 0.3098 | |
| SPA103 | -0.2796 | -0.3661 | -0.3052 | -0.3052 | -0.304 | -0.3449 | -0.3816 | -0.4169 | -0.4081 | -0.4358 | -0.3717 | -0.3497 | -0.0402 | -0.2022 | -0.3473 | -0.3782 | -0.3393 | -0.3663 | -0.0249 | 0.0196 |
| SPA104 | 0.4311 | 0.4416 | 0.3084 | 0.3084 | 0.399 | 0.3797 | 0.4362 | 0.3734 | 0.4496 | 0.447 | 0.4518 | 0.4109 | 0.4111 | 0.3607 | 0.4503 | 0.4263 | 0.3995 | 0.3701 | 0.4076 | 0.3786 |
| SPA105 | 0.2124 | 0.175 | 0.0889 | 0.0889 | 0.0008 | 0.085 | 0.1213 | 0.1142 | 0.0932 | 0.083 | 0.0515 | 0.0453 | 0.1736 | 0.1186 | 0.125 | 0.1232 | 0.2349 | 0.211 | 0.0999 | 0.0584 |
| SPA106 | 0.1325 | 0.1657 | 0.1303 | 0.1303 | 0.1468 | 0.0657 | 0.0729 | 0.0991 | | 0.016 | 0.0478 | 0.0739 | 0.0724 | 0.0657 | 0.0274 | 0.0176 | 0.1152 | 0.1521 | 0.0219 | 0.0207 |
| SPA107 | 0.7098 | 0.6764 | 0.5726 | 0.5726 | 0.4695 | 0.5104 | 0.3541 | 0.5386 | 0.5688 | 0.6175 | 0.3637 | 0.4545 | 0.1253 | 0.1906 | 0.3653 | 0.525 | 0.6716 | 0.6722 | -0.1144 | 0.0801 |
| SPA108 | 0.6753 | 0.6554 | 0.6498 | 0.6498 | 0.6232 | 0.5517 | 0.6477 | 0.6573 | 0.6538 | 0.6566 | 0.6622 | 0.6504 | 0.6131 | 0.5628 | 0.6466 | 0.6226 | 0.6295 | 0.623 | 0.6246 | 0.6288 |
| SPA109 | 0.2514 | 0.2353 | 0.2611 | 0.2611 | 0.2159 | 0.1845 | 0.2203 | 0.2043 | 0.2038 | 0.2038 | | 0.2245 | 0.2132 | 0.2056 | 0.2246 | 0.2303 | 0.2126 | 0.2254 | 0.1608 | 0.1666 |
| SPA111 | 0.6582 | 0.6552 | 0.5817 | 0.5817 | 0.4654 | 0.4083 | 0.6098 | 0.6245 | 0.6541 | 0.6266 | 0.5427 | 0.5728 | 0.5139 | 0.5148 | 0.5469 | 0.4544 | 0.4807 | 0.5533 | 0.5488 | 0.5753 |
| SPA112 | 0.6808 | 0.6572 | 0.6061 | 0.6061 | 0.433 | 0.4638 | 0.6116 | 0.5915 | 0.6234 | 0.6057 | 0.6049 | 0.5805 | 0.45 | 0.4388 | 0.5395 | 0.3558 | 0.4203 | 0.5656 | 0.6175 | 0.5722 |
| SPA114 | 0.1448 | 0.047 | -0.0393 | -0.0393 | -0.038 | 0.0877 | -0.0509 | -0.0707 | -0.041 | 0.0136 | -0.1296 | -0.0382 | -0.0226 | 0.1027 | 0.0715 | 0.0546 | 0.1194 | 0.0368 | -0.0867 | -0.1462 |
| SPA115 | 0.1051 | 0.096 | 0.1169 | 0.1169 | 0.0906 | 0.0281 | 0.0756 | 0.0824 | 0.0388 | 0.048 | 0.0655 | 0.055 | 0.154 | 0.1349 | 0.0748 | 0.0693 | 0.0978 | 0.0743 | 0.1158 | 0.1278 |
| SPA116 | 0.5123 | 0.3561 | 0.1917 | 0.1917 | 0.1741 | -0.477 | -0.5197 | -0.429 | 0.547 | 0.6178 | -0.1159 | -0.256 | -0.6509 | -0.6645 | 0.9123 | 0.8889 | 0.8588 | 0.827 | -0.5516 | -0.4123 |
| SPA117 | -0.0688 | 0.0648 | 0.5675 | 0.5675 | 0.403 | 0.2184 | -0.3082 | -0.1602 | 0.3379 | 0.118 | -0.2689 | -0.245 | -0.2561 | -0.3838 | -0.0525 | -0.0407 | 0.7525 | 0.7086 | -0.304 | -0.2726 |
| SPA118 | 0.4907 | 0.4625 | 0.5087 | 0.5087 | 0.3282 | 0.3768 | 0.4441 | 0.4445 | 0.4658 | 0.4683 | 0.489 | 0.4522 | 0.4661 | 0.4929 | 0.496 | 0.5063 | 0.5527 | 0.5107 | 0.3848 | 0.4372 |
| SPA55 | 0.3528 | 0.2705 | 0.0716 | 0.0716 | 0.3702 | 0.314 | 0.3804 | 0.3705 | 0.22 | 0.0307 | 0.3387 | 0.223 | -0.0638 | -0.1494 | 0.4254 | 0.4336 | 0.2269 | 0.0676 | 0.3651 | 0.3036 |
| SPA57 | 0.3982 | 0.4651 | 0.2981 | 0.2981 | 0.3293 | 0.2787 | 0.4358 | 0.3881 | 0.4799 | 0.4555 | 0.423 | 0.4432 | 0.2187 | 0.1704 | 0.4 | 0.4329 | 0.3067 | 0.0772 | 0.1937 | 0.1039 |
| SPA60 | 0.4109 | 0.5733 | 0.3331 | 0.3331 | -0.1056 | 0.1224 | 0.481 | 0.5244 | 0.5127 | 0.4949 | 0.3247 | 0.4942 | 0.097 | 0.0561 | 0.3719 | 0.0608 | 0.5136 | 0.5866 | 0.0529 | 0.0281 |
| SPA61 | 0.6919 | 0.6296 | 0.3269 | 0.3269 | 0.0918 | 0.2482 | 0.4936 | 0.5422 | 0.5498 | 0.5755 | 0.3516 | 0.4662 | 0.2225 | 0.1803 | 0.0279 | 0.0754 | 0.7269 | 0.7459 | 0.2363 | 0.3112 |
| SPA64 | 0.6573 | 0.6699 | 0.5243 | 0.5243 | -0.357 | -0.2116 | -0.1401 | -0.1914 | 0.6497 | 0.6448 | 0.6865 | 0.6801 | -0.0892 | -0.2108 | 0.6265 | 0.5856 | 0.2026 | 0.5202 | -0.309 | 0.0511 |
| SPA66 | 0.564 | 0.4687 | -0.434 | -0.434 | -0.3864 | -0.4801 | -0.0253 | -0.3528 | 0.4358 | 0.429 | 0.6105 | 0.504 | -0.3846 | -0.2932 | -0.0441 | -0.3285 | 0.3401 | 0.066 | -0.3688 | -0.298 |
| SPA67 | 0.7629 | 0.5921 | 0.2713 | 0.2713 | -0.3697 | -0.3444 | -0.3296 | -0.3236 | 0.7276 | 0.5705 | 0.7587 | 0.6648 | -0.2812 | -0.2337 | 0.5126 | 0.46 | -0.2151 | -0.2115 | -0.0174 | 0.0055 |
| SPA68 | 0.7791 | 0.7801 | 0.1909 | 0.1909 | -0.044 | -0.1587 | 0.1552 | 0.1792 | 0.7169 | 0.6848 | 0.7218 | 0.7107 | 0.016 | -0.0691 | 0.523 | 0.6217 | 0.2055 | 0.0738 | -0.1369 | -0.0363 |
| SPA69 | 0.5527 | 0.4554 | 0.4585 | 0.4585 | 0.4298 | 0.3441 | 0.5599 | 0.5811 | 0.5265 | 0.4918 | 0.3421 | 0.284 | 0.4225 | 0.3915 | 0.6027 | 0.5707 | 0.6171 | 0.5291 | 0.4305 | 0.3757 |
| SPA73 | 0.9145 | 0.912 | 0.9177 | 0.9177 | 0.9176 | 0.9028 | 0.9015 | 0.9042 | 0.9189 | 0.922 | 0.9204 | 0.9202 | 0.8763 | 0.8007 | 0.9118 | 0.9155 | 0.9203 | 0.9215 | 0.8832 | 0.8939 |
| SPA80 | 0.0339 | 0.0478 | 0.033 | 0.033 | 0.0995 | 0.0724 | 0.0651 | 0.0662 | 0.0292 | 0.0218 | 0.0415 | 0.0381 | 0.0009 | -0.0126 | 0.1718 | 0.06 | 0.3023 | -0.0212 | 0.0148 | -0.0371 |
| SPA82 | 0.1503 | 0.2003 | 0.0785 | 0.0785 | 0.1391 | 0.1029 | 0.2615 | 0.279 | 0.1761 | 0.1387 | 0.1115 | 0.1249 | 0.2522 | 0.361 | 0.2229 | 0.0902 | 0.1445 | 0.0633 | 0.0259 | 0.2167 |
| SPA83 | 0.2971 | -0.0408 | -0.3274 | -0.3274 | -0.1514 | -0.3728 | -0.3141 | -0.418 | 0.2819 | 0.1582 | 0.5702 | 0.4804 | -0.3173 | -0.2967 | 0.1312 | -0.0052 | -0.5002 | -0.5058 | -0.5533 | -0.5437 |
| SPA84 | 0.5466 | 0.5234 | 0.4078 | 0.4078 | 0.2711 | 0.5013 | 0.5748 | 0.556 | 0.3466 | 0.3222 | 0.4237 | 0.3842 | 0.5094 | 0.4889 | 0.3797 | 0.2704 | 0.0593 | 0.2046 | 0.2732 | 0.1447 |
| SPA90 | 0.4637 | 0.5984 | 0.5506 | 0.5506 | 0.4285 | 0.3322 | 0.5499 | 0.6374 | 0.5004 | 0.5617 | 0.4185 | 0.4896 | 0.2171 | 0.2879 | 0.5069 | 0.6132 | 0.6736 | 0.553 | 0.3982 | 0.376 |
| SPA91 | 0.5945 | 0.4986 | 0.4513 | 0.4513 | 0.3577 | 0.3124 | 0.5125 | 0.4989 | 0.221 | 0.4097 | 0.4164 | 0.1183 | 0.5467 | 0.5148 | 0.4166 | 0.3739 | 0.6642 | 0.4969 | 0.4053 | 0.3153 |
| SPA92 | 0.2837 | 0.103 | -0.3247 | -0.3247 | -0.0016 | 0.02 | 0.3333 | 0.357 | 0.4111 | 0.3627 | 0.1801 | 0.2205 | 0.2932 | 0.3334 | 0.573 | 0.5809 | 0.4825 | 0.4186 | -0.1382 | 0.2735 |
| SPA93 | 0.3599 | 0.3695 | 0.3158 | 0.3158 | 0.2338 | 0.172 | 0.1238 | 0.1106 | 0.3164 | 0.2943 | 0.441 | 0.3991 | 0.0418 | 0.0219 | 0.2908 | 0.3382 | | 0.3394 | 0.2714 | 0.2618 |
| SPA94 | 0.392 | 0.3719 | -0.052 | -0.052 | -0.0108 | 0.1007 | -0.2373 | -0.2193 | 0.3397 | 0.3424 | 0.3276 | 0.3407 | -0.2324 | -0.1487 | 0.1648 | 0.2202 | 0.2936 | 0.2581 | -0.0558 | -0.0207 |
| SPA95 | 0.5206 | 0.5955 | 0.4181 | 0.4181 | 0.5001 | 0.4856 | 0.5844 | 0.6365 | 0.5665 | 0.5973 | 0.5362 | 0.5044 | 0.6402 | 0.5937 | 0.5485 | 0.5486 | | 0.5275 | 0.4951 | 0.5122 |
| SPA96 | 0.5228 | 0.4795 | 0.5283 | 0.5283 | 0.2921 | 0.2767 | 0.3683 | 0.404 | 0.5834 | 0.4521 | 0.4878 | 0.446 | 0.3564 | 0.4527 | 0.4026 | 0.4273 | 0.6844 | 0.617 | 0.3837 | 0.4867 |
| SPA97 | 0.6266 | 0.6161 | 0.6017 | 0.6017 | 0.4545 | 0.3943 | 0.5023 | 0.5471 | 0.7189 | 0.6856 | 0.5689 | 0.506 | 0.4681 | 0.4601 | 0.5581 | 0.5258 | 0.6809 | 0.7031 | 0.0864 | 0.1313 |
| SPA98 | 0.698 | 0.6114 | 0.5953 | 0.5953 | 0.3023 | 0.3975 | 0.5834 | 0.5476 | 0.6573 | 0.6442 | 0.6651 | 0.708 | 0.4823 | 0.5208 | 0.2047 | 0.2767 | 0.6309 | 0.5739 | 0.1999 | 0.2973 |
| SPA99 | 0.6098 | 0.5916 | 0.5198 | 0.5198 | 0.3708 | 0.3051 | 0.3831 | 0.4047 | 0.6078 | 0.5787 | 0.6095 | 0.5739 | 0.1034 | 0.1353 | 0.6122 | 0.5877 | 0.4919 | 0.4928 | 0.2995 | 0.3513 |
| TENX128 | 0.578 | | | | 0.3679 | | 0.5341 | | | | 0.5787 | | 0.5103 | | | | 0.532 | | 0.4696 | |
| TENX147 | 0.6863 | 0.656 | 0.6893 | 0.6893 | 0.5953 | 0.6495 | 0.7039 | 0.7012 | 0.5967 | 0.5615 | 0.7081 | 0.6925 | 0.0594 | 0.0423 | 0.7328 | 0.6404 | 0.4871 | 0.4475 | 0.4943 | -0.0608 |
| TENX148 | 0.5222 | 0.5922 | 0.5277 | 0.5277 | 0.3609 | 0.4161 | 0.6384 | 0.3721 | 0.4875 | 0.4409 | 0.3928 | 0.5729 | 0.5877 | 0.3617 | | 0.4661 | 0.244 | 0.4051 | 0.375 | 0.0654 |
| TENX149 | 0.3089 | 0.381 | 0.4584 | 0.4584 | 0.3143 | 0.3465 | 0.4301 | 0.4173 | 0.3549 | 0.4237 | | 0.4518 | 0.1868 | 0.2234 | 0.3008 | 0.3681 | 0.332 | 0.3734 | 0.4191 | 0.1514 |
| TENX152 | 0.7223 | 0.7264 | 0.5822 | 0.5822 | 0.7787 | 0.6314 | 0.6355 | 0.5379 | 0.7218 | 0.7205 | 0.7346 | 0.6911 | 0.1712 | 0.4445 | | 0.2523 | 0.8415 | 0.5187 | 0.5273 | 0.2826 |
| TENX157 | 0.7652 | 0.5942 | 0.5361 | 0.5361 | 0.7414 | 0.4285 | 0.747 | 0.2513 | 0.6065 | 0.4516 | 0.7922 | 0.654 | 0.1839 | 0.0203 | 0.7278 | 0.3632 | 0.8052 | 0.3993 | 0.253 | 0.1339 |
| TENX30 | 0.1155 | 0.1275 | 0.1955 | 0.1955 | 0.1445 | 0.1436 | 0.1495 | 0.0905 | 0.1316 | 0.1679 | 0.112 | 0.2121 | 0.1153 | 0.0581 | 0.1221 | 0.0205 | 0.1979 | 0.1181 | 0.1062 | 0.0977 |
| TENX31 | 0.1129 | 0.0528 | 0.1047 | 0.1047 | 0.1142 | 0.0414 | 0.1799 | 0.0907 | 0.2284 | 0.181 | 0.1259 | 0.1898 | 0.1606 | 0.0728 | 0.1584 | -0.0544 | 0.2244 | 0.0295 | 0.1304 | 0.0258 |
| TENX46 | 0.4086 | 0.043 | 0.4664 | 0.5082 | 0.5563 | 0.3372 | 0.3478 | 0.0903 | | | 0.4107 | | -0.0494 | -0.0081 | 0.3893 | 0.1697 | 0.1707 | | 0.1998 | 0.0506 |
| TENX70 | 0.5749 | | 0.6643 | 0.6632 | 0.7802 | 0.659 | 0.5587 | 0.5764 | 0.5798 | 0.5694 | 0.6246 | 0.5416 | 0.3114 | 0.3985 | | 0.4171 | 0.7138 | 0.5057 | 0.5753 | 0.2893 |
| TENX90 | 0.4006 | 0.4589 | 0.5103 | 0.5103 | 0.4175 | 0.4943 | 0.3123 | 0.4768 | 0.3794 | 0.572 | 0.4372 | 0.5474 | 0.3426 | 0.0558 | 0.45 | 0.2774 | 0.4162 | 0.4059 | 0.3078 | 0.0871 |
| TENX91 | 0.7689 | 0.6362 | 0.626 | 0.626 | -0.0942 | -0.1844 | 0.5037 | 0.3963 | 0.7445 | 0.6768 | 0.7171 | 0.8065 | -0.1703 | 0.1319 | 0.6518 | 0.2206 | 0.6935 | 0.7008 | -0.2033 | 0.1557 |
| TENX92 | 0.7244 | 0.6807 | 0.577 | 0.577 | 0.7098 | 0.5398 | 0.721 | 0.5364 | 0.7004 | 0.406 | 0.7446 | 0.7392 | 0.5139 | 0.4097 | 0.6842 | 0.4058 | 0.6953 | 0.62 | 0.3347 | 0.2475 |
| ZEN38 | 0.6206 | 0.8076 | 0.8268 | 0.8268 | 0.8033 | 0.7967 | 0.8472 | 0.7523 | 0.7053 | 0.7957 | 0.697 | 0.8432 | 0.6371 | -0.1179 | 0.7086 | 0.8439 | 0.623 | 0.565 | 0.5845 | 0.0695 |
| ZEN42 | 0.6927 | 0.6764 | 0.6709 | 0.671 | 0.7018 | 0.6712 | 0.722 | 0.7128 | 0.7054 | 0.6473 | 0.6555 | 0.6751 | 0.5831 | 0.4288 | 0.6731 | 0.6961 | 0.6411 | 0.6805 | 0.6883 | 0.4121 |
| ZEN44 | 0.7862 | 0.7245 | 0.687 | 0.687 | 0.7047 | 0.6668 | 0.6868 | 0.7003 | 0.7385 | 0.714 | 0.7178 | 0.7153 | 0.4789 | 0.4438 | 0.7617 | 0.7358 | 0.7556 | 0.6205 | 0.516 | 0.5922 |
| ZEN45 | 0.5145 | 0.7363 | 0.6785 | 0.6785 | 0.7149 | 0.6728 | 0.6385 | 0.7379 | 0.6963 | 0.7127 | 0.5365 | 0.7016 | 0.6635 | 0.4637 | 0.7182 | 0.6905 | 0.6998 | 0.7126 | 0.4305 | 0.6547 |
| ZEN78 | 0.3808 | 0.5498 | 0.4747 | 0.4747 | 0.4416 | 0.2796 | 0.2169 | 0.5016 | 0.3592 | 0.5651 | 0.2817 | 0.3004 | 0.5357 | 0.5558 | 0.4278 | 0.4603 | 0.0393 | 0.5353 | 0.5896 | 0.5029 |
| ZEN80 | 0.7382 | 0.7323 | 0.723 | 0.723 | 0.6973 | 0.6539 | 0.7832 | 0.6862 | 0.8209 | 0.772 | 0.8398 | 0.7584 | 0.6661 | 0.6496 | 0.7839 | 0.7223 | 0.8384 | 0.7213 | 0.7148 | 0.6604 |
| ZEN81 | 0.6829 | 0.7031 | 0.6026 | 0.6026 | 0.5545 | 0.4745 | 0.6976 | 0.6923 | 0.7044 | 0.6981 | 0.6895 | 0.6505 | -0.5885 | 0.6628 | 0.6455 | 0.6566 | | 0.5528 | 0.5684 | 0.6228 |
| INT12 | | 0.3572 | 0.1855 | 0.1855 | 0.1336 | 0.1637 | 0.2253 | -0.0339 | 0.5375 | 0.4921 | 0.8099 | 0.0998 | 0.056 | 0.0303 | 0.8211 | 0.1742 | | -0.0383 | 0.0654 | -0.0801 |
| MEND148 | | 0.6919 | 0.6131 | 0.613 | 0.6071 | 0.6634 | 0.4122 | 0.6471 | 0.6338 | 0.7729 | 0.5596 | 0.6989 | 0.4139 | 0.5927 | 0.6926 | 0.7377 | 0.6475 | 0.7508 | 0.5367 | 0.5628 |
| MEND65 | | 0.8315 | 0.8423 | 0.8423 | 0.8429 | 0.832 | 0.828 | 0.8184 | | 0.8359 | 0.8574 | 0.8313 | 0.8028 | 0.7765 | 0.8492 | 0.8304 | 0.8526 | 0.835 | 0.7885 | 0.7869 |
| MISC42 | | 0.6519 | 0.5941 | 0.5941 | 0.7937 | 0.6307 | 0.5794 | 0.4421 | | 0.4577 | | 0.7691 | 0.3746 | 0.3645 | 0.8092 | 0.5282 | 0.754 | 0.5902 | 0.5931 | 0.3461 |
| MISC57 | | -0.2166 | 0.3356 | 0.3356 | 0.4603 | 0.4536 | -0.3661 | -0.299 | -0.4599 | 0.5231 | | -0.4397 | 0.3237 | 0.4558 | | 0.4371 | | -0.0416 | -0.2033 | 0.3884 |
| MISC58 | | 0.577 | -0.0101 | -0.0101 | 0.4085 | 0.0219 | 0.6662 | 0.2634 | 0.7971 | 0.317 | 0.6804 | 0.7113 | -0.2138 | -0.1405 | 0.7911 | 0.2728 | | 0.5203 | 0.3216 | -0.0358 |
| TENX40 | | 0.7229 | 0.7083 | 0.6776 | 0.7274 | 0.6537 | 0.712 | 0.6417 | 0.7929 | 0.7487 | 0.7697 | 0.7397 | 0.5729 | 0.4689 | 0.8168 | 0.7174 | | 0.7025 | 0.4294 | 0.2859 |
| ZEN79 | | 0.6779 | 0.7024 | 0.7025 | 0.6258 | 0.628 | 0.712 | 0.7003 | 0.7455 | 0.7288 | 0.7716 | 0.7245 | | 0.6979 | | 0.7557 | 0.732 | 0.6887 | 0.574 | 0.6444 |
| MEND156 | | | 0.6143 | 0.6143 | 0.639 | 0.6691 | 0.6148 | 0.7571 | 0.6839 | | 0.7216 | 0.6671 | 0.6879 | 0.6537 | 0.7487 | 0.6909 | | 0.505 | 0.6914 | 0.5965 |
| MEND141 | | | | | | 0.7513 | | | | | | | 0.5488 | 0.5419 | | | | 0.6314 | | 0.2615 |

### Table.S2

| kendall_tau | UNI_big_region | UNI_small_region | CONCH_big_region | CONCH_small_region | CONCH15_big_region | CONCH15_small_region | CTransPath_big_region | CTransPath_small_region | GigaPath_big_region | GigaPath_small_region | UNI2h_big_region | UNI2h_small_region | RN18_big_region | RN18_small_region | Virchow_big_region | Virchow_small_region | Virchow2_big_region | Virchow2_small_region | ViTB16_big_region | ViTB16_small_region |
|---|---|---|---|---|---|---|---|---|---|---|---|---|---|---|---|---|---|---|---|---|
| INT13 | 0.373 | 0.2901 | 0.3386 | 0.3621 | 0.1625 | 0.1115 | 0.3025 | 0.2841 | 0.4254 | 0.3798 | 0.4612 | 0.3591 | 0.2306 | 0.1314 | 0.3852 | 0.2886 | 0.3001 | 0.3861 | 0.2339 | 0.2126 |
| INT14 | 0.6378 | 0.5543 | 0.298 | 0.298 | 0.5207 | 0.2659 | 0.6295 | 0.4801 | 0.5806 | 0.5091 | 0.6086 | 0.4608 | 0.5061 | 0.3864 | 0.6091 | 0.5383 | | 0.3207 | 0.4925 | 0.3711 |
| INT15 | 0.3875 | 0.4446 | 0.4255 | 0.4256 | 0.4063 | 0.3695 | 0.4326 | 0.1753 | 0.4325 | 0.464 | 0.389 | 0.4129 | 0.3341 | 0.1951 | 0.4946 | 0.5033 | 0.4469 | 0.4474 | 0.3629 | 0.286 |
| INT16 | 0.6262 | 0.5448 | 0.4747 | 0.4747 | 0.5094 | 0.4424 | 0.5946 | 0.3954 | 0.6309 | 0.5481 | 0.6531 | 0.5081 | 0.1203 | 0.1076 | 0.6496 | 0.5906 | 0.5964 | 0.4729 | 0.1565 | 0.2021 |
| INT19 | 0.3777 | 0.36 | 0.1917 | 0.3835 | 0.3442 | 0.2118 | 0.2585 | 0.2015 | 0.4085 | 0.4078 | 0.4181 | 0.3752 | 0.257 | 0.2812 | 0.3831 | 0.3856 | 0.3738 | 0.3811 | 0.2854 | 0.2817 |
| INT21 | 0.603 | 0.4796 | 0.4559 | 0.4559 | 0.5236 | 0.4168 | 0.5645 | 0.0509 | 0.5837 | 0.4789 | 0.5754 | 0.3967 | 0.4407 | -0.0576 | 0.5428 | 0.4487 | 0.5004 | 0.375 | 0.398 | 0.3173 |
| INT22 | 0.1637 | 0.1445 | 0.1931 | 0.1932 | 0.0182 | 0.1384 | 0.0021 | 0.1035 | 0.1202 | 0.0436 | 0.2163 | 0.0869 | 0.1651 | 0.0944 | 0.1308 | 0.1697 | 0.2003 | -0.033 | -0.0135 | 0.0788 |
| INT24 | 0.5408 | 0.3045 | 0.283 | 0.283 | 0.4682 | 0.3445 | 0.2929 | -0.1849 | 0.5301 | 0.3656 | 0.5984 | 0.549 | -0.0278 | -0.0861 | 0.4301 | 0.4646 | 0.4857 | 0.234 | 0.2293 | 0.0361 |
| INT25 | 0.335 | 0.2949 | 0.3029 | 0.2561 | 0.3061 | 0.1881 | 0.2959 | 0.2831 | 0.3544 | 0.3046 | 0.4209 | 0.3147 | 0.2695 | 0.2111 | 0.364 | 0.2797 | 0.3359 | 0.2801 | 0.2866 | 0.2839 |
| INT26 | 0.4157 | 0.3857 | 0.4332 | 0.4332 | 0.4526 | 0.35 | 0.4617 | 0.4767 | 0.5047 | 0.5008 | 0.3862 | 0.4011 | 0.4819 | 0.4153 | 0.5252 | 0.4294 | 0.4133 | 0.4687 | 0.4851 | 0.239 |
| INT27 | 0.3985 | 0.607 | 0.5739 | 0.5739 | 0.6215 | 0.6135 | 0.601 | 0.6358 | 0.6401 | 0.645 | 0.38 | 0.5951 | 0.4355 | 0.3652 | 0.7056 | 0.5548 | 0.6027 | | 0.506 | 0.5502 |
| INT28 | 0.6632 | 0.5729 | 0.52 | 0.52 | 0.5686 | 0.5717 | 0.6536 | 0.5865 | 0.6885 | 0.5996 | 0.6582 | 0.5018 | 0.3502 | 0.3566 | 0.6663 | 0.548 | 0.6111 | 0.5379 | 0.5469 | 0.4952 |
| INT35 | 0.4707 | 0.4551 | 0.4144 | 0.4144 | 0.4903 | 0.482 | 0.4853 | 0.3709 | 0.5681 | 0.5359 | | 0.4773 | 0.3254 | 0.3632 | 0.5493 | 0.2878 | 0.5092 | 0.4985 | 0.3835 | 0.3423 |
| INT4 | 0.1059 | -0.1832 | -0.159 | -0.159 | -0.25 | -0.2489 | -0.1177 | -0.0232 | 0.0718 | -0.0542 | -0.0525 | -0.2178 | -0.2056 | -0.2005 | -0.0764 | 0.0151 | -0.2028 | -0.1501 | -0.035 | -0.1195 |
| INT5 | 0.1101 | -0.0818 | -0.1326 | -0.1326 | 0.0542 | -0.2418 | 0.0618 | -0.0819 | 0.1702 | 0.1338 | 0.2968 | -0.1553 | 0.1847 | -0.1436 | 0.1351 | -0.0047 | 0.3226 | 0.0471 | 0.1295 | -0.0351 |
| INT7 | 0.2749 | 0.312 | -0.0277 | -0.0277 | 0.0548 | -0.0504 | 0.2924 | 0.254 | 0.3057 | 0.2625 | 0.2229 | 0.2722 | -0.1256 | -0.1294 | 0.254 | 0.1907 | 0.2529 | 0.1995 | -0.0619 | -0.107 |
| INT8 | 0.1234 | -0.0166 | 0.0677 | 0.0677 | 0.0283 | 0.087 | 0.0658 | 0.0237 | 0.1514 | 0.0999 | 0.184 | 0.0669 | 0.0315 | -0.0038 | 0.0978 | 0.1136 | 0.1163 | 0.0372 | 0.1096 | 0.0204 |
| MEND140 | 0.4442 | 0.534 | 0.3068 | 0.3068 | 0.3647 | 0.0877 | 0.2577 | 0.3969 | 0.4945 | 0.4057 | 0.3892 | 0.5531 | 0.3501 | 0.4023 | 0.4954 | 0.5491 | 0.568 | 0.5268 | 0.2783 | -0.0428 |
| MEND142 | 0.5499 | 0.4087 | 0.4749 | 0.4749 | 0.5859 | 0.3915 | 0.3858 | 0.4013 | 0.5306 | 0.5232 | 0.5289 | 0.52 | 0.051 | 0.184 | 0.4398 | 0.3641 | 0.4748 | 0.357 | 0.3173 | 0.2198 |
| MEND143 | 0.3858 | 0.4383 | 0.2622 | 0.2622 | 0.1398 | 0.2294 | 0.3616 | 0.0307 | 0.4704 | 0.2287 | 0.4219 | 0.3901 | 0.064 | 0.0537 | 0.2911 | 0.3449 | 0.3993 | 0.4301 | 0.0348 | -0.0991 |
| MEND145 | 0.5629 | 0.4918 | 0.4545 | 0.4545 | 0.5312 | 0.2451 | 0.5075 | 0.5103 | 0.5871 | 0.3591 | 0.5773 | 0.4067 | 0.343 | 0.1572 | 0.5881 | 0.3433 | 0.5642 | 0.3507 | 0.188 | 0.1068 |
| MEND146 | 0.4145 | 0.5116 | 0.4285 | 0.4285 | 0.5025 | 0.1041 | 0.3672 | 0.492 | 0.5508 | 0.509 | 0.4619 | 0.3963 | 0.3106 | 0.3052 | 0.4583 | 0.3692 | 0.5103 | 0.4171 | 0.3835 | 0.4199 |
| MEND147 | 0.5628 | 0.3647 | 0.3727 | 0.3727 | 0.4261 | 0.4237 | 0.466 | 0.3758 | 0.5725 | 0.4143 | 0.5728 | 0.4367 | 0.3152 | 0.3417 | 0.5527 | 0.3121 | 0.439 | 0.3199 | 0.2749 | 0.1276 |
| MEND150 | 0.4591 | 0.6315 | 0.6382 | 0.6382 | 0.3198 | 0.5903 | 0.4137 | 0.6123 | 0.648 | 0.6468 | 0.4611 | 0.6467 | 0.489 | 0.4533 | 0.6241 | 0.6051 | 0.5145 | 0.6418 | 0.1585 | 0.5543 |
| MEND151 | 0.4849 | 0.5516 | 0.5715 | 0.5558 | 0.5088 | 0.4658 | 0.469 | | 0.5634 | 0.592 | 0.5059 | 0.5849 | 0.4264 | 0.4187 | 0.5505 | 0.5141 | 0.4833 | 0.582 | 0.2268 | 0.3447 |
| MEND153 | 0.0741 | -0.0145 | -0.0629 | -0.0629 | -0.0391 | -0.2717 | 0.3454 | 0.0804 | 0.3029 | 0.0978 | 0.1791 | -0.0486 | 0.1654 | 0.0329 | 0.0028 | -0.1273 | 0.0095 | -0.0874 | 0.1298 | 0.0497 |
| MEND154 | -0.0075 | 0.3737 | 0.4915 | 0.3899 | -0.0809 | 0.3341 | 0.0655 | 0.4136 | 0.1616 | 0.4116 | 0.049 | 0.1931 | -0.079 | 0.3193 | 0.2078 | 0.407 | 0.0558 | 0.401 | -0.0013 | 0.3138 |
| MEND157 | 0.5446 | 0.5166 | 0.4256 | 0.5456 | 0.5327 | 0.4852 | 0.4915 | 0.4773 | 0.5351 | 0.5384 | 0.557 | 0.5437 | 0.4823 | 0.4341 | 0.5505 | 0.549 | 0.5487 | 0.473 | 0.3599 | 0.4737 |
| MEND158 | 0.3573 | 0.5362 | -0.1508 | 0.5043 | 0.3503 | 0.2929 | 0.4809 | 0.5132 | 0.5214 | 0.5396 | 0.347 | 0.5379 | 0.3323 | 0.3051 | 0.2793 | 0.3706 | 0.5652 | 0.4523 | 0.2274 | 0.3317 |
| MEND160 | 0.5732 | 0.6297 | 0.6014 | 0.5782 | 0.5764 | 0.4092 | 0.5682 | 0.6038 | 0.599 | 0.6282 | 0.572 | 0.6012 | 0.5286 | 0.4286 | 0.6044 | 0.6034 | 0.5604 | 0.5991 | 0.5051 | 0.4695 |
| MEND162 | 0.4179 | 0.2507 | 0.1549 | 0.1549 | 0.2877 | 0.1672 | 0.2757 | 0.2016 | 0.282 | 0.157 | 0.4423 | 0.3107 | 0.1002 | 0.1919 | 0.2967 | 0.1997 | 0.2577 | 0.2015 | 0.142 | 0.1336 |
| MEND18 | -0.0213 | 0.052 | 0.0748 | 0.0749 | -0.128 | -0.0897 | -0.0908 | -0.0234 | -0.0117 | -0.005 | 0.0637 | 0.0898 | -0.0017 | 0.0766 | 0.0214 | 0.0251 | 0.1048 | 0.1362 | -0.0817 | -0.0317 |
| MEND19 | 0.5403 | 0.5231 | 0.4898 | 0.4898 | 0.5005 | 0.4708 | 0.5247 | 0.5267 | 0.5587 | 0.5515 | 0.5313 | 0.5437 | 0.2426 | 0.2948 | 0.4554 | 0.4268 | 0.4492 | 0.4986 | 0.2414 | 0.3207 |
| MEND24 | -0.1447 | -0.1375 | 0.0736 | 0.1208 | -0.2334 | -0.1843 | -0.0824 | -0.1253 | -0.066 | -0.0373 | -0.1549 | -0.1258 | -0.1616 | 0.0407 | 0.0175 | 0.0366 | 0.1218 | 0.1745 | -0.1795 | -0.1657 |
| MEND37 | 0.5628 | 0.507 | 0.4793 | 0.4793 | 0.4919 | 0.4426 | 0.5563 | 0.4715 | 0.5765 | 0.5001 | 0.5492 | 0.5034 | 0.4293 | 0.287 | 0.5537 | 0.3969 | 0.5496 | 0.4577 | 0.3664 | 0.2564 |
| MEND38 | 0.4046 | 0.2912 | 0.2302 | 0.2302 | 0.346 | 0.2606 | 0.3682 | 0.2946 | 0.3439 | 0.2128 | 0.4118 | 0.2576 | 0.0084 | 0.0795 | 0.3608 | 0.1164 | 0.2626 | 0.2538 | 0.0992 | 0.0902 |
| MEND39 | 0.539 | 0.4428 | 0.3652 | 0.4844 | 0.4767 | 0.4174 | 0.442 | 0.4183 | 0.477 | 0.4107 | 0.5397 | 0.3997 | 0.2692 | 0.1204 | 0.4898 | 0.3945 | 0.5084 | 0.391 | 0.1983 | 0.2017 |
| MEND40 | 0.5318 | 0.4752 | 0.4278 | 0.4278 | 0.413 | 0.3711 | 0.4856 | 0.4681 | 0.5264 | 0.4447 | 0.4999 | 0.4607 | 0.3709 | 0.3609 | 0.5062 | 0.1824 | 0.4719 | 0.4218 | 0.3885 | 0.264 |
| MEND59 | 0.5566 | 0.2658 | 0.1412 | 0.1411 | 0.3879 | 0.1785 | 0.515 | 0.2469 | 0.5087 | 0.2674 | 0.5598 | 0.2688 | 0.4591 | 0.2448 | | 0.301 | 0.5211 | 0.1944 | 0.4368 | 0.2437 |
| MEND60 | 0.5749 | 0.5366 | 0.422 | 0.422 | 0.5335 | 0.5271 | 0.5907 | 0.5654 | 0.5726 | 0.5226 | 0.5881 | 0.529 | 0.4656 | 0.5085 | 0.5945 | 0.4575 | 0.6246 | 0.5206 | 0.4389 | 0.172 |
| MEND61 | 0.5931 | 0.5423 | 0.4565 | 0.4565 | 0.5187 | 0.4691 | 0.5815 | 0.5109 | 0.6145 | 0.4489 | 0.5807 | 0.5499 | 0.5203 | 0.4652 | 0.5875 | 0.4829 | 0.5784 | 0.4056 | 0.5211 | 0.3968 |
| MEND62 | 0.6733 | 0.6422 | 0.5581 | 0.5581 | 0.6347 | 0.5475 | 0.678 | 0.6439 | 0.6818 | 0.6119 | 0.6724 | 0.6204 | 0.5987 | 0.555 | 0.6688 | 0.566 | 0.6654 | 0.5947 | 0.582 | 0.5025 |
| MEND66 | -0.0278 | 0.0422 | -0.0284 | -0.0284 | 0.0047 | 0.0221 | -0.023 | 0.0505 | 0.0188 | 0.0494 | 0.0153 | 0.0259 | 0.0055 | 0.069 | 0.0488 | 0.0685 | 0.0425 | 0.036 | 0.006 | 0.0301 |
| MISC33 | 0.584 | | 0.5221 | 0.4939 | 0.3737 | 0.0606 | 0.5519 | 0.3803 | 0.5429 | 0.5001 | 0.5711 | 0.564 | 0.0732 | -0.2708 | 0.5755 | 0.4934 | 0.5731 | 0.5564 | 0.3301 | -0.2165 |
| MISC35 | 0.522 | 0.4233 | 0.4176 | 0.4176 | 0.4708 | 0.4018 | 0.499 | 0.4375 | 0.5172 | 0.4183 | 0.5229 | 0.4385 | 0.1693 | 0.0889 | 0.5008 | 0.3894 | | 0.3966 | 0.4093 | 0.0206 |
| MISC36 | 0.432 | 0.4265 | 0.3142 | 0.3142 | 0.4423 | 0.3242 | 0.484 | 0.4513 | 0.5752 | 0.4074 | 0.412 | 0.3385 | 0.2483 | 0.2149 | 0.3099 | 0.3651 | 0.2463 | 0.1366 | 0.176 | 0.0612 |
| MISC40 | 0.393 | 0.0753 | 0.3745 | 0.3745 | 0.3351 | 0.3397 | 0.3603 | 0.3649 | 0.3609 | 0.3646 | 0.378 | 0.3465 | 0.3012 | 0.1224 | 0.3917 | 0.2828 | 0.4494 | 0.2988 | 0.0013 | 0.1877 |
| MISC41 | 0.5575 | 0.3486 | 0.4718 | 0.4718 | 0.4922 | 0.4636 | 0.4506 | -0.0443 | 0.5381 | 0.0672 | 0.5359 | 0.4728 | 0.0359 | -0.0115 | 0.5231 | 0.0157 | | 0.4096 | 0.2157 | 0.1828 |
| MISC45 | 0.3678 | 0.2855 | 0.2428 | 0.2428 | 0.3175 | 0.2565 | 0.3023 | 0.2721 | 0.3884 | 0.312 | 0.3856 | 0.3484 | 0.0601 | 0.072 | 0.3997 | 0.3012 | 0.3987 | 0.3135 | 0.1522 | 0.1368 |
| MISC47 | 0.498 | 0.2932 | 0.0727 | 0.0727 | 0.0674 | -0.0016 | 0.2024 | 0.0151 | 0.4863 | 0.2933 | 0.449 | 0.3771 | 0.0474 | 0.0357 | 0.3645 | 0.2011 | 0.404 | 0.0822 | -0.0282 | -0.0011 |
| MISC48 | 0.434 | 0.389 | 0.3682 | 0.3682 | 0.3946 | 0.3751 | 0.4529 | 0.4534 | 0.4078 | 0.3968 | 0.404 | 0.3873 | 0.3676 | 0.3406 | 0.4477 | 0.377 | 0.4255 | 0.3748 | 0.41 | 0.3536 |
| MISC49 | 0.4628 | 0.4539 | 0.379 | 0.379 | 0.4092 | 0.3424 | 0.4506 | 0.3889 | 0.5232 | 0.341 | 0.4864 | 0.4554 | 0.1194 | -0.1153 | 0.4601 | 0.408 | 0.5011 | 0.4848 | 0.3568 | 0.3352 |
| MISC50 | 0.549 | 0.3921 | 0.276 | 0.276 | 0.3764 | 0.2412 | 0.5203 | 0.2538 | 0.5722 | 0.3078 | 0.6011 | 0.4494 | 0.1739 | 0.0919 | 0.5544 | 0.2421 | 0.5373 | 0.3279 | 0.18 | 0.1971 |
| MISC62 | 0.3117 | 0.069 | 0.0427 | 0.2263 | 0.3633 | 0.2262 | 0.2232 | 0.1294 | 0.4439 | 0.1813 | 0.4033 | 0.2022 | 0.1591 | 0.1406 | 0.3975 | 0.1589 | 0.3873 | 0.1874 | 0.1437 | 0.1973 |
| MISC63 | 0.1326 | 0.0041 | 0.0988 | 0.0988 | 0.1704 | 0.0741 | 0.0921 | 0.0743 | 0.2365 | 0.0983 | 0.2247 | 0.1064 | 0.0747 | 0.0272 | 0.1807 | 0.1209 | 0.2129 | 0.1016 | 0.0525 | 0.0221 |
| MISC66 | 0.5153 | 0.257 | 0.236 | 0.1545 | 0.3296 | 0.0498 | 0.2923 | 0.1502 | 0.3931 | 0.1974 | 0.5123 | 0.3951 | -0.1785 | -0.0935 | 0.2935 | 0.0501 | 0.5162 | 0.2239 | -0.0297 | 0.0356 |
| MISC67 | 0.6593 | 0.4417 | 0.3619 | 0.3619 | 0.487 | 0.3483 | 0.4074 | 0.404 | 0.6247 | 0.4485 | 0.666 | 0.6199 | 0.0134 | 0.1214 | 0.577 | 0.5025 | 0.6682 | 0.5018 | 0.2535 | 0.1409 |
| MISC72 | 0.4219 | 0.3409 | 0.3243 | 0.3243 | 0.3779 | 0.3215 | 0.4153 | 0.3124 | 0.43 | 0.3256 | 0.392 | 0.3986 | 0.2125 | 0.076 | 0.3793 | 0.1835 | 0.383 | 0.3462 | 0.1724 | 0.1106 |
| MISC73 | 0.5587 | 0.5463 | 0.5254 | 0.5254 | 0.537 | 0.4902 | 0.5698 | 0.4284 | 0.5321 | 0.4709 | 0.5687 | 0.5734 | 0.3771 | 0.2881 | 0.5217 | 0.5081 | 0.5806 | 0.5457 | 0.1809 | 0.1563 |
| NCBI629 | 0.1864 | 0.1688 | 0.1122 | 0.1122 | 0.0294 | 0.1853 | 0.0402 | 0.1658 | 0.1242 | 0.1575 | 0.1484 | 0.098 | 0.0747 | 0.1416 | 0.1984 | 0.1659 | 0.2186 | 0.1775 | 0.2243 | 0.1452 |
| NCBI631 | 0.1917 | 0.1571 | 0.1952 | 0.185 | 0.0967 | 0.1543 | 0.1687 | 0.1609 | 0.1426 | 0.1356 | 0.1582 | 0.1884 | 0.0988 | 0.0695 | 0.1989 | 0.1672 | 0.1601 | 0.1201 | 0.0702 | 0.1725 |
| NCBI632 | 0.326 | 0.2981 | 0.2987 | 0.2987 | 0.3169 | 0.3364 | 0.334 | 0.335 | 0.3551 | 0.3033 | 0.3218 | 0.2073 | 0.2223 | 0.2476 | 0.338 | 0.2581 | 0.2959 | 0.3312 | 0.3309 | 0.3264 |
| NCBI635 | 0.3074 | 0.2983 | 0.2624 | 0.2624 | 0.2943 | 0.2791 | 0.2553 | 0.2672 | 0.3574 | 0.3209 | 0.3289 | 0.2905 | 0.2327 | 0.231 | 0.3138 | 0.2458 | 0.2598 | 0.3144 | 0.1886 | 0.2368 |
| NCBI638 | -0.0697 | -0.0267 | -0.0599 | -0.0599 | -0.0299 | -0.0623 | -0.0146 | -0.1035 | -0.0568 | -0.0358 | -0.094 | -0.0016 | 0.0893 | 0.0472 | -0.01 | 0.0467 | -0.0938 | -0.0298 | -0.022 | -0.018 |
| NCBI640 | 0.2164 | 0.1338 | 0.1562 | 0.1562 | 0.1537 | 0.1829 | 0.0316 | 0.1068 | 0.2028 | 0.169 | 0.281 | 0.2472 | 0.1415 | 0.071 | 0.229 | 0.1527 | 0.2079 | 0.1294 | 0.2246 | 0.1994 |
| NCBI681 | 0.5203 | 0.5287 | 0.5113 | 0.5113 | 0.4811 | | 0.5222 | 0.3429 | 0.5165 | 0.5369 | 0.5136 | 0.5436 | 0.3507 | 0.1827 | 0.5153 | 0.5028 | 0.5476 | 0.4991 | 0.4162 | 0.263 |
| NCBI682 | 0.5318 | 0.5037 | 0.2727 | 0.4795 | 0.4502 | 0.2872 | 0.5118 | 0.4972 | 0.5694 | 0.5293 | 0.5385 | 0.5161 | 0.4382 | 0.3527 | 0.5358 | 0.4756 | 0.539 | 0.4865 | 0.3959 | 0.4149 |
| NCBI683 | 0.4536 | 0.446 | 0.3714 | 0.3714 | 0.3865 | 0.3008 | 0.399 | 0.3697 | 0.408 | 0.4351 | 0.3833 | 0.4039 | 0.1747 | 0.0359 | 0.4243 | 0.3826 | 0.4043 | 0.4693 | 0.3636 | 0.2539 |
| NCBI760 | 0.2541 | 0.2516 | 0.1797 | 0.1797 | 0.2029 | 0.2127 | 0.2925 | 0.2631 | 0.2682 | 0.25 | 0.2464 | 0.2663 | 0.2043 | 0.2119 | 0.2266 | 0.2778 | 0.2277 | 0.2259 | 0.2299 | 0.1932 |
| NCBI764 | 0.1243 | 0.0991 | 0.1041 | 0.1041 | 0.0922 | 0.1126 | 0.1059 | 0.1027 | 0.1255 | 0.1041 | 0.1165 | 0.1056 | 0.0703 | 0.0444 | 0.1002 | 0.1035 | 0.1146 | 0.1074 | 0.0201 | -0.031 |
| NCBI765 | 0.3722 | 0.3794 | 0.399 | 0.399 | 0.337 | 0.3347 | 0.3947 | 0.3924 | 0.3739 | 0.3906 | 0.3765 | 0.3978 | 0.3146 | 0.3066 | 0.3792 | 0.3775 | 0.3441 | 0.4046 | 0.2914 | 0.3293 |
| NCBI766 | 0.3128 | 0.3044 | 0.327 | 0.327 | 0.3199 | 0.3388 | 0.3046 | 0.2924 | 0.2935 | 0.2895 | 0.2867 | 0.2937 | 0.2818 | 0.3042 | 0.3235 | 0.3321 | 0.1487 | 0.2834 | 0.3037 | 0.2895 |
| NCBI768 | 0.4891 | 0.4733 | 0.4995 | 0.4995 | 0.5339 | 0.5314 | 0.4933 | 0.4717 | 0.4921 | 0.4769 | 0.5184 | 0.491 | 0.4119 | 0.4051 | 0.3753 | 0.4311 | 0.5688 | 0.4991 | 0.4404 | 0.4599 |
| NCBI769 | 0.5301 | | 0.5531 | 0.5531 | 0.4481 | 0.5771 | 0.5492 | 0.5189 | 0.557 | 0.5357 | 0.5114 | 0.4959 | 0.4407 | 0.4208 | 0.5391 | 0.5196 | 0.5727 | 0.5513 | 0.3776 | 0.5047 |
| NCBI770 | -0.3017 | -0.2327 | -0.2531 | -0.2531 | -0.0909 | -0.1264 | -0.3215 | -0.263 | -0.1226 | -0.2492 | -0.2408 | -0.2417 | -0.2047 | -0.1776 | -0.2825 | -0.2542 | -0.2523 | -0.2531 | -0.195 | -0.175 |
| SPA100 | 0.3145 | 0.4396 | 0.0291 | 0.0291 | 0.2277 | 0.2333 | 0.3447 | 0.2936 | 0.4666 | 0.4834 | 0.4315 | 0.415 | -0.0239 | -0.0299 | 0.4592 | 0.4628 | 0.4456 | 0.4288 | 0.0987 | 0.1304 |
| SPA101 | 0.3181 | 0.2816 | 0.2678 | 0.2678 | 0.1431 | 0.183 | 0.3058 | 0.3121 | 0.2888 | 0.2825 | 0.3099 | 0.2984 | 0.1796 | 0.2407 | 0.3407 | 0.3453 | 0.2766 | 0.3219 | 0.2066 | 0.2233 |
| SPA102 | 0.2012 | 0.1798 | 0.153 | 0.153 | 0.2181 | 0.1754 | 0.1407 | 0.1362 | -0.0591 | 0.0464 | 0.1924 | 0.2287 | 0.1962 | 0.1756 | 0.2755 | 0.2761 | 0.2732 | 0.2862 | 0.2009 | |
| SPA103 | -0.1915 | -0.2099 | -0.2133 | -0.2133 | -0.2367 | -0.2743 | -0.2259 | -0.2389 | -0.2296 | -0.2318 | -0.242 | -0.231 | -0.0723 | -0.1382 | -0.2334 | -0.2412 | -0.2362 | -0.2483 | -0.0958 | -0.0072 |
| SPA104 | 0.2936 | 0.3053 | 0.1627 | 0.1627 | 0.2187 | 0.1877 | 0.2862 | 0.2588 | 0.288 | 0.2621 | 0.2712 | 0.2554 | 0.2549 | 0.2117 | 0.28 | 0.2417 | 0.2509 | 0.2044 | 0.2469 | 0.2269 |
| SPA105 | 0.1465 | 0.1483 | 0.104 | 0.104 | 0.0046 | 0.0364 | 0.1489 | 0.1379 | 0.1221 | 0.1302 | 0.0884 | 0.0563 | 0.1414 | 0.0979 | 0.1402 | 0.1132 | 0.1651 | 0.1593 | 0.0629 | 0.0642 |
| SPA106 | 0.1081 | 0.1249 | 0.0909 | 0.0909 | 0.0864 | 0.0134 | 0.0103 | 0.0245 | | -0.0275 | -0.0152 | -0.0207 | 0.036 | 0.0207 | -0.0808 | -0.0487 | 0.1034 | 0.0996 | -0.0059 | -0.0212 |
| SPA107 | 0.5622 | 0.5466 | 0.3854 | 0.3854 | 0.2969 | 0.3371 | 0.4707 | 0.5098 | 0.5743 | 0.5781 | 0.4145 | 0.4942 | 0.1615 | 0.1713 | 0.3989 | 0.4735 | 0.5135 | 0.5046 | -0.0678 | 0.0682 |
| SPA108 | 0.3558 | 0.3298 | 0.2815 | 0.2815 | 0.2748 | 0.1857 | 0.2477 | 0.2528 | 0.2953 | 0.2751 | 0.2557 | 0.2547 | 0.2886 | 0.3147 | 0.2868 | 0.272 | 0.2966 | 0.2701 | 0.2693 | 0.3099 |
| SPA109 | 0.1646 | 0.1995 | 0.1963 | 0.1963 | 0.0741 | 0.0548 | 0.1183 | 0.0568 | 0.0926 | 0.0486 | | 0.0518 | 0.075 | 0.1145 | 0.1381 | 0.1829 | 0.152 | 0.1862 | 0.0058 | 0.0158 |
| SPA111 | 0.4046 | 0.3883 | 0.361 | 0.361 | 0.3001 | 0.2424 | 0.3939 | 0.4006 | 0.4014 | 0.3771 | 0.3484 | 0.3647 | 0.3287 | 0.3056 | 0.3579 | 0.3177 | 0.3959 | 0.4041 | 0.3178 | 0.36 |
| SPA112 | 0.4649 | 0.4487 | 0.3759 | 0.3759 | 0.2712 | 0.2868 | 0.4317 | 0.412 | 0.4358 | 0.4323 | 0.4436 | 0.4247 | 0.2499 | 0.2618 | 0.3202 | 0.2326 | 0.3888 | 0.3805 | 0.3952 | 0.3052 |
| SPA114 | 0.0015 | -0.0077 | 0.003 | 0.003 | -0.0096 | 0.0174 | -0.0636 | -0.0903 | -0.0495 | 0.0917 | -0.1359 | 0.0044 | 0.0436 | 0.0068 | -0.0067 | 0.0577 | 0.0651 | 0.0441 | -0.0292 | -0.1275 |
| SPA115 | 0.0703 | 0.0629 | 0.065 | 0.065 | 0.0611 | 0.0257 | 0.0413 | 0.0351 | 0.0388 | 0.0462 | 0.0422 | 0.0367 | 0.0742 | 0.0554 | 0.0759 | 0.0541 | 0.0798 | 0.0646 | 0.0751 | 0.0767 |
| SPA116 | 0.4218 | 0.2697 | 0.1042 | 0.1042 | 0.09 | -0.2798 | -0.2921 | -0.2801 | 0.4123 | 0.4266 | -0.1518 | -0.2196 | -0.3238 | -0.3216 | 0.4966 | 0.4829 | 0.4463 | 0.4487 | -0.2783 | -0.2371 |
| SPA117 | -0.0558 | -0.0314 | 0.361 | 0.361 | 0.2314 | 0.12 | -0.1156 | -0.066 | 0.2334 | 0.0068 | -0.0894 | -0.0828 | -0.1764 | -0.1883 | -0.0561 | -0.0299 | 0.436 | 0.4344 | -0.158 | -0.1356 |
| SPA118 | 0.3568 | 0.3519 | 0.3432 | 0.3432 | 0.2453 | 0.236 | 0.3216 | 0.3067 | 0.3555 | 0.3576 | 0.3421 | 0.3368 | 0.2837 | 0.3086 | 0.3362 | 0.3333 | 0.3394 | 0.3346 | 0.2576 | 0.2808 |
| SPA55 | 0.2287 | 0.1764 | 0.0518 | 0.0518 | 0.2682 | 0.2155 | 0.2556 | 0.2629 | 0.1491 | 0.025 | 0.2284 | 0.1472 | -0.0556 | -0.1124 | 0.2969 | 0.301 | 0.148 | 0.0564 | 0.2474 | 0.203 |
| SPA57 | 0.3564 | 0.4044 | 0.1764 | 0.1764 | 0.2665 | 0.1953 | 0.3507 | 0.3669 | 0.3731 | 0.3604 | 0.3804 | 0.406 | 0.1681 | 0.104 | 0.3382 | 0.3402 | 0.2941 | -0.0675 | 0.1126 | 0.0404 |
| SPA60 | 0.2309 | 0.4101 | 0.2342 | 0.2342 | -0.1205 | 0.0516 | 0.3088 | 0.3422 | 0.3177 | 0.3776 | 0.1942 | 0.3088 | 0.0251 | -0.0145 | 0.2828 | 0.1507 | 0.3739 | 0.4253 | -0.0295 | -0.1165 |
| SPA61 | 0.602 | 0.4303 | 0.2905 | 0.2905 | 0.0645 | 0.2003 | 0.1146 | 0.2573 | 0.3679 | 0.3882 | 0.1875 | 0.3577 | 0.0647 | 0.0601 | 0.0124 | 0.1003 | 0.5412 | 0.5324 | 0.1199 | 0.1486 |
| SPA64 | 0.2681 | 0.234 | 0.179 | 0.179 | -0.1408 | -0.034 | 0.0852 | 0.0367 | 0.224 | 0.1884 | 0.2163 | 0.2218 | -0.017 | -0.1525 | 0.2456 | 0.2194 | 0.1214 | 0.2021 | -0.0284 | 0.0573 |
| SPA66 | 0.3865 | 0.3149 | -0.4144 | -0.4144 | -0.3974 | -0.4624 | 0.0274 | -0.3498 | 0.3062 | 0.3045 | 0.4417 | 0.3648 | -0.3513 | -0.259 | 0.11 | -0.2978 | 0.2742 | 0.1561 | -0.359 | -0.3402 |
| SPA67 | 0.621 | 0.4743 | 0.2061 | 0.2061 | -0.4903 | -0.4364 | -0.5027 | -0.4897 | 0.5979 | 0.4827 | 0.6135 | 0.5578 | -0.4021 | -0.3783 | 0.4075 | 0.3403 | -0.2676 | -0.3055 | -0.1207 | -0.1388 |
| SPA68 | 0.5572 | 0.5589 | -0.0657 | -0.0657 | -0.337 | -0.4262 | -0.2469 | -0.1461 | 0.465 | 0.4581 | 0.4674 | 0.4331 | -0.3663 | -0.3473 | 0.3594 | 0.3364 | 0.1071 | -0.1236 | -0.3815 | -0.2482 |
| SPA69 | 0.4171 | 0.3344 | 0.3136 | 0.3136 | 0.3208 | 0.2677 | 0.3971 | 0.4214 | 0.4322 | 0.4093 | 0.3305 | 0.2999 | 0.2841 | 0.2559 | 0.449 | 0.4131 | 0.4713 | 0.4078 | 0.2809 | 0.2564 |
| SPA73 | 0.4617 | 0.4556 | 0.4815 | 0.4815 | 0.5101 | 0.4764 | 0.5204 | 0.5141 | 0.4825 | 0.5021 | 0.4927 | 0.5015 | 0.4923 | 0.466 | 0.4334 | 0.4603 | 0.4974 | 0.5061 | 0.5326 | 0.5154 |
| SPA80 | 0.0703 | 0.1115 | 0.0909 | 0.0909 | 0.0852 | 0.0635 | 0.095 | 0.1206 | 0.0638 | 0.1158 | 0.087 | 0.1169 | -0.0161 | -0.0187 | 0.1878 | 0.1123 | 0.2808 | 0.0786 | 0.0031 | -0.0245 |
| SPA82 | -0.1262 | -0.0386 | -0.1406 | -0.1406 | -0.0515 | -0.0747 | -0.0735 | -0.0787 | -0.115 | -0.1129 | -0.1034 | -0.1253 | 0.0229 | 0.0386 | -0.0729 | -0.1655 | -0.1099 | -0.1559 | -0.0841 | -0.0203 |
| SPA83 | 0.1779 | -0.0543 | -0.212 | -0.212 | -0.1822 | -0.217 | -0.2587 | -0.3189 | 0.0293 | -0.0494 | 0.3298 | 0.2979 | -0.1642 | -0.1943 | 0.1067 | 0.003 | -0.3245 | -0.3348 | -0.283 | -0.3099 |
| SPA84 | 0.3803 | 0.3679 | 0.2846 | 0.2846 | 0.1408 | 0.3516 | 0.3876 | 0.3635 | 0.2387 | 0.2206 | 0.2945 | 0.2808 | 0.3634 | 0.3569 | 0.2558 | 0.213 | 0.039 | 0.1783 | 0.1513 | 0.0721 |
| SPA90 | 0.3948 | 0.4902 | 0.3824 | 0.3824 | 0.2817 | 0.1738 | 0.4042 | 0.4516 | 0.3655 | 0.4177 | 0.3308 | 0.3509 | 0.1735 | 0.2092 | 0.3679 | 0.4292 | 0.5018 | 0.392 | 0.2464 | 0.2607 |
| SPA91 | 0.4586 | 0.3859 | 0.3607 | 0.3607 | 0.2432 | 0.2348 | 0.3842 | 0.3927 | 0.3342 | 0.3648 | 0.3591 | 0.1997 | 0.398 | 0.3648 | 0.2685 | 0.252 | 0.5288 | 0.3825 | 0.2908 | 0.2468 |
| SPA92 | 0.3833 | 0.287 | -0.2212 | -0.2212 | 0.0299 | 0.0252 | 0.3517 | 0.3089 | 0.3752 | 0.2646 | 0.265 | 0.233 | 0.219 | 0.238 | 0.4049 | 0.415 | 0.3305 | 0.3031 | -0.1793 | 0.1843 |
| SPA93 | 0.2721 | 0.2788 | 0.1518 | 0.1518 | 0.1562 | 0.1223 | 0.1075 | 0.0987 | 0.2457 | 0.2188 | 0.3102 | 0.2935 | 0.0526 | 0.0521 | 0.1531 | 0.1454 | | 0.1112 | 0.1465 | 0.1883 |
| SPA94 | 0.3059 | 0.2685 | 0.0217 | 0.0216 | 0.066 | 0.0662 | -0.0638 | -0.0617 | 0.2341 | 0.2289 | 0.2504 | 0.2454 | -0.0769 | -0.0424 | 0.1523 | 0.157 | 0.1697 | 0.1541 | 0.0247 | 0.0311 |
| SPA95 | 0.3787 | 0.3657 | 0.2774 | 0.2774 | 0.3214 | 0.2948 | 0.3427 | 0.3515 | 0.4058 | 0.4111 | 0.3891 | 0.3628 | 0.3194 | 0.3118 | 0.3861 | 0.3572 | | 0.304 | 0.3052 | 0.2963 |
| SPA96 | 0.2961 | 0.2359 | 0.2862 | 0.2862 | 0.1633 | 0.1594 | 0.1804 | 0.1862 | 0.33 | 0.2356 | 0.2999 | 0.2531 | 0.1123 | 0.1542 | 0.1875 | 0.1985 | 0.3513 | 0.2811 | 0.1782 | 0.2357 |
| SPA97 | 0.3237 | 0.347 | 0.4091 | 0.4091 | 0.2729 | 0.287 | 0.336 | 0.3582 | 0.353 | 0.4266 | 0.3249 | 0.3244 | 0.301 | 0.2858 | 0.2359 | 0.1994 | 0.4055 | 0.4057 | 0.0168 | 0.0762 |
| SPA98 | 0.4047 | 0.3371 | 0.3218 | 0.3218 | 0.1693 | 0.215 | 0.29 | 0.2807 | 0.3793 | 0.3276 | 0.4635 | 0.4144 | 0.2271 | 0.2181 | 0.1362 | 0.1688 | 0.36 | 0.2876 | 0.0879 | 0.146 |
| SPA99 | 0.3356 | 0.3513 | 0.3394 | 0.3394 | 0.243 | 0.1988 | 0.2678 | 0.2572 | 0.369 | 0.3649 | 0.3684 | 0.3706 | 0.0697 | 0.0932 | 0.3828 | 0.3717 | 0.3412 | 0.3542 | 0.2273 | 0.2915 |
| TENX128 | 0.2785 | | | | 0.2707 | | 0.2728 | | | | 0.268 | | 0.2255 | | | | 0.3194 | | 0.1812 | |
| TENX147 | 0.3947 | 0.4456 | 0.5114 | 0.5114 | 0.439 | 0.4782 | 0.379 | 0.4599 | 0.3898 | 0.4277 | 0.4306 | 0.4785 | 0.0974 | 0.0644 | 0.5902 | 0.5232 | 0.2925 | 0.3097 | 0.3229 | 0.0354 |
| TENX148 | 0.5497 | 0.5032 | 0.4084 | 0.4084 | 0.4719 | 0.423 | 0.5578 | 0.202 | 0.5233 | 0.2872 | 0.5631 | 0.3875 | 0.3233 | 0.1637 | | 0.3153 | 0.528 | 0.5109 | -0.0386 | -0.1502 |
| TENX149 | 0.3373 | 0.5001 | 0.5478 | 0.5478 | 0.3233 | 0.4581 | 0.5768 | 0.543 | 0.4329 | 0.5312 | | 0.5686 | 0.0409 | 0.1759 | 0.3443 | 0.5346 | 0.3898 | 0.4957 | 0.2766 | 0.1919 |
| TENX152 | 0.5744 | 0.4742 | 0.3397 | 0.3397 | 0.5111 | 0.3832 | 0.5217 | 0.3958 | 0.5866 | 0.5032 | 0.5426 | 0.4647 | 0.3217 | 0.2544 | | 0.1927 | 0.5748 | 0.3176 | 0.4004 | 0.2141 |
| TENX157 | 0.5057 | 0.4199 | 0.3989 | 0.3989 | 0.4815 | 0.3259 | 0.413 | 0.2962 | 0.47 | 0.3504 | 0.4969 | 0.44 | 0.183 | 0.1513 | 0.5191 | 0.3271 | 0.4904 | 0.3355 | 0.1624 | 0.1145 |
| TENX30 | 0.0963 | 0.0924 | 0.1203 | 0.1203 | 0.1171 | 0.0741 | 0.084 | 0.0456 | 0.0745 | 0.1167 | 0.031 | 0.1255 | 0.053 | 0.0337 | 0.0468 | 0.0185 | 0.1596 | 0.0669 | 0.0524 | 0.0501 |
| TENX31 | 0.0286 | -0.0289 | 0.0395 | 0.0395 | 0.0274 | -0.0276 | 0.0398 | 0.0008 | 0.0531 | 0.0305 | 0.066 | 0.108 | 0.0344 | 0.01 | 0.0577 | -0.0299 | 0.0897 | -0.035 | -0.0075 | -0.0286 |
| TENX46 | 0.3487 | 0.3177 | 0.3234 | 0.3407 | 0.3554 | 0.2313 | 0.4606 | 0.1968 | | | 0.4325 | | 0.0399 | 0.0475 | 0.5122 | 0.2763 | 0.2733 | | 0.1778 | 0.0293 |
| TENX70 | 0.5797 | | 0.5476 | 0.5468 | 0.6267 | 0.554 | 0.5379 | 0.5311 | 0.5464 | 0.5551 | 0.592 | 0.5774 | 0.3716 | 0.4084 | | 0.4236 | 0.6238 | 0.4992 | 0.4434 | 0.2018 |
| TENX90 | 0.4138 | 0.3939 | 0.4338 | 0.4338 | 0.345 | 0.3636 | 0.3234 | 0.3389 | 0.4021 | 0.4452 | 0.3981 | 0.4519 | 0.2148 | 0.036 | 0.3567 | 0.1591 | 0.4186 | 0.3493 | 0.2188 | 0.0824 |
| TENX91 | 0.6658 | 0.4322 | 0.4381 | 0.4381 | -0.0672 | -0.1269 | 0.3861 | 0.307 | 0.5774 | 0.455 | 0.5594 | 0.5683 | -0.0963 | 0.0473 | 0.483 | 0.1861 | 0.5113 | 0.5006 | -0.1229 | 0.1302 |
| TENX92 | 0.5958 | 0.4999 | 0.4675 | 0.4675 | 0.5533 | 0.423 | 0.5661 | 0.3852 | 0.6133 | 0.2955 | 0.5974 | 0.5787 | 0.2545 | 0.1903 | 0.4872 | 0.2414 | 0.5512 | 0.4779 | 0.1857 | 0.1829 |
| ZEN38 | 0.4531 | 0.5897 | 0.607 | 0.607 | 0.6146 | 0.6094 | 0.6416 | 0.5314 | 0.531 | 0.5799 | 0.5245 | 0.6299 | 0.4783 | -0.0674 | 0.515 | 0.6239 | 0.4922 | 0.3706 | 0.4055 | 0.0476 |
| ZEN42 | 0.5557 | 0.501 | 0.5023 | 0.5023 | 0.5299 | 0.5074 | 0.5595 | 0.5388 | 0.5539 | 0.4874 | 0.5165 | 0.5225 | 0.4367 | 0.2587 | 0.5751 | 0.5212 | 0.5052 | 0.5194 | 0.5141 | 0.2715 |
| ZEN44 | 0.5973 | 0.5268 | 0.5291 | 0.5291 | 0.5262 | 0.5103 | 0.5187 | 0.5167 | 0.5701 | 0.5283 | 0.5643 | 0.5168 | 0.3862 | 0.331 | 0.565 | 0.5554 | 0.5708 | 0.4398 | 0.3781 | 0.4457 |
| ZEN45 | 0.4086 | 0.5653 | 0.4801 | 0.4801 | 0.5116 | 0.5143 | 0.4432 | 0.5745 | 0.4895 | 0.5492 | 0.3674 | 0.5701 | 0.4981 | 0.3107 | 0.4526 | 0.4999 | 0.4778 | 0.5063 | 0.3237 | 0.469 |
| ZEN78 | 0.2522 | 0.4456 | 0.309 | 0.309 | 0.2947 | 0.1706 | 0.1296 | 0.3469 | 0.2273 | 0.3906 | 0.1928 | 0.2061 | 0.3765 | 0.3775 | 0.2631 | 0.3016 | 0.0448 | 0.3712 | 0.4553 | 0.4001 |
| ZEN80 | 0.5452 | 0.5384 | 0.5581 | 0.5581 | 0.5314 | 0.507 | 0.5702 | 0.5071 | 0.6039 | 0.569 | 0.6012 | 0.5469 | 0.5076 | 0.5171 | 0.5835 | 0.5114 | 0.5767 | 0.5092 | 0.5349 | 0.5319 |
| ZEN81 | 0.4765 | 0.4916 | 0.4463 | 0.4463 | 0.3936 | 0.3815 | 0.4841 | 0.4865 | 0.491 | 0.4844 | 0.4717 | 0.4586 | -0.3589 | 0.4637 | 0.4463 | 0.4712 | | 0.4054 | 0.3916 | 0.4257 |
| INT12 | | 0.283 | 0.3428 | 0.3428 | 0.2594 | 0.3168 | 0.2516 | 0.0484 | 0.4716 | 0.4171 | 0.3284 | 0.261 | 0.1166 | 0.0395 | 0.4694 | 0.2753 | | 0.2193 | 0.133 | -0.0145 |
| MEND148 | | 0.3977 | 0.1761 | 0.1761 | 0.2088 | 0.2498 | 0.3466 | 0.3133 | 0.4228 | 0.4085 | 0.4482 | 0.3827 | 0.1334 | 0.344 | 0.4086 | 0.3611 | 0.3237 | 0.3948 | 0.3567 | 0.3773 |
| MEND65 | | 0.2097 | 0.1093 | 0.1093 | 0.0085 | 0.104 | 0.1826 | 0.021 | | 0.1107 | 0.2455 | 0.1698 | 0.0521 | 0.071 | 0.2193 | 0.1291 | 0.1717 | 0.1203 | 0.0281 | 0.0503 |
| MISC42 | | 0.5036 | 0.4906 | 0.4906 | 0.5972 | 0.488 | 0.4023 | 0.2679 | | 0.277 | | 0.5632 | 0.3227 | 0.2835 | 0.6025 | 0.4358 | 0.5728 | 0.4498 | 0.4122 | 0.3205 |
| MISC57 | | -0.0496 | 0.22 | 0.22 | 0.3278 | 0.3327 | -0.1534 | -0.1739 | -0.2304 | 0.3883 | | -0.1468 | 0.2163 | 0.3312 | | 0.3225 | | 0.086 | -0.0779 | 0.3127 |
| MISC58 | | 0.2667 | -0.0087 | -0.0087 | 0.2138 | 0.0252 | 0.2759 | 0.1448 | 0.5697 | 0.1317 | 0.5801 | 0.3346 | -0.3724 | -0.2129 | 0.4522 | 0.1098 | | 0.2123 | 0.2259 | -0.1063 |
| TENX40 | | 0.655 | 0.4297 | 0.5359 | 0.6311 | 0.5369 | 0.6571 | 0.5543 | 0.6933 | 0.6494 | 0.6768 | 0.6715 | 0.4959 | 0.3843 | 0.6866 | 0.6218 | | 0.6047 | 0.3718 | 0.2348 |
| ZEN79 | | 0.5176 | 0.5354 | 0.5355 | 0.5041 | 0.4798 | 0.5421 | 0.5414 | 0.5614 | 0.5428 | 0.589 | 0.5304 | | 0.5328 | | 0.565 | 0.5749 | 0.5462 | 0.4643 | 0.4996 |
| MEND156 | | | 0.5269 | 0.5269 | 0.5765 | 0.5561 | 0.2398 | 0.5708 | 0.5918 | | 0.509 | 0.6322 | 0.443 | 0.4316 | 0.5987 | 0.6085 | | 0.4626 | 0.5323 | 0.4584 |
| MEND141 | | | | | | 0.3072 | | | | | | | 0.2392 | 0.2878 | | | | 0.3568 | | 0.1282 |

### Table.S3

| spearman | UNI_big_region | UNI_small_region | CONCH_big_region | CONCH_small_region | CONCH15_big_region | CONCH15_small_region | CTransPath_big_region | CTransPath_small_region | GigaPath_big_region | GigaPath_small_region | UNI2h_big_region | UNI2h_small_region | RN18_big_region | RN18_small_region | Virchow_big_region | Virchow_small_region | Virchow2_big_region | Virchow2_small_region | ViTB16_big_region | ViTB16_small_region |
|---|---|---|---|---|---|---|---|---|---|---|---|---|---|---|---|---|---|---|---|---|
| INT13 | 0.5565 | 0.4189 | 0.479 | 0.5207 | 0.2301 | 0.1488 | 0.4358 | 0.4104 | 0.611 | 0.5444 | 0.647 | 0.5162 | 0.3365 | 0.1943 | 0.5498 | 0.416 | 0.4321 | 0.5529 | 0.3443 | 0.3093 |
| INT14 | 0.8456 | 0.7787 | 0.4355 | 0.4355 | 0.7262 | 0.391 | 0.8402 | 0.6786 | 0.7996 | 0.7366 | 0.8219 | 0.6697 | 0.704 | 0.5564 | 0.8249 | 0.7613 | | 0.4699 | 0.6795 | 0.5275 |
| INT15 | 0.5373 | 0.6079 | 0.5841 | 0.5842 | 0.5598 | 0.5147 | 0.5908 | 0.2554 | 0.5871 | 0.629 | 0.5384 | 0.5698 | 0.4647 | 0.2782 | 0.6627 | 0.6732 | 0.604 | 0.6098 | 0.5045 | 0.4052 |
| INT16 | 0.8129 | 0.7474 | 0.6645 | 0.6645 | 0.7114 | 0.6312 | 0.7816 | 0.544 | 0.8197 | 0.7506 | 0.827 | 0.692 | 0.1761 | 0.1561 | 0.8396 | 0.7896 | 0.7962 | 0.6645 | 0.2208 | 0.2935 |
| INT19 | 0.5395 | 0.5109 | 0.2669 | 0.5386 | 0.4947 | 0.3006 | 0.3733 | 0.2977 | 0.5742 | 0.5666 | 0.5859 | 0.5304 | 0.3746 | 0.4078 | 0.5409 | 0.5433 | 0.5346 | 0.5425 | 0.4115 | 0.4082 |
| INT21 | 0.7955 | 0.675 | 0.6584 | 0.6584 | 0.7214 | 0.6049 | 0.7585 | 0.0845 | 0.7741 | 0.6743 | 0.765 | 0.5795 | 0.618 | -0.0871 | 0.7272 | 0.6393 | 0.6845 | 0.5432 | 0.5742 | 0.4768 |
| INT22 | 0.2254 | 0.2091 | 0.2862 | 0.2864 | 0.0247 | 0.2071 | -0.0032 | 0.1535 | 0.1603 | 0.0608 | 0.2922 | 0.1306 | 0.2443 | 0.138 | 0.1823 | 0.2489 | 0.2894 | -0.0471 | -0.0216 | 0.1177 |
| INT24 | 0.7168 | 0.4181 | 0.3904 | 0.3904 | 0.6332 | 0.4732 | 0.407 | -0.298 | 0.7073 | 0.4945 | 0.7808 | 0.7324 | -0.0532 | -0.1217 | 0.5843 | 0.628 | 0.6493 | 0.3405 | 0.3451 | 0.0659 |
| INT25 | 0.4741 | 0.426 | 0.4433 | 0.3749 | 0.4382 | 0.2832 | 0.4203 | 0.4114 | 0.4953 | 0.4359 | 0.5855 | 0.4482 | 0.3909 | 0.3108 | 0.5132 | 0.405 | 0.4796 | 0.4073 | 0.4141 | 0.4138 |
| INT26 | 0.5859 | 0.5436 | 0.6128 | 0.6129 | 0.6361 | 0.5099 | 0.6491 | 0.6758 | 0.6954 | 0.699 | 0.5221 | 0.5543 | 0.6736 | 0.5965 | 0.7215 | 0.6122 | 0.5801 | 0.6542 | 0.6789 | 0.3561 |
| INT27 | 0.5303 | 0.815 | 0.7692 | 0.7692 | 0.8319 | 0.8227 | 0.8062 | 0.8403 | 0.8447 | 0.8484 | 0.5044 | 0.7936 | 0.617 | 0.528 | 0.8948 | 0.7637 | 0.8144 | | 0.7018 | 0.7603 |
| INT28 | 0.8563 | 0.7809 | 0.7193 | 0.7193 | 0.7868 | 0.7871 | 0.8525 | 0.7966 | 0.8798 | 0.8129 | 0.8537 | 0.6967 | 0.5106 | 0.514 | 0.8574 | 0.7623 | 0.8191 | 0.7464 | 0.7561 | 0.6986 |
| INT35 | 0.656 | 0.634 | 0.6056 | 0.6055 | 0.6873 | 0.6883 | 0.6704 | 0.5409 | 0.7687 | 0.7453 | | 0.6683 | 0.4653 | 0.5332 | 0.7563 | 0.4212 | 0.7072 | 0.705 | 0.5464 | 0.4929 |
| INT4 | 0.1648 | -0.295 | -0.235 | -0.235 | -0.3786 | -0.3746 | -0.1894 | -0.0424 | 0.0901 | -0.0862 | -0.0832 | -0.3264 | -0.3272 | -0.3008 | -0.1181 | 0.0178 | -0.3169 | -0.2277 | -0.0701 | -0.1811 |
| INT5 | 0.151 | -0.1181 | -0.194 | -0.194 | 0.0684 | -0.3488 | 0.0654 | -0.1208 | 0.2166 | 0.1859 | 0.3918 | -0.2286 | 0.2589 | -0.2105 | 0.1837 | -0.0079 | 0.4344 | 0.0721 | 0.181 | -0.0525 |
| INT7 | 0.4105 | 0.465 | -0.039 | -0.0391 | 0.0847 | -0.0719 | 0.4332 | 0.377 | 0.4484 | 0.3905 | 0.3228 | 0.3935 | -0.187 | -0.1915 | 0.3624 | 0.2832 | 0.3694 | 0.2951 | -0.0913 | -0.1581 |
| INT8 | 0.1771 | -0.0269 | 0.1005 | 0.1005 | 0.042 | 0.1308 | 0.0987 | 0.0342 | 0.2162 | 0.1442 | 0.2654 | 0.0998 | 0.0469 | -0.005 | 0.1448 | 0.1683 | 0.1741 | 0.0547 | 0.1618 | 0.0322 |
| MEND140 | 0.6287 | 0.7312 | 0.4621 | 0.4621 | 0.5317 | 0.1378 | 0.3825 | 0.5724 | 0.6861 | 0.5867 | 0.56 | 0.7478 | 0.4988 | 0.5829 | 0.6867 | 0.7422 | 0.7657 | 0.7248 | 0.4218 | -0.0601 |
| MEND142 | 0.7287 | 0.5806 | 0.6543 | 0.6543 | 0.7825 | 0.5508 | 0.5476 | 0.568 | 0.7205 | 0.7147 | 0.7041 | 0.7058 | 0.0762 | 0.2675 | 0.6178 | 0.5281 | 0.6453 | 0.5141 | 0.4602 | 0.3231 |
| MEND143 | 0.5393 | 0.603 | 0.3766 | 0.3766 | 0.2102 | 0.3355 | 0.5163 | 0.0618 | 0.6455 | 0.3311 | 0.5881 | 0.5515 | 0.0962 | 0.0816 | 0.4232 | 0.4914 | 0.5653 | 0.5982 | 0.0529 | -0.144 |
| MEND145 | 0.7537 | 0.6746 | 0.63 | 0.63 | 0.7145 | 0.365 | 0.6934 | 0.6943 | 0.7674 | 0.5109 | 0.769 | 0.577 | 0.4893 | 0.2276 | 0.7853 | 0.4919 | 0.7676 | 0.5031 | 0.2828 | 0.161 |
| MEND146 | 0.5913 | 0.6976 | 0.6082 | 0.6082 | 0.6915 | 0.1601 | 0.5364 | 0.6769 | 0.7422 | 0.6977 | 0.6477 | 0.5632 | 0.46 | 0.4485 | 0.6593 | 0.5374 | 0.7116 | 0.5906 | 0.548 | 0.5959 |
| MEND147 | 0.7575 | 0.5385 | 0.5298 | 0.5298 | 0.6218 | 0.6085 | 0.6459 | 0.5662 | 0.7749 | 0.5881 | 0.7692 | 0.6181 | 0.4756 | 0.5093 | 0.7513 | 0.4614 | 0.6198 | 0.4715 | 0.416 | 0.1944 |
| MEND150 | 0.642 | 0.8427 | 0.8432 | 0.8432 | 0.4588 | 0.8006 | 0.5874 | 0.825 | 0.8466 | 0.8522 | 0.6253 | 0.8523 | 0.6837 | 0.6397 | 0.8327 | 0.8165 | 0.7229 | 0.8481 | 0.2171 | 0.7603 |
| MEND151 | 0.6757 | 0.7487 | 0.7725 | 0.7606 | 0.7074 | 0.6532 | 0.6606 | | 0.7543 | 0.797 | 0.6991 | 0.786 | 0.6157 | 0.5937 | 0.7431 | 0.7036 | 0.6793 | 0.7933 | 0.3198 | 0.4885 |
| MEND153 | 0.1141 | -0.0209 | -0.0916 | -0.0916 | -0.059 | -0.4087 | 0.4962 | 0.1124 | 0.4265 | 0.1393 | 0.26 | -0.0802 | 0.2416 | 0.0502 | 0.0004 | -0.1968 | 0.0062 | -0.1298 | 0.1918 | 0.0732 |
| MEND154 | -0.0153 | 0.5213 | 0.7017 | 0.5814 | -0.1317 | 0.4777 | 0.0971 | 0.5959 | 0.2419 | 0.6053 | 0.0806 | 0.279 | -0.1212 | 0.4756 | 0.3021 | 0.6005 | 0.0943 | 0.5937 | -0.0026 | 0.4512 |
| MEND157 | 0.7523 | 0.7275 | 0.6116 | 0.7548 | 0.7418 | 0.6894 | 0.6931 | 0.6774 | 0.7363 | 0.7493 | 0.7684 | 0.7573 | 0.6885 | 0.6222 | 0.7555 | 0.761 | 0.7578 | 0.6758 | 0.5247 | 0.6798 |
| MEND158 | 0.4984 | 0.729 | -0.2483 | 0.698 | 0.513 | 0.4357 | 0.6786 | 0.7244 | 0.7028 | 0.7366 | 0.4815 | 0.7286 | 0.4945 | 0.4427 | 0.394 | 0.5317 | 0.7645 | 0.6368 | 0.3364 | 0.4933 |
| MEND160 | 0.7805 | 0.8365 | 0.8085 | 0.7884 | 0.7828 | 0.5786 | 0.7746 | 0.8122 | 0.8064 | 0.8346 | 0.7784 | 0.8119 | 0.7309 | 0.6085 | 0.8093 | 0.8086 | 0.775 | 0.8099 | 0.7033 | 0.6519 |
| MEND162 | 0.5944 | 0.3667 | 0.228 | 0.228 | 0.4206 | 0.25 | 0.4052 | 0.2969 | 0.4091 | 0.2313 | 0.6253 | 0.4505 | 0.1411 | 0.2837 | 0.4261 | 0.2951 | 0.369 | 0.2959 | 0.2091 | 0.1967 |
| MEND18 | -0.0266 | 0.0854 | 0.127 | 0.1272 | -0.1893 | -0.1356 | -0.1285 | -0.0203 | -0.0098 | 0.0033 | 0.1098 | 0.14 | -0.0024 | 0.114 | 0.0391 | 0.0529 | 0.1608 | 0.2026 | -0.1114 | -0.0419 |
| MEND19 | 0.7467 | 0.7314 | 0.6932 | 0.6932 | 0.6944 | 0.6829 | 0.7165 | 0.7291 | 0.7649 | 0.7526 | 0.7334 | 0.7475 | 0.3655 | 0.4472 | 0.6394 | 0.6056 | 0.6418 | 0.6881 | 0.3717 | 0.4771 |
| MEND24 | -0.2302 | -0.2002 | 0.115 | 0.1708 | -0.3461 | -0.2749 | -0.121 | -0.1988 | -0.1236 | -0.0667 | -0.2559 | -0.2155 | -0.2353 | 0.0715 | 0.0228 | 0.0455 | 0.1888 | 0.2796 | -0.2657 | -0.2503 |
| MEND37 | 0.7438 | 0.6793 | 0.6595 | 0.6594 | 0.6694 | 0.6155 | 0.7406 | 0.6433 | 0.7583 | 0.675 | 0.732 | 0.676 | 0.6056 | 0.4146 | 0.7395 | 0.5596 | 0.731 | 0.6255 | 0.5231 | 0.3706 |
| MEND38 | 0.553 | 0.4318 | 0.3436 | 0.3436 | 0.487 | 0.3758 | 0.5059 | 0.4406 | 0.4783 | 0.3297 | 0.5757 | 0.3897 | 0.0218 | 0.1192 | 0.5019 | 0.1722 | 0.3476 | 0.3632 | 0.1305 | 0.1397 |
| MEND39 | 0.7206 | 0.6122 | 0.5101 | 0.6635 | 0.653 | 0.5835 | 0.6157 | 0.5823 | 0.6525 | 0.5733 | 0.7197 | 0.5606 | 0.411 | 0.1819 | 0.6657 | 0.5577 | 0.6882 | 0.5483 | 0.3114 | 0.3032 |
| MEND40 | 0.715 | 0.6487 | 0.591 | 0.591 | 0.5792 | 0.5183 | 0.664 | 0.637 | 0.7105 | 0.6154 | 0.679 | 0.6371 | 0.5298 | 0.5143 | 0.6926 | 0.2807 | 0.644 | 0.5897 | 0.5539 | 0.3815 |
| MEND59 | 0.7898 | 0.3642 | 0.1869 | 0.1869 | 0.5608 | 0.2405 | 0.72 | 0.3454 | 0.748 | 0.3922 | 0.793 | 0.3502 | 0.663 | 0.3711 | | 0.4629 | 0.7575 | 0.2591 | 0.6192 | 0.3586 |
| MEND60 | 0.7914 | 0.7641 | 0.6215 | 0.6215 | 0.7604 | 0.7436 | 0.8131 | 0.7892 | 0.7985 | 0.7498 | 0.8091 | 0.7568 | 0.6948 | 0.7255 | 0.8134 | 0.6754 | 0.8387 | 0.7328 | 0.6485 | 0.2546 |
| MEND61 | 0.81 | 0.7593 | 0.6612 | 0.6612 | 0.7395 | 0.6733 | 0.8009 | 0.7234 | 0.8287 | 0.6458 | 0.7999 | 0.7717 | 0.7424 | 0.6665 | 0.8047 | 0.6996 | 0.7927 | 0.5913 | 0.7402 | 0.5775 |
| MEND62 | 0.8687 | 0.8469 | 0.7671 | 0.7671 | 0.8421 | 0.759 | 0.8727 | 0.8481 | 0.875 | 0.8201 | 0.8675 | 0.8264 | 0.8119 | 0.7643 | 0.863 | 0.7792 | 0.8639 | 0.8077 | 0.7919 | 0.7102 |
| MEND66 | -0.0358 | 0.0619 | -0.0442 | -0.0442 | 0.0042 | 0.0335 | -0.0295 | 0.0729 | 0.0325 | 0.0723 | 0.0229 | 0.0364 | 0.0058 | 0.1031 | 0.0742 | 0.1007 | 0.062 | 0.0514 | 0.0071 | 0.0444 |
| MISC33 | 0.7941 | | 0.7369 | 0.7077 | 0.5352 | 0.0908 | 0.7506 | 0.5421 | 0.7612 | 0.6992 | 0.7927 | 0.7815 | 0.1075 | -0.4216 | 0.7936 | 0.7006 | 0.7909 | 0.771 | 0.4972 | -0.3284 |
| MISC35 | 0.6949 | 0.5937 | 0.5828 | 0.5828 | 0.6467 | 0.5708 | 0.6757 | 0.6095 | 0.6919 | 0.5917 | 0.7032 | 0.6142 | 0.2398 | 0.1308 | 0.6746 | 0.549 | | 0.5617 | 0.5808 | 0.025 |
| MISC36 | 0.5981 | 0.6084 | 0.4754 | 0.4754 | 0.6085 | 0.4814 | 0.6558 | 0.6039 | 0.7741 | 0.5877 | 0.5765 | 0.5176 | 0.3646 | 0.3251 | 0.4641 | 0.5279 | 0.3987 | 0.2156 | 0.2525 | 0.0973 |
| MISC40 | 0.5562 | 0.114 | 0.5414 | 0.5414 | 0.4955 | 0.5023 | 0.5175 | 0.5273 | 0.5172 | 0.54 | 0.5117 | 0.4965 | 0.4358 | 0.1796 | 0.5552 | 0.4191 | 0.6154 | 0.4402 | 0.0005 | 0.274 |
| MISC41 | 0.7737 | 0.5447 | 0.6854 | 0.6854 | 0.7181 | 0.683 | 0.6782 | -0.0644 | 0.7596 | 0.1064 | 0.7569 | 0.7034 | 0.0619 | -0.0163 | 0.7471 | 0.0207 | | 0.6287 | 0.3279 | 0.2762 |
| MISC45 | 0.5001 | 0.4029 | 0.3545 | 0.3545 | 0.4426 | 0.3693 | 0.4215 | 0.3905 | 0.5283 | 0.4374 | 0.5252 | 0.4801 | 0.0966 | 0.1115 | 0.5457 | 0.4249 | 0.5407 | 0.4438 | 0.2287 | 0.207 |
| MISC47 | 0.6682 | 0.4191 | 0.1095 | 0.1095 | 0.1249 | -0.0014 | 0.3159 | 0.03 | 0.6507 | 0.4179 | 0.6055 | 0.5217 | 0.0734 | 0.056 | 0.5143 | 0.2934 | 0.5543 | 0.1247 | -0.0227 | -0.0028 |
| MISC48 | 0.5931 | 0.5391 | 0.5183 | 0.5182 | 0.5495 | 0.5261 | 0.617 | 0.6192 | 0.5634 | 0.5475 | 0.5604 | 0.5417 | 0.5177 | 0.4817 | 0.6116 | 0.5278 | 0.5885 | 0.5243 | 0.5672 | 0.4954 |
| MISC49 | 0.6438 | 0.6323 | 0.545 | 0.545 | 0.59 | 0.5014 | 0.6381 | 0.5579 | 0.7204 | 0.4941 | 0.6702 | 0.6356 | 0.1748 | -0.1851 | 0.6334 | 0.5819 | 0.6796 | 0.6632 | 0.5167 | 0.4838 |
| MISC50 | 0.7411 | 0.5544 | 0.4067 | 0.4067 | 0.5343 | 0.3556 | 0.7099 | 0.3765 | 0.7612 | 0.447 | 0.8012 | 0.6282 | 0.261 | 0.1421 | 0.7451 | 0.3545 | 0.7276 | 0.4795 | 0.2711 | 0.2967 |
| MISC62 | 0.4504 | 0.1053 | 0.064 | 0.3464 | 0.5147 | 0.3424 | 0.3258 | 0.1968 | 0.6058 | 0.2697 | 0.5645 | 0.3113 | 0.2344 | 0.2059 | 0.5579 | 0.239 | 0.5457 | 0.2925 | 0.2158 | 0.2938 |
| MISC63 | 0.2187 | 0.0289 | 0.1761 | 0.1761 | 0.2652 | 0.1452 | 0.1676 | 0.1536 | 0.3339 | 0.183 | 0.3257 | 0.1916 | 0.1181 | 0.0533 | 0.2835 | 0.1948 | 0.315 | 0.1861 | 0.0953 | 0.0463 |
| MISC66 | 0.6783 | 0.3434 | 0.3543 | 0.2111 | 0.4587 | 0.0621 | 0.3951 | 0.2124 | 0.5455 | 0.2639 | 0.6866 | 0.5304 | -0.2673 | -0.134 | 0.4209 | 0.0687 | 0.7069 | 0.3131 | -0.0474 | 0.0541 |
| MISC67 | 0.8462 | 0.6095 | 0.5158 | 0.5157 | 0.6527 | 0.4932 | 0.5695 | 0.5656 | 0.8126 | 0.6061 | 0.8513 | 0.7838 | 0.0149 | 0.179 | 0.7581 | 0.6787 | 0.8562 | 0.6716 | 0.3675 | 0.2034 |
| MISC72 | 0.561 | 0.4802 | 0.4626 | 0.4626 | 0.5189 | 0.4574 | 0.5671 | 0.4448 | 0.5719 | 0.4631 | 0.5269 | 0.5537 | 0.3055 | 0.1137 | 0.5189 | 0.27 | 0.5223 | 0.4992 | 0.2607 | 0.1677 |
| MISC73 | 0.7614 | 0.7571 | 0.7292 | 0.7292 | 0.7308 | 0.6861 | 0.7749 | 0.6062 | 0.7297 | 0.6653 | 0.7661 | 0.7767 | 0.5497 | 0.4294 | 0.7054 | 0.7058 | 0.7842 | 0.7522 | 0.2532 | 0.2264 |
| NCBI629 | 0.2786 | 0.2541 | 0.1699 | 0.1699 | 0.044 | 0.2779 | 0.0569 | 0.25 | 0.1855 | 0.2365 | 0.2253 | 0.1486 | 0.1109 | 0.2104 | 0.2936 | 0.2509 | 0.3274 | 0.265 | 0.3329 | 0.2181 |
| NCBI631 | 0.2803 | 0.2313 | 0.2884 | 0.274 | 0.1447 | 0.2282 | 0.2541 | 0.2463 | 0.2028 | 0.1969 | 0.2272 | 0.2777 | 0.1537 | 0.1039 | 0.2933 | 0.2494 | 0.2316 | 0.1742 | 0.0971 | 0.2597 |
| NCBI632 | 0.4789 | 0.4343 | 0.4467 | 0.4467 | 0.468 | 0.4939 | 0.4864 | 0.4887 | 0.5212 | 0.4495 | 0.4753 | 0.3074 | 0.3299 | 0.3633 | 0.4993 | 0.3839 | 0.4431 | 0.4854 | 0.4869 | 0.4838 |
| NCBI635 | 0.4532 | 0.4378 | 0.3919 | 0.3919 | 0.4344 | 0.4089 | 0.3738 | 0.3918 | 0.5133 | 0.4683 | 0.4777 | 0.4236 | 0.3379 | 0.3475 | 0.4533 | 0.3571 | 0.3806 | 0.4554 | 0.2692 | 0.3496 |
| NCBI638 | -0.1006 | -0.0425 | -0.0839 | -0.0839 | -0.0436 | -0.0927 | -0.0171 | -0.1498 | -0.085 | -0.052 | -0.1339 | -0.005 | 0.1376 | 0.0718 | -0.0129 | 0.067 | -0.1391 | -0.0478 | -0.0273 | -0.0278 |
| NCBI640 | 0.324 | 0.202 | 0.233 | 0.233 | 0.2311 | 0.28 | 0.0453 | 0.1619 | 0.3015 | 0.2494 | 0.4167 | 0.3726 | 0.208 | 0.1048 | 0.3458 | 0.2327 | 0.3039 | 0.191 | 0.3315 | 0.2956 |
| NCBI681 | 0.6709 | 0.7078 | 0.7037 | 0.7037 | 0.6543 | | 0.6883 | 0.4838 | 0.6732 | 0.7185 | 0.6663 | 0.7233 | 0.4875 | 0.2625 | 0.6868 | 0.6885 | 0.7246 | 0.6916 | 0.5758 | 0.3736 |
| NCBI682 | 0.7138 | 0.6815 | 0.3965 | 0.653 | 0.6189 | 0.4214 | 0.6879 | 0.6766 | 0.7504 | 0.7089 | 0.7249 | 0.6962 | 0.6085 | 0.505 | 0.7152 | 0.6528 | 0.7177 | 0.6541 | 0.5609 | 0.5875 |
| NCBI683 | 0.6188 | 0.6121 | 0.5192 | 0.5192 | 0.5376 | 0.4287 | 0.5509 | 0.5154 | 0.5636 | 0.6014 | 0.5327 | 0.5625 | 0.2532 | 0.0555 | 0.5856 | 0.5333 | 0.5593 | 0.6421 | 0.511 | 0.3613 |
| NCBI760 | 0.3675 | 0.3758 | 0.2919 | 0.2919 | 0.3138 | 0.3255 | 0.4289 | 0.4005 | 0.3955 | 0.3809 | 0.3634 | 0.3964 | 0.3039 | 0.3246 | 0.3384 | 0.4118 | 0.3331 | 0.3423 | 0.3424 | 0.3067 |
| NCBI764 | 0.1851 | 0.1479 | 0.1542 | 0.1542 | 0.1376 | 0.1668 | 0.1575 | 0.1534 | 0.1845 | 0.1511 | 0.1715 | 0.157 | 0.1058 | 0.0669 | 0.1492 | 0.1532 | 0.1708 | 0.1599 | 0.0304 | -0.0465 |
| NCBI765 | 0.546 | 0.5567 | 0.5798 | 0.5798 | 0.4979 | 0.5055 | 0.5809 | 0.5866 | 0.5599 | 0.5862 | 0.5532 | 0.5862 | 0.4688 | 0.4496 | 0.5587 | 0.557 | 0.5071 | 0.5969 | 0.4273 | 0.4906 |
| NCBI766 | 0.4556 | 0.4431 | 0.4684 | 0.4684 | 0.4576 | 0.4788 | 0.445 | 0.4252 | 0.4341 | 0.4257 | 0.4234 | 0.4291 | 0.4137 | 0.4433 | 0.4673 | 0.4741 | 0.2199 | 0.4181 | 0.4406 | 0.4178 |
| NCBI768 | 0.6754 | 0.6632 | 0.694 | 0.694 | 0.7293 | 0.7275 | 0.6804 | 0.6537 | 0.684 | 0.6653 | 0.7108 | 0.6806 | 0.5791 | 0.5709 | 0.5332 | 0.6006 | 0.7733 | 0.6994 | 0.6208 | 0.6436 |
| NCBI769 | 0.6892 | | 0.7334 | 0.7334 | 0.6227 | 0.7707 | 0.7142 | 0.6773 | 0.7307 | 0.6977 | 0.6764 | 0.6535 | 0.5915 | 0.568 | 0.7249 | 0.6894 | 0.7561 | 0.7303 | 0.512 | 0.6617 |
| NCBI770 | -0.3857 | -0.2701 | -0.334 | -0.334 | -0.1143 | -0.1518 | -0.4147 | -0.3209 | -0.1426 | -0.3215 | -0.2888 | -0.2896 | -0.2893 | -0.2558 | -0.3897 | -0.3307 | -0.3147 | -0.3129 | -0.2806 | -0.2562 |
| SPA100 | 0.4413 | 0.6341 | 0.0205 | 0.0205 | 0.3316 | 0.3382 | 0.4874 | 0.4156 | 0.6666 | 0.6875 | 0.6193 | 0.5968 | -0.0261 | -0.0413 | 0.6546 | 0.6608 | 0.6434 | 0.6207 | 0.1477 | 0.192 |
| SPA101 | 0.4537 | 0.4045 | 0.3843 | 0.3843 | 0.2038 | 0.2648 | 0.4356 | 0.4483 | 0.4123 | 0.3981 | 0.4508 | 0.427 | 0.2622 | 0.3521 | 0.4761 | 0.4806 | 0.3955 | 0.4579 | 0.3034 | 0.3246 |
| SPA102 | 0.2924 | 0.2647 | 0.2281 | 0.2281 | 0.3141 | 0.259 | 0.2072 | 0.2032 | -0.0938 | 0.0678 | 0.2779 | 0.3288 | 0.2887 | 0.259 | 0.4 | 0.3974 | 0.4021 | 0.4171 | 0.2964 | |
| SPA103 | -0.2694 | -0.2966 | -0.3117 | -0.3117 | -0.3435 | -0.4015 | -0.3187 | -0.3399 | -0.3244 | -0.3297 | -0.3427 | -0.3313 | -0.1089 | -0.2082 | -0.3367 | -0.3478 | -0.3358 | -0.3555 | -0.14 | -0.0127 |
| SPA104 | 0.4312 | 0.44 | 0.2398 | 0.2398 | 0.321 | 0.2829 | 0.4197 | 0.3821 | 0.4191 | 0.3793 | 0.395 | 0.3697 | 0.3707 | 0.3109 | 0.3964 | 0.3468 | 0.3622 | 0.2976 | 0.3654 | 0.3322 |
| SPA105 | 0.2105 | 0.2144 | 0.1447 | 0.1447 | 0.0094 | 0.0498 | 0.2119 | 0.1995 | 0.176 | 0.1856 | 0.1264 | 0.0861 | 0.2053 | 0.146 | 0.2111 | 0.1691 | 0.232 | 0.228 | 0.0881 | 0.0929 |
| SPA106 | 0.163 | 0.1888 | 0.1418 | 0.1418 | 0.1274 | 0.0241 | 0.0132 | 0.0317 | | -0.0416 | -0.0248 | -0.0325 | 0.0522 | 0.0287 | -0.1212 | -0.0727 | 0.1561 | 0.1455 | -0.0105 | -0.0317 |
| SPA107 | 0.7687 | 0.7542 | 0.5573 | 0.5573 | 0.4341 | 0.4905 | 0.6682 | 0.711 | 0.7783 | 0.785 | 0.6005 | 0.694 | 0.2333 | 0.2455 | 0.5802 | 0.6621 | 0.719 | 0.7031 | -0.113 | 0.0885 |
| SPA108 | 0.5078 | 0.4757 | 0.4113 | 0.4113 | 0.4027 | 0.2834 | 0.3702 | 0.3794 | 0.4344 | 0.4056 | 0.3883 | 0.3859 | 0.4201 | 0.4542 | 0.4173 | 0.4009 | 0.4296 | 0.394 | 0.3972 | 0.4497 |
| SPA109 | 0.2476 | 0.2996 | 0.2934 | 0.2934 | 0.1159 | 0.083 | 0.1814 | 0.0853 | 0.137 | 0.0737 | | 0.0788 | 0.1148 | 0.1725 | 0.2097 | 0.2749 | 0.2271 | 0.2764 | 0.0089 | 0.0226 |
| SPA111 | 0.571 | 0.5485 | 0.514 | 0.514 | 0.4402 | 0.3599 | 0.5591 | 0.5647 | 0.5647 | 0.536 | 0.496 | 0.5197 | 0.4714 | 0.4361 | 0.5081 | 0.4464 | 0.5596 | 0.5717 | 0.4596 | 0.5195 |
| SPA112 | 0.6522 | 0.6308 | 0.5446 | 0.5446 | 0.4005 | 0.415 | 0.6078 | 0.579 | 0.6092 | 0.6059 | 0.6288 | 0.5999 | 0.3417 | 0.3708 | 0.4607 | 0.3342 | 0.5733 | 0.5561 | 0.5659 | 0.446 |
| SPA114 | 0.0043 | -0.0192 | 0.0017 | 0.0017 | -0.012 | 0.0298 | -0.0985 | -0.1393 | -0.0733 | 0.1382 | -0.2035 | 0.0064 | 0.067 | 0.0137 | -0.0016 | 0.0846 | 0.1018 | 0.0706 | -0.0445 | -0.1977 |
| SPA115 | 0.1018 | 0.0937 | 0.0966 | 0.0966 | 0.0914 | 0.0385 | 0.0645 | 0.0562 | 0.0566 | 0.0666 | 0.0661 | 0.051 | 0.1125 | 0.0896 | 0.1149 | 0.0781 | 0.1194 | 0.0974 | 0.1102 | 0.114 |
| SPA116 | 0.6164 | 0.4068 | 0.1575 | 0.1575 | 0.1331 | -0.4394 | -0.443 | -0.4263 | 0.6037 | 0.6306 | -0.2487 | -0.3382 | -0.4953 | -0.5058 | 0.6983 | 0.687 | 0.6486 | 0.6524 | -0.4463 | -0.3725 |
| SPA117 | -0.092 | -0.0546 | 0.516 | 0.516 | 0.3463 | 0.1798 | -0.184 | -0.1172 | 0.3449 | 0.0101 | -0.1407 | -0.1438 | -0.2772 | -0.2881 | -0.0897 | -0.053 | 0.616 | 0.6095 | -0.2547 | -0.2228 |
| SPA118 | 0.518 | 0.5109 | 0.5093 | 0.5093 | 0.3631 | 0.3533 | 0.4839 | 0.4689 | 0.5158 | 0.5217 | 0.5182 | 0.5054 | 0.4214 | 0.4544 | 0.499 | 0.4975 | 0.5075 | 0.5013 | 0.3821 | 0.414 |
| SPA55 | 0.3388 | 0.262 | 0.074 | 0.074 | 0.4014 | 0.325 | 0.3765 | 0.3834 | 0.2192 | 0.0276 | 0.3418 | 0.2188 | -0.0829 | -0.1673 | 0.4383 | 0.448 | 0.2166 | 0.075 | 0.3719 | 0.3066 |
| SPA57 | 0.5136 | 0.5709 | 0.2587 | 0.2587 | 0.3952 | 0.2879 | 0.5175 | 0.5389 | 0.5392 | 0.5246 | 0.5439 | 0.5775 | 0.2464 | 0.154 | 0.4899 | 0.4965 | 0.4323 | -0.0885 | 0.1675 | 0.0618 |
| SPA60 | 0.3368 | 0.5827 | 0.3486 | 0.3486 | -0.1962 | 0.0775 | 0.4364 | 0.4862 | 0.4566 | 0.529 | 0.2867 | 0.4449 | 0.0394 | -0.0203 | 0.4052 | 0.2151 | 0.5381 | 0.6157 | -0.0443 | -0.1609 |
| SPA61 | 0.8118 | 0.6071 | 0.4154 | 0.4154 | 0.0749 | 0.2594 | 0.1724 | 0.3743 | 0.5092 | 0.5546 | 0.2667 | 0.475 | 0.1031 | 0.091 | 0.0294 | 0.1431 | 0.7557 | 0.7433 | 0.1772 | 0.2234 |
| SPA64 | 0.3822 | 0.3363 | 0.2638 | 0.2638 | -0.2046 | -0.0518 | 0.117 | 0.0502 | 0.3214 | 0.2744 | 0.3123 | 0.3174 | -0.0277 | -0.2269 | 0.3491 | 0.3175 | 0.1772 | 0.3036 | -0.0471 | 0.0839 |
| SPA66 | 0.5527 | 0.4301 | -0.6031 | -0.6031 | -0.5824 | -0.6718 | 0.0005 | -0.5163 | 0.4274 | 0.4203 | 0.6193 | 0.4977 | -0.5126 | -0.3774 | 0.0992 | -0.4634 | 0.3897 | 0.2044 | -0.5228 | -0.4846 |
| SPA67 | 0.8022 | 0.6317 | 0.2891 | 0.2891 | -0.6997 | -0.6653 | -0.7248 | -0.7216 | 0.774 | 0.6398 | 0.7935 | 0.7352 | -0.6066 | -0.5676 | 0.5469 | 0.4741 | -0.4461 | -0.5041 | -0.1929 | -0.2163 |
| SPA68 | 0.7242 | 0.7357 | -0.1188 | -0.1188 | -0.475 | -0.6027 | -0.3553 | -0.2236 | 0.6061 | 0.5965 | 0.6106 | 0.5716 | -0.5299 | -0.5063 | 0.483 | 0.4482 | 0.1441 | -0.1862 | -0.5591 | -0.3752 |
| SPA69 | 0.5906 | 0.4942 | 0.4543 | 0.4543 | 0.4579 | 0.3762 | 0.5562 | 0.5912 | 0.6127 | 0.5832 | 0.4649 | 0.4114 | 0.4212 | 0.3848 | 0.634 | 0.5866 | 0.6588 | 0.5757 | 0.4132 | 0.3837 |
| SPA73 | 0.6572 | 0.6508 | 0.674 | 0.674 | 0.7068 | 0.6684 | 0.717 | 0.7117 | 0.6835 | 0.7036 | 0.6912 | 0.7022 | 0.6983 | 0.6762 | 0.6258 | 0.6547 | 0.6935 | 0.7023 | 0.734 | 0.7174 |
| SPA80 | 0.1057 | 0.1691 | 0.1335 | 0.1335 | 0.1285 | 0.0937 | 0.1517 | 0.1849 | 0.097 | 0.1762 | 0.1322 | 0.1703 | -0.0284 | -0.028 | 0.2822 | 0.1706 | 0.4148 | 0.1132 | 0.0008 | -0.0452 |
| SPA82 | -0.1686 | -0.0399 | -0.1956 | -0.1956 | -0.0793 | -0.1063 | -0.095 | -0.0977 | -0.157 | -0.1524 | -0.1383 | -0.1722 | 0.0391 | 0.0624 | -0.0915 | -0.2119 | -0.1521 | -0.2098 | -0.1202 | -0.0283 |
| SPA83 | 0.2554 | -0.0757 | -0.3118 | -0.3118 | -0.2664 | -0.3202 | -0.3751 | -0.4582 | 0.0477 | -0.0734 | 0.4602 | 0.4153 | -0.2319 | -0.2779 | 0.1547 | 0.0022 | -0.471 | -0.4841 | -0.4098 | -0.445 |
| SPA84 | 0.5396 | 0.5198 | 0.4108 | 0.4108 | 0.2126 | 0.5095 | 0.5561 | 0.5257 | 0.3459 | 0.3223 | 0.4229 | 0.4092 | 0.5265 | 0.5157 | 0.3782 | 0.3124 | 0.0565 | 0.2592 | 0.223 | 0.1081 |
| SPA90 | 0.5573 | 0.6846 | 0.5536 | 0.5536 | 0.4076 | 0.2599 | 0.5819 | 0.6448 | 0.5143 | 0.5942 | 0.4744 | 0.4893 | 0.2551 | 0.3151 | 0.5446 | 0.6062 | 0.6975 | 0.5642 | 0.3467 | 0.3764 |
| SPA91 | 0.6125 | 0.5274 | 0.5112 | 0.5112 | 0.356 | 0.3397 | 0.5504 | 0.564 | 0.4444 | 0.4846 | 0.4917 | 0.2751 | 0.5734 | 0.5322 | 0.3671 | 0.3452 | 0.7371 | 0.5168 | 0.4323 | 0.3697 |
| SPA92 | 0.522 | 0.4052 | -0.3377 | -0.3377 | 0.0483 | 0.039 | 0.507 | 0.4525 | 0.5143 | 0.3664 | 0.3712 | 0.3377 | 0.3286 | 0.3602 | 0.5751 | 0.5899 | 0.4841 | 0.4356 | -0.2816 | 0.2731 |
| SPA93 | 0.3931 | 0.4009 | 0.2232 | 0.2232 | 0.2283 | 0.1623 | 0.1488 | 0.1336 | 0.3487 | 0.3118 | 0.4379 | 0.4186 | 0.0749 | 0.0676 | 0.2363 | 0.2232 | | 0.1769 | 0.208 | 0.2752 |
| SPA94 | 0.4473 | 0.3944 | 0.0365 | 0.0365 | 0.1049 | 0.1044 | -0.1022 | -0.0952 | 0.348 | 0.3392 | 0.3717 | 0.3665 | -0.1181 | -0.0585 | 0.2241 | 0.2347 | 0.2584 | 0.2343 | 0.0361 | 0.0571 |
| SPA95 | 0.565 | 0.5459 | 0.4126 | 0.4126 | 0.4714 | 0.4437 | 0.5152 | 0.5127 | 0.5903 | 0.5906 | 0.5754 | 0.5351 | 0.4691 | 0.4462 | 0.5666 | 0.5282 | | 0.4502 | 0.4484 | 0.4329 |
| SPA96 | 0.4269 | 0.339 | 0.423 | 0.423 | 0.2464 | 0.2419 | 0.2688 | 0.2784 | 0.4631 | 0.3404 | 0.4502 | 0.3804 | 0.1621 | 0.2293 | 0.2726 | 0.2825 | 0.5043 | 0.4088 | 0.2656 | 0.3504 |
| SPA97 | 0.4618 | 0.4944 | 0.5682 | 0.5682 | 0.3888 | 0.4088 | 0.4789 | 0.5057 | 0.5033 | 0.5928 | 0.4623 | 0.4621 | 0.4342 | 0.4112 | 0.3361 | 0.2884 | 0.5624 | 0.562 | 0.0219 | 0.1078 |
| SPA98 | 0.5742 | 0.487 | 0.4625 | 0.4625 | 0.2537 | 0.3134 | 0.4229 | 0.4116 | 0.532 | 0.4571 | 0.6479 | 0.5813 | 0.3256 | 0.3141 | 0.1899 | 0.2385 | 0.5133 | 0.4113 | 0.1233 | 0.2195 |
| SPA99 | 0.4949 | 0.5137 | 0.495 | 0.495 | 0.3568 | 0.2937 | 0.3844 | 0.3718 | 0.5364 | 0.5328 | 0.5334 | 0.5405 | 0.1054 | 0.1373 | 0.5537 | 0.5403 | 0.5053 | 0.5222 | 0.335 | 0.4236 |
| TENX128 | 0.3925 | | | | 0.396 | | 0.3835 | | | | 0.3716 | | 0.3266 | | | | 0.4404 | | 0.2615 | |
| TENX147 | 0.6462 | 0.672 | 0.7258 | 0.7258 | 0.6698 | 0.6984 | 0.6271 | 0.6782 | 0.6418 | 0.6446 | 0.6747 | 0.6958 | 0.0961 | 0.0638 | 0.8003 | 0.7375 | 0.496 | 0.4282 | 0.4687 | 0.0583 |
| TENX148 | 0.7307 | 0.6806 | 0.5576 | 0.5576 | 0.6531 | 0.5797 | 0.7315 | 0.266 | 0.7033 | 0.3734 | 0.7421 | 0.5091 | 0.4582 | 0.236 | | 0.4265 | 0.7072 | 0.6928 | -0.039 | -0.2129 |
| TENX149 | 0.5021 | 0.6925 | 0.7422 | 0.7422 | 0.5224 | 0.6636 | 0.7669 | 0.7354 | 0.6119 | 0.7212 | | 0.7556 | 0.0756 | 0.2585 | 0.5089 | 0.7312 | 0.5677 | 0.6941 | 0.3941 | 0.2804 |
| TENX152 | 0.7305 | 0.6473 | 0.4836 | 0.4836 | 0.675 | 0.5487 | 0.6862 | 0.5526 | 0.7479 | 0.6858 | 0.6991 | 0.6314 | 0.4471 | 0.3711 | | 0.2876 | 0.7508 | 0.4631 | 0.5577 | 0.3133 |
| TENX157 | 0.6777 | 0.5912 | 0.5741 | 0.5741 | 0.6592 | 0.4872 | 0.5763 | 0.4204 | 0.6178 | 0.4868 | 0.668 | 0.6214 | 0.2612 | 0.216 | 0.6978 | 0.477 | 0.6703 | 0.4877 | 0.2422 | 0.1678 |
| TENX30 | 0.1389 | 0.1369 | 0.1786 | 0.1786 | 0.1717 | 0.1096 | 0.1245 | 0.068 | 0.1099 | 0.171 | 0.0487 | 0.1874 | 0.0801 | 0.0501 | 0.0699 | 0.0278 | 0.2361 | 0.0996 | 0.0802 | 0.0752 |
| TENX31 | 0.054 | -0.038 | 0.0613 | 0.0613 | 0.0463 | -0.0396 | 0.067 | 0.0046 | 0.0875 | 0.0585 | 0.1024 | 0.1586 | 0.0542 | 0.0159 | 0.0952 | -0.0412 | 0.1366 | -0.0513 | -0.0059 | -0.043 |
| TENX46 | 0.4933 | 0.4501 | 0.4663 | 0.4849 | 0.5019 | 0.339 | 0.6257 | 0.2868 | | | 0.5963 | | 0.0573 | 0.0703 | 0.6825 | 0.4041 | 0.3945 | | 0.2593 | 0.0438 |
| TENX70 | 0.7611 | | 0.7595 | 0.7588 | 0.8229 | 0.7598 | 0.7297 | 0.7415 | 0.7277 | 0.7643 | 0.7842 | 0.7838 | 0.5325 | 0.5793 | | 0.6061 | 0.8163 | 0.6833 | 0.6297 | 0.3 |
| TENX90 | 0.5766 | 0.5502 | 0.6184 | 0.6184 | 0.5105 | 0.5274 | 0.4802 | 0.4962 | 0.5611 | 0.6244 | 0.5509 | 0.6104 | 0.3126 | 0.0512 | 0.497 | 0.2435 | 0.5718 | 0.5022 | 0.3353 | 0.1264 |
| TENX91 | 0.8591 | 0.6118 | 0.6099 | 0.6099 | -0.1149 | -0.1851 | 0.5474 | 0.4394 | 0.7561 | 0.6333 | 0.7396 | 0.7642 | -0.1361 | 0.091 | 0.6626 | 0.2731 | 0.6962 | 0.6904 | -0.2 | 0.1977 |
| TENX92 | 0.7741 | 0.6893 | 0.6427 | 0.6427 | 0.7454 | 0.5988 | 0.7401 | 0.5381 | 0.7868 | 0.4153 | 0.7705 | 0.7788 | 0.3778 | 0.2871 | 0.6446 | 0.3586 | 0.7318 | 0.6639 | 0.2937 | 0.283 |
| ZEN38 | 0.6161 | 0.7988 | 0.8234 | 0.8234 | 0.8234 | 0.813 | 0.849 | 0.7423 | 0.7339 | 0.7944 | 0.7256 | 0.8345 | 0.6604 | -0.111 | 0.7165 | 0.8333 | 0.6687 | 0.5251 | 0.5962 | 0.0719 |
| ZEN42 | 0.7642 | 0.7113 | 0.6917 | 0.6917 | 0.7318 | 0.6999 | 0.779 | 0.7435 | 0.7614 | 0.6897 | 0.7192 | 0.7301 | 0.6213 | 0.3754 | 0.7871 | 0.7257 | 0.7237 | 0.7181 | 0.7183 | 0.3949 |
| ZEN44 | 0.8074 | 0.7433 | 0.7424 | 0.7424 | 0.7342 | 0.7263 | 0.7269 | 0.7319 | 0.7771 | 0.7483 | 0.7768 | 0.7305 | 0.5731 | 0.4877 | 0.7721 | 0.7662 | 0.7839 | 0.6585 | 0.5655 | 0.6519 |
| ZEN45 | 0.5722 | 0.7637 | 0.6673 | 0.6673 | 0.7058 | 0.7042 | 0.6211 | 0.7733 | 0.6785 | 0.7475 | 0.5294 | 0.7699 | 0.6824 | 0.4505 | 0.6335 | 0.6796 | 0.6542 | 0.6955 | 0.4681 | 0.6625 |
| ZEN78 | 0.3668 | 0.6231 | 0.4688 | 0.4688 | 0.429 | 0.2592 | 0.1782 | 0.5029 | 0.3262 | 0.5642 | 0.2737 | 0.3094 | 0.5382 | 0.5411 | 0.3981 | 0.4522 | 0.0207 | 0.5321 | 0.6378 | 0.572 |
| ZEN80 | 0.7396 | 0.7496 | 0.7689 | 0.7688 | 0.7353 | 0.7139 | 0.776 | 0.718 | 0.8044 | 0.7762 | 0.7952 | 0.758 | 0.7195 | 0.7203 | 0.7811 | 0.7178 | 0.7719 | 0.7181 | 0.7344 | 0.7407 |
| ZEN81 | 0.6718 | 0.6858 | 0.6274 | 0.6274 | 0.558 | 0.5509 | 0.679 | 0.6861 | 0.6847 | 0.6803 | 0.6642 | 0.6478 | -0.5305 | 0.6567 | 0.6465 | 0.6678 | | 0.5708 | 0.5721 | 0.6113 |
| INT12 | | 0.386 | 0.4945 | 0.4945 | 0.3815 | 0.4638 | 0.3662 | 0.0711 | 0.6495 | 0.5791 | 0.4532 | 0.3904 | 0.1756 | 0.059 | 0.652 | 0.3914 | | 0.3223 | 0.1975 | -0.0221 |
| MEND148 | | 0.5531 | 0.2641 | 0.2641 | 0.3081 | 0.361 | 0.4721 | 0.4395 | 0.5778 | 0.5684 | 0.6029 | 0.5307 | 0.198 | 0.491 | 0.5631 | 0.5106 | 0.4588 | 0.5569 | 0.5103 | 0.5347 |
| MEND65 | | 0.3012 | 0.1613 | 0.1613 | 0.0178 | 0.1541 | 0.2677 | 0.0297 | | 0.1598 | 0.3514 | 0.2458 | 0.0773 | 0.1035 | 0.3155 | 0.1864 | 0.2508 | 0.1733 | 0.046 | 0.0801 |
| MISC42 | | 0.6828 | 0.6785 | 0.6785 | 0.7937 | 0.6842 | 0.5615 | 0.413 | | 0.4082 | | 0.7574 | 0.4733 | 0.4177 | 0.8127 | 0.6231 | 0.8017 | 0.6356 | 0.5776 | 0.4781 |
| MISC57 | | -0.1106 | 0.2936 | 0.2936 | 0.5015 | 0.4872 | -0.2577 | -0.2595 | -0.3624 | 0.5551 | | -0.2429 | 0.3168 | 0.4787 | | 0.4745 | | 0.0776 | -0.1345 | 0.455 |
| MISC58 | | 0.413 | 0.0084 | 0.0084 | 0.3297 | 0.0525 | 0.4163 | 0.2512 | 0.7562 | 0.2307 | 0.7986 | 0.5046 | -0.6011 | -0.3219 | 0.6284 | 0.2044 | | 0.3399 | 0.3408 | -0.1544 |
| TENX40 | | 0.8496 | 0.6173 | 0.7351 | 0.8288 | 0.7366 | 0.8489 | 0.7533 | 0.8798 | 0.8452 | 0.871 | 0.8649 | 0.6904 | 0.5521 | 0.8673 | 0.8189 | | 0.8043 | 0.5106 | 0.3353 |
| ZEN79 | | 0.7019 | 0.7458 | 0.7458 | 0.7054 | 0.679 | 0.7349 | 0.7538 | 0.757 | 0.7502 | 0.7951 | 0.7419 | | 0.7403 | | 0.7741 | 0.7774 | 0.7502 | 0.646 | 0.7087 |
| MEND156 | | | 0.7266 | 0.7266 | 0.758 | 0.748 | 0.3692 | 0.7599 | 0.7658 | | 0.6827 | 0.8114 | 0.6101 | 0.6047 | 0.7802 | 0.7992 | | 0.6397 | 0.721 | 0.6395 |
| MEND141 | | | | | | 0.4462 | | | | | | | 0.354 | 0.4188 | | | | 0.5052 | | 0.1942 |

### Table.S4

| wasserstein_distance | UNI_big_region | UNI_small_region | CONCH_big_region | CONCH_small_region | CONCH15_big_region | CONCH15_small_region | CTransPath_big_region | CTransPath_small_region | GigaPath_big_region | GigaPath_small_region | UNI2h_big_region | UNI2h_small_region | RN18_big_region | RN18_small_region | Virchow_big_region | Virchow_small_region | Virchow2_big_region | Virchow2_small_region | ViTB16_big_region | ViTB16_small_region |
|---|---|---|---|---|---|---|---|---|---|---|---|---|---|---|---|---|---|---|---|---|
| INT13 | 0.2132 | 0.2541 | 0.1526 | 0.2208 | 0.423 | 0.1963 | 0.414 | 0.3825 | 0.2322 | 0.2241 | 0.1438 | 0.1841 | 0.434 | 0.3136 | 0.2511 | 0.2713 | 0.253 | 0.1627 | 0.3253 | 0.2435 |
| INT14 | 0.4154 | 0.3388 | 0.3408 | 0.3408 | 0.4151 | 0.3214 | 0.3608 | 0.2569 | 0.3414 | 0.3153 | 0.3576 | 0.3694 | 0.4416 | 0.4382 | 0.3069 | 0.2746 |  | 0.2706 | 0.4012 | 0.4063 |
| INT15 | 0.2882 | 0.264 | 0.3665 | 0.3664 | 0.3822 | 0.2969 | 0.3065 | 0.1618 | 0.2713 | 0.2626 | 0.311 | 0.2923 | 0.2621 | 0.2317 | 0.2559 | 0.2274 | 0.3037 | 0.2941 | 0.2893 | 0.2219 |
| INT16 | 0.2392 | 0.2423 | 0.1522 | 0.1522 | 0.2516 | 0.1479 | 0.1063 | 0.4811 | 0.2664 | 0.2397 | 0.2737 | 0.2451 | 0.2909 | 0.3701 | 0.1759 | 0.2415 | 0.2575 | 0.1924 | 0.2618 | 0.2241 |
| INT19 | 0.5972 | 0.6346 | 0.4751 | 0.5833 | 0.5028 | 0.5135 | 0.6176 | 0.6484 | 0.622 | 0.6439 | 0.554 | 0.5386 | 0.6052 | 0.4794 | 0.5471 | 0.5258 | 0.5853 | 0.5952 | 0.6882 | 0.6817 |
| INT21 | 0.325 | 0.3676 | 0.2932 | 0.2932 | 0.3666 | 0.1838 | 0.3512 | 0.5874 | 0.3393 | 0.3385 | 0.3143 | 0.3265 | 0.3342 | 0.477 | 0.3134 | 0.2899 | 0.3377 | 0.2209 | 0.3545 | 0.2525 |
| INT22 | 0.3453 | 0.2178 | 0.1884 | 0.1878 | 0.3909 | 0.3853 | 0.4433 | 0.3935 | 0.3858 | 0.3263 | 0.1849 | 0.1637 | 0.0674 | 0.2206 | 0.3314 | 0.1427 | 0.3323 | 0.4422 | 0.3887 | 0.3626 |
| INT24 | 0.2012 | 0.1505 | 0.3743 | 0.3743 | 0.2425 | 0.2609 | 0.0813 | 0.2929 | 0.2052 | 0.1491 | 0.208 | 0.2157 | 0.4552 | 0.4028 | 0.19 | 0.1312 | 0.2136 | 0.2382 | 0.3122 | 0.4171 |
| INT25 | 0.2329 | 0.423 | 0.2941 | 0.4504 | 0.3423 | 0.323 | 0.2126 | 0.4078 | 0.1876 | 0.379 | 0.1529 | 0.3102 | 0.2013 | 0.2892 | 0.2047 | 0.5122 | 0.21 | 0.289 | 0.2722 | 0.2949 |
| INT26 | 0.1883 | 0.2568 | 0.2006 | 0.2005 | 0.1263 | 0.1688 | 0.1641 | 0.2088 | 0.1207 | 0.1702 | 0.1491 | 0.1742 | 0.1159 | 0.1851 | 0.1308 | 0.3978 | 0.2591 | 0.1324 | 0.1037 | 0.2021 |
| INT27 | 0.2614 | 0.2931 | 0.3625 | 0.3625 | 0.2842 | 0.263 | 0.2281 | 0.2617 | 0.2748 | 0.2036 | 0.356 | 0.3156 | 0.3813 | 0.3357 | 0.1224 | 0.4642 | 0.1441 |  | 0.2569 | 0.2218 |
| INT28 | 0.3425 | 0.1168 | 0.2416 | 0.2416 | 0.215 | 0.1486 | 0.1085 | 0.1331 | 0.2661 | 0.1775 | 0.3607 | 0.2086 | 0.2591 | 0.2372 | 0.1166 | 0.1781 | 0.1878 | 0.207 | 0.1798 | 0.1424 |
| INT35 | 0.7067 | 0.2936 | 0.2108 | 0.2106 | 0.4535 | 0.116 | 0.6445 | 0.0861 | 0.6417 | 0.2182 |  | 0.4222 | 0.0785 | 0.1765 | 0.4728 | 0.2101 | 0.4866 | 0.2168 | 0.3596 | 0.2269 |
| INT4 | 0.1073 | 0.1454 | 0.1415 | 0.1415 | 0.1821 | 0.1809 | 0.1061 | 0.0608 | 0.1939 | 0.2017 | 0.1052 | 0.1871 | 0.1701 | 0.1748 | 0.0817 | 0.1892 | 0.2799 | 0.2237 | 0.1292 | 0.1782 |
| INT5 | 0.4035 | 0.3789 | 0.3945 | 0.3945 | 0.2709 | 0.2557 | 0.4207 | 0.402 | 0.4244 | 0.3605 | 0.3755 | 0.3803 | 0.1923 | 0.2817 | 0.3128 | 0.3171 | 0.232 | 0.464 | 0.294 | 0.3271 |
| INT7 | 0.4636 | 0.5746 | 0.0913 | 0.0913 | 0.4567 | 0.1395 | 0.5667 | 0.443 | 0.6305 | 0.5515 | 0.317 | 0.2985 | 0.4733 | 0.1523 | 0.5033 | 0.3755 | 0.3215 | 0.1265 | 0.578 | 0.2678 |
| INT8 | 0.6307 | 0.6411 | 0.4678 | 0.4678 | 0.5102 | 0.4551 | 0.5063 | 0.5254 | 0.6343 | 0.5664 | 0.5908 | 0.5511 | 0.3265 | 0.2546 | 0.5398 | 0.4767 | 0.5785 | 0.5516 | 0.5149 | 0.4869 |
| MEND140 | 0.4973 | 0.3287 | 0.2255 | 0.2255 | 0.4889 | 0.3076 | 0.4561 | 0.266 | 0.2653 | 0.2033 | 0.5375 | 0.2503 | 0.3174 | 0.2762 | 0.4446 | 0.2159 | 0.4464 | 0.2495 | 0.5118 | 0.4457 |
| MEND142 | 0.301 | 0.1282 | 0.1631 | 0.1631 | 0.2916 | 0.0818 | 0.19 | 0.161 | 0.2989 | 0.0876 | 0.2793 | 0.1169 | 0.1308 | 0.1146 | 0.1716 | 0.0927 | 0.2767 | 0.0798 | 0.0877 | 0.1745 |
| MEND143 | 0.2173 | 0.2617 | 0.2937 | 0.2937 | 0.2981 | 0.4813 | 0.416 | 0.1042 | 0.3356 | 0.2875 | 0.3618 | 0.2858 | 0.5063 | 0.6032 | 0.2758 | 0.421 | 0.234 | 0.3703 | 0.5835 | 0.5954 |
| MEND145 | 0.1887 | 0.1305 | 0.0811 | 0.0811 | 0.1804 | 0.1123 | 0.1459 | 0.1046 | 0.1715 | 0.1428 | 0.1871 | 0.1244 | 0.365 | 0.1357 | 0.1491 | 0.0904 | 0.1705 | 0.0811 | 0.3909 | 0.102 |
| MEND146 | 0.1724 | 0.3318 | 0.1532 | 0.1532 | 0.4099 | 0.2447 | 0.1157 | 0.3283 | 0.393 | 0.2954 | 0.231 | 0.3692 | 0.4563 | 0.2745 | 0.4153 | 0.4332 | 0.3745 | 0.4399 | 0.3082 | 0.4326 |
| MEND147 | 0.3533 | 0.2204 | 0.2596 | 0.2596 | 0.196 | 0.2776 | 0.2875 | 0.1379 | 0.24 | 0.1868 | 0.3343 | 0.316 | 0.4203 | 0.3717 | 0.3357 | 0.15 | 0.2374 | 0.1332 | 0.1816 | 0.2269 |
| MEND150 | 0.2876 | 0.1846 | 0.1845 | 0.1845 | 0.3033 | 0.1463 | 0.247 | 0.2039 | 0.2332 | 0.158 | 0.2154 | 0.2123 | 0.3273 | 0.284 | 0.2262 | 0.2417 | 0.2098 | 0.2032 | 0.2512 | 0.3263 |
| MEND151 | 0.5422 | 0.3791 | 0.252 | 0.3663 | 0.4716 | 0.3251 | 0.583 |  | 0.4639 | 0.495 | 0.5203 | 0.3791 | 0.4622 | 0.2726 | 0.4985 | 0.4666 | 0.5292 | 0.3505 | 0.4012 | 0.3768 |
| MEND153 | 0.2197 | 0.1819 | 0.3554 | 0.3554 | 0.1676 | 0.2998 | 0.155 | 0.1687 | 0.2141 | 0.443 | 0.2432 | 0.22 | 0.229 | 0.3761 | 0.183 | 0.2213 | 0.214 | 0.1501 | 0.26 | 0.2272 |
| MEND154 | 0.1053 | 0.1437 | 0.1609 | 0.3266 | 0.2836 | 0.2408 | 0.2273 | 0.0639 | 0.1169 | 0.3237 | 0.1788 | 0.2385 | 0.1541 | 0.2276 | 0.1395 | 0.313 | 0.3402 | 0.3061 | 0.2944 | 0.1539 |
| MEND157 | 0.4459 | 0.113 | 0.1252 | 0.1643 | 0.4043 | 0.1662 | 0.3333 | 0.1682 | 0.2772 | 0.0843 | 0.4768 | 0.1533 | 0.2903 | 0.143 | 0.2903 | 0.0798 | 0.4326 | 0.1491 | 0.2847 | 0.1468 |
| MEND158 | 0.2697 | 0.183 | 0.1056 | 0.1782 | 0.2119 | 0.185 | 0.3721 | 0.2503 | 0.2761 | 0.0921 | 0.1616 | 0.1324 | 0.3106 | 0.0487 | 0.1225 | 0.1798 | 0.4188 | 0.2749 | 0.5606 | 0.1165 |
| MEND160 | 0.3763 | 0.2395 | 0.2696 | 0.262 | 0.364 | 0.1703 | 0.343 | 0.2218 | 0.3371 | 0.2376 | 0.4438 | 0.2931 | 0.3556 | 0.1573 | 0.3266 | 0.1994 | 0.4154 | 0.2914 | 0.3224 | 0.1979 |
| MEND162 | 0.3532 | 0.4338 | 0.3054 | 0.3055 | 0.3307 | 0.3293 | 0.4095 | 0.4048 | 0.4471 | 0.3353 | 0.3405 | 0.2996 | 0.2279 | 0.2759 | 0.3549 | 0.2453 | 0.3056 | 0.2146 | 0.2376 | 0.1931 |
| MEND18 | 0.1426 | 0.0831 | 0.1877 | 0.1877 | 0.1878 | 0.1489 | 0.3146 | 0.4335 | 0.1686 | 0.1686 | 0.1736 | 0.1584 | 0.095 | 0.0708 | 0.1677 | 0.2354 | 0.1388 | 0.1499 | 0.3288 | 0.3351 |
| MEND19 | 0.1216 | 0.1329 | 0.0742 | 0.0742 | 0.1422 | 0.1369 | 0.148 | 0.1492 | 0.1305 | 0.1329 | 0.19 | 0.243 | 0.102 | 0.1126 | 0.0995 | 0.115 | 0.1007 | 0.0962 | 0.1405 | 0.2159 |
| MEND24 | 0.2347 | 0.2096 | 0.5333 | 0.5781 | 0.4666 | 0.5789 | 0.6948 | 0.6793 | 0.5669 | 0.5227 | 0.3402 | 0.3631 | 0.3874 | 0.3374 | 0.2123 | 0.3011 | 0.2101 | 0.2574 | 0.6172 | 0.6678 |
| MEND37 | 0.5668 | 0.5528 | 0.572 | 0.572 | 0.6024 | 0.6306 | 0.583 | 0.6163 | 0.5873 | 0.4299 | 0.6205 | 0.4388 | 0.4251 | 0.5634 | 0.6413 | 0.5684 | 0.545 | 0.5633 | 0.5648 | 0.5632 |
| MEND38 | 0.3631 | 0.1509 | 0.2422 | 0.2422 | 0.2995 | 0.3155 | 0.229 | 0.1759 | 0.3594 | 0.3118 | 0.5429 | 0.3566 | 0.2592 | 0.2715 | 0.3987 | 0.2153 | 0.2253 | 0.2066 | 0.3313 | 0.3183 |
| MEND39 | 0.3375 | 0.3615 | 0.4114 | 0.4476 | 0.3119 | 0.3461 | 0.4235 | 0.4233 | 0.4078 | 0.3418 | 0.4441 | 0.3402 | 0.2787 | 0.4006 | 0.4847 | 0.3936 | 0.3426 | 0.2464 | 0.4448 | 0.5883 |
| MEND40 | 0.4677 | 0.3223 | 0.3487 | 0.3487 | 0.3206 | 0.3682 | 0.44 | 0.3943 | 0.5123 | 0.3236 | 0.5513 | 0.1844 | 0.246 | 0.4812 | 0.4671 | 0.2811 | 0.3322 | 0.2376 | 0.3057 | 0.3713 |
| MEND59 | 0.1638 | 0.3119 | 0.3389 | 0.339 | 0.122 | 0.3111 | 0.1096 | 0.3206 | 0.1715 | 0.3299 | 0.149 | 0.3813 | 0.1942 | 0.3409 |  | 0.3164 | 0.1783 | 0.4573 | 0.2794 | 0.368 |
| MEND60 | 0.1669 | 0.2687 | 0.2559 | 0.2559 | 0.1541 | 0.4548 | 0.1807 | 0.2452 | 0.2269 | 0.2814 | 0.1827 | 0.3344 | 0.2094 | 0.2119 | 0.1358 | 0.4078 | 0.2746 | 0.2914 | 0.2724 | 0.3629 |
| MEND61 | 0.0398 | 0.2387 | 0.5543 | 0.5543 | 0.2662 | 0.4211 | 0.1042 | 0.3305 | 0.065 | 0.3461 | 0.0619 | 0.3425 | 0.3347 | 0.3826 | 0.0691 | 0.2246 | 0.1608 | 0.4892 | 0.312 | 0.3882 |
| MEND62 | 0.1172 | 0.2438 | 0.4471 | 0.4471 | 0.3097 | 0.3081 | 0.1388 | 0.3115 | 0.1233 | 0.2878 | 0.2067 | 0.3683 | 0.2623 | 0.401 | 0.1291 | 0.2203 | 0.1347 | 0.263 | 0.3561 | 0.3744 |
| MEND66 | 0.3023 | 0.3507 | 0.4257 | 0.4257 | 0.3962 | 0.4513 | 0.3232 | 0.5141 | 0.247 | 0.3402 | 0.2659 | 0.3871 | 0.5541 | 0.4793 | 0.3731 | 0.4986 | 0.303 | 0.5638 | 0.5504 | 0.5888 |
| MISC33 | 0.278 |  | 0.3941 | 0.2071 | 0.312 | 0.2159 | 0.1514 | 0.096 | 0.2482 | 0.2402 | 0.2331 | 0.2941 | 0.2363 | 0.2773 | 0.1751 | 0.1896 | 0.3056 | 0.1324 | 0.1183 | 0.1565 |
| MISC35 | 0.3232 | 0.1793 | 0.2367 | 0.2367 | 0.3015 | 0.185 | 0.2167 | 0.4921 | 0.3195 | 0.1247 | 0.3334 | 0.2373 | 0.6143 | 0.5115 | 0.2993 | 0.1455 |  | 0.2134 | 0.2247 | 0.4378 |
| MISC36 | 0.2735 | 0.1844 | 0.2113 | 0.2113 | 0.2436 | 0.1576 | 0.3076 | 0.3355 | 0.2726 | 0.1674 | 0.2661 | 0.1488 | 0.394 | 0.4407 | 0.2582 | 0.233 | 0.327 | 0.2335 | 0.3754 | 0.4033 |
| MISC40 | 0.3409 | 0.2988 | 0.2916 | 0.2916 | 0.2329 | 0.2151 | 0.5778 | 0.5391 | 0.2503 | 0.3254 | 0.3502 | 0.3858 | 0.2934 | 0.4554 | 0.3124 | 0.4564 | 0.4272 | 0.395 | 0.4003 | 0.3919 |
| MISC41 | 0.4766 | 0.3483 | 0.4219 | 0.4219 | 0.5029 | 0.4326 | 0.479 | 0.2593 | 0.4353 | 0.3496 | 0.4595 | 0.4358 | 0.258 | 0.2707 | 0.3914 | 0.3693 |  | 0.2361 | 0.3451 | 0.2724 |
| MISC45 | 0.3183 | 0.2285 | 0.3353 | 0.3353 | 0.3532 | 0.3038 | 0.4003 | 0.2568 | 0.4022 | 0.3334 | 0.3802 | 0.3655 | 0.2334 | 0.2236 | 0.4169 | 0.3331 | 0.4364 | 0.4644 | 0.3847 | 0.248 |
| MISC47 | 0.3887 | 0.5818 | 0.5771 | 0.5771 | 0.326 | 0.6138 | 0.267 | 0.6022 | 0.3204 | 0.6205 | 0.3742 | 0.4543 | 0.6056 | 0.6071 | 0.3203 | 0.479 | 0.5087 | 0.5565 | 0.2899 | 0.6574 |
| MISC48 | 0.2905 | 0.2559 | 0.2376 | 0.2376 | 0.3373 | 0.3494 | 0.3022 | 0.2366 | 0.2886 | 0.2289 | 0.3301 | 0.2623 | 0.2451 | 0.2196 | 0.4026 | 0.326 | 0.301 | 0.3317 | 0.1911 | 0.1611 |
| MISC49 | 0.3708 | 0.4393 | 0.4569 | 0.4569 | 0.5269 | 0.4621 | 0.3749 | 0.3163 | 0.4191 | 0.4741 | 0.4091 | 0.4973 | 0.4753 | 0.4144 | 0.3444 | 0.3652 | 0.3361 | 0.3085 | 0.5585 | 0.4619 |
| MISC50 | 0.5369 | 0.3577 | 0.1455 | 0.1455 | 0.5457 | 0.3086 | 0.4733 | 0.3207 | 0.4339 | 0.2492 | 0.5052 | 0.3257 | 0.1735 | 0.4272 | 0.4524 | 0.3595 | 0.472 | 0.1703 | 0.3982 | 0.4571 |
| MISC62 | 0.3173 | 0.2664 | 0.4689 | 0.241 | 0.3634 | 0.2434 | 0.3658 | 0.291 | 0.377 | 0.3543 | 0.4693 | 0.3117 | 0.2721 | 0.2068 | 0.2791 | 0.2782 | 0.3179 | 0.2911 | 0.245 | 0.2553 |
| MISC63 | 0.4769 | 0.2594 | 0.2605 | 0.2605 | 0.4105 | 0.268 | 0.2864 | 0.2825 | 0.3373 | 0.2312 | 0.5025 | 0.3486 | 0.1456 | 0.134 | 0.3033 | 0.1601 | 0.3753 | 0.3123 | 0.3741 | 0.3455 |
| MISC66 | 0.2499 | 0.4573 | 0.2528 | 0.2363 | 0.3222 | 0.1798 | 0.4729 | 0.3066 | 0.2798 | 0.2365 | 0.1897 | 0.3151 | 0.1692 | 0.218 | 0.3283 | 0.3707 | 0.1464 | 0.2297 | 0.2061 | 0.2032 |
| MISC67 | 0.0751 | 0.3053 | 0.49 | 0.49 | 0.408 | 0.3973 | 0.4622 | 0.5041 | 0.2125 | 0.4041 | 0.1012 | 0.2326 | 0.5477 | 0.4539 | 0.2604 | 0.3989 | 0.0946 | 0.2805 | 0.4058 | 0.4741 |
| MISC72 | 0.2626 | 0.1609 | 0.1161 | 0.1161 | 0.2317 | 0.0572 | 0.1995 | 0.1288 | 0.2364 | 0.0765 | 0.2759 | 0.1671 | 0.1613 | 0.1348 | 0.2341 | 0.1286 | 0.2634 | 0.1479 | 0.2815 | 0.1166 |
| MISC73 | 0.1493 | 0.2185 | 0.2794 | 0.2794 | 0.2141 | 0.2152 | 0.1441 | 0.3379 | 0.1512 | 0.199 | 0.143 | 0.1951 | 0.3851 | 0.351 | 0.1417 | 0.2289 | 0.2042 | 0.1352 | 0.4181 | 0.3547 |
| NCBI629 | 0.3715 | 0.4301 | 0.3301 | 0.3301 | 0.144 | 0.3254 | 0.4056 | 0.4479 | 0.3433 | 0.3929 | 0.3314 | 0.3898 | 0.2666 | 0.227 | 0.328 | 0.2974 | 0.3826 | 0.3971 | 0.3088 | 0.2688 |
| NCBI631 | 0.5966 | 0.5886 | 0.6112 | 0.5928 | 0.6546 | 0.6815 | 0.5157 | 0.6128 | 0.6026 | 0.6608 | 0.6329 | 0.5913 | 0.5336 | 0.6203 | 0.5599 | 0.6478 | 0.5934 | 0.5756 | 0.552 | 0.5948 |
| NCBI632 | 0.416 | 0.4022 | 0.4074 | 0.4074 | 0.3649 | 0.451 | 0.4362 | 0.4067 | 0.4384 | 0.4127 | 0.4217 | 0.3847 | 0.4245 | 0.4103 | 0.4319 | 0.4423 | 0.4431 | 0.3822 | 0.4368 | 0.4428 |
| NCBI635 | 0.492 | 0.3326 | 0.4427 | 0.4427 | 0.3574 | 0.2498 | 0.2285 | 0.2652 | 0.558 | 0.4522 | 0.5465 | 0.4572 | 0.2887 | 0.0925 | 0.4923 | 0.3791 | 0.3965 | 0.289 | 0.3424 | 0.2731 |
| NCBI638 | 0.4683 | 0.5651 | 0.3828 | 0.3828 | 0.3392 | 0.4925 | 0.3227 | 0.4938 | 0.5519 | 0.571 | 0.35 | 0.5305 | 0.4393 | 0.4112 | 0.5581 | 0.5651 | 0.3785 | 0.5524 | 0.4249 | 0.496 |
| NCBI640 | 0.2831 | 0.266 | 0.1412 | 0.1412 | 0.1168 | 0.1263 | 0.3672 | 0.1852 | 0.1128 | 0.157 | 0.179 | 0.1725 | 0.2162 | 0.1868 | 0.2076 | 0.1872 | 0.3192 | 0.2283 | 0.3969 | 0.2394 |
| NCBI681 | 0.313 | 0.2512 | 0.2807 | 0.2807 | 0.3395 |  | 0.2884 | 0.3464 | 0.3307 | 0.221 | 0.3684 | 0.2471 | 0.3413 | 0.5031 | 0.2267 | 0.3095 | 0.1378 | 0.3294 | 0.3939 | 0.3945 |
| NCBI682 | 0.2128 | 0.1087 | 0.0685 | 0.1296 | 0.2206 | 0.1124 | 0.1655 | 0.1636 | 0.1141 | 0.1183 | 0.2093 | 0.1765 | 0.2967 | 0.2693 | 0.2277 | 0.1469 | 0.3882 | 0.1845 | 0.1616 | 0.2437 |
| NCBI683 | 0.7063 | 0.6927 | 0.6778 | 0.6778 | 0.6932 | 0.6836 | 0.7057 | 0.7106 | 0.7027 | 0.6975 | 0.7136 | 0.7045 | 0.7345 | 0.6766 | 0.6825 | 0.6486 | 0.6753 | 0.6132 | 0.6663 | 0.5953 |
| NCBI760 | 0.3815 | 0.4273 | 0.4377 | 0.4377 | 0.4821 | 0.5035 | 0.3999 | 0.3515 | 0.3561 | 0.4278 | 0.4329 | 0.4558 | 0.3756 | 0.3735 | 0.5514 | 0.483 | 0.3409 | 0.3588 | 0.4375 | 0.3412 |
| NCBI764 | 0.1626 | 0.2131 | 0.0882 | 0.0882 | 0.231 | 0.158 | 0.0792 | 0.1266 | 0.1283 | 0.1323 | 0.1139 | 0.172 | 0.111 | 0.0729 | 0.088 | 0.1455 | 0.095 | 0.1472 | 0.097 | 0.1494 |
| NCBI765 | 0.3083 | 0.3621 | 0.3082 | 0.3082 | 0.1453 | 0.2465 | 0.2611 | 0.2514 | 0.2403 | 0.2327 | 0.3013 | 0.3258 | 0.2293 | 0.2597 | 0.2292 | 0.1976 | 0.2752 | 0.2013 | 0.1994 | 0.2083 |
| NCBI766 | 0.4309 | 0.4373 | 0.2848 | 0.2848 | 0.3184 | 0.3019 | 0.2913 | 0.265 | 0.2267 | 0.3122 | 0.4274 | 0.3667 | 0.433 | 0.3907 | 0.3477 | 0.3522 | 0.3605 | 0.366 | 0.2849 | 0.2921 |
| NCBI768 | 0.2398 | 0.2948 | 0.2521 | 0.2521 | 0.2645 | 0.2736 | 0.3089 | 0.2427 | 0.3165 | 0.3267 | 0.3245 | 0.3095 | 0.379 | 0.33 | 0.3679 | 0.3313 | 0.3444 | 0.3491 | 0.2483 | 0.3081 |
| NCBI769 | 0.1529 |  | 0.1718 | 0.1718 | 0.1959 | 0.1461 | 0.2206 | 0.1924 | 0.2051 | 0.1965 | 0.2127 | 0.154 | 0.1361 | 0.1776 | 0.247 | 0.2369 | 0.2077 | 0.2306 | 0.1831 | 0.2573 |
| NCBI770 | 0.4241 | 0.3834 | 0.4255 | 0.4255 | 0.5216 | 0.4852 | 0.4578 | 0.3397 | 0.4294 | 0.375 | 0.448 | 0.451 | 0.4159 | 0.4137 | 0.5041 | 0.4655 | 0.4125 | 0.388 | 0.1587 | 0.2023 |
| SPA100 | 0.1115 | 0.1031 | 0.3496 | 0.3496 | 0.18 | 0.1827 | 0.1794 | 0.1826 | 0.0466 | 0.0429 | 0.2332 | 0.2816 | 0.452 | 0.4301 | 0.1 | 0.0961 | 0.1146 | 0.1216 | 0.2669 | 0.3189 |
| SPA101 | 0.1034 | 0.1308 | 0.0553 | 0.0553 | 0.2779 | 0.1814 | 0.1725 | 0.1093 | 0.0607 | 0.0602 | 0.1801 | 0.1634 | 0.092 | 0.2044 | 0.0961 | 0.0937 | 0.0789 | 0.1309 | 0.1551 | 0.0796 |
| SPA102 | 0.6393 | 0.6297 | 0.6603 | 0.6603 | 0.5697 | 0.3967 | 0.51 | 0.4938 | 0.6004 | 0.6239 | 0.57 | 0.5703 | 0.6307 | 0.6256 | 0.6618 | 0.6454 | 0.7051 | 0.6179 | 0.5749 |  |
| SPA103 | 0.6749 | 0.7234 | 0.6957 | 0.6957 | 0.6797 | 0.643 | 0.6763 | 0.7021 | 0.7161 | 0.6877 | 0.6837 | 0.7151 | 0.641 | 0.6206 | 0.7492 | 0.7656 | 0.6484 | 0.6907 | 0.4185 | 0.4863 |
| SPA104 | 0.658 | 0.643 | 0.607 | 0.607 | 0.3701 | 0.5018 | 0.4637 | 0.4792 | 0.6794 | 0.6621 | 0.6577 | 0.6548 | 0.5802 | 0.6054 | 0.6432 | 0.6276 | 0.6028 | 0.5798 | 0.575 | 0.5872 |
| SPA105 | 0.1173 | 0.1096 | 0.169 | 0.169 | 0.2053 | 0.1398 | 0.2133 | 0.2231 | 0.1664 | 0.2979 | 0.2114 | 0.172 | 0.1829 | 0.0947 | 0.2931 | 0.1814 | 0.2317 | 0.0567 | 0.1162 | 0.2397 |
| SPA106 | 0.5636 | 0.5065 | 0.5249 | 0.5249 | 0.1168 | 0.1572 | 0.5792 | 0.5983 |  | 0.5301 | 0.6101 | 0.5629 | 0.2157 | 0.2397 | 0.6723 | 0.6051 | 0.6071 | 0.5625 | 0.2564 | 0.3343 |
| SPA107 | 0.1639 | 0.165 | 0.1928 | 0.1928 | 0.1702 | 0.1425 | 0.5521 | 0.345 | 0.4186 | 0.3261 | 0.5251 | 0.4679 | 0.2838 | 0.2816 | 0.5041 | 0.303 | 0.2016 | 0.233 | 0.3031 | 0.1344 |
| SPA108 | 0.3864 | 0.3712 | 0.3776 | 0.3776 | 0.3159 | 0.2855 | 0.4214 | 0.4148 | 0.3775 | 0.3756 | 0.3004 | 0.2896 | 0.3124 | 0.2516 | 0.3957 | 0.3601 | 0.3807 | 0.351 | 0.3301 | 0.3441 |
| SPA109 | 0.5597 | 0.5714 | 0.5412 | 0.5412 | 0.5852 | 0.5505 | 0.6261 | 0.6154 | 0.5448 | 0.5756 |  | 0.5544 | 0.5253 | 0.5056 | 0.5655 | 0.5903 | 0.5579 | 0.5156 | 0.4592 | 0.5108 |
| SPA111 | 0.1691 | 0.1568 | 0.1395 | 0.1395 | 0.2788 | 0.1855 | 0.1665 | 0.1723 | 0.1599 | 0.1723 | 0.2079 | 0.1922 | 0.1091 | 0.1137 | 0.1181 | 0.1097 | 0.3076 | 0.2503 | 0.1359 | 0.0824 |
| SPA112 | 0.2954 | 0.2844 | 0.2281 | 0.2281 | 0.3076 | 0.1946 | 0.3683 | 0.3274 | 0.331 | 0.3115 | 0.4647 | 0.4553 | 0.204 | 0.1112 | 0.2791 | 0.3044 | 0.5839 | 0.275 | 0.1921 | 0.1645 |
| SPA114 | 0.3768 | 0.3169 | 0.2132 | 0.2132 | 0.3291 | 0.3558 | 0.3698 | 0.2991 | 0.1986 | 0.1313 | 0.2073 | 0.1804 | 0.2923 | 0.4256 | 0.1491 | 0.1374 | 0.2388 | 0.2628 | 0.2254 | 0.3493 |
| SPA115 | 0.428 | 0.4007 | 0.5423 | 0.5423 | 0.1341 | 0.2834 | 0.3305 | 0.4413 | 0.168 | 0.1534 | 0.2925 | 0.1934 | 0.3235 | 0.3114 | 0.4161 | 0.4129 | 0.5028 | 0.5424 | 0.2581 | 0.1466 |
| SPA116 | 0.6014 | 0.693 | 0.5884 | 0.5884 | 0.5491 | 0.4805 | 0.4391 | 0.4154 | 0.5505 | 0.4945 | 0.5811 | 0.5326 | 0.5443 | 0.5656 | 0.0802 | 0.108 | 0.1101 | 0.1667 | 0.4262 | 0.3959 |
| SPA117 | 0.4439 | 0.529 | 0.3412 | 0.3412 | 0.403 | 0.424 | 0.4095 | 0.3635 | 0.5936 | 0.5169 | 0.3946 | 0.4008 | 0.4255 | 0.4177 | 0.4509 | 0.4156 | 0.1335 | 0.1905 | 0.2587 | 0.3036 |
| SPA118 | 0.3699 | 0.4021 | 0.3216 | 0.3216 | 0.2917 | 0.3227 | 0.3419 | 0.3701 | 0.4048 | 0.4071 | 0.3971 | 0.4272 | 0.2563 | 0.2209 | 0.3702 | 0.3713 | 0.2765 | 0.3285 | 0.2905 | 0.282 |
| SPA55 | 0.2018 | 0.192 | 0.1152 | 0.1152 | 0.0492 | 0.0625 | 0.0811 | 0.1095 | 0.2078 | 0.1687 | 0.1326 | 0.1914 | 0.1712 | 0.1353 | 0.1546 | 0.0962 | 0.2048 | 0.1124 | 0.2521 | 0.2966 |
| SPA57 | 0.4806 | 0.4324 | 0.3826 | 0.3826 | 0.4885 | 0.4503 | 0.4455 | 0.5357 | 0.3669 | 0.4058 | 0.5281 | 0.5148 | 0.3958 | 0.3978 | 0.5245 | 0.484 | 0.4722 | 0.4872 | 0.508 | 0.4882 |
| SPA60 | 0.3472 | 0.3573 | 0.2785 | 0.2785 | 0.4258 | 0.5181 | 0.2232 | 0.1897 | 0.301 | 0.1857 | 0.2659 | 0.2525 | 0.4617 | 0.3687 | 0.2731 | 0.4903 | 0.434 | 0.336 | 0.5575 | 0.4506 |
| SPA61 | 0.4294 | 0.3735 | 0.353 | 0.353 | 0.3664 | 0.3653 | 0.3942 | 0.3701 | 0.4493 | 0.4211 | 0.4046 | 0.2467 | 0.3168 | 0.311 | 0.5753 | 0.4867 | 0.1915 | 0.2738 | 0.4127 | 0.619 |
| SPA64 | 0.2325 | 0.2212 | 0.2504 | 0.2504 | 0.6988 | 0.4615 | 0.4349 | 0.4649 | 0.1793 | 0.2048 | 0.1357 | 0.1538 | 0.5491 | 0.5048 | 0.1823 | 0.2455 | 0.1778 | 0.2276 | 0.6483 | 0.3788 |
| SPA66 | 0.19 | 0.1995 | 0.3846 | 0.3846 | 0.4182 | 0.3741 | 0.3829 | 0.4387 | 0.2565 | 0.2221 | 0.1951 | 0.2523 | 0.424 | 0.4283 | 0.4301 | 0.5542 | 0.2624 | 0.3707 | 0.3073 | 0.3824 |
| SPA67 | 0.2707 | 0.3508 | 0.475 | 0.475 | 0.467 | 0.5281 | 0.5155 | 0.5129 | 0.3178 | 0.3594 | 0.2901 | 0.3514 | 0.4513 | 0.5319 | 0.3389 | 0.3763 | 0.5673 | 0.546 | 0.5555 | 0.5654 |
| SPA68 | 0.2626 | 0.2886 | 0.5599 | 0.5599 | 0.5703 | 0.5803 | 0.5509 | 0.5658 | 0.272 | 0.2958 | 0.2552 | 0.2785 | 0.4109 | 0.5425 | 0.3801 | 0.3115 | 0.5802 | 0.6124 | 0.5569 | 0.5459 |
| SPA69 | 0.1612 | 0.0979 | 0.1517 | 0.1517 | 0.2888 | 0.3183 | 0.1217 | 0.1981 | 0.1747 | 0.1804 | 0.305 | 0.3136 | 0.2188 | 0.2753 | 0.2911 | 0.2836 | 0.3125 | 0.4115 | 0.201 | 0.2037 |
| SPA73 | 0.1509 | 0.1336 | 0.126 | 0.126 | 0.1293 | 0.083 | 0.1172 | 0.1444 | 0.1596 | 0.1486 | 0.1736 | 0.1653 | 0.0698 | 0.0774 | 0.1831 | 0.1693 | 0.1785 | 0.158 | 0.0886 | 0.063 |
| SPA80 | 0.258 | 0.2708 | 0.1835 | 0.1835 | 0.1775 | 0.149 | 0.3529 | 0.3386 | 0.3393 | 0.3405 | 0.2876 | 0.2818 | 0.1719 | 0.1164 | 0.3033 | 0.2233 | 0.2293 | 0.3203 | 0.1403 | 0.1128 |
| SPA82 | 0.2612 | 0.279 | 0.2584 | 0.2584 | 0.3205 | 0.2698 | 0.1835 | 0.1626 | 0.2361 | 0.2726 | 0.2681 | 0.2569 | 0.2245 | 0.1108 | 0.1873 | 0.2813 | 0.2269 | 0.3692 | 0.1581 | 0.1902 |
| SPA83 | 0.1768 | 0.1601 | 0.076 | 0.076 | 0.1629 | 0.1252 | 0.0725 | 0.1077 | 0.1846 | 0.1976 | 0.1261 | 0.1172 | 0.2643 | 0.2013 | 0.142 | 0.1539 | 0.2935 | 0.3326 | 0.315 | 0.2916 |
| SPA84 | 0.2564 | 0.2492 | 0.3646 | 0.3646 | 0.3629 | 0.2263 | 0.2927 | 0.2928 | 0.3499 | 0.3373 | 0.2666 | 0.3043 | 0.1993 | 0.322 | 0.341 | 0.3835 | 0.2149 | 0.3101 | 0.2401 | 0.2533 |
| SPA90 | 0.4342 | 0.3003 | 0.2142 | 0.2142 | 0.1471 | 0.2084 | 0.1445 | 0.1298 | 0.1366 | 0.1361 | 0.1496 | 0.0827 | 0.2388 | 0.2203 | 0.1578 | 0.0561 | 0.1284 | 0.1521 | 0.1324 | 0.1797 |
| SPA91 | 0.0685 | 0.0739 | 0.0936 | 0.0936 | 0.1496 | 0.2556 | 0.1164 | 0.1001 | 0.2688 | 0.1323 | 0.1607 | 0.2726 | 0.1546 | 0.1498 | 0.2488 | 0.2116 | 0.1449 | 0.1643 | 0.2796 | 0.296 |
| SPA92 | 0.299 | 0.4497 | 0.2376 | 0.2376 | 0.3703 | 0.1955 | 0.2493 | 0.2123 | 0.256 | 0.1372 | 0.333 | 0.1995 | 0.151 | 0.1439 | 0.0905 | 0.1197 | 0.224 | 0.1144 | 0.2814 | 0.2687 |
| SPA93 | 0.2469 | 0.1725 | 0.3014 | 0.3014 | 0.3471 | 0.36 | 0.438 | 0.4913 | 0.2505 | 0.2208 | 0.2089 | 0.1629 | 0.4021 | 0.3821 | 0.2388 | 0.3103 |  | 0.4637 | 0.2293 | 0.2169 |
| SPA94 | 0.2122 | 0.2379 | 0.4391 | 0.4391 | 0.2874 | 0.2427 | 0.4961 | 0.5054 | 0.22 | 0.2187 | 0.2713 | 0.2437 | 0.454 | 0.4399 | 0.3084 | 0.4043 | 0.3515 | 0.376 | 0.3341 | 0.3515 |
| SPA95 | 0.2256 | 0.2651 | 0.2684 | 0.2684 | 0.1856 | 0.2557 | 0.3449 | 0.2894 | 0.1848 | 0.1878 | 0.2098 | 0.2004 | 0.2892 | 0.3415 | 0.2304 | 0.34 |  | 0.199 | 0.449 | 0.3658 |
| SPA96 | 0.4457 | 0.4021 | 0.3368 | 0.3368 | 0.5556 | 0.6326 | 0.499 | 0.4538 | 0.3301 | 0.441 | 0.4589 | 0.4562 | 0.3198 | 0.2079 | 0.4538 | 0.4168 | 0.1548 | 0.2557 | 0.3246 | 0.3818 |
| SPA97 | 0.4526 | 0.4799 | 0.5291 | 0.5291 | 0.6154 | 0.6553 | 0.6369 | 0.5944 | 0.3113 | 0.4473 | 0.409 | 0.5314 | 0.6359 | 0.6392 | 0.3967 | 0.3597 | 0.3875 | 0.3686 | 0.5046 | 0.3023 |
| SPA98 | 0.3422 | 0.3529 | 0.3429 | 0.3429 | 0.3105 | 0.2994 | 0.3862 | 0.3996 | 0.3467 | 0.2786 | 0.4404 | 0.3435 | 0.3277 | 0.28 | 0.2475 | 0.2848 | 0.2804 | 0.279 | 0.6435 | 0.3832 |
| SPA99 | 0.2191 | 0.2549 | 0.3089 | 0.3089 | 0.3104 | 0.3482 | 0.2875 | 0.2862 | 0.2345 | 0.2356 | 0.3211 | 0.3618 | 0.6767 | 0.6253 | 0.2575 | 0.2578 | 0.2959 | 0.3495 | 0.4663 | 0.4271 |
| TENX128 | 0.37 |  |  |  | 0.5714 |  | 0.3316 |  |  |  | 0.3648 |  | 0.2061 |  |  |  | 0.3336 |  | 0.2505 |  |
| TENX147 | 0.3876 | 0.4218 | 0.3199 | 0.3199 | 0.2265 | 0.344 | 0.342 | 0.2345 | 0.4264 | 0.4878 | 0.4147 | 0.3841 | 0.2206 | 0.2425 | 0.4435 | 0.4502 | 0.5383 | 0.5713 | 0.3545 | 0.4152 |
| TENX148 | 0.6974 | 0.6594 | 0.6258 | 0.6258 | 0.8995 | 0.7659 | 0.6324 | 0.3405 | 0.7259 | 0.382 | 0.7897 | 0.5208 | 0.3854 | 0.3007 |  | 0.6136 | 0.8963 | 0.7473 | 0.2817 | 0.3426 |
| TENX149 | 0.777 | 0.6645 | 0.6068 | 0.6068 | 0.6532 | 0.5149 | 0.6691 | 0.5231 | 0.7105 | 0.5821 |  | 0.6205 | 0.2403 | 0.2893 | 0.741 | 0.6797 | 0.6908 | 0.5357 | 0.3312 | 0.3318 |
| TENX152 | 0.384 | 0.2917 | 0.1905 | 0.1905 | 0.1285 | 0.1297 | 0.4639 | 0.4522 | 0.37 | 0.305 | 0.366 | 0.2695 | 0.4699 | 0.3289 |  | 0.4789 | 0.187 | 0.251 | 0.2662 | 0.2786 |
| TENX157 | 0.1948 | 0.2372 | 0.3306 | 0.3306 | 0.1449 | 0.3547 | 0.1789 | 0.2984 | 0.1139 | 0.2313 | 0.2111 | 0.2479 | 0.3061 | 0.4395 | 0.0652 | 0.3713 | 0.159 | 0.2626 | 0.6805 | 0.7636 |
| TENX30 | 0.693 | 0.462 | 0.4863 | 0.4863 | 0.2619 | 0.4551 | 0.5293 | 0.3471 | 0.6627 | 0.457 | 0.6777 | 0.3933 | 0.4399 | 0.3576 | 0.6754 | 0.2154 | 0.6186 | 0.4287 | 0.5058 | 0.6356 |
| TENX31 | 0.4435 | 0.3355 | 0.3854 | 0.3854 | 0.4019 | 0.3744 | 0.3994 | 0.4708 | 0.3505 | 0.3234 | 0.4235 | 0.3741 | 0.2878 | 0.3909 | 0.4176 | 0.4589 | 0.3759 | 0.5034 | 0.2676 | 0.4775 |
| TENX46 | 0.2297 | 0.5563 | 0.3353 | 0.2187 | 0.14 | 0.133 | 0.3119 | 0.4102 |  |  | 0.2702 |  | 0.4347 | 0.1857 | 0.3494 | 0.4168 | 0.2959 |  | 0.3133 | 0.2327 |
| TENX70 | 0.3683 |  | 0.2473 | 0.2481 | 0.2472 | 0.261 | 0.3373 | 0.2725 | 0.3367 | 0.303 | 0.3791 | 0.3308 | 0.4386 | 0.3707 |  | 0.3915 | 0.2538 | 0.3313 | 0.2855 | 0.4472 |
| TENX90 | 0.0996 | 0.2646 | 0.1306 | 0.1306 | 0.1369 | 0.1031 | 0.1582 | 0.1629 | 0.1004 | 0.1452 | 0.0657 | 0.2599 | 0.1758 | 0.2381 | 0.3027 | 0.285 | 0.4294 | 0.2403 | 0.0706 | 0.0994 |
| TENX91 | 0.2563 | 0.1878 | 0.2111 | 0.2111 | 0.14 | 0.2052 | 0.1652 | 0.1286 | 0.3415 | 0.2941 | 0.3274 | 0.1531 | 0.2531 | 0.3653 | 0.2988 | 0.1357 | 0.4546 | 0.3769 | 0.2092 | 0.178 |
| TENX92 | 0.4856 | 0.3516 | 0.4118 | 0.4118 | 0.2933 | 0.3888 | 0.4002 | 0.3661 | 0.5197 | 0.1631 | 0.4352 | 0.3907 | 0.4004 | 0.4435 | 0.4956 | 0.4873 | 0.547 | 0.5261 | 0.1304 | 0.1956 |
| ZEN38 | 0.3188 | 0.1664 | 0.1843 | 0.1843 | 0.2481 | 0.22 | 0.1522 | 0.1541 | 0.3023 | 0.1874 | 0.2839 | 0.1656 | 0.2729 | 0.319 | 0.2489 | 0.1973 | 0.3627 | 0.3131 | 0.2298 | 0.3428 |
| ZEN42 | 0.2017 | 0.1586 | 0.0957 | 0.0957 | 0.1042 | 0.1577 | 0.16 | 0.1037 | 0.1931 | 0.1496 | 0.2189 | 0.1773 | 0.1226 | 0.1813 | 0.3004 | 0.1169 | 0.3108 | 0.1452 | 0.0542 | 0.1632 |
| ZEN44 | 0.1981 | 0.1977 | 0.2891 | 0.2891 | 0.2102 | 0.3318 | 0.1611 | 0.2231 | 0.2295 | 0.2423 | 0.2736 | 0.2121 | 0.4159 | 0.3644 | 0.1534 | 0.2475 | 0.199 | 0.3061 | 0.2962 | 0.3722 |
| ZEN45 | 0.3 | 0.1588 | 0.1515 | 0.1515 | 0.1461 | 0.253 | 0.151 | 0.2381 | 0.1644 | 0.2055 | 0.2581 | 0.2068 | 0.1253 | 0.1334 | 0.1494 | 0.1253 | 0.16 | 0.1357 | 0.1572 | 0.1177 |
| ZEN78 | 0.1166 | 0.278 | 0.1427 | 0.1427 | 0.1239 | 0.1133 | 0.1139 | 0.1402 | 0.1107 | 0.1518 | 0.0793 | 0.2006 | 0.1294 | 0.1185 | 0.096 | 0.3031 | 0.086 | 0.1395 | 0.2911 | 0.4279 |
| ZEN80 | 0.1914 | 0.1888 | 0.1728 | 0.1729 | 0.1056 | 0.1709 | 0.2091 | 0.1797 | 0.2077 | 0.2616 | 0.1438 | 0.2436 | 0.2342 | 0.285 | 0.2202 | 0.1702 | 0.1594 | 0.2025 | 0.2147 | 0.2325 |
| ZEN81 | 0.1877 | 0.1819 | 0.2047 | 0.2047 | 0.0577 | 0.1833 | 0.1851 | 0.1527 | 0.1961 | 0.1346 | 0.1856 | 0.1273 | 0.2026 | 0.126 | 0.0604 | 0.2192 |  | 0.0631 | 0.1206 | 0.2437 |
| INT12 |  | 0.3175 | 0.5779 | 0.5779 | 0.6463 | 0.5949 | 0.3412 | 0.3277 | 0.5077 | 0.5034 | 0.1262 | 0.6118 | 0.4069 | 0.5113 | 0.2388 | 0.5459 |  | 0.5531 | 0.5891 | 0.646 |
| MEND148 |  | 0.3596 | 0.3936 | 0.3937 | 0.4016 | 0.3392 | 0.3256 | 0.3507 | 0.4211 | 0.326 | 0.3559 | 0.29 | 0.3965 | 0.3027 | 0.4353 | 0.3391 | 0.4753 | 0.2614 | 0.4974 | 0.3992 |
| MEND65 |  | 0.1891 | 0.1831 | 0.1831 | 0.169 | 0.169 | 0.1585 | 0.1689 |  | 0.2036 | 0.1775 | 0.1589 | 0.1462 | 0.1859 | 0.1438 | 0.1341 | 0.1278 | 0.1503 | 0.1753 | 0.1379 |
| MISC42 |  | 0.5068 | 0.3425 | 0.3425 | 0.4171 | 0.4062 | 0.5776 | 0.5429 |  | 0.5048 |  | 0.3881 | 0.505 | 0.5849 | 0.4547 | 0.5065 | 0.4971 | 0.6202 | 0.5646 | 0.4376 |
| MISC57 |  | 0.3068 | 0.3869 | 0.3869 | 0.4964 | 0.3799 | 0.3468 | 0.1786 | 0.5287 | 0.4264 |  | 0.4904 | 0.4507 | 0.3372 |  | 0.3598 |  | 0.2036 | 0.1894 | 0.3386 |
| MISC58 |  | 0.3577 | 0.5924 | 0.5924 | 0.5045 | 0.6156 | 0.3036 | 0.6345 | 0.3596 | 0.6026 | 0.6804 | 0.2813 | 0.7598 | 0.6781 | 0.3312 | 0.6169 |  | 0.4392 | 0.6912 | 0.7468 |
| TENX40 |  | 0.5042 | 0.4302 | 0.48 | 0.4568 | 0.4627 | 0.546 | 0.5252 | 0.4466 | 0.4326 | 0.4727 | 0.4695 | 0.3937 | 0.4069 | 0.3491 | 0.4262 |  | 0.4609 | 0.3348 | 0.2478 |
| ZEN79 |  | 0.3021 | 0.2469 | 0.2469 | 0.2498 | 0.1794 | 0.2843 | 0.2489 | 0.3286 | 0.3012 | 0.2615 | 0.225 |  | 0.2118 |  | 0.1887 | 0.3195 | 0.3113 | 0.3057 | 0.288 |
| MEND156 |  |  | 0.3817 | 0.3817 | 0.4058 | 0.3915 | 0.3009 | 0.224 | 0.2259 |  | 0.1646 | 0.4715 | 0.1925 | 0.2523 | 0.1448 | 0.41 |  | 0.3987 | 0.2708 | 0.2927 |
| MEND141 |  |  |  |  |  | 0.0924 |  |  |  |  |  |  | 0.3489 | 0.2974 |  |  |  | 0.1832 |  | 0.4335 |

### Table.S5

| cosine_similarity | UNI_big_region | UNI_small_region | CONCH_big_region | CONCH_small_region | CONCH15_big_region | CONCH15_small_region | CTransPath_big_region | CTransPath_small_region | GigaPath_big_region | GigaPath_small_region | UNI2h_big_region | UNI2h_small_region | RN18_big_region | RN18_small_region | Virchow_big_region | Virchow_small_region | Virchow2_big_region | Virchow2_small_region | ViTB16_big_region | ViTB16_small_region |
|---|---|---|---|---|---|---|---|---|---|---|---|---|---|---|---|---|---|---|---|---|
| INT13 | 0.4905 | 0.351 | 0.4385 | 0.4961 | 0.0748 | 0.0249 | 0.2625 | 0.431 | 0.4898 | 0.4599 | 0.5968 | 0.4447 | 0.216 | 0.2589 | 0.4617 | 0.3435 | 0.3771 | 0.527 | 0.302 | 0.3778 |
| INT14 | 0.8364 | 0.8672 | 0.7575 | 0.7575 | 0.8028 | 0.7311 | 0.8618 | 0.8176 | 0.8672 | 0.8448 | 0.8565 | 0.8372 | 0.7836 | 0.7275 | 0.8875 | 0.8827 | | 0.806 | 0.8206 | 0.7397 |
| INT15 | 0.8332 | 0.8381 | 0.6913 | 0.6913 | 0.7346 | 0.7083 | 0.8225 | 0.7 | 0.855 | 0.8451 | 0.8347 | 0.8311 | 0.5898 | 0.3013 | 0.8401 | 0.834 | 0.8187 | 0.8019 | 0.6305 | 0.5417 |
| INT16 | 0.7746 | 0.699 | 0.625 | 0.625 | 0.6576 | 0.5737 | 0.6826 | 0.3355 | 0.7558 | 0.6444 | 0.7817 | 0.6138 | 0.2577 | 0.248 | 0.8063 | 0.7489 | 0.7409 | 0.6131 | 0.2996 | 0.3752 |
| INT19 | 0.3613 | 0.3591 | 0.312 | 0.379 | 0.3625 | 0.2055 | 0.3189 | 0.3194 | 0.3487 | 0.3436 | 0.4102 | 0.3884 | 0.349 | 0.3474 | 0.373 | 0.3872 | 0.3719 | 0.3776 | 0.3463 | 0.3382 |
| INT21 | 0.8539 | 0.6319 | 0.6008 | 0.6008 | 0.813 | 0.6423 | 0.7978 | 0.2247 | 0.8289 | 0.7259 | 0.8405 | 0.7666 | 0.6003 | -0.029 | 0.8219 | 0.7411 | 0.8419 | 0.6105 | 0.6277 | 0.5092 |
| INT22 | 0.2314 | 0.3386 | 0.3356 | 0.3356 | 0.0181 | 0.228 | 0.0541 | 0.1921 | 0.1745 | 0.1332 | 0.3807 | 0.2323 | 0.3038 | 0.2017 | 0.2925 | 0.337 | 0.3103 | 0.023 | 0.0602 | 0.152 |
| INT24 | 0.8311 | 0.4413 | 0.0626 | 0.0625 | 0.6506 | 0.2459 | 0.463 | -0.3187 | 0.7763 | 0.4758 | 0.8314 | 0.7935 | -0.1728 | -0.2066 | 0.7208 | 0.6141 | 0.8399 | 0.4213 | 0.1307 | -0.1079 |
| INT25 | 0.5191 | 0.3761 | 0.3839 | 0.322 | 0.3755 | 0.2768 | 0.4209 | 0.3758 | 0.5734 | 0.4111 | 0.6251 | 0.4343 | 0.4251 | 0.3111 | 0.5668 | 0.2939 | 0.5474 | 0.3967 | 0.3946 | 0.4074 |
| INT26 | 0.4776 | 0.4804 | 0.4865 | 0.4869 | 0.5358 | 0.4279 | 0.6032 | 0.6127 | 0.6054 | 0.5448 | 0.4434 | 0.4731 | 0.5944 | 0.515 | 0.6326 | 0.326 | 0.5614 | 0.5082 | 0.5483 | 0.3213 |
| INT27 | 0.5501 | 0.7302 | 0.7374 | 0.7374 | 0.8026 | 0.7908 | 0.7433 | 0.7235 | 0.8131 | 0.8376 | 0.5208 | 0.7606 | 0.5952 | 0.5163 | 0.9091 | 0.6566 | 0.8333 | | 0.7153 | 0.7769 |
| INT28 | 0.7627 | 0.7617 | 0.7044 | 0.7044 | 0.7783 | 0.787 | 0.8536 | 0.7964 | 0.8562 | 0.8093 | 0.7747 | 0.6825 | 0.4916 | 0.5009 | 0.8598 | 0.7313 | 0.8036 | 0.7248 | 0.7236 | 0.6934 |
| INT35 | 0.47 | 0.6115 | 0.6047 | 0.6047 | 0.594 | 0.6496 | 0.4907 | 0.5139 | 0.5195 | 0.6982 | | 0.6486 | 0.4114 | 0.4784 | 0.5989 | 0.413 | 0.5988 | 0.641 | 0.4284 | 0.3892 |
| INT4 | 0.1723 | -0.2142 | -0.1356 | -0.1356 | -0.3005 | -0.3216 | -0.1145 | -0.0099 | 0.1473 | 0.0008 | -0.0259 | -0.2641 | -0.2797 | -0.2786 | -0.0772 | 0.0181 | -0.3052 | -0.1267 | -0.0377 | -0.1733 |
| INT5 | 0.0935 | -0.2235 | -0.1663 | -0.1663 | -0.0452 | -0.3149 | -0.1134 | -0.1144 | 0.0996 | 0.1013 | 0.283 | -0.1941 | 0.1744 | -0.2211 | 0.0796 | -0.0614 | 0.3156 | 0.0617 | 0.0646 | -0.0338 |
| INT7 | 0.214 | 0.1807 | 0.0269 | 0.0269 | 0.081 | -0.0473 | 0.1793 | 0.2061 | 0.1351 | 0.1618 | 0.1939 | 0.1423 | -0.0187 | -0.0835 | 0.1568 | 0.183 | 0.2346 | 0.2511 | 0.0833 | -0.0155 |
| INT8 | 0.0547 | -0.0209 | 0.0714 | 0.0714 | 0.0445 | 0.0735 | 0.1003 | 0.0148 | 0.067 | 0.0522 | 0.1202 | 0.0387 | 0.1018 | 0.0232 | 0.119 | 0.0792 | 0.0942 | 0.0144 | 0.0605 | 0.0336 |
| MEND140 | 0.4502 | 0.6763 | 0.5405 | 0.5405 | 0.3616 | 0.2261 | 0.2749 | 0.5448 | 0.5824 | 0.634 | 0.4512 | 0.6906 | 0.4156 | 0.5373 | 0.5727 | 0.6864 | 0.6275 | 0.6899 | 0.2601 | 0.0089 |
| MEND142 | 0.7103 | 0.7527 | 0.8106 | 0.8106 | 0.7773 | 0.7587 | 0.4151 | 0.6692 | 0.7066 | 0.788 | 0.7464 | 0.8165 | 0.1371 | 0.326 | 0.6467 | 0.6877 | 0.6367 | 0.7592 | 0.6259 | 0.4186 |
| MEND143 | 0.8232 | 0.8207 | 0.7017 | 0.7017 | 0.7403 | 0.5158 | 0.6765 | 0.8265 | 0.827 | 0.7669 | 0.7385 | 0.7986 | 0.1106 | 0.124 | 0.8067 | 0.7306 | 0.8089 | 0.737 | 0.216 | 0.0381 |
| MEND145 | 0.7568 | 0.6558 | 0.6842 | 0.6842 | 0.7676 | 0.4582 | 0.5078 | 0.6597 | 0.8105 | 0.6039 | 0.7592 | 0.6211 | 0.3868 | 0.2806 | 0.7641 | 0.5681 | 0.763 | 0.6172 | 0.2498 | 0.1895 |
| MEND146 | 0.5008 | 0.4758 | 0.2822 | 0.2822 | 0.5187 | 0.1866 | 0.2487 | 0.3988 | 0.5952 | 0.5143 | 0.567 | 0.5785 | 0.2506 | 0.1826 | 0.4881 | 0.3856 | 0.4544 | 0.499 | 0.266 | 0.2907 |
| MEND147 | 0.5438 | 0.4645 | 0.4311 | 0.4311 | 0.5377 | 0.4321 | 0.4932 | 0.4674 | 0.6785 | 0.5729 | 0.5306 | 0.391 | 0.2539 | 0.285 | 0.534 | 0.3974 | 0.5487 | 0.432 | 0.3164 | 0.179 |
| MEND150 | 0.6985 | 0.8634 | 0.8557 | 0.8557 | 0.5309 | 0.8463 | 0.6449 | 0.8301 | 0.8549 | 0.8727 | 0.6764 | 0.8607 | 0.5897 | 0.6266 | 0.8237 | 0.7905 | 0.7813 | 0.8663 | 0.2394 | 0.7173 |
| MEND151 | 0.4458 | 0.675 | 0.7187 | 0.6998 | 0.5609 | 0.5417 | 0.5123 | | 0.5878 | 0.6539 | 0.4875 | 0.6911 | 0.5311 | 0.5522 | 0.4152 | 0.5582 | 0.585 | 0.7058 | 0.2405 | 0.3622 |
| MEND153 | 0.2038 | 0.0041 | -0.039 | -0.039 | 0.0298 | -0.3557 | 0.5414 | 0.0516 | 0.3626 | 0.002 | 0.3166 | -0.0133 | 0.264 | 0.131 | 0.1258 | -0.1193 | -0.0166 | -0.1849 | 0.2176 | 0.1137 |
| MEND154 | -0.0612 | 0.3486 | 0.7218 | 0.4888 | -0.1209 | 0.3608 | 0.0719 | 0.4778 | 0.1668 | 0.5109 | 0.0862 | 0.2299 | -0.1591 | 0.3477 | 0.25 | 0.5003 | 0.0053 | 0.5071 | -0.0413 | 0.3414 |
| MEND157 | 0.6906 | 0.7341 | 0.6241 | 0.7479 | 0.6804 | 0.697 | 0.6241 | 0.6377 | 0.7261 | 0.7477 | 0.6872 | 0.7439 | 0.6475 | 0.6068 | 0.733 | 0.7117 | 0.6913 | 0.6144 | 0.4984 | 0.6744 |
| MEND158 | 0.4973 | 0.6981 | -0.1932 | 0.6468 | 0.4359 | 0.3707 | 0.5325 | 0.6647 | 0.7003 | 0.6957 | 0.3975 | 0.6795 | 0.3085 | 0.4014 | 0.3682 | 0.518 | 0.6989 | 0.5593 | 0.0466 | 0.3979 |
| MEND160 | 0.734 | 0.8212 | 0.7804 | 0.7718 | 0.7555 | 0.6614 | 0.7472 | 0.8012 | 0.778 | 0.8085 | 0.7146 | 0.7684 | 0.7169 | 0.6213 | 0.7971 | 0.7706 | 0.7142 | 0.726 | 0.6781 | 0.6534 |
| MEND162 | 0.6268 | 0.3568 | 0.2422 | 0.2422 | 0.4716 | 0.2215 | 0.3857 | 0.2825 | 0.439 | 0.1868 | 0.6588 | 0.5189 | 0.0974 | 0.2985 | 0.5053 | 0.2185 | 0.3566 | 0.2783 | 0.1369 | 0.1559 |
| MEND18 | -0.0061 | 0.0996 | 0.151 | 0.151 | -0.1361 | -0.0917 | -0.0125 | 0.011 | 0.1576 | 0.1943 | 0.2264 | 0.2463 | 0.013 | 0.1471 | 0.0516 | 0.1232 | 0.1948 | 0.2433 | -0.0499 | 0.0396 |
| MEND19 | 0.7453 | 0.732 | 0.6828 | 0.6828 | 0.6864 | 0.6713 | 0.6884 | 0.7116 | 0.7507 | 0.7366 | 0.6941 | 0.6818 | 0.374 | 0.4608 | 0.6422 | 0.6 | 0.6508 | 0.6881 | 0.4116 | 0.459 |
| MEND24 | -0.2302 | -0.2012 | 0.1306 | 0.2086 | -0.3479 | -0.2443 | -0.1453 | -0.1823 | -0.1578 | -0.0885 | -0.343 | -0.2838 | -0.1149 | 0.1283 | 0.0118 | 0.0726 | 0.1919 | 0.2655 | -0.1891 | -0.1671 |
| MEND37 | 0.6194 | 0.6575 | 0.595 | 0.595 | 0.5717 | 0.5709 | 0.5114 | 0.572 | 0.5665 | 0.7239 | 0.52 | 0.7531 | 0.2967 | 0.3753 | 0.4912 | 0.4512 | 0.6024 | 0.6305 | 0.4701 | 0.3902 |
| MEND38 | 0.569 | 0.5947 | 0.4805 | 0.4805 | 0.4885 | 0.4019 | 0.5082 | 0.5485 | 0.4194 | 0.4828 | 0.4079 | 0.4969 | 0.1022 | 0.2703 | 0.3464 | 0.3756 | 0.5142 | 0.5946 | 0.2944 | 0.1859 |
| MEND39 | 0.7244 | 0.6918 | 0.6092 | 0.653 | 0.7032 | 0.6633 | 0.6014 | 0.6404 | 0.6198 | 0.6965 | 0.6632 | 0.691 | 0.402 | 0.3222 | 0.5955 | 0.5362 | 0.6867 | 0.754 | 0.5099 | 0.3176 |
| MEND40 | 0.6236 | 0.6458 | 0.5509 | 0.5508 | 0.6208 | 0.5739 | 0.5462 | 0.6073 | 0.5844 | 0.5997 | 0.5408 | 0.6913 | 0.3224 | 0.407 | 0.5953 | 0.394 | 0.6349 | 0.6475 | 0.5866 | 0.4653 |
| MEND59 | 0.8945 | 0.4785 | 0.1604 | 0.1604 | 0.672 | 0.2161 | 0.8455 | 0.3679 | 0.8545 | 0.4507 | 0.91 | 0.34 | 0.7016 | 0.3916 | | 0.4563 | 0.8394 | 0.1552 | 0.6612 | 0.3666 |
| MEND60 | 0.8991 | 0.796 | 0.6509 | 0.6509 | 0.8251 | 0.6801 | 0.8967 | 0.822 | 0.8825 | 0.7782 | 0.9085 | 0.7553 | 0.7557 | 0.7318 | 0.8856 | 0.6484 | 0.8839 | 0.7013 | 0.7371 | 0.3323 |
| MEND61 | 0.9632 | 0.8163 | 0.6022 | 0.6022 | 0.8126 | 0.6324 | 0.9472 | 0.7357 | 0.9592 | 0.6268 | 0.9557 | 0.8014 | 0.814 | 0.6238 | 0.9501 | 0.7772 | 0.8845 | 0.5121 | 0.7853 | 0.511 |
| MEND62 | 0.937 | 0.8775 | 0.7008 | 0.7008 | 0.8452 | 0.7486 | 0.9349 | 0.8407 | 0.9389 | 0.8209 | 0.9083 | 0.7869 | 0.8495 | 0.7299 | 0.9276 | 0.8125 | 0.9334 | 0.8178 | 0.7685 | 0.6992 |
| MEND66 | 0.0174 | 0.0646 | -0.0144 | -0.0144 | 0.0027 | 0.0357 | 0.0061 | 0.1196 | 0.0544 | 0.075 | -0.0157 | 0.0503 | 0.0597 | 0.1067 | 0.1963 | 0.2165 | 0.1066 | 0.1047 | 0.0987 | 0.0837 |
| MISC33 | 0.8118 | | 0.6218 | 0.6527 | 0.4815 | 0.0211 | 0.6638 | 0.4251 | 0.7715 | 0.6298 | 0.8046 | 0.7623 | 0.0765 | -0.4443 | 0.7912 | 0.6399 | 0.8175 | 0.7458 | 0.4994 | -0.3167 |
| MISC35 | 0.8407 | 0.7886 | 0.6194 | 0.6194 | 0.8079 | 0.6025 | 0.8011 | 0.4175 | 0.8402 | 0.7704 | 0.8383 | 0.7964 | 0.1465 | 0.1291 | 0.8434 | 0.6261 | | 0.676 | 0.598 | -0.0084 |
| MISC36 | 0.8608 | 0.7868 | 0.6311 | 0.6311 | 0.8665 | 0.6744 | 0.7249 | 0.6135 | 0.8632 | 0.7518 | 0.8679 | 0.7771 | 0.3579 | 0.334 | 0.8253 | 0.7655 | 0.6201 | 0.2531 | 0.4607 | 0.2815 |
| MISC40 | 0.5126 | 0.1357 | 0.5397 | 0.5397 | 0.5618 | 0.5668 | 0.2555 | 0.3177 | 0.5169 | 0.4806 | 0.4745 | 0.407 | 0.4171 | 0.073 | 0.5279 | 0.3237 | 0.4873 | 0.3685 | 0.0101 | 0.2777 |
| MISC41 | 0.7068 | 0.4552 | 0.5303 | 0.5303 | 0.6204 | 0.5278 | 0.6084 | 0.0856 | 0.7513 | 0.1756 | 0.7362 | 0.6622 | 0.1322 | 0.0736 | 0.77 | 0.1768 | | 0.6577 | 0.2994 | 0.239 |
| MISC45 | 0.4585 | 0.3006 | 0.2458 | 0.2458 | 0.3869 | 0.2568 | 0.3179 | 0.2304 | 0.5278 | 0.3205 | 0.5217 | 0.4771 | -0.0944 | -0.0736 | 0.5736 | 0.3788 | 0.533 | 0.3934 | 0.1069 | 0.0484 |
| MISC47 | 0.5058 | 0.1549 | 0.0421 | 0.0421 | 0.2932 | -0.0334 | 0.4382 | 0.0502 | 0.5875 | 0.1503 | 0.51 | 0.3665 | 0.123 | 0.0718 | 0.54 | 0.2393 | 0.3827 | 0.0832 | 0.1422 | 0.0186 |
| MISC48 | 0.6799 | 0.6761 | 0.5564 | 0.5564 | 0.6052 | 0.4997 | 0.6772 | 0.669 | 0.6926 | 0.6014 | 0.6426 | 0.6368 | 0.5214 | 0.4076 | 0.6 | 0.5491 | 0.6489 | 0.5637 | 0.5291 | 0.4731 |
| MISC49 | 0.6858 | 0.6425 | 0.5361 | 0.5361 | 0.5947 | 0.4962 | 0.6553 | 0.5431 | 0.7307 | 0.5143 | 0.7068 | 0.6744 | 0.0757 | -0.2555 | 0.6684 | 0.5801 | 0.7172 | 0.6693 | 0.3985 | 0.3953 |
| MISC50 | 0.6779 | 0.6273 | 0.4786 | 0.4786 | 0.6201 | 0.5429 | 0.6364 | 0.418 | 0.7028 | 0.5588 | 0.7055 | 0.6856 | 0.3499 | 0.1485 | 0.656 | 0.3361 | 0.6476 | 0.5131 | 0.287 | 0.226 |
| MISC62 | 0.5853 | 0.1097 | 0.0013 | 0.4082 | 0.5104 | 0.377 | 0.4975 | 0.3067 | 0.6476 | 0.3068 | 0.6614 | 0.457 | 0.3335 | 0.2441 | 0.637 | 0.2524 | 0.6455 | 0.3378 | 0.2174 | 0.2684 |
| MISC63 | -0.0631 | -0.0735 | -0.0154 | -0.0153 | -0.0354 | -0.0502 | -0.0518 | -0.069 | 0.0303 | -0.0099 | -0.04 | -0.0234 | -0.0056 | -0.0104 | 0.0549 | 0.0255 | 0.0116 | -0.0025 | -0.0907 | -0.0778 |
| MISC66 | 0.8362 | 0.2812 | 0.157 | 0.1536 | 0.5509 | -0.0359 | 0.2675 | 0.0184 | 0.7342 | 0.0965 | 0.9179 | 0.582 | -0.2701 | -0.1869 | 0.6053 | 0.0401 | 0.8713 | 0.3123 | -0.1398 | -0.0609 |
| MISC67 | 0.8979 | 0.5294 | 0.352 | 0.352 | 0.603 | 0.334 | 0.4484 | 0.3353 | 0.8123 | 0.4227 | 0.8923 | 0.7539 | -0.094 | 0.0491 | 0.651 | 0.5302 | 0.9085 | 0.6141 | 0.201 | -0.0116 |
| MISC72 | 0.6146 | 0.5457 | 0.5162 | 0.5162 | 0.6241 | 0.5325 | 0.6336 | 0.4873 | 0.6288 | 0.5505 | 0.5798 | 0.59 | 0.3354 | 0.1734 | 0.6046 | 0.4798 | 0.5995 | 0.5652 | 0.3301 | 0.2578 |
| MISC73 | 0.7901 | 0.6787 | 0.6833 | 0.6833 | 0.6804 | 0.6166 | 0.7413 | 0.4346 | 0.6948 | 0.5791 | 0.7848 | 0.7602 | 0.3835 | 0.283 | 0.7606 | 0.5903 | 0.7929 | 0.7353 | 0.0737 | 0.1406 |
| NCBI629 | 0.1944 | 0.1758 | 0.1346 | 0.1346 | 0.0683 | 0.1793 | 0.036 | 0.1796 | 0.1445 | 0.1688 | 0.1614 | 0.1355 | 0.0905 | 0.1534 | 0.1973 | 0.1718 | 0.2032 | 0.1861 | 0.2054 | 0.1362 |
| NCBI631 | 0.0919 | -0.0519 | 0.1848 | 0.1518 | 0.0348 | 0.0072 | 0.0243 | -0.0356 | -0.0326 | -0.0563 | -0.009 | 0.0072 | 0.1153 | -0.1243 | 0.0808 | 0.0208 | -0.0391 | -0.0767 | -0.0996 | 0.0152 |
| NCBI632 | 0.385 | 0.3376 | 0.3934 | 0.3934 | 0.4024 | 0.3993 | 0.3629 | 0.4146 | 0.4323 | 0.3867 | 0.4289 | 0.2536 | 0.2511 | 0.2809 | 0.4264 | 0.3108 | 0.4108 | 0.4037 | 0.389 | 0.4076 |
| NCBI635 | 0.3176 | 0.3175 | 0.2589 | 0.2589 | 0.3455 | 0.285 | 0.3303 | 0.3425 | 0.312 | 0.3308 | 0.2964 | 0.3106 | 0.3137 | 0.3937 | 0.3315 | 0.3126 | 0.3217 | 0.3983 | 0.23 | 0.3259 |
| NCBI638 | -0.0882 | -0.0375 | -0.1164 | -0.1164 | 0.0081 | -0.0682 | -0.0702 | -0.1281 | -0.0619 | -0.0397 | -0.158 | -0.0066 | 0.0815 | 0.0186 | -0.0012 | 0.0735 | -0.1425 | -0.0413 | -0.0448 | -0.0232 |
| NCBI640 | 0.2398 | 0.1497 | 0.1353 | 0.1353 | 0.1573 | 0.2001 | 0.0257 | 0.08 | 0.2171 | 0.2009 | 0.348 | 0.3004 | 0.0898 | 0.0615 | 0.2694 | 0.1536 | 0.2917 | 0.1487 | 0.2555 | 0.2502 |
| NCBI681 | 0.79 | 0.7509 | 0.6673 | 0.6673 | 0.6899 | | 0.8028 | 0.5559 | 0.7816 | 0.7708 | 0.7728 | 0.7712 | 0.4493 | 0.2578 | 0.7885 | 0.6461 | 0.8421 | 0.6438 | 0.5709 | 0.4108 |
| NCBI682 | 0.6119 | 0.7103 | 0.4657 | 0.7008 | 0.5802 | 0.4575 | 0.6612 | 0.6603 | 0.7069 | 0.7024 | 0.6601 | 0.6534 | 0.4095 | 0.35 | 0.5793 | 0.5871 | 0.6447 | 0.6406 | 0.486 | 0.5394 |
| NCBI683 | 0.5412 | 0.5436 | 0.5416 | 0.5416 | 0.5635 | 0.5121 | 0.5592 | 0.5064 | 0.5524 | 0.5306 | 0.5417 | 0.5381 | 0.4845 | 0.4285 | 0.5521 | 0.5525 | 0.5802 | 0.5899 | 0.5276 | 0.4844 |
| NCBI760 | 0.3717 | 0.3749 | 0.3651 | 0.3651 | 0.3908 | 0.3893 | 0.3855 | 0.4026 | 0.4158 | 0.415 | 0.3725 | 0.3857 | 0.3851 | 0.3849 | 0.3828 | 0.4047 | 0.3027 | 0.2993 | 0.3265 | 0.2658 |
| NCBI764 | 0.2323 | 0.1949 | 0.1937 | 0.1937 | 0.1469 | 0.1622 | 0.1708 | 0.1562 | 0.2413 | 0.2058 | 0.2247 | 0.206 | 0.1296 | 0.1099 | 0.2004 | 0.186 | 0.2247 | 0.1588 | 0.0434 | -0.0266 |
| NCBI765 | 0.5045 | 0.4914 | 0.5457 | 0.5457 | 0.484 | 0.4948 | 0.552 | 0.556 | 0.5205 | 0.5341 | 0.509 | 0.5224 | 0.451 | 0.4308 | 0.5381 | 0.5323 | 0.4508 | 0.5549 | 0.425 | 0.4778 |
| NCBI766 | 0.4558 | 0.444 | 0.4514 | 0.4514 | 0.448 | 0.4489 | 0.4044 | 0.3588 | 0.3704 | 0.3907 | 0.442 | 0.4107 | 0.4413 | 0.4477 | 0.4641 | 0.4551 | 0.2525 | 0.425 | 0.4677 | 0.4315 |
| NCBI768 | 0.5423 | 0.5439 | 0.6093 | 0.6093 | 0.6552 | 0.6566 | 0.6053 | 0.5685 | 0.5994 | 0.5763 | 0.605 | 0.5675 | 0.5076 | 0.5136 | 0.4975 | 0.5242 | 0.685 | 0.6092 | 0.537 | 0.5521 |
| NCBI769 | 0.5001 | | 0.6326 | 0.6326 | 0.5566 | 0.6822 | 0.5838 | 0.558 | 0.598 | 0.548 | 0.4609 | 0.4686 | 0.5031 | 0.4983 | 0.5782 | 0.5433 | 0.6306 | 0.6184 | 0.4426 | 0.5332 |
| NCBI770 | -0.1996 | -0.0904 | -0.2631 | -0.2631 | -0.0994 | -0.0227 | -0.2183 | -0.1633 | 0.0678 | -0.159 | -0.1167 | -0.1483 | -0.2312 | -0.1781 | -0.2551 | -0.2375 | -0.1477 | -0.1666 | -0.2004 | -0.1857 |
| SPA100 | 0.4046 | 0.5788 | -0.1386 | -0.1386 | 0.2951 | 0.3088 | 0.3931 | 0.3295 | 0.6149 | 0.6731 | 0.5761 | 0.5571 | 0.0277 | -0.0143 | 0.5912 | 0.6114 | 0.5791 | 0.5755 | 0.0827 | 0.1206 |
| SPA101 | 0.4192 | 0.373 | 0.3731 | 0.3731 | 0.0927 | 0.232 | 0.366 | 0.422 | 0.388 | 0.3457 | 0.4524 | 0.4297 | 0.3041 | 0.3426 | 0.4294 | 0.4352 | 0.358 | 0.4416 | 0.2902 | 0.3061 |
| SPA102 | 0.4029 | 0.4161 | 0.3377 | 0.3377 | 0.3687 | 0.3479 | 0.4654 | 0.4571 | 0.0467 | 0.1713 | 0.399 | 0.4424 | 0.35 | 0.3297 | 0.4013 | 0.4199 | 0.3591 | 0.3973 | 0.3098 | |
| SPA103 | -0.2796 | -0.3661 | -0.3052 | -0.3052 | -0.304 | -0.3449 | -0.3816 | -0.4169 | -0.4081 | -0.4358 | -0.3717 | -0.3497 | -0.0402 | -0.2022 | -0.3473 | -0.3782 | -0.3393 | -0.3663 | -0.0249 | 0.0196 |
| SPA104 | 0.4311 | 0.4416 | 0.3084 | 0.3084 | 0.399 | 0.3797 | 0.4362 | 0.3734 | 0.4496 | 0.447 | 0.4518 | 0.4109 | 0.4111 | 0.3607 | 0.4503 | 0.4263 | 0.3995 | 0.3701 | 0.4076 | 0.3786 |
| SPA105 | 0.2124 | 0.175 | 0.0889 | 0.0889 | 0.0008 | 0.085 | 0.1213 | 0.1142 | 0.0932 | 0.083 | 0.0515 | 0.0453 | 0.1736 | 0.1186 | 0.125 | 0.1232 | 0.2349 | 0.211 | 0.0999 | 0.0584 |
| SPA106 | 0.1325 | 0.1657 | 0.1303 | 0.1303 | 0.1468 | 0.0657 | 0.0729 | 0.0991 | | 0.016 | 0.0478 | 0.0739 | 0.0724 | 0.0657 | 0.0274 | 0.0176 | 0.1152 | 0.1521 | 0.0219 | 0.0207 |
| SPA107 | 0.7098 | 0.6764 | 0.5726 | 0.5726 | 0.4695 | 0.5104 | 0.3541 | 0.5386 | 0.5688 | 0.6175 | 0.3637 | 0.4545 | 0.1253 | 0.1906 | 0.3653 | 0.525 | 0.6716 | 0.6722 | -0.1144 | 0.0801 |
| SPA108 | 0.6753 | 0.6554 | 0.6498 | 0.6498 | 0.6232 | 0.5517 | 0.6477 | 0.6573 | 0.6538 | 0.6566 | 0.6622 | 0.6504 | 0.6131 | 0.5628 | 0.6466 | 0.6226 | 0.6295 | 0.623 | 0.6246 | 0.6288 |
| SPA109 | 0.2514 | 0.2353 | 0.2611 | 0.2611 | 0.2159 | 0.1845 | 0.2203 | 0.2043 | 0.2038 | 0.2038 | | 0.2245 | 0.2132 | 0.2056 | 0.2246 | 0.2303 | 0.2126 | 0.2254 | 0.1608 | 0.1666 |
| SPA111 | 0.6582 | 0.6552 | 0.5817 | 0.5817 | 0.4654 | 0.4083 | 0.6098 | 0.6245 | 0.6541 | 0.6266 | 0.5427 | 0.5728 | 0.5139 | 0.5148 | 0.5469 | 0.4544 | 0.4807 | 0.5533 | 0.5488 | 0.5753 |
| SPA112 | 0.6808 | 0.6572 | 0.6061 | 0.6061 | 0.433 | 0.4638 | 0.6116 | 0.5915 | 0.6234 | 0.6057 | 0.6049 | 0.5805 | 0.45 | 0.4388 | 0.5395 | 0.3558 | 0.4203 | 0.5656 | 0.6175 | 0.5722 |
| SPA114 | 0.1448 | 0.047 | -0.0393 | -0.0393 | -0.038 | 0.0877 | -0.0509 | -0.0707 | -0.041 | 0.0136 | -0.1296 | -0.0382 | -0.0226 | 0.1027 | 0.0715 | 0.0546 | 0.1194 | 0.0368 | -0.0867 | -0.1462 |
| SPA115 | 0.1051 | 0.096 | 0.1169 | 0.1169 | 0.0906 | 0.0281 | 0.0756 | 0.0824 | 0.0388 | 0.048 | 0.0655 | 0.055 | 0.154 | 0.1349 | 0.0748 | 0.0693 | 0.0978 | 0.0743 | 0.1158 | 0.1278 |
| SPA116 | 0.5123 | 0.3561 | 0.1917 | 0.1917 | 0.1741 | -0.477 | -0.5197 | -0.429 | 0.547 | 0.6178 | -0.1159 | -0.256 | -0.6509 | -0.6645 | 0.9123 | 0.8889 | 0.8588 | 0.827 | -0.5516 | -0.4123 |
| SPA117 | -0.0688 | 0.0648 | 0.5675 | 0.5675 | 0.403 | 0.2184 | -0.3082 | -0.1602 | 0.3379 | 0.118 | -0.2689 | -0.245 | -0.2561 | -0.3838 | -0.0525 | -0.0407 | 0.7525 | 0.7086 | -0.304 | -0.2726 |
| SPA118 | 0.4907 | 0.4625 | 0.5087 | 0.5087 | 0.3282 | 0.3768 | 0.4441 | 0.4445 | 0.4658 | 0.4683 | 0.489 | 0.4522 | 0.4661 | 0.4929 | 0.496 | 0.5063 | 0.5527 | 0.5107 | 0.3848 | 0.4372 |
| SPA55 | 0.3528 | 0.2705 | 0.0716 | 0.0716 | 0.3702 | 0.314 | 0.3804 | 0.3705 | 0.22 | 0.0307 | 0.3387 | 0.223 | -0.0638 | -0.1494 | 0.4254 | 0.4336 | 0.2269 | 0.0676 | 0.3651 | 0.3036 |
| SPA57 | 0.3982 | 0.4651 | 0.2981 | 0.2981 | 0.3293 | 0.2787 | 0.4358 | 0.3881 | 0.4799 | 0.4555 | 0.423 | 0.4432 | 0.2187 | 0.1704 | 0.4 | 0.4329 | 0.3067 | 0.0772 | 0.1937 | 0.1039 |
| SPA60 | 0.4109 | 0.5733 | 0.3331 | 0.3331 | -0.1056 | 0.1224 | 0.481 | 0.5244 | 0.5127 | 0.4949 | 0.3247 | 0.4942 | 0.097 | 0.0561 | 0.3719 | 0.0608 | 0.5136 | 0.5866 | 0.0529 | 0.0281 |
| SPA61 | 0.6919 | 0.6296 | 0.3269 | 0.3269 | 0.0918 | 0.2482 | 0.4936 | 0.5422 | 0.5498 | 0.5755 | 0.3516 | 0.4662 | 0.2225 | 0.1803 | 0.0279 | 0.0754 | 0.7269 | 0.7459 | 0.2363 | 0.3112 |
| SPA64 | 0.6573 | 0.6699 | 0.5243 | 0.5243 | -0.357 | -0.2116 | -0.1401 | -0.1914 | 0.6497 | 0.6448 | 0.6865 | 0.6801 | -0.0892 | -0.2108 | 0.6265 | 0.5856 | 0.2026 | 0.5202 | -0.309 | 0.0511 |
| SPA66 | 0.564 | 0.4687 | -0.434 | -0.434 | -0.3864 | -0.4801 | -0.0253 | -0.3528 | 0.4358 | 0.429 | 0.6105 | 0.504 | -0.3846 | -0.2932 | -0.0441 | -0.3285 | 0.3401 | 0.066 | -0.3688 | -0.298 |
| SPA67 | 0.7629 | 0.5921 | 0.2713 | 0.2713 | -0.3697 | -0.3444 | -0.3296 | -0.3236 | 0.7276 | 0.5705 | 0.7587 | 0.6648 | -0.2812 | -0.2337 | 0.5126 | 0.46 | -0.2151 | -0.2115 | -0.0174 | 0.0055 |
| SPA68 | 0.7791 | 0.7801 | 0.1909 | 0.1909 | -0.044 | -0.1587 | 0.1552 | 0.1792 | 0.7169 | 0.6848 | 0.7218 | 0.7107 | 0.016 | -0.0691 | 0.523 | 0.6217 | 0.2055 | 0.0738 | -0.1369 | -0.0363 |
| SPA69 | 0.5527 | 0.4554 | 0.4585 | 0.4585 | 0.4298 | 0.3441 | 0.5599 | 0.5811 | 0.5265 | 0.4918 | 0.3421 | 0.284 | 0.4225 | 0.3915 | 0.6027 | 0.5707 | 0.6171 | 0.5291 | 0.4305 | 0.3757 |
| SPA73 | 0.9145 | 0.912 | 0.9177 | 0.9177 | 0.9176 | 0.9028 | 0.9015 | 0.9042 | 0.9189 | 0.922 | 0.9204 | 0.9202 | 0.8763 | 0.8007 | 0.9118 | 0.9155 | 0.9203 | 0.9215 | 0.8832 | 0.8939 |
| SPA80 | 0.0339 | 0.0478 | 0.033 | 0.033 | 0.0995 | 0.0724 | 0.0651 | 0.0662 | 0.0292 | 0.0218 | 0.0415 | 0.0381 | 0.0009 | -0.0126 | 0.1718 | 0.06 | 0.3023 | -0.0212 | 0.0148 | -0.0371 |
| SPA82 | 0.1503 | 0.2003 | 0.0785 | 0.0785 | 0.1391 | 0.1029 | 0.2615 | 0.279 | 0.1761 | 0.1387 | 0.1115 | 0.1249 | 0.2522 | 0.361 | 0.2229 | 0.0902 | 0.1445 | 0.0633 | 0.0259 | 0.2167 |
| SPA83 | 0.2971 | -0.0408 | -0.3274 | -0.3274 | -0.1514 | -0.3728 | -0.3141 | -0.418 | 0.2819 | 0.1582 | 0.5702 | 0.4804 | -0.3173 | -0.2967 | 0.1312 | -0.0052 | -0.5002 | -0.5058 | -0.5533 | -0.5437 |
| SPA84 | 0.5466 | 0.5234 | 0.4078 | 0.4078 | 0.2711 | 0.5013 | 0.5748 | 0.556 | 0.3466 | 0.3222 | 0.4237 | 0.3842 | 0.5094 | 0.4889 | 0.3797 | 0.2704 | 0.0593 | 0.2046 | 0.2732 | 0.1447 |
| SPA90 | 0.4637 | 0.5984 | 0.5506 | 0.5506 | 0.4285 | 0.3322 | 0.5499 | 0.6374 | 0.5004 | 0.5617 | 0.4185 | 0.4896 | 0.2171 | 0.2879 | 0.5069 | 0.6132 | 0.6736 | 0.553 | 0.3982 | 0.376 |
| SPA91 | 0.5945 | 0.4986 | 0.4513 | 0.4513 | 0.3577 | 0.3124 | 0.5125 | 0.4989 | 0.221 | 0.4097 | 0.4164 | 0.1183 | 0.5467 | 0.5148 | 0.4166 | 0.3739 | 0.6642 | 0.4969 | 0.4053 | 0.3153 |
| SPA92 | 0.2837 | 0.103 | -0.3247 | -0.3247 | -0.0016 | 0.02 | 0.3333 | 0.357 | 0.4111 | 0.3627 | 0.1801 | 0.2205 | 0.2932 | 0.3334 | 0.573 | 0.5809 | 0.4825 | 0.4186 | -0.1382 | 0.2735 |
| SPA93 | 0.3599 | 0.3695 | 0.3158 | 0.3158 | 0.2338 | 0.172 | 0.1238 | 0.1106 | 0.3164 | 0.2943 | 0.441 | 0.3991 | 0.0418 | 0.0219 | 0.2908 | 0.3382 | | 0.3394 | 0.2714 | 0.2618 |
| SPA94 | 0.392 | 0.3719 | -0.052 | -0.052 | -0.0108 | 0.1007 | -0.2373 | -0.2193 | 0.3397 | 0.3424 | 0.3276 | 0.3407 | -0.2324 | -0.1487 | 0.1648 | 0.2202 | 0.2936 | 0.2581 | -0.0558 | -0.0207 |
| SPA95 | 0.5206 | 0.5955 | 0.4181 | 0.4181 | 0.5001 | 0.4856 | 0.5844 | 0.6365 | 0.5665 | 0.5973 | 0.5362 | 0.5044 | 0.6402 | 0.5937 | 0.5485 | 0.5486 | | 0.5275 | 0.4951 | 0.5122 |
| SPA96 | 0.5228 | 0.4795 | 0.5283 | 0.5283 | 0.2921 | 0.2767 | 0.3683 | 0.404 | 0.5834 | 0.4521 | 0.4878 | 0.446 | 0.3564 | 0.4527 | 0.4026 | 0.4273 | 0.6844 | 0.617 | 0.3837 | 0.4867 |
| SPA97 | 0.6266 | 0.6161 | 0.6017 | 0.6017 | 0.4545 | 0.3943 | 0.5023 | 0.5471 | 0.7189 | 0.6856 | 0.5689 | 0.506 | 0.4681 | 0.4601 | 0.5581 | 0.5258 | 0.6809 | 0.7031 | 0.0864 | 0.1313 |
| SPA98 | 0.698 | 0.6114 | 0.5953 | 0.5953 | 0.3023 | 0.3975 | 0.5834 | 0.5476 | 0.6573 | 0.6442 | 0.6651 | 0.708 | 0.4823 | 0.5208 | 0.2047 | 0.2767 | 0.6309 | 0.5739 | 0.1999 | 0.2973 |
| SPA99 | 0.6098 | 0.5916 | 0.5198 | 0.5198 | 0.3708 | 0.3051 | 0.3831 | 0.4047 | 0.6078 | 0.5787 | 0.6095 | 0.5739 | 0.1034 | 0.1353 | 0.6122 | 0.5877 | 0.4919 | 0.4928 | 0.2995 | 0.3513 |
| TENX128 | 0.578 | | | | 0.3679 | | 0.5341 | | | | 0.5787 | | 0.5103 | | | | 0.532 | | 0.4696 | |
| TENX147 | 0.6863 | 0.656 | 0.6893 | 0.6893 | 0.5953 | 0.6495 | 0.7039 | 0.7012 | 0.5967 | 0.5615 | 0.7081 | 0.6925 | 0.0594 | 0.0423 | 0.7328 | 0.6404 | 0.4871 | 0.4475 | 0.4943 | -0.0608 |
| TENX148 | 0.5222 | 0.5922 | 0.5277 | 0.5277 | 0.3609 | 0.4161 | 0.6384 | 0.3721 | 0.4875 | 0.4409 | 0.3928 | 0.5729 | 0.5877 | 0.3617 | | 0.4661 | 0.244 | 0.4051 | 0.375 | 0.0654 |
| TENX149 | 0.3089 | 0.381 | 0.4584 | 0.4584 | 0.3143 | 0.3465 | 0.4301 | 0.4173 | 0.3549 | 0.4237 | | 0.4518 | 0.1868 | 0.2234 | 0.3008 | 0.3681 | 0.332 | 0.3734 | 0.4191 | 0.1514 |
| TENX152 | 0.7223 | 0.7264 | 0.5822 | 0.5822 | 0.7787 | 0.6314 | 0.6355 | 0.5379 | 0.7218 | 0.7205 | 0.7346 | 0.6911 | 0.1712 | 0.4445 | | 0.2523 | 0.8415 | 0.5187 | 0.5273 | 0.2826 |
| TENX157 | 0.7652 | 0.5942 | 0.5361 | 0.5361 | 0.7414 | 0.4285 | 0.747 | 0.2513 | 0.6065 | 0.4516 | 0.7922 | 0.654 | 0.1839 | 0.0203 | 0.7278 | 0.3632 | 0.8052 | 0.3993 | 0.253 | 0.1339 |
| TENX30 | 0.1155 | 0.1275 | 0.1955 | 0.1955 | 0.1445 | 0.1436 | 0.1495 | 0.0905 | 0.1316 | 0.1679 | 0.112 | 0.2121 | 0.1153 | 0.0581 | 0.1221 | 0.0205 | 0.1979 | 0.1181 | 0.1062 | 0.0977 |
| TENX31 | 0.1129 | 0.0528 | 0.1047 | 0.1047 | 0.1142 | 0.0414 | 0.1799 | 0.0907 | 0.2284 | 0.181 | 0.1259 | 0.1898 | 0.1606 | 0.0728 | 0.1584 | -0.0544 | 0.2244 | 0.0295 | 0.1304 | 0.0258 |
| TENX46 | 0.4086 | 0.043 | 0.4664 | 0.5082 | 0.5563 | 0.3372 | 0.3478 | 0.0903 | | | 0.4107 | | -0.0494 | -0.0081 | 0.3893 | 0.1697 | 0.1707 | | 0.1998 | 0.0506 |
| TENX70 | 0.5749 | | 0.6643 | 0.6632 | 0.7802 | 0.659 | 0.5587 | 0.5764 | 0.5798 | 0.5694 | 0.6246 | 0.5416 | 0.3114 | 0.3985 | | 0.4171 | 0.7138 | 0.5057 | 0.5753 | 0.2893 |
| TENX90 | 0.4006 | 0.4589 | 0.5103 | 0.5103 | 0.4175 | 0.4943 | 0.3123 | 0.4768 | 0.3794 | 0.572 | 0.4372 | 0.5474 | 0.3426 | 0.0558 | 0.45 | 0.2774 | 0.4162 | 0.4059 | 0.3078 | 0.0871 |
| TENX91 | 0.7689 | 0.6362 | 0.626 | 0.626 | -0.0942 | -0.1844 | 0.5037 | 0.3963 | 0.7445 | 0.6768 | 0.7171 | 0.8065 | -0.1703 | 0.1319 | 0.6518 | 0.2206 | 0.6935 | 0.7008 | -0.2033 | 0.1557 |
| TENX92 | 0.7244 | 0.6807 | 0.577 | 0.577 | 0.7098 | 0.5398 | 0.721 | 0.5364 | 0.7004 | 0.406 | 0.7446 | 0.7392 | 0.5139 | 0.4097 | 0.6842 | 0.4058 | 0.6953 | 0.62 | 0.3347 | 0.2475 |
| ZEN38 | 0.6206 | 0.8076 | 0.8268 | 0.8268 | 0.8033 | 0.7967 | 0.8472 | 0.7523 | 0.7053 | 0.7957 | 0.697 | 0.8432 | 0.6371 | -0.1179 | 0.7086 | 0.8439 | 0.623 | 0.565 | 0.5845 | 0.0695 |
| ZEN42 | 0.6927 | 0.6764 | 0.6709 | 0.671 | 0.7018 | 0.6712 | 0.722 | 0.7128 | 0.7054 | 0.6473 | 0.6555 | 0.6751 | 0.5831 | 0.4288 | 0.6731 | 0.6961 | 0.6411 | 0.6805 | 0.6883 | 0.4121 |
| ZEN44 | 0.7862 | 0.7245 | 0.687 | 0.687 | 0.7047 | 0.6668 | 0.6868 | 0.7003 | 0.7385 | 0.714 | 0.7178 | 0.7153 | 0.4789 | 0.4438 | 0.7617 | 0.7358 | 0.7556 | 0.6205 | 0.516 | 0.5922 |
| ZEN45 | 0.5145 | 0.7363 | 0.6785 | 0.6785 | 0.7149 | 0.6728 | 0.6385 | 0.7379 | 0.6963 | 0.7127 | 0.5365 | 0.7016 | 0.6635 | 0.4637 | 0.7182 | 0.6905 | 0.6998 | 0.7126 | 0.4305 | 0.6547 |
| ZEN78 | 0.3808 | 0.5498 | 0.4747 | 0.4747 | 0.4416 | 0.2796 | 0.2169 | 0.5016 | 0.3592 | 0.5651 | 0.2817 | 0.3004 | 0.5357 | 0.5558 | 0.4278 | 0.4603 | 0.0393 | 0.5353 | 0.5896 | 0.5029 |
| ZEN80 | 0.7382 | 0.7323 | 0.723 | 0.723 | 0.6973 | 0.6539 | 0.7832 | 0.6862 | 0.8209 | 0.772 | 0.8398 | 0.7584 | 0.6661 | 0.6496 | 0.7839 | 0.7223 | 0.8384 | 0.7213 | 0.7148 | 0.6604 |
| ZEN81 | 0.6829 | 0.7031 | 0.6026 | 0.6026 | 0.5545 | 0.4745 | 0.6976 | 0.6923 | 0.7044 | 0.6981 | 0.6895 | 0.6505 | -0.5885 | 0.6628 | 0.6455 | 0.6566 | | 0.5528 | 0.5684 | 0.6228 |
| INT12 | | 0.3572 | 0.1855 | 0.1855 | 0.1336 | 0.1637 | 0.2253 | -0.0339 | 0.5375 | 0.4921 | 0.8099 | 0.0998 | 0.056 | 0.0303 | 0.8211 | 0.1742 | | -0.0383 | 0.0654 | -0.0801 |
| MEND148 | | 0.6919 | 0.6131 | 0.613 | 0.6071 | 0.6634 | 0.4122 | 0.6471 | 0.6338 | 0.7729 | 0.5596 | 0.6989 | 0.4139 | 0.5927 | 0.6926 | 0.7377 | 0.6475 | 0.7508 | 0.5367 | 0.5628 |
| MEND65 | | 0.8315 | 0.8423 | 0.8423 | 0.8429 | 0.832 | 0.828 | 0.8184 | | 0.8359 | 0.8574 | 0.8313 | 0.8028 | 0.7765 | 0.8492 | 0.8304 | 0.8526 | 0.835 | 0.7885 | 0.7869 |
| MISC42 | | 0.6519 | 0.5941 | 0.5941 | 0.7937 | 0.6307 | 0.5794 | 0.4421 | | 0.4577 | | 0.7691 | 0.3746 | 0.3645 | 0.8092 | 0.5282 | 0.754 | 0.5902 | 0.5931 | 0.3461 |
| MISC57 | | -0.2166 | 0.3356 | 0.3356 | 0.4603 | 0.4536 | -0.3661 | -0.299 | -0.4599 | 0.5231 | | -0.4397 | 0.3237 | 0.4558 | | 0.4371 | | -0.0416 | -0.2033 | 0.3884 |
| MISC58 | | 0.577 | -0.0101 | -0.0101 | 0.4085 | 0.0219 | 0.6662 | 0.2634 | 0.7971 | 0.317 | 0.6804 | 0.7113 | -0.2138 | -0.1405 | 0.7911 | 0.2728 | | 0.5203 | 0.3216 | -0.0358 |
| TENX40 | | 0.7229 | 0.7083 | 0.6776 | 0.7274 | 0.6537 | 0.712 | 0.6417 | 0.7929 | 0.7487 | 0.7697 | 0.7397 | 0.5729 | 0.4689 | 0.8168 | 0.7174 | | 0.7025 | 0.4294 | 0.2859 |
| ZEN79 | | 0.6779 | 0.7024 | 0.7025 | 0.6258 | 0.628 | 0.712 | 0.7003 | 0.7455 | 0.7288 | 0.7716 | 0.7245 | | 0.6979 | | 0.7557 | 0.732 | 0.6887 | 0.574 | 0.6444 |
| MEND156 | | | 0.6143 | 0.6143 | 0.639 | 0.6691 | 0.6148 | 0.7571 | 0.6839 | | 0.7216 | 0.6671 | 0.6879 | 0.6537 | 0.7487 | 0.6909 | | 0.505 | 0.6914 | 0.5965 |
| MEND141 | | | | | | 0.7513 | | | | | | | 0.5488 | 0.5419 | | | | 0.6314 | | 0.2615 |

### Table.S6

| Large Region | Positively Correlated Genes (late-enriched genes) | GigaPath | UNI2h | ViTB16 | RN18 |
|---|---|---|---|---|---|
| | Colon | 61<br>ACTG1,ASS1,ATP5F1B,ATP8B1,C19orf33,CDH1,CEACAM5,CEACAM6,CFL1,CLDN3,CLDN4,CLDN7,COX7A2,EEF1B2,ELF3,EPCAM,ERBB3,FABP1,FERMT1,FXYD3,GPX2,HNF4A,HSPA5,ITGB4,KLF5,KRT8,LGALS3,LGALS3BP,LGALS4,LRATD1,LSR,MGAT4B,MUC13,MUC5B,MYH14,NQO1,PHGR1,PKM,PKP3,PLPP2,PPDPF,PPP1R1B,PRR15L,RAB25,RNF43,S100A10,S100A14,S100A6,S100P,SELENBP1,SERF2,SLC12A2,SOX9,SPINT2,ST14,TFF1,TFF3,TMEM54,TMPRSS4,TSPAN8,USH1C | 60<br>ACTG1,ASS1,ATP8B1,C19orf33,CDH1,CEACAM5,CEACAM6,CHCHD2,CLDN3,CLDN4,CLDN7,COX7A2,DDR1,ELF3,EPCAM,ERBB3,FABP1,FXYD3,GPX2,HNF4A,HSPA5,ITGB4,KRT8,LGALS3,LGALS3BP,LGALS4,LRATD1,LSR,MAL2,MGAT4B,MUC13,MYH14,NQO1,P4HB,PHGR1,PKP3,PLPP2,PPDPF,PPP1R1B,PRR15L,RAB25,RNF43,S100A10,S100A14,S100A6,S100P,SELENBP1,SERF2,SLC12A2,SOX9,SPINT2,ST14,STARD10,TFF3,TMEM54,TMPRSS4,TMSB10,TPD52,TSPAN8,VDAC1 | 37<br>ATP8B1,C19orf33,CDCA7,CDH1,CEACAM1,CEACAM5,CEACAM6,CLDN3,CLDN4,ELF3,EPCAM,FERMT1,GPRC5A,GPX2,HIST1H1C,KRT8,LGALS4,LRATD1,MUC13,MUC5B,MYH14,NQO1,PHGR1,PLPP2,PPDPF,PPP1R1B,RAB25,RNF43,S100A14,S100A6,SELENBP1,SERF2,SLC12A2,SOX9,TFF1,TFF3,TSPAN8 | 30<br>ACTG1,CEACAM1,CEACAM5,CEACAM6,CLDN3,CLDN7,CTNND1,EPCAM,ERBB3,GPX2,KRT8,LGALS4,LSR,MGAT4B,MUC13,MYH14,NQO1,P4HB,PKM,PKP3,PPDPF,PRR15L,RAB25,RNF43,S100A6,SERF2,SOX9,ST14,TAGLN2,TFF3 |
| | Prostate | 74<br>ABCC4,ACLY,ACSL3,ADI1,ARG2,ATF4,ATP1A1,ATP5MC2,CANX,CD9,CDH1,CIRBP,CREB3L4,DBI,DCXR,DDX5,DHRS7,EEF2,EIF4A1,EPCAM,FASN,FLNB,FOXA1,FTH1,GOLPH3,H2AFJ,HINT1,HNRNPA2B1,HOXB13,HSPA8,KDELR2,KLK2,KLK3,KLK4,KRT18,KRT8,MLPH,MPC2,NEDD4L,NKX3-1,NPY,ODC1,P4HB,PABPC1,PARK7,PLPP1,PMEPA1,PPP3CA,PTGR1,PTPRF,RAB11A,RACK1,RDH11,SEC11C,SERF2,SERP1,SLC45A3,SMS,SPDEF,SPOCK1,SPON2,STEAP1,STEAP2,SYNGR2,TACSTD2,TMEM258,TMPRSS2,TPD52,TRPM4,TSPAN1,TSTD1,UBA52,UQCRQ,XBP1 | 68<br>ABCC4,ACLY,ACPP,ADI1,APLP2,ARF1,ATF4,ATP1A1,ATP5MC2,CD9,CDH1,CREB3L4,DBI,DCXR,DHRS7,EEF2,EIF4A1,EPCAM,FASN,FLNB,FOXA1,FTH1,GOLPH3,H2AFJ,HINT1,HOXB13,HSP90AA1,HSPA8,HSPE1,KDELR2,KLK2,KLK3,KLK4,KRT18,KRT8,MARCKSL1,MLPH,MPC2,NEDD4L,NKX3-1,ODC1,P4HB,PABPC1,PARK7,PMEPA1,PPP3CA,PTPRF,RAB11A,RACK1,RDH11,SEC11C,SERP1,SLC45A3,SMS,SPDEF,SPOCK1,SPON2,STEAP1,SYNGR2,TACSTD2,TMEM258,TMPRSS2,TPD52,TRPM4,TSPAN1,TSTD1,UQCRQ,XBP1 | 51<br>ABCC4,ACLY,APLP2,ATF4,CANX,CIRBP,CREB3L4,DHRS7,EIF4A1,FAAH,FASN,FLNB,FOXA1,FTH1,H2AFJ,HINT1,HNRNPA2B1,HOXB13,HSP90AA1,HSPA8,HSPE1,KLK2,KLK3,KRT18,MARCKSL1,MLPH,MPC2,NEDD4L,NKX3-1,NPY,ODC1,PABPC1,PARK7,PLPP1,PMEPA1,PPP3CA,PTGR1,RACK1,RDH11,SEC11C,SLC39A6,SMS,SPOCK1,SPON2,STEAP1,STEAP2,TACSTD2,TMPRSS2,TSPAN1,UBA52,UQCRQ | 65<br>ABCC4,ACLY,ACPP,ACSL3,APLP2,ARF1,ARG2,ATP1A1,CANX,CD9,CIRBP,CLDN3,CPNE4,CREB3L4,DCXR,DDX5,DHRS7,ELOVL5,EPCAM,FASN,FLNB,FOXA1,GOLPH3,H2AFJ,HINT1,HNRNPA2B1,HNRNPC,HOXB13,HSPA8,HSPE1,KDELR2,KLK2,KLK3,KLK4,KRT18,MLPH,MPC2,NEDD4L,NKX3-1,NPY,ODC1,P4HB,PABPC1,PARK7,PLPP1,PMEPA1,PPP3CA,RAB11A,RACK1,RDH11,SEC11C,SERP1,SLC39A6,SMS,SPOCK1,STEAP1,STEAP2,TACSTD2,TMEM258,TMPRSS2,TPD52,TRPM4,TSPAN1,UQCRQ,XBP1 |
| | Breast | 44<br>ACTB,ACTG1,ASAH1,ATP5F1A,ATP6AP2,BEST1,CALR,CD24,CD46,CDH1,DDX5,DUSP4,EIF4G2,ERBB2,FASN,GATA3,GNAS,H1-0,H3-3B,HNRNPA2B1,HSP90AB1,HSPA1A,HSPB1,KRT19,MORF4L2,PRDX1,PSMD3,PTMA,RPL23,RPL31,RPL5,S100A11,SARAF,SLC39A6,SLC7A2,SPINT2,SREBF1,SRRM2,TACSTD2,TAGLN2,TMSB10,VMP1,XBP1,YWHAZ | 40<br>ACTB,ACTG1,AREG,ASAH1,ATP5F1A,ATP6AP2,CALR,CD24,CD46,COX6C,DDX5,DUSP4,EIF4G2,ERBB2,FASN,GATA3,GNAS,H1-0,H3-3B,HNRNPA2B1,HSP90AB1,HSPB1,KRT19,MORF4L2,PRDX1,PSMD3,PTMA,PTPRF,RP11-53O19.3,RPL23,S100A11,SDC1,SLC7A2,SPINT2,SREBF1,SRRM2,TACSTD2,TAGLN2,XBP1,YWHAZ | 27<br>ACTB,ACTG1,ATP5F1A,ATP6AP2,CD46,CDH1,COX6C,DDX5,DUSP4,ESR1,FASN,GATA3,GNAS,H3-3B,HNRNPA2B1,HSP90AB1,HSPA1A,HSPB1,KRT19,MAGED2,MCCC2,MORF4L2,NEAT1,PTMA,SLC39A6,TACSTD2,XBP1 | 25<br>ASAH1,ATP5F1A,ATP6AP2,BEST1,CD46,COX6C,DDX5,DUSP4,EIF4A1,FASN,H3-3B,HNRNPA2B1,HNRNPH1,MAGED2,MCCC2,MORF4L2,NFKBIZ,PLXNB2,PSMD3,PTMA,RPL31,SARAF,SRRM2,TRPS1,XBP1 |
| | Lymph | 41<br>ANGPTL4,ASS1,ATP1B1,BCAP31,CANX,CD24,CLU,CP,CRYAB,CXCL14,CYB5A,ENO1,EPCAM,ERO1A,FTH1,FXYD2,HILPDA,HSPB1,ITM2B,MGST1,MIF,NDRG1,P4HB,PABPC1,PDZK1IP1,PGRMC1,PLOD2,PPDPF,PPIB,PTTG1IP,S100A10,SCD,SERF2,SERINC2,SERPINA1,TMBIM6,TXN,UBC,UQCRQ,VEGFA,WFDC2 | 31<br>ASS1,ATP1B1,AZGP1,CD24,CLU,CP,CRYAB,CXCL14,DDR1,ENO1,EPCAM,FXYD2,HSPB1,MGST1,MIF,MT1G,NDRG1,NETO2,PDZK1IP1,PGRMC1,PLOD2,PPIB,S100A10,S100A11,SCD,SERF2,SERINC2,SPINT2,UBC,VEGFA,WFDC2 | 37<br>APOL1,ATP1B1,AZGP1,C3,CA12,CALR,CLU,CP,CRYAB,CYB5A,DHCR24,EPCAM,ERO1A,FKBP5,FTH1,GAPDH,GPX3,HILPDA,HNRNPA2B1,HSPB1,MIF,P4HB,PABPC1,PDE10A,PPDPF,S100A10,S100A11,SCD,SERF2,SERPINA1,SPINT2,SPP1,TMBIM6,TMSB10,TPD52,UBC,VEGFA | 29<br>APOL1,ATP1B1,AZGP1,C3,CLU,CP,CRYAB,CXCL14,DHCR24,EPCAM,GPX3,HILPDA,MGST1,MIF,NDRG1,P4HB,PDZK1IP1,PIGR,PLOD2,PPDPF,PPIB,S100A10,S100A11,SCD,SOD2,SPINT2,SPP1,TMBIM6,UBC |
| | Negatively Correlated Genes (early-enriched genes) | GigaPath | UNI2h | ViTB16 | RN18 |
| | Colon | 90<br>A2M,ACTA2,ACTG2,ADAM33,AEBP1,BGN,BOC,C11orf96,C1R,C1S,C3,CALD1,CAVIN1,CCDC80,CCN1,CHRDL2,CLU,CNN1,COL14A1,COL16A1,COL1A1,COL1A2,COL3A1,COL6A2,COL6A3,COL8A1,COMP,CRISPLD2,CRYAB,CXCL12,DCN,DES,DPYSL3,EFEMP1,ELN,EMILIN1,FBLN1,FBLN2,FBN1,FERMT2,FLNA,FN1,FNDC1,FSTL1,FXYD6,GREM1,HSPB6,HTRA1,IGFBP5,IGFBP7,IGHG1,ISLR,LIMS2,LMOD1,LUM,MFAP4,MFAP5,MGP,MMP2,MRVI1,MXRA8,MYH11,MYL9,MYLK,NNMT,PDLIM3,PLN,PODN,PRELP,PTGDS,PTGIS,SERPINF1,SERPING1,SFRP2,SFRP4,SPARC,SPARCL1,SSC5D,SULF1,SYNM,SYNPO2,TAGLN,THBS2,THBS4,THY1,TIMP2,TNS1,TNXB,TPM2,VIM | 87<br>A2M,ACTA2,ACTG2,AEBP1,ANTXR1,BGN,BOC,C11orf96,C1R,C1S,C3,CALD1,CAVIN1,CCDC80,CCN1,CHRDL2,CLU,CNN1,COL14A1,COL16A1,COL1A1,COL1A2,COL3A1,COL6A2,COL6A3,COL8A1,COMP,CRISPLD2,CRYAB,DCN,DES,DPYSL3,EFEMP1,ELN,EMILIN1,FBLN1,FBLN2,FBN1,FERMT2,FLNA,FN1,FNDC1,FSTL1,FXYD6,GREM1,HSPB6,HTRA1,IGFBP5,IGFBP7,IGHG1,ISLR,LIMS2,LMOD1,LUM,MFAP4,MGP,MMP2,MRVI1,MXRA8,MYH11,MYL9,MYLK,NNMT,PDLIM3,PLN,PODN,PRELP,PTGDS,PTGIS,SERPINF1,SERPING1,SFRP2,SFRP4,SPARC,SPARCL1,SSC5D,SULF1,SYNPO2,TAGLN,THBS2,THBS4,THY1,TIMP2,TNS1,TNXB,TPM2,VIM | 77<br>ACTA2,ACTG2,AEBP1,ANTXR1,BGN,BOC,C11orf96,C1R,C1S,C3,CALD1,CAVIN1,CCDC80,CCN1,CNN1,COL14A1,COL16A1,COL1A1,COL1A2,COL3A1,COL6A2,COL6A3,COL8A1,COMP,CRISPLD2,DCN,DES,DPYSL3,EFEMP1,ELN,EMILIN1,FBLN1,FBLN2,FBN1,FERMT2,FLNA,FN1,FNDC1,FSTL1,GREM1,HTRA1,IGFBP5,IGFBP7,ISLR,LIMS2,LMOD1,LUM,MFAP4,MGP,MMP2,MRVI1,MXRA8,MYH11,MYL9,MYLK,NNMT,PDLIM3,PODN,PRELP,PTGIS,SERPINF1,SERPING1,SFRP2,SFRP4,SPARC,SPARCL1,SSC5D,SULF1,SYNPO2,TAGLN,THBS2,THY1,TIMP2,TNS1,TPM2,VCAN,VIM | 64<br>A2M,ACTA2,ACTG2,AEBP1,ANTXR1,BGN,C11orf96,C1R,C1S,C3,CALD1,CAVIN1,CCDC80,CCN1,CNN1,COL16A1,COL1A1,COL1A2,COL3A1,COL6A2,COL6A3,COL8A1,COMP,CRISPLD2,DCN,DPYSL3,EMILIN1,FBLN1,FBLN2,FBN1,FLNA,FN1,FSTL1,GREM1,IGFBP5,IGFBP7,ISLR,LIMS2,LMOD1,LUM,MGP,MMP2,MXRA8,MYH11,MYL9,MYLK,NNMT,PDLIM3,SERPINF1,SERPING1,SFRP2,SFRP4,SPARC,SPARCL1,SULF1,SYNPO2,TAGLN,THBS2,THY1,TIMP2,TNS1,TPM2,VCAN,VIM |
| | Prostate | 92<br>ACTA2,ACTC1,ACTG2,ADGRA2,AEBP1,AL078639.1,AOC3,C1R,C1S,CALD1,CARMN,CAV1,CAVIN1,CCDC80,CHRDL1,CLU,CNN1,COL16A1,COL1A2,COL4A2,COL6A1,COL6A2,COL6A3,COX7A1,CRYAB,CSRP1,DCN,DES,EMILIN1,FBLN1,FHL1,FILIP1L,FLNA,FLNC,FXYD1,FXYD6,GSN,HSPB6,HSPB8,IGFBP6,INMT,ITGA5,JPH2,KCNMB1,LGALS1,LIMS2,LMOD1,MAP1B,MASP1,MFGE8,MGP,MRGPRF,MRVI1,MSRB3,MXRA7,MYH11,MYL9,MYLK,NEXN,OGN,PAGE4,PALLD,PCP4,PDE5A,PDLIM3,PDLIM7,PGM5,PGM5-AS1,PLN,PPP1R14A,PRELP,PTGDS,PTGIS,RARRES2,RASL12,RBPMS,S100A6,SELENOM,SERPINF1,SERPING1,SMOC1,SMTN,SORBS1,SPARCL1,SYNM,SYNPO2,TAGLN,TGFB1I1,TNS1,TPM1,TPM2,WFDC1 | 91<br>ACTA2,ACTC1,ACTG2,ADGRA2,AEBP1,AOC3,C1R,C1S,C2orf40,CALD1,CARMN,CAVIN1,CCDC80,CHRDL1,CLU,CNN1,COL16A1,COL1A2,COL4A2,COL6A1,COL6A2,COL6A3,COX7A1,CRYAB,CSRP1,DCN,DES,EMILIN1,FBLN1,FHL1,FLNA,FLNC,FXYD1,FXYD6,GSN,HSPB6,HSPB8,IGFBP6,INMT,ITGA5,JPH2,KCNMB1,LGALS1,LIMS2,LMOD1,MAP1B,MASP1,MFAP4,MFGE8,MGP,MRGPRF,MRVI1,MSRB3,MXRA7,MYH11,MYL9,MYLK,NEXN,OGN,PAGE4,PALLD,PCP4,PDE5A,PDLIM3,PDLIM7,PGM5,PGM5-AS1,PLN,PPP1R14A,PRELP,PTGDS,PTGIS,RARRES2,RASL12,RBPMS,S100A6,SELENOM,SERPINF1,SERPING1,SMOC1,SMTN,SORBS1,SPARCL1,SYNM,SYNPO2,TAGLN,TGFB1I1,TNS1,TPM1,TPM2,WFDC1 | 85<br>ACTA2,ACTG2,ADGRA2,AEBP1,AOC3,C1R,C1S,C2orf40,CALD1,CARMN,CAVIN1,CCDC80,CHRDL1,CLU,CNN1,COL16A1,COL1A2,COL4A2,COL6A1,COL6A2,COL6A3,COX7A1,CRYAB,CSRP1,DCN,DES,EMILIN1,FBLN1,FHL1,FLNA,FLNC,FXYD1,GSN,HSPB6,HSPB8,IGFBP6,INMT,ITGA5,JPH2,KCNMB1,LGALS1,LIMS2,LMOD1,MAP1B,MASP1,MFGE8,MGP,MRGPRF,MRVI1,MSRB3,MXRA7,MYH11,MYL9,MYLK,NEXN,OGN,PAGE4,PALLD,PCP4,PDLIM3,PDLIM7,PGM5,PLN,PPP1R14A,PRELP,PTGDS,PTGIS,RARRES2,RASL12,RBPMS,S100A6,SELENOM,SERPINF1,SERPING1,SMTN,SORBS1,SPARCL1,SYNM,SYNPO2,TAGLN,TGFB1I1,TNS1,TPM1,TPM2,WFDC1 | 87<br>ACTA2,ACTC1,ACTG2,ADGRA2,AEBP1,C1R,C1S,C2orf40,CALD1,CARMN,CAV1,CAVIN1,CCDC80,CHRDL1,CLU,CNN1,COL16A1,COL1A2,COL4A2,COL6A1,COL6A2,COL6A3,COX7A1,CSRP1,DCN,DES,EMILIN1,FBLN1,FHL1,FLNA,FLNC,FXYD1,FXYD6,GSN,HSPB6,HSPB8,IGFBP6,INMT,ITGA5,JPH2,KCNMB1,LAMA4,LGALS1,LIMS2,LMOD1,MAP1B,MFAP4,MFGE8,MGP,MRGPRF,MRVI1,MSRB3,MXRA7,MYH11,MYL9,MYLK,NEXN,OGN,PAGE4,PALLD,PCP4,PDLIM3,PDLIM7,PGM5,PGM5-AS1,PPP1R14A,PRELP,PTGDS,PTGIS,RARRES2,RASL12,RBPMS,S100A6,SELENOM,SERPINF1,SERPING1,SMTN,SORBS1,SPARCL1,SYNM,SYNPO2,TAGLN,TGFB1I1,TNS1,TPM1,TPM2,WFDC1 |
| | Breast | 78<br>AEBP1,APOC1,APOE,BGN,C1QA,C1QB,C1QC,C1R,C3,CAVIN1,CCL19,CCL21,CCL5,CD14,CD2,CD27,CD52,CD6,CD68,CD74,CD79A,CLDN5,COL6A2,COMP,CORO1A,CSF1R,CTSD,CYBA,CYTH4,DCN,DERL3,EMILIN1,EMP3,ENG,GPSM3,GSTP1,HCST,HLA-DMA,HLA-DPA1,HLA-DPB1,HLA-DQB1,HLA-DRB1,HLA-E,IGFBP4,IGLL1,IGLL5,IL32,JAK3,JCHAIN,LAPTM5,LGALS1,LSP1,LTB,MMP2,MMP9,MYL9,MZB1,NNMT,PCOLCE,PI16,PLEKHO1,POU2AF1,PTGDS,RARRES2,RPL18A,RPLP1,S100A4,SELENOM,SERPINF1,SSR4,STAB1,TAGLN,THY1,TIMP1,TMEM119,TPSB2,VIM,VWF | 73<br>AEBP1,APOC1,APOE,BGN,C1QA,C1QB,C1QC,C1R,C3,CAVIN1,CCL19,CCL21,CCL5,CD14,CD2,CD27,CD52,CD6,CD68,CD74,CD79A,CLDN5,COL6A2,COMP,CORO1A,CSF1R,CTSD,CYBA,DCN,EMILIN1,EMP3,ENG,GPSM3,GSTP1,HLA-DMA,HLA-DPA1,HLA-DPB1,HLA-DQB1,HLA-DRB1,HLA-E,IGFBP4,IGLL1,IGLL5,IL32,JAK3,JCHAIN,LAPTM5,LGALS1,LSP1,LTB,MMP2,MMP9,MYL9,MZB1,PCOLCE,PECAM1,PTGDS,RARRES2,RPL18A,RPLP1,S100A4,SELENOM,SERPINF1,SFRP4,SSR4,STAB1,TAGLN,TGFB1,THY1,TIMP1,TMEM119,VIM,VWF | 36<br>AEBP1,APOC1,APOE,BGN,C1QA,C3,CAVIN1,CCL5,CD52,CD6,CD68,CD74,CD79A,CTSD,CYBA,ENG,GSTP1,HLA-DMA,HLA-DPA1,HLA-E,IGFBP4,IGLL1,IGLL5,LAPTM5,LSP1,LTB,MMP9,MYL9,MZB1,RPL18A,RPLP1,SELENOM,SSR4,TAGLN,TIMP1,VWF | 41<br>AEBP1,APOC1,APOE,BGN,C1QA,C3,CAVIN1,CCL19,CCL5,CD2,CD6,CD68,CD79A,COL6A2,COMP,CTSD,CYBA,ENG,GPSM3,GSTP1,HLA-DPA1,IGFBP4,IGLL1,IGLL5,IL32,JAK3,JCHAIN,LAPTM5,LSP1,MMP9,MYL9,MYO1F,MZB1,RPL18A,RPLP1,S100A4,SELENOM,SSR4,TAGLN,VWF,ZAP70 |
| | Lymph | 72<br>A2M,ACTA2,AEBP1,BGN,C7,CAVIN1,CCDC3,CCDC80,CCN1,CCN2,CD163,CD93,COL15A1,COL16A1,COL1A1,COL1A2,COL3A1,COL4A1,COL4A2,COL5A1,COL6A2,COL6A3,CRISPLD2,DCN,EFEMP1,ELN,EMILIN1,ENG,FBLN1,FBLN2,FBLN5,FBN1,FN1,FOS,FOSB,FSTL1,GGT5,HSPG2,ID3,IGFBP7,IGHG1,IGHM,IGKC,IL7R,JCHAIN,LHFPL6,LTBP2,LUM,MCAM,MFAP4,MFGE8,MGP,MMP2,MYL9,OLFML2B,PDGFRB,PECAM1,PPP1R14A,PRELP,PRSS23,PTPRC,SERPINE1,SERPINF1,SLIT3,SPARC,TAGLN,THBS1,THBS2,TIMP2,TPM2,VCAN,VWF | 72<br>A2M,ACTA2,AEBP1,BGN,C7,CAVIN1,CCDC3,CCDC80,CCN1,CCN2,CD163,CD93,CDH11,COL15A1,COL1A1,COL1A2,COL3A1,COL4A1,COL4A2,COL6A2,COL6A3,CRISPLD2,CXCL12,DCN,EFEMP1,ELN,EMILIN1,ENG,FBLN1,FBLN2,FBLN5,FBN1,FN1,FOS,FOSB,FSTL1,GGT5,HSPG2,ID3,IGFBP7,IGHG1,IGHM,IGKC,IL7R,LHFPL6,LTBP2,LUM,MCAM,MFAP4,MFGE8,MGP,MMP2,MYL9,OLFML2B,PDGFRA,PDGFRB,PECAM1,PPP1R14A,PRELP,PRSS23,PTPRC,SERPINE1,SERPINF1,SLIT3,SPARC,TAGLN,THBS1,THBS2,TIMP2,TPM2,VCAN,VWF | 68<br>A2M,ACTA2,ADAMTS2,AEBP1,ANTXR1,BGN,CAVIN1,CCDC3,CCDC80,CCN1,CCN2,CD163,CD93,CDH11,CNN1,COL12A1,COL15A1,COL16A1,COL1A1,COL1A2,COL3A1,COL4A1,COL4A2,COL5A1,COL5A2,COL6A2,COL6A3,CRISPLD2,DCN,DKK3,EMILIN1,FBLN1,FBLN2,FBN1,FN1,FOS,FSTL1,GGT5,HSPG2,ID3,IGFBP7,IL7R,INHBA,LHFPL6,LTBP2,LUM,MCAM,MFGE8,MMP2,MYL9,OLFML2B,PDGFRB,PDLIM3,PECAM1,PPP1R14A,PRELP,PRSS23,SERPINE1,SLIT3,SPARC,SULF1,TAGLN,THBS2,TIMP2,TIMP3,TPM2,VCAN,VWF | 67<br>A2M,ACTA2,AEBP1,ANTXR1,BGN,C7,CAVIN1,CCDC80,CCN1,CCN2,CD163,CD93,CNN1,COL15A1,COL16A1,COL1A1,COL1A2,COL3A1,COL4A1,COL4A2,COL5A1,COL5A2,COL6A2,COL6A3,CRISPLD2,DCN,DKK3,EMILIN1,ENG,FBLN2,FBLN5,FBN1,FN1,FOS,FSTL1,GGT5,HSPG2,ID3,IGFBP7,IL7R,INHBA,LHFPL6,LTBP2,LUM,MCAM,MFGE8,MMP2,MYL9,OLFML2B,PDGFRB,PECAM1,PPP1R14A,PRELP,PRSS23,SERPINE1,SLIT3,SPARC,SULF1,SYNPO2,TAGLN,THBS1,THBS2,TIMP2,TIMP3,TPM2,VCAN,VWF |
| Small Region | Positively Correlated Genes (late-enriched genes) | GigaPath | UNI2h | ViTB16 | RN18 |
| | Colon | 52<br>ACTG1,C19orf33,CBLC,CDC42EP5,CDH1,CDH17,CEACAM5,CEACAM6,CFL1,CLDN3,CLDN4,CLDN7,COX6B1,DDR1,EEF1B2,ELF3,EPCAM,ERBB3,FERMT1,FXYD3,GPX2,HMGN1,HNF4A,KLF5,KRT8,LGALS3BP,LGALS4,LRATD1,LSR,MUC13,MYH14,PHGR1,PLPP2,PPDPF,PPP1R1B,RAB25,RNF43,S100A10,S100A14,S100A6,SELENBP1,SLC12A2,SOX9,SPINT2,ST14,STARD10,TFF3,TMEM54,TMSB10,TPD52,TSPAN8,UBA52 | 61<br>ARL6IP1,ATP5F1B,C19orf33,CDH1,CDH17,CEACAM5,CEACAM6,CHCHD2,CLDN3,CLDN4,CLDN7,CLIC1,DDR1,EEF1B2,ELF3,EPCAM,ERBB3,FERMT1,FXYD3,GPX2,HDGF,HMGN1,HNF4A,HSPA8,INAVA,ITGB4,KLF5,KRT8,LGALS3,LGALS3BP,LGALS4,LRATD1,LSR,MAL2,MUC13,MYH14,PHGR1,PKM,PKP3,PLPP2,POF1B,PPDPF,PPP1R1B,PRR15L,RAB25,RNF43,S100A10,S100A14,S100P,SLC12A2,SOX9,SPINT2,ST14,STARD10,TAGLN2,TFF3,TMEM54,TMPRSS4,TMSB10,TSPAN8,USH1C | 33<br>ATP5IF1,CAPN1,CDC42EP5,CDCA7,CDH1,CEACAM5,CEACAM6,CLDN3,CLDN4,CLIC1,DDR1,EEF1B2,ELF3,EPCAM,FERMT1,GPX2,HDGF,HIST1H1C,HSPA5,KRT8,LGALS3BP,LGALS4,MGAT4B,MUC13,NME2,P4HB,RNF43,S100A10,S100A14,S100A6,TMSB10,TPD52,TSPAN8 | 41<br>ACTG1,BSG,CDC42EP5,CDH1,CEACAM5,CEACAM6,CLDN3,CLDN4,CLDN7,COX6B1,EEF1B2,EPCAM,ERBB3,FXYD3,GPX2,HNF4A,HSPA8,ITGB4,KRT8,LGALS3BP,LGALS4,LSR,MUC13,MYH14,NME2,NQO1,PKM,PKP3,PPDPF,PRR15L,RNF43,S100A14,SOX9,SPINT2,ST14,STARD10,TFF1,TFF3,TMPRSS4,UBA52,USH1C |
| | Prostate | 74<br>ABCC4,ACPP,ATF4,ATP1A1,ATP5MC2,CD9,CDH1,CIRBP,CREB3L4,DBI,DCXR,DDX5,DHRS7,EEF2,EPCAM,FAAH,FASN,FLNB,FOXA1,FTH1,H2AFJ,HINT1,HNRNPA2B1,HOXB13,HSP90AA1,HSP90B1,HSPA8,HSPE1,KDELR2,KLK2,KLK3,KLK4,KRT18,KRT8,MLPH,MPC2,NDRG1,NEDD4L,NKX3-1,NPY,ODC1,P4HB,PARK7,PLPP1,PMEPA1,PRAC1,PTPRF,RACK1,RDH11,RPL11,RPN2,SEC11C,SERF2,SERP1,SLC44A4,SLC45A3,SMS,SPDEF,SPINT2,SPOCK1,SPON2,STEAP1,STEAP2,SYNGR2,TACSTD2,TMEM258,TMPRSS2,TPD52,TRPM4,TSPAN1,TSTD1,UBA52,UQCRQ,XBP1 | 68<br>ABCC4,ACLY,ACPP,ACSL3,ARG2,ATP1A1,CANX,CD9,CDH1,CREB3L4,DBI,DCXR,DHRS7,EEF2,EIF4A1,EPCAM,FAAH,FASN,FLNB,FOXA1,GLO1,GOLPH3,H2AFJ,HINT1,HNRNPA2B1,HOXB13,HSP90B1,HSPE1,KLK2,KLK3,KLK4,KRT18,KRT8,MLPH,MPC2,NEDD4L,NKX3-1,P4HB,PABPC1,PARK7,PLPP1,PMEPA1,PRAC1,PTGR1,PTPRF,RAB11A,RACK1,RDH11,RPL11,RPN2,RPS24,SEC11C,SERF2,SLC45A3,SMS,SPOCK1,SRSF5,STEAP1,STEAP2,SYNGR2,TACSTD2,TMEM258,TMPRSS2,TPD52,TRPM4,TSPAN1,TSTD1,XBP1 | 46<br>ACPP,APLP2,ATP1A1,ATP2C1,CDH1,CIRBP,EPCAM,FAAH,FASN,FLNB,FOXA1,GOLPH3,H2AFJ,HOXB13,HSP90AA1,HSPA8,KLK2,KLK3,KRT18,MARCKSL1,MLPH,MPC2,NDRG1,NEDD4L,NKX3-1,P4HB,PMEPA1,PRAC1,PTPRF,RACK1,RDH11,SERF2,SERP1,SLC44A4,SLC45A3,SPOCK1,STEAP1,STEAP2,SYNGR2,TMPRSS2,TPD52,TRPM4,TSPAN1,UBA52,UQCRQ,XBP1 | 55<br>ACPP,ATP1A1,CD9,CDH1,CIRBP,CREB3L4,DBI,DCXR,DHCR24,DHRS7,FASN,FLNB,FOXA1,FTH1,H2AFJ,HNRNPA2B1,HOXB13,HSP90AA1,HSPA5,HSPE1,KDELR2,KLK2,KLK3,KLK4,KRT18,KRT8,MLPH,MPC2,NDRG1,NKX3-1,ODC1,P4HB,PLPP1,PMEPA1,PTGR1,PTPRF,RDH11,RPS24,SEC11C,SERP1,SLC39A6,SLC44A4,SLC45A3,SMS,SPDEF,SPINT2,STEAP1,STEAP2,TACSTD2,TMPRSS2,TPD52,TSPAN1,TSTD1,UBA52,XBP1 |
| | Breast | 40<br>ACTB,ACTG1,ASAH1,ATP5F1A,ATP6AP2,CALR,CD24,CD46,CHCHD2,DDX5,DUSP4,EIF4G2,ERBB2,ESR1,FASN,GNAS,H1-0,HNRNPA2B1,HSP90AB1,HSPA1A,HSPB1,KRT19,MORF4L2,PSMD3,PTMA,RP11-53O19.3,RPL23,RPL31,SARAF,SLC39A6,SLC7A2,SPINT2,SREBF1,SRRM2,TACSTD2,TAGLN2,TMSB10,VMP1,XBP1,YWHAZ | 31<br>ACTG1,ASAH1,ATP5F1A,ATP6AP2,CALR,CD24,CD46,DDX5,DUSP4,ERBB2,FASN,GATA3,GNAS,H1-0,HNRNPA2B1,HSP90AB1,HSPB1,KRT19,MORF4L2,PTMA,RPL31,SDC1,SLC39A6,SLC7A2,SPINT2,SRRM2,TAGLN2,TMSB10,VMP1,XBP1,YWHAZ | 24<br>ACTB,ACTG1,ASAH1,ATP6AP2,CALR,CD24,CD46,COX6C,DUSP4,EIF4G2,ERBB2,FASN,GATA3,HNRNPA2B1,HNRNPH1,KRT19,MORF4L2,NEAT1,PRDX1,PTMA,RPL31,SPINT2,SREBF1,XBP1 | 23<br>ACTB,ACTG1,CD24,CD46,CHCHD2,COX6C,DDX5,EIF4A1,ERBB2,ESR1,FASN,HNRNPA2B1,HSPA1A,MORF4L2,NEAT1,PLXNB2,PRDX1,PTMA,RP11-53O19.3,RPL23,TSPAN13,UBE2B,XBP1 |
| | Lymph | 40<br>AK4,ATP1B1,AZGP1,BEX3,CD9,CLU,CMBL,CP,CRYAB,CXCL14,CYB5A,DHCR24,ENO1,ERO1A,FABP7,FXYD2,KRT18,MGST1,MIF,MUC1,NDRG1,P4HB,PABPC1,PDE10A,PDZK1IP1,PGGHG,PLOD2,PPDPF,S100A10,S100A11,SCD,SERF2,SPINK13,SPP1,TMBIM6,TXN,UBC,UQCRQ,VEGFA,WFDC2 | 39<br>ASS1,ATP1B1,AZGP1,BEX3,CD24,CD9,CLU,CP,CSTB,CXCL14,CYB5A,DHCR24,ENO1,FABP7,FTH1,FXYD2,GPX3,KRT18,MGST1,MIF,MUC1,NDRG1,P4HB,PDE10A,PDZK1IP1,PPDPF,S100A10,S100A11,SCD,SDC4,SERF2,SERINC2,SPINK13,SPP1,TMBIM6,TXN,UBC,UQCRQ,VEGFA | 34<br>AK4,ANGPTL4,APOL1,ATP1B1,C3,CA12,CLU,CP,CRYAB,ENO1,ERO1A,FXYD2,GPX3,IGFBP3,KRT18,MGST1,MUC1,NDRG1,NNMT,NUPR1,P4HB,PDE10A,PDZK1IP1,PLOD2,S100A11,SCD,SERINC2,SERPINA1,SOD2,SPINK13,SPP1,TMBIM6,UBC,VEGFA | 18<br>APOL1,ATP1B1,C3,CALR,CLU,CP,ENO1,FABP7,FXYD2,KRT18,NNMT,P4HB,PDZK1IP1,PIGR,PPIB,S100A11,SOD2,WFDC2 |
| | Negatively Correlated Genes (early-enriched genes) | GigaPath | UNI2h | ViTB16 | RN18 |
| | Colon | 85<br>A2M,ACTA2,ACTG2,AEBP1,BGN,BOC,C11orf96,C1R,C1S,C3,CALD1,CAVIN1,CCDC80,CCN1,CHRDL2,CLU,CNN1,COL14A1,COL16A1,COL1A1,COL1A2,COL3A1,COL6A2,COL6A3,COL8A1,COMP,CRISPLD2,CRYAB,CXCL12,DCN,DES,DPYSL3,EFEMP1,ELN,EMILIN1,FBLN1,FBLN2,FBN1,FERMT2,FGF7,FLNA,FN1,FNDC1,FSTL1,FXYD6,GREM1,HSPB6,IGFBP5,IGFBP7,IGHG1,LIMS2,LMOD1,LUM,MFAP4,MGP,MMP2,MXRA8,MYH11,MYL9,MYLK,NNMT,PDLIM3,PLN,PODN,PRELP,PTGDS,PTGIS,SERPINF1,SERPING1,SFRP2,SFRP4,SPARC,SPARCL1,SSC5D,SULF1,SYNM,SYNPO2,TAGLN,THBS2,TIMP2,TNS1,TNXB,TPM2,VCAN,VIM | 84<br>A2M,ACTA2,ACTG2,ADAM33,AEBP1,BGN,BOC,C11orf96,C1R,C1S,C3,CALD1,CAVIN1,CCDC80,CCN1,CHRDL2,CLU,CNN1,COL14A1,COL16A1,COL1A1,COL1A2,COL3A1,COL6A2,COL6A3,COMP,CRISPLD2,CRYAB,CXCL12,DCN,DES,DPT,DPYSL3,EFEMP1,ELN,EMILIN1,FBLN1,FBLN2,FBN1,FERMT2,FGF7,FHL1,FLNA,FNDC1,FSTL1,FXYD6,GREM1,HSPB6,IGFBP5,IGFBP7,LIMS2,LMOD1,LUM,MFAP4,MGP,MMP2,MXRA8,MYH11,MYL9,MYLK,NNMT,PDLIM3,PLN,PODN,PRELP,PTGDS,PTGIS,SERPINF1,SERPING1,SFRP2,SFRP4,SPARC,SPARCL1,SSC5D,SULF1,SYNM,SYNPO2,TAGLN,THBS4,TIMP2,TNS1,TNXB,TPM2,VIM | 75<br>A2M,ACTA2,ACTG2,AEBP1,BGN,BOC,C11orf96,C1R,C1S,C3,CALD1,CAVIN1,CCDC80,CCN1,CHRDL2,CNN1,COL16A1,COL1A1,COL1A2,COL3A1,COL6A2,COL6A3,COL8A1,COMP,CRISPLD2,DCN,DES,DPYSL3,ELN,EMILIN1,FBLN1,FBLN2,FBN1,FERMT2,FGF7,FLNA,FN1,FSTL1,FXYD6,GREM1,IGFBP5,IGFBP7,ISLR,LIMS2,LMOD1,LUM,MFAP4,MGP,MMP2,MXRA8,MYH11,MYL9,MYLK,NNMT,PDLIM3,PLN,PODN,PRELP,PTGIS,SERPING1,SFRP2,SFRP4,SPARC,SPARCL1,SSC5D,SULF1,SYNM,SYNPO2,TAGLN,THBS2,TIMP2,TNS1,TNXB,TPM2,VIM | 58<br>A2M,ACTA2,ACTG2,AEBP1,BGN,BOC,C11orf96,C1R,C1S,C3,CALD1,CAVIN1,CCDC80,CCN1,CNN1,COL16A1,COL1A1,COL1A2,COL3A1,COL6A2,COL6A3,COL8A1,COMP,CRISPLD2,DCN,DPYSL3,EMILIN1,FBLN1,FBLN2,FBN1,FLNA,FN1,FSTL1,GREM1,IGFBP5,IGFBP7,ISLR,LIMS2,LUM,MMP2,MXRA8,MYH11,MYL9,MYLK,NNMT,PDLIM3,SERPING1,SPARC,SPARCL1,SULF1,TAGLN,THBS2,THY1,TIMP2,TNS1,TPM2,VCAN,VIM |
| | Prostate | 90<br>ACTA2,ACTC1,ACTG2,ADGRA2,AEBP1,AL078639.1,C1R,C1S,C2orf40,CALD1,CARMN,CAVIN1,CCDC80,CHRDL1,CLU,CNN1,COL16A1,COL1A2,COL4A2,COL6A1,COL6A2,COL6A3,COX7A1,CRYAB,CSRP1,DCN,DES,EMILIN1,FBLN1,FHL1,FILIP1L,FLNA,FLNC,FXYD1,FXYD6,GSN,HSPB6,HSPB8,IGFBP6,INMT,ITGA5,JPH2,KCNMB1,LGALS1,LIMS2,LMOD1,MAP1B,MASP1,MFGE8,MGP,MRGPRF,MRVI1,MSRB3,MXRA7,MYH11,MYL9,MYLK,NEXN,OGN,PAGE4,PALLD,PCP4,PDE5A,PDLIM7,PGM5,PGM5-AS1,PLN,PPP1R14A,PRELP,PTGDS,PTGIS,RARRES2,RASL12,RBPMS,S100A6,SELENOM,SERPINF1,SERPING1,SMOC1,SMTN,SORBS1,SPARCL1,SYNM,SYNPO2,TAGLN,TGFB1I1,TNS1,TPM1,TPM2,WFDC1 | 91<br>ACTA2,ACTC1,ACTG2,ADGRA2,AEBP1,AL078639.1,C1R,C1S,C2orf40,CALD1,CARMN,CAVIN1,CCDC80,CHRDL1,CLU,CNN1,COL16A1,COL1A2,COL4A2,COL6A1,COL6A2,COL6A3,COX7A1,CSRP1,DCN,DES,EMILIN1,FBLN1,FHL1,FILIP1L,FLNA,FLNC,FXYD1,FXYD6,GSN,HSPB6,HSPB8,IGFBP6,INMT,ITGA5,JPH2,KCNMB1,LGALS1,LIMS2,LMOD1,MAP1B,MASP1,MFAP4,MFGE8,MGP,MRGPRF,MRVI1,MSRB3,MXRA7,MYH11,MYL9,MYLK,NEXN,OGN,PAGE4,PALLD,PCP4,PDE5A,PDLIM3,PDLIM7,PGM5,PGM5-AS1,PLN,PPP1R14A,PRELP,PTGDS,PTGIS,RARRES2,RASL12,RBPMS,S100A6,SELENOM,SERPINF1,SERPING1,SMOC1,SMTN,SORBS1,SPARCL1,SYNM,SYNPO2,TAGLN,TGFB1I1,TNS1,TPM1,TPM2,WFDC1 | 83<br>ACTA2,ACTG2,ADGRA2,AEBP1,AL078639.1,AOC3,C1R,C1S,C2orf40,CALD1,CARMN,CAVIN1,CCDC80,CHRDL1,CLU,CNN1,COL16A1,COL1A2,COL4A2,COL6A1,COL6A2,COL6A3,COX7A1,CSRP1,DCN,DES,EMILIN1,FBLN1,FHL1,FILIP1L,FLNA,FLNC,FXYD1,GSN,HSPB6,HSPB8,IGFBP6,INMT,ITGA5,JPH2,KCNMB1,LGALS1,LIMS2,LMOD1,MAP1B,MFGE8,MGP,MRGPRF,MRVI1,MSRB3,MXRA7,MYH11,MYL9,MYLK,NEXN,OGN,PAGE4,PALLD,PCP4,PDLIM7,PGM5,PLN,PPP1R14A,PRELP,PTGDS,PTGIS,RARRES2,RBPMS,S100A6,SELENOM,SERPINF1,SERPING1,SMTN,SORBS1,SPARCL1,SYNM,SYNPO2,TAGLN,TGFB1I1,TNS1,TPM1,TPM2,WFDC1 | 89<br>ACTA2,ACTC1,ACTG2,ADGRA2,AEBP1,AL078639.1,AOC3,C1R,C2orf40,CALD1,CARMN,CAVIN1,CCDC80,CHRDL1,CLU,CNN1,COL16A1,COL1A2,COL4A2,COL6A1,COL6A2,COL6A3,COX7A1,CSRP1,DCN,DES,EMILIN1,FBLN1,FHL1,FLNA,FLNC,FXYD1,FXYD6,GSN,HSPB6,HSPB8,IGFBP6,INMT,ITGA5,JPH2,KCNMB1,LAMA4,LGALS1,LIMS2,LMOD1,MAP1B,MFGE8,MGP,MRGPRF,MRVI1,MSRB3,MXRA7,MYH11,MYL9,MYLK,NEXN,OGN,PAGE4,PALLD,PCP4,PDE5A,PDLIM7,PGM5,PGM5-AS1,PLN,PPP1R14A,PRELP,PTGDS,PTGIS,RARRES2,RASL12,RBPMS,S100A6,SELENOM,SERPINF1,SERPING1,SGCA,SMOC1,SMTN,SORBS1,SPARCL1,SYNM,SYNPO2,TAGLN,TGFB1I1,TNS1,TPM1,TPM2,WFDC1 |
| | Breast | 76<br>AEBP1,APOC1,APOE,BGN,C1QA,C1QB,C1QC,C1R,C3,CAVIN1,CCL19,CCL5,CD14,CD2,CD27,CD52,CD6,CD68,CD74,CD79A,CLDN5,COL6A2,COMP,CORO1A,CSF1R,CTSD,CYBA,DCN,DERL3,EMILIN1,EMP3,ENG,GPSM3,GSTP1,HCST,HLA-DMA,HLA-DPA1,HLA-DPB1,HLA-DQB1,HLA-DRB1,HLA-E,IGFBP4,IGLL1,IGLL5,IL32,JAK3,JCHAIN,LAPTM5,LGALS1,LSP1,LTB,MMP2,MMP9,MYL9,MZB1,NNMT,PCOLCE,PLEKHO1,POU2AF1,PTGDS,RARRES2,RPL18A,RPLP1,S100A4,SERPINF1,SSR4,STAB1,TAGLN,TBC1D10C,TGFB1,THY1,TIMP1,TMEM119,TPSB2,VIM,VWF | 74<br>AEBP1,APOC1,APOE,BGN,C1QA,C1QB,C1QC,C1R,C3,CAVIN1,CCL19,CCL21,CCL5,CD14,CD2,CD27,CD52,CD6,CD68,CD74,CD79A,CLDN5,COL6A2,COMP,CORO1A,CSF1R,CTSD,CYBA,DCN,EMILIN1,EMP3,ENG,GPSM3,GSTP1,HCST,HLA-DMA,HLA-DPA1,HLA-DPB1,HLA-DQB1,HLA-DRB1,HLA-E,IGFBP4,IGLL1,IGLL5,IL32,JAK3,JCHAIN,LAPTM5,LGALS1,LSP1,LTB,MMP9,MYL9,MZB1,PCOLCE,PI16,PLEKHO1,PTGDS,RARRES2,RPL18A,RPLP1,S100A4,SELENOM,SERPINF1,SSR4,STAB1,TAGLN,TGFB1,THY1,TIMP1,TMEM119,TPSB2,VIM,VWF | 51<br>AEBP1,APOC1,APOE,BGN,C1QA,C1QB,C3,CAVIN1,CCL19,CCL5,CD14,CD2,CD27,CD52,CD74,CD79A,CLDN5,COL6A2,CSF1R,CTSD,CYBA,ENG,GSTP1,HLA-DMA,HLA-DPA1,HLA-E,IGFBP4,IGLL1,IGLL5,IL32,JAK3,JCHAIN,LAPTM5,LGALS1,LSP1,LTB,MMP9,MYL9,MYO1F,MZB1,PTGDS,RARRES2,RPL18A,RPLP1,SERPINF1,SSR4,TAGLN,TIMP1,TMEM119,VIM,VWF | 29<br>AEBP1,APOC1,APOE,BGN,C1QA,CAVIN1,CCL19,CCL5,CD2,CD79A,CTSD,ENG,GSTP1,HLA-DPA1,IGFBP4,IGLL1,IGLL5,IL32,JAK3,JCHAIN,LAPTM5,LSP1,MMP9,MZB1,NKG7,POU2AF1,SSR4,TAGLN,VWF |
| | Lymph | 67<br>A2M,ACTA2,ADAMTS2,AEBP1,BGN,C7,CAVIN1,CCDC80,CCN1,CCN2,CD163,CD93,CDH11,COL15A1,COL1A1,COL1A2,COL3A1,COL4A1,COL4A2,COL5A1,COL5A2,COL6A2,COL6A3,CRISPLD2,DCN,EFEMP1,ELN,EMILIN1,ENG,FBLN1,FBLN2,FBLN5,FBN1,FN1,FOS,FSTL1,GGT5,HSPG2,ID3,IGFBP7,IGKC,IL7R,LHFPL6,LTBP2,LUM,MCAM,MFGE8,MGP,MMP2,MYL9,OLFML2B,PECAM1,PPP1R14A,PRELP,PRSS23,PTPRC,SERPINE1,SERPINF1,SLIT3,SPARC,TAGLN,THBS1,THBS2,TIMP2,TPM2,VCAN,VWF | 64<br>A2M,ACTA2,ADAMTS2,AEBP1,BGN,C7,CAVIN1,CCDC80,CCN1,CCN2,CD163,CD93,COL15A1,COL16A1,COL1A1,COL1A2,COL3A1,COL4A1,COL4A2,COL5A1,COL6A2,COL6A3,DCN,EFEMP1,ELN,EMILIN1,ENG,FBLN1,FBLN2,FBLN5,FBN1,FN1,FOS,FOSB,FSTL1,GGT5,HSPG2,ID3,IGFBP7,IGKC,IL7R,LHFPL6,LTBP2,LUM,MCAM,MFGE8,MMP2,MYL9,OLFML2B,PECAM1,PPP1R14A,PRELP,PRSS23,PTPRC,SERPINE1,SERPINF1,SLIT3,SPARC,TAGLN,THBS2,TIMP2,TPM2,VCAN,VWF | 61<br>A2M,ACTA2,ADAMTS2,AEBP1,ANTXR1,BGN,CAVIN1,CCDC3,CCDC80,CCN1,CCN2,CD93,CNN1,COL16A1,COL1A1,COL1A2,COL3A1,COL4A1,COL4A2,COL5A1,COL5A2,COL6A2,COL6A3,CRISPLD2,DCN,DKK3,EMILIN1,ENG,FBLN5,FBN1,FN1,FSTL1,GGT5,HSPG2,ID3,IGFBP7,IL7R,INHBA,LHFPL6,LUM,MCAM,MFAP4,MFGE8,MMP2,MYL9,OLFML2B,PDGFRB,PDLIM3,PPP1R14A,PRELP,PRSS23,SERPINE1,SLIT3,SPARC,SULF1,SYNPO2,TAGLN,THBS2,TIMP2,TIMP3,TPM2 | 27<br>A2M,ACTA2,AEBP1,BGN,CAVIN1,CCN1,CCN2,CD93,CNN1,COL1A2,COL3A1,COL4A1,FN1,FSTL1,IGFBP7,LHFPL6,MCAM,MYL9,PDGFRB,PPP1R14A,PRSS23,SERPINE1,SPARC,SULF1,TAGLN,THBS2,TPM2 |

### Table.S7

| Region | Gene set | Model | Genes |
|---|---|---|---|
| Large Region | Positively Correlated Genes (late- enriched genes) | CONCH | AC007272.1,AC008937.3,AC026471.5,AC092490.1,AC093843.2,AC104035.1,AC127024.5,AC130650.2,AD000090.1,AL022323.4,AL121970.1,AL158835.1,AL356259.1,AL449363.2,AL596325.2,AP000708.1,AP000873.4,APOD,BEND2,BOK,CCDC80,CDH3,CEMIP,CNKSR1,COL10A1,COL11A1,COL12A1,COL14A1,COL8A1,COMP,CRISPLD2,DCANP1,DCN,DHCR24-DT,DKK2,EFNB3,FAP,FGF7,FNDC1,GASAL1,GNAL,HECW1,HPD,HSD17B6,IGF1,IGHG1,IGHG2,IGHG3,IGHG4,IGHGP,IGKC,IGKV1-5,IGKV3-15,IGLC2,IGLC3,INHBA,ISM1,ITGA11,ITGBL1,KIAA1841,KLHL35,LIMD1-AS1,LUM,LYRM7,MFAP5,MMP1,MYH10,NGEF,OMD,OR11G2,OR51D1,P2RY10,PCDHA7,PDE6A,PLN,PNOC,PRELP,PTGIS,RIC3,RPS6KA6,RSPO3,SFRP4,SHISA3,SLC28A2,SLCO2A1,SPOCD1,SPON1,SSTR5-AS1,SULF1,TEKT4,TIMP3,TMEM163,TMEM178A,TRBV28,TSACC,WT1-AS,Z98257.1,ZFPM2,ZNF329,ZNF682 |
| | | CONCH15 | AC008915.2,AC093843.2,AC127024.5,ACTA2,AD000090.1,AEBP1,AFAP1,AL049838.1,ANTXR1,APOD,ARL17A,BGN,CCDC80,CCN1,CCN2,CD52,CEMIP,COL10A1,COL11A1,COL12A1,COL14A1,COL1A1,COL1A2,COL3A1,COL5A1,COL8A1,COMP,CPXM2,CRISPLD2,DCN,DKK2,EFEMP1,ELN,EMILIN1,FAP,FGF7,FNDC1,GJA5,GREM1,HDGFL3,IGF1,IGHG1,IGHG2,IGHG3,IGHG4,IGHGP,IGKC,IGLC2,IGLC3,IL7R,INHBA,ISLR,ISM1,ITGA11,ITGBL1,KLRB1,LIMD1-AS1,LRRC17,LTBP2,LUM,MFAP4,MFAP5,MGP,MMP2,MRGPRF,MRVI1,MXRA5,MYH10,MZB1,NEXN,NUDT10,OMD,P2RY10,PALLD,PCDHA7,PDE6A,PNOC,POSTN,PPP1R14A,PRELP,PTGDS,PTGIS,RAD51B,RAMP1,RFLNA,SFRP2,SFRP4,SHISA3,SPON1,SULF1,TAGLN,THBS1,THBS2,TIMP3,TNN,TPM2,UCHL1,WT1-AS,ZNF582-AS1,ZNF682 |
| | | CTransPath | AC005041.4,AC005072.1,AC007272.1,AC009955.4,AC010226.1,AC016907.2,AC025271.4,AC080005.1,AC087071.1,AC093426.1,AC093849.4,AC096733.2,AC099066.2,AC211486.2,AC245052.4,AD000090.1,ADGRB2,AL022323.4,AL031963.3,AL034550.2,AL121970.1,AL136962.1,AL137002.1,AL158835.1,AL356259.1,AL391684.1,AL596325.2,ALDH1L2,AP000873.4,ATP1A2,BEND2,BTBD11,CCDC26,CCDC80,CEND1,CNKSR1,COL11A1,COL14A1,COMP,DCANP1,DCN,DHCR24-DT,EFNB3,EREG,FGF7,FNDC1,GASAL1,GLI2,GOLGA6L7,HOXC-AS1,HPD,HSD17B6,IGF1,IGHV4-34,IGKV1-5,IGKV3-15,INHBA,ISM1,ITGA11,ITGAD,KCNQ4,KIF26B,LDLRAD2,LGI1,LINC01048,LINC01500,LINC02037,LINC02432,LUM,LYRM7,MFAP5,MROH7,MYH10,NGEF,NLGN2,NPIPB3,OMD,PDE6A,PTGIS,RIC3,RPS6KA6,RSPO3,RUNX2,SFRP4,SH2D4A,SHISA3,SLC28A2,SPOCD1,SPON1,SULF1,TCF23,TEKT4,TIMP3,WFDC1,WT1-AS,Z98257.1,ZNF561,ZNF682,ZNF749,ZNF823 |
| | | GigaPath | ABHD4,AC006001.2,AC008875.3,AC092171.4,AC093843.2,AC093849.4,AC096733.2,AC097662.1,AC100810.1,ACKR4,AKAP11,AL136962.1,AL645933.2,ALDH1L2,ALOX15B,ANOS1,APOD,ARF4,ARHGEF26,ARL11,ARL17A,ATP1A2,BACE1,BCAN,C12orf29,CALM1,CCDC80,CD209,CD36,CEP63,CHCHD6,CHIT1,COL3A1,COLEC12,CRYBG2,CSF2RA,DCAF4,DCN,DCTN3,DPY30,DYNC1I1,EPHB2,FAM111A-DT,FCGR2A,FILIP1L,FNDC1,FOXD2,HIST1H2BN,HSCB,INHBA,ISM1,ITGAD,ITGB1BP2,KIAA1586,KIF26B,KREMEN1,LINC02432,LIPA,ME1,MIR4458HG,NAV2,NCOA3,NGEF,NLGN2,P2RX7,PABPC1,PDE6A,PIP4K2B,PLA2G7,PLAC9,PPM1M,PRMT9,PRR12,PRSS35,PTGIS,RASGRF2,RIC3,RP1,SLC16A12-AS1,SLC20A2,SLC46A3,SMIM30,SPOCD1,SRPRA,STXBP5,SULF1,TCEAL3,TMEM154,TMEM248,TMEM9B-AS1,TRDMT1,TRNT1,UBE2Q2,UTP11,YBX1,YIPF5,ZMYM6,ZNF134,ZNF44,ZNF726 |
| | | RN18 | AARD,AC004771.4,AC044839.1,ACTA2,AD000090.1,AEBP1,AP000873.4,APOD,ARHGEF26,CCDC80,CCN1,CCN2,CCN4,CDH3,CFAP206,CLCNKB,COL10A1,COL11A1,COL12A1,COL14A1,COL16A1,COL1A1,COL1A2,COL3A1,COL5A1,COL5A2,COL8A1,COMP,CRISPLD2,CSRP1,CSTF2T,CTSK,DCN,DKK2,ELN,F3,FBN1,FNDC1,FZD3,GGTLC2,HMBOX1,IGF1,IGHG1,IGHG3,IGHG4,IGHGP,IGKC,IGLC2,IGLC3,IL7R,INHBA,ISLR,ISM1,ITGBL1,KCNMB1,KCNMB3,KL,LIMD1-AS1,LMTK3,LTBP2,LUM,LYRM7,MATN3,MFAP4,MFAP5,MGP,MMP2,MT1E,MT1M,MXRA8,MYH10,NEXN,NGEF,NLGN2,OMD,OR1I1,PCBP4,PLA2R1,PLXNA4,PRELP,PRRX1,PTGDS,PTGFR,PTGIS,RNASEH2C,RPL39L,SCUBE2,SFRP2,SFRP4,SHISA3,SMOC1,SPON1,SULF1,TAGLN,THBS2,TIMP3,TPM2,UCHL1,ZNF641,ZNF682 |
| | | UNI | ABHD4,AC002059.1,AC005072.1,AC006001.2,AC007272.1,AC008759.3,AC013270.1,AC020661.1,AC026979.4,AC073225.1,AC080005.1,AC087071.1,AC093388.1,AC093426.1,AC093849.4,AC114271.1,AC138781.1,ADPGK-AS1,AJAP1,AL008729.1,AL031963.3,AL121772.1,AL136962.1,AL137002.1,AL353600.2,AL390195.2,ANGPTL6,AP000873.2,AQP7,ARL11,ASAP1-IT2,ATP1A2,BCAN,BCL2L15,BMP8B,BUB1B,C12orf29,CCNB2,CD36,CEND1,CRYBG2,CTAGE1,DNAH6,DPY30,FAM90A26,FANCI,FBXO27,FILNC1,GASAL1,GPRIN1,GSTA1,HILPDA,HIST1H1C,HOXC-AS1,HP,HSCB,IGHV4-34,IMMP2L,ITGAD,KCND3,KCNQ4,KEL,KIAA1586,LCTL,LGI1,LINC00173,LINC02256,LINC02432,LINC02482,LRRC4C,LRRD1,MAGEL2,MELTF-AS1,MIR4458HG,MT-ATP8,MYRF,NDRG1,NPIPB3,PAK4,PART1,PIFO,PPP4R4,RAB39B,RASL11A,RIC3,RMST,RP1,RSPO3,SAA1,SERHL2,SLC25A23,SNAPC1,TPBGL,TPRN,TRAIP,TRNT1,TTYH1,ZMYM6,ZNF682,ZNF726 |
| | | UNI2h | AC000403.1,AC007272.1,AC007448.5,AC007952.3,AC020661.1,AC044839.1,AC080005.1,AC087878.1,AC090192.2,AC093426.1,AC093849.4,AC096733.2,AC099066.2,AC099796.2,AC138781.1,AD000090.1,ADGRB2,AL008729.2,AL035587.3,AL137002.1,AL139393.3,AL161782.1,AL356417.2,AL391684.1,ALDH1L2,AP000873.4,ARHGEF26,ATP1A2,BMP8B,C5orf64,CBWD2,CCDC26,CCDC80,CDH24,CEND1,CNKSR1,COL12A1,COMP,CSPG5,CTAGE1,DCN,DKK2,DUSP26,DYNC1I1,EFNB3,F8,GASAL1,GLI2,GPR171,GREM2,HOXA10-AS,HOXC-AS1,HPD,HSD17B6,IGF1,IGHV1-46,IGKV2-30,IGLV3-1,ISM1,ITGAD,ITK,KCNJ14,KCNQ4,KIF26B,LCTL,LGI1,LIMD1-AS1,LINC01048,LINC01277,LINC02432,LRRD1,MAGEL2,MFAP5,MROH7,MYH10,NGEF,NLGN2,OGN,OMD,PCBP4,PDE6A,PRSS35,PTGIS,RSPO3,SFRP4,SFTPD-AS1,SNAI3-AS1,SOX2-OT,SULF1,TEX11,TMEM9B-AS1,TNN,TNNC2,TRAV23DV6,TRDMT1,WFDC1,WT1-AS,Z98257.1,ZNF749,ZNF823 |
| | | Virchow | ABCB5,AC005072.1,AC006001.2,AC007272.1,AC020661.1,AC039056.2,AC044839.1,AC053527.2,AC080005.1,AC087071.1,AC090192.2,AC092171.4,AC092445.1,AC093426.1,AC093843.2,AC104035.1,AC104561.4,AC105137.2,AC113383.1,ACKR4,AD000090.1,AJAP1,AL031963.3,AL136962.1,AL137002.1,AL161782.1,AL662884.1,AP000708.1,AP000873.2,APOD,ARHGEF26,ARL17A,ATP1A2,BMX,C5orf64,CA6,CCDC26,CCDC80,CCNA1,CEND1,CLCN1,CMTM2,COL14A1,COMP,DKK2,DNAH6,DYNC1I1,FAM43B,FBXL2,FNDC1,GASAL1,GNAL,GPR78,HCG9,HOXC-AS1,HPD,HSD17B6,IGF1,IGHV1-24,IGHV4-34,IGKV1-5,INHBA,ISM1,ITGBL1,ITK,KCNQ4,KCNRG,KHDRBS2,KIF26B,LGI1,LINC02432,LMCD1-AS1,LRRD1,MFAP5,NGEF,OMD,OR1I1,PDE6A,PLBD1-AS1,PLEKHD1,PPY,PRDM5,PRSS35,PTGIS,RASL11B,RIC3,RSPO3,SFTPD-AS1,SLC16A12-AS1,SLC17A4,SNAI3-AS1,SPON1,SULF1,TBX1,TDRD9,TMEM9B-AS1,TPH1,VIPR2,WT1-AS,XKR3 |
| | | Virchow2 | AC004865.2,AC005041.4,AC025280.1,AC044839.1,AC093426.1,AC104035.1,AC104561.4,ACKR4,ACTA2,AD000090.1,ADGRD1,AL022323.4,AL137002.1,AL158835.1,AL513548.4,ALDH1L2,ANTXR1,AP000708.1,AP000873.2,APOD,ARHGEF26,ARL17A,ATP1A2,CCDC80,CCN1,CCNA1,CDH3,CLCN1,COL10A1,COL11A1,COL12A1,COL14A1,COL1A1,COL1A2,COL3A1,COL8A1,COMP,CPXM2,CRISPLD2,DCN,DKK2,FAM180A,FAP,FGF7,FNDC1,GNAL,GSPT2,HOXA5,HTR2B,IGF1,IGHG1,IGHG3,IGHG4,IGHV1-24,IGKC,IGLC2,INHBA,ISM1,ITGAD,ITGBL1,KCNQ4,KIF26B,LIMD1-AS1,LINC01424,LRRD1,LUM,MFAP5,MYH10,NEXN,NGEF,NLGN2,NUDT10,OMD,OR1I1,P2RY10,PCBP4,PDE6A,PLN,PPP1R14A,PRELP,PRSS35,PTGFR,PTGIS,RIC3,RSPO3,SFRP4,SHISA3,SIX4,SPON1,SULF1,TAGLN,TIMP3,TPM2,UCHL1,ULBP2,VGLL3,WT1-AS,ZFHX4,ZFPM2,ZNF682 |
| | | ViTB16 | ACTA2,AEBP1,ANKH,ANTXR1,ANXA2,B2M,BCAT1,BGN,CALD1,CAPG,CCDC80,CCL18,CCN1,CCN2,CCN4,CD14,CD84,CHCHD6,CHIT1,COL11A1,COL12A1,COL14A1,COL1A1,COL1A2,COL3A1,COL5A1,COL5A2,COL6A2,COL6A3,COL8A1,COMP,CRISPLD2,CSRP1,CTSB,DCN,EFEMP1,EMILIN1,FABP5,FCGR2A,FCGR3A,FN1,FNDC1,FSTL1,FTL,GPNMB,HLA-DMB,HLA-DPB1,HLA-DQA1,HLA-DRB1,IFI27,IFI30,IGF1,INHBA,LGMN,LIPA,LTBP2,LUM,LYZ,MFAP5,MGP,MMP2,MS4A4A,MSR1,MT1E,MT2A,MXRA5,MXRA8,MYL9,OLFML2B,PALLD,POSTN,PRELP,PRSS23,PTGIS,RPL36A,RPL36AL,RPS8,S100A6,SELENOP,SERPINF1,SFRP2,SFRP4,SPARC,SULF1,TAGLN,TALDO1,THBS1,THBS2,TIMP1,TIMP2,TIMP3,TLR4,TMSB4X,TNC,TPM2,TREM2,TYROBP,VCAN,VIM,ZYX |
| | Negatively Correlated Genes (early- enriched genes) | CONCH | ACTB,AKR1C3,ANGPTL4,ATP1B1,B2M,BIRC3,C1QA,C1S,C3,CA12,CALR,CD24,CD74,CFL1,CLU,CNDP2,CP,CRYAB,CSTB,CTSD,DDIT4,EEF1A1,EEF1G,EEF2,ENO1,FGB,FGG,FTH1,FTL,FXYD2,GAPDH,GPX3,GSDMD,HLA-A,HLA-B,HLA-E,HP,HSP90B1,HSPB1,IGFBP3,IGFBP4,ITM2B,LBP,LDHA,MALAT1,MAP7D2,MGST1,MIF,MT-ATP6,MT-CO1,MT-CO2,MT-CO3,MT-CYB,MT-ND2,MT-ND3,MT-ND4,MT-ND4L,MT-ND5,MYOCOS,NDRG1,NDUFA4L2,NEAT1,NNMT,P4HB,PABPC1,PDIA3,PDK4,PDZK1IP1,PFN1,PGK1,PKM,RPL13,RPL19,RPL21,RPL39,RPL4,RPL41,RPL7,RPL8,RPLP0,RPLP1,RPS17,RPS19,RPS2,RPS20,RPS21,S100A10,S100A11,SAT1,SERPINA1,SLC16A3,SOD2,SPP1,TAGLN2,TMBIM6,TMSB10,TPI1,TPT1,TRAM1,UBA52 |
| | | CONCH15 | AC090578.1,ADGRG1,AKR1C3,ATP1B1,BIRC3,C1S,C3,CA12,CALR,CANX,CD24,CERNA2,CERS2,CKB,CLU,CNDP2,COX6A1,COX6C,CP,CRYAB,CSTB,CXCL14,CYB5R3,CYC1,DDIT4,DEGS1,EFNA1,FGA,FGB,FGG,FXYD2,GAPDH,GPX3,GSDMD,GUK1,HINT1,HLA-A,HP,HSP90B1,HSPB1,IGFBP3,IGFBP4,ITIH5,ITM2B,KRT18,KRT8,LBP,LDHA,LOX,MALAT1,MAP7D2,MGST1,MIF,MLEC,MT-ATP6,MT-ATP8,MT-CO1,MT-CO2,MT-CO3,MT-CYB,MT-ND2,MT-ND3,MT-ND4,MT-ND4L,MT-ND5,MUC1,MYOCOS,NDRG1,NDUFA4L2,NEAT1,NNMT,NUPR1,P4HB,PABPC1,PDK4,PDZK1IP1,PGK1,PKM,PLOD2,RNASET2,RPL23A,RPL39,RPL4,RPL8,RPLP0,RPLP1,RPN2,RPS20,S100A10,SAA1,SERPINA1,SERPINF2,SLC16A3,SNHG25,SOD2,SPON2,SPP1,TMBIM6,TPI1,TRAM1 |
| | | CTransPath | ADGRG1,AKR1C3,ATP1B1,BIRC3,C1R,C1S,C3,CALR,CANX,CD24,CD74,CKB,CLU,CNDP2,CP,CRYAB,DDIT4,EEF1A1,EEF1G,EEF2,ENO1,FGB,FGG,FXYD2,GAPDH,GPX3,GSDMD,HLA-A,HLA-B,HP,HSP90B1,HSPB1,IGFBP3,IGFBP4,IGFBP5,ITM2B,LBP,LDHA,MALAT1,MAP7D2,MCL1,MIF,MT-ATP6,MT-CO1,MT-CO2,MT-CO3,MT-CYB,MT-ND2,MT-ND3,MT-ND4,MT-ND4L,MT-ND5,MUC1,MYOCOS,NDRG1,NDUFA4L2,NEAT1,NNMT,P4HB,PABPC1,PDK4,PDZK1IP1,PFN1,PGK1,PKM,RPL13,RPL13A,RPL19,RPL21,RPL28,RPL3,RPL4,RPL41,RPL7,RPL9,RPLP0,RPLP1,RPLP2,RPN2,RPS17,RPS19,RPS2,RPS20,RPS21,RPS27,RPS28,SEC61A1,SERPINA1,SERPINF2,SERPING1,SLC16A3,SNHG25,SOD2,SPP1,TAGLN2,TAPBP,TMSB10,TPI1,TRAM1,UBA52 |
| | | GigaPath | AC004687.1,ADA,ADAMTS1,AL627171.2,APOL1,ARHGDIA,ATP2A3,BTG2,C1QA,C1orf122,C3AR1,CCND3,CCR7,CD27,CD79A,CKB,CRIP1,CSK,CST3,CXCL9,DDX39A,DNAJB1,DNAJC13,EEF1G,EEF2,ENAH,EPAS1,EPS8L2,ERGIC1,ERO1B,FBLN1,FJX1,FUS,GIGYF1,GSDMD,H19,HBA2,HDAC10,HLA-DRA,HNRNPK,IER2,IGHA1,IGHG1,IGHG2,IGHG3,IGHG4,IGHGP,IGHM,IGKC,IGLC1,IGLC2,ILF3,INPP5D,ISG15,JCHAIN,JUN,LFNG,LINC02422,LTBP3,MBD3,MCL1,MDC1,MLLT1,MT-ND1,MT-ND4,MYCT1,MYL9,MZB1,NDUFV1,NLRC5,NR4A1,NUDCD3,PABPN1,PML,POU2AF1,PSMB9,RAD1,RBM6,RFNG,RGS5,RHOB,RPS5,SEC23B,SEC61A1,SPIB,SRRM2,ST6GAL1,STAT2,TAGLN2,TAPBP,TBC1D10C,TMEM94,TNRC18,TRABD,TRIM52,TSC2,TYMP,UBA52,VWF,WARS |
| | | RN18 | AC090578.1,ADGRG1,AKR1C3,ALPL,ATP1B1,BIRC3,C1S,C3,CALR,CANX,CD24,CERNA2,CKB,CLU,CNDP2,COX6A1,CP,CRYAB,CSTB,CXCL14,CYC1,DDIT4,DEGS1,FGB,FGG,FXYD2,GAPDH,GPX3,GSDMD,HLA-A,HLA-B,HP,HSP90B1,HSPB1,IGFBP3,IGFBP4,ITIH5,ITM2B,KCNJ16,KRT18,LBP,LDHA,LRP2,MALAT1,MAP7D2,MGST1,MIF,MLEC,MT-ATP6,MT-ATP8,MT-CO1,MT-CO2,MT-CO3,MT-CYB,MT-ND2,MT-ND3,MT-ND4,MT-ND4L,MT-ND5,MUC1,MYOCOS,NDRG1,NDUFA4L2,NEAT1,NNMT,NUPR1,P4HB,PABPC1,PDK4,PDZK1IP1,PGK1,PKM,PLOD2,RNASET2,RPL19,RPL23,RPL35A,RPL39,RPL4,RPL7,RPLP0,RPLP1,RPN2,RPS17,RPS20,RPS3,S100A10,SAA1,SERPINA1,SERPINF2,SLC16A3,SNHG25,SOD2,SPP1,TMBIM6,TMEM176B,TMSB10,TPI1,TPT1,YBX3 |
| | | UNI | A2M,ACTA2,ACTB,ACTG1,ACTN4,AEBP1,ANXA2,ARHGDIA,B2M,BGN,C1QA,C1QB,CALD1,CCN2,CD74,COL1A1,COL1A2,COL3A1,COL4A2,COL6A2,COMP,CST3,CTSB,CXCL9,DUSP1,EEF1A1,EEF2,FLNA,FOS,FTL,HLA-B,HLA-C,HLA-DPA1,HLA-DPB1,HLA-DQB1,HLA-DRA,HSPA8,IGHA1,IGHG1,IGHG2,IGHG3,IGHG4,IGHGP,IGHM,IGKC,IGLC1,IGLC2,IGLC3,JCHAIN,KLF2,LITAF,LTBP2,LUM,MGP,MLLT1,MMP2,MYL9,NR4A1,PFN1,RGS1,RPL10,RPL10A,RPL13,RPL13A,RPL18A,RPL19,RPL21,RPL26,RPL27A,RPL28,RPL32,RPL34,RPL35,RPL41,RPL9,RPS11,RPS12,RPS13,RPS15A,RPS27,RPS27A,RPS28,RPS29,RPS5,S100A4,SERPINF1,SFRP2,SPARC,STAT2,TAGLN,THBS1,THY1,TIMP1,TIMP2,TMSB10,TMSB4X,TRBC2,UBA52,VWF,ZFP36L2 |
| | | UNI2h | ACTB,ACTG1,ARHGDIA,ATP5F1B,ATP6V0C,B2M,BIRC3,BTG2,C1QA,C1QB,C1S,C3,CD24,CD74,CKB,CNDP2,CP,CRYAB,CST3,CXCL9,DDIT4,DUSP1,EEF1A1,EEF1D,EEF1G,EEF2,ENO1,EPAS1,FOS,FSTL3,FUS,GPX3,GSDMD,HBA2,HLA-A,HLA-B,HLA-C,HLA-DRA,HLA-E,HNRNPK,HSP90B1,IGFBP5,ILF3,LDHA,MALAT1,MCL1,MGST1,MIF,MT-ATP6,MT-CO1,MT-CO2,MT-CO3,MT-CYB,MT-ND1,MT-ND2,MT-ND3,MT-ND4,MT-ND4L,NNMT,P4HB,PDZK1IP1,PFN1,PGK1,PKM,PLOD2,RACK1,RHOB,RPL13,RPL13A,RPL18A,RPL19,RPL21,RPL27A,RPL28,RPL3,RPL36,RPL41,RPL7,RPL9,RPN2,RPS11,RPS19,RPS2,RPS27,RPS28,RPS29,RPS5,SCPEP1,SEC61A1,SERPINA1,SERPING1,SLC16A3,SOD2,SULF2,TAGLN2,TAPBP,TMSB10,TPI1,TYMP,UBA52 |
| | | Virchow | ACTB,ACTG1,ARHGDIA,ATP5F1B,ATP6V0C,B2M,BIRC3,BTG2,C1QA,C1QB,C1S,C3,CD24,CD74,CFL1,CKB,CNDP2,CP,CRYAB,CST3,CTSD,DDIT4,DUSP1,EEF1A1,EEF1D,EEF1G,EEF2,ENO1,EPAS1,FTL,FUS,GPX3,HBA2,HLA-A,HLA-B,HLA-DPA1,HLA-DRA,HLA-E,HNRNPA2B1,HNRNPK,ICAM1,IGFBP5,IGFBP7,IL32,ITM2B,JUN,LDHA,LITAF,MALAT1,MCL1,MT-ATP6,MT-CO1,MT-CO2,MT-CO3,MT-CYB,MT-ND2,MT-ND3,MT-ND4,MT-ND4L,NEAT1,NNMT,P4HB,PFN1,PGK1,PSAP,RACK1,RPL10,RPL13,RPL13A,RPL18A,RPL19,RPL21,RPL27A,RPL28,RPL3,RPL32,RPL35,RPL41,RPL7,RPL9,RPS11,RPS19,RPS2,RPS25,RPS27,RPS28,RPS29,RPS5,SEC61A1,SERPING1,SLC16A3,SOD2,TAGLN2,TAPBP,TMBIM6,TMEM176B,TMSB10,TYMP,UBA52,UBC |
| | | Virchow2 | ACTB,AKR1C3,ATP1B1,BIRC3,C1S,C3,CA12,CALR,CANX,CD24,CD74,CFL1,CKB,CLU,CNDP2,COX6C,CP,CRYAB,CSTB,CTSD,CYB5R3,DDIT4,EEF1A1,EEF1G,EEF2,ENO1,FGB,FGG,FTH1,FXYD2,GAPDH,GPX3,GSDMD,HLA-A,HLA-B,HP,HSP90B1,HSPB1,IGFBP3,IGFBP4,ITM2B,LBP,LDHA,MALAT1,MAP7D2,MGST1,MIF,MT-ATP6,MT-CO1,MT-CO2,MT-CO3,MT-CYB,MT-ND2,MT-ND3,MT-ND4,MT-ND4L,MT-ND5,MUC1,MYOCOS,NDRG1,NDUFA4L2,NEAT1,NNMT,P4HB,PABPC1,PDK4,PDZK1IP1,PFN1,PGK1,PKM,PLOD2,RNASET2,RPL13,RPL19,RPL3,RPL39,RPL4,RPL41,RPL7,RPL8,RPLP0,RPLP1,RPS19,RPS2,RPS20,RPS21,S100A10,SERPINA1,SERPING1,SLC16A3,SOD2,SPP1,TAPBP,TGFBI,TMBIM6,TMSB10,TPI1,TPT1,TRAM1,UBA52 |
| | | ViTB16 | AC015722.2,ADAMTS9,AGT,AIMP1,AL136295.2,AL162458.1,AL583785.1,ALG10B,ALPL,ANKZF1,ARGFX,ARMC2,ATG16L1,BARX2,C11orf54,CAPN6,CCR7,CD24,CGREF1,CP,CPTP,CXCL14,CXCL2,CYC1,DDIT4,DNAJC13,DPYS,DTHD1,EPB41,EPS8L2,ERO1B,FAM199X,FARSB,FLNC,FXYD2,GATM,GPX3,GSDMD,GSTA1,GSTA2,HBA2,HIPK2,HP,IGSF8,IL17RD,INAVA,KRT18,LINC00887,LINC01320,MAP7D2,MEGF11,MGST1,MPV17L,MT-ATP6,MT-CO1,MT-CO2,MT-CO3,MT-CYB,MT-ND1,MT-ND2,MT-ND3,MT-ND4,MT-ND4L,MT-ND5,MT3,MTHFSD,MVK,MYCT1,NLRC5,NUDT5,OSBPL11,PDIA5,PDK4,PDZK1,PDZK1IP1,PGAM2,PIK3R1,PLA1A,PPL,PPP1R16A,PRPF39,RAD50,SAA1,SDHAF3,SHISA8,SLC5A12,SLC6A8,SLC9B1,SMIM24,SORBS2,STXBP5-AS1,TARBP2,TMEM191B,TMEM9,TRIB3,TRMT13,TTC21A,YPEL2,ZBED6CL,ZNF487 |
| Small Region | Positively Correlated Genes (late- enriched genes) | CONCH | AC245407.2,ACTA2,ADGRD1,ADH1B,AEBP1,ANKS1B,ANTXR1,AP001160.2,CCDC80,CCN1,CCN2,CDH3,CEMIP,CNKSR1,CNN1,COL10A1,COL11A1,COL12A1,COL14A1,COL1A1,COL1A2,COL3A1,COL5A2,COL8A1,COMP,CORO2B,CRISPLD2,CSTF2T,CXCL12,DCN,DKK2,DKK3,DPYSL3,EFEMP1,EMILIN1,FAP,FGF7,FMO2,FNDC1,FOXC2,FST,GJA5,GLI2,GSDMA,HDGFL3,HPSE2,IGF1,IGHG1,IGHG3,IGKC,IGLC2,IGLC3,INHBA,ISM1,ITGBL1,KIF26B,LRRC15,LTBP2,LUM,MEDAG,MFAP4,MFAP5,MGP,MMP2,MRGPRF,MT1E,MYH10,NEXN,NGEF,OMD,P2RY10,PALLD,PCDHA7,PDE6A,PDPN,PID1,PLN,PNOC,PPP1R14A,PRELP,PRRX1,PTGIS,RUNX2,SFRP1,SFRP2,SFRP4,SHISA3,SPARC,SPON1,SULF1,TAGLN,THBS1,THBS2,TIMP3,TPM2,UCHL1,VGLL3,WT1-AS,Z98257.1,ZFPM2 |
| | | CONCH15 | AC004865.2,AC130895.1,ACTA2,AD000090.1,AEBP1,ANTXR1,APOD,ARL17A,CBWD2,CCDC80,CCN1,CCN2,CEMIP,CNKSR1,COL10A1,COL11A1,COL12A1,COL14A1,COL16A1,COL1A1,COL1A2,COL3A1,COL5A1,COL5A2,COL8A1,COMP,CRISPLD2,DCN,DKK2,EFEMP1,ELN,EMILIN1,FAM180A,FAP,FGF7,FMO2,FNDC1,FSTL1,GJA5,GLI2,HDGFL3,IGF1,IGHG1,IGHG2,IGHG3,IGHG4,IGHGP,IGKC,IGLC2,IGLC3,IL7R,INHBA,ISLR,ISM1,ITGBL1,JCHAIN,LIMD1-AS1,LRRC17,LTBP2,LUM,LYRM7,MFAP4,MFAP5,MGP,MMP2,MRGPRF,MT1M,MXRA8,MYL9,MZB1,NEXN,NUDT10,OMD,PALLD,PCDHA7,PDE6A,PNOC,PODN,POSTN,PRELP,PTGIS,RFLNA,SCRG1,SCUBE2,SFRP2,SFRP4,SHISA3,SPARC,SPON1,SULF1,TAGLN,THBS1,THBS2,TIMP3,TPM2,UCHL1,WARS2-AS1,WDR74,Z98257.1,ZBP1 |
| | | CTransPath | AC010186.3,AC099066.2,AC107373.2,ACTA2,AL021978.1,AL031963.3,AL139393.3,AL160006.1,ALCAM,ALDH1L2,ARNTL2,C3orf80,CCDC80,CCL18,CCL2,CCN1,CCN2,CCN4,CD300LF,CHCHD6,COL12A1,COL14A1,COL1A1,COL1A2,COL3A1,COL5A1,COL8A1,COLEC12,COMP,COX7A1,CYP27A1,DCN,EMILIN1,ESPN,FCGR2B,FNDC1,GLT8D1,GPNMB,GPR1,GPR171,GUSB,HDGFL3,HECW1,HMBOX1,HSCB,IGF1,IGKV3-15,IL16,INHBA,ITGAD,ITGBL1,KIAA1586,LILRB4,LIPA,LUM,LYRM7,LYZ,ME1,MFAP5,MPP1,MSR1,MT1E,MYH10,NPR2,NUDT22,PALLD,PCBP4,PDE4A,PHF23,PHYHD1,POSTN,PPP1R14A,PRMT9,PTGIS,PTPRO,PTX3,RBP1,RMST,RRP9,RSPH14,S100A6,SFRP4,SHISA3,SLC16A5,SLC20A2,SLC22A18AS,SLC46A3,SPON1,SULF1,TAGLN,TIMP3,TNC,TNFAIP8L2,TPBGL,TPM2,UCHL1,VGLL3,WLS,XYLT1,Z97192.3 |
| | | GigaPath | AC000403.1,AC004490.1,AC005072.1,AC006001.2,AC007952.3,AC020661.1,AC022306.2,AC025280.1,AC039056.2,AC080005.1,AC092171.4,AC093426.1,AC097662.1,AC104035.1,AC106739.1,AC113383.1,AC138781.1,ACKR4,ACTN3,AD000090.1,ADGRB2,AJAP1,AL049796.1,AL078587.2,AL137002.1,AL353600.2,ANKRD20A3,AP000708.1,AP000873.2,APOD,ARHGEF26,ARL17A,BTBD11,CCDC26,CCDC80,CCN1,CDH3,CEND1,CNKSR1,COL10A1,COL14A1,COMP,CRYBG2,CTAGE1,CTSG,DCN,DKK2,F7,FAM92B,FNDC1,FREM1,HIST1H2AE,HPSE2,HSD17B2,HTR2A,IGF1,IL1RL2,INHBA,ISM1,ITGBL1,KCNQ4,KIF26B,LGI1,LRRD1,LUM,MAP6,MATN3,MFAP5,MRGPRF,NEGR1,NEXN,NPB,NPR3,PCBP4,PDE6A,PRELP,PRSS35,PTGFR,PTGIS,RAB23,RAB39B,RASL11B,RSPO3,SFRP4,SFTPD-AS1,SHISA3,SLC16A12-AS1,SPOCD1,STXBP5,SULF1,TIMP3,TMEM9B-AS1,TRAIP,TRIT1,TSEN2,UBXN10-AS1,VEGFD,WFDC1,ZNF662,ZNF704 |
| | | RN18 | AC008915.2,AC016723.1,AC016831.1,AC016831.5,AC106791.1,ACTA2,ADAM12,ADGRB2,AEBP1,AL133330.1,ANTXR1,AP001160.2,BLK,CCDC80,CCL21,CCN1,CCN2,CD79A,COL10A1,COL11A1,COL12A1,COL1A1,COL1A2,COL3A1,COL5A1,COL8A1,COMP,CPEB1,CRISPLD2,DCN,DKK2,DLC1,EFEMP1,ELN,F3,FAP,FBN1,FCRL3,FILIP1,FMO2,FNDC1,GMNC,GREM1,HOXD10,HRK,IGF1,IGHG1,IGHG2,IGHG3,IGHG4,IGHGP,IGKC,IGLC2,IGLC3,INHBA,ISM1,ITGBL1,KLF2,LOXL1-AS1,LTBP2,LUM,MEF2B,MEMO1,MFAP4,MFAP5,MGP,MMP2,NEXN,OR7D2,P2RY10,PALLD,PDE6A,PID1,PLAT,PLSCR4,PODN,PRRX1,PTGDS,PTGIS,PWWP2B,RUNX2,SCRG1,SFRP2,SFRP4,SHISA3,SLIT3,SPON1,SRGAP2,STK38L,SULF1,TAGLN,THBS1,THBS2,TIMP3,TMEM169,TNFRSF10C,TPM2,TWIST2,VCL,ZNF117 |
| | | UNI | AC004771.4,AC005072.1,AC025280.1,AC044839.1,AC090579.1,AC093426.1,AC113383.1,ACKR4,ACTA2,AD000090.1,AL160006.1,AL353600.2,AL356310.1,AL391684.1,ANTXR1,AP000873.2,APOD,ARL17A,BMX,CCDC80,CCN1,CCN2,CD1E,CDH3,CMTM2,CNKSR1,COL10A1,COL11A1,COL12A1,COL14A1,COL1A1,COL1A2,COL3A1,COL8A1,COMP,CRYBG2,CTSG,DCN,DKK2,DKK3,FAM92B,FAP,FGF7,FNDC1,GNAL,HOXC8,HPD,IGF1,IGHG1,IGHG3,IGKC,IGKV1-5,IGKV3-15,IGLC2,IGLC3,INHBA,ISLR,ISM1,ITGA11,ITGBL1,KCNQ4,KIAA1841,LIMD1-AS1,LRRD1,LUM,LYRM7,MAP6,MATN3,MFAP4,MFAP5,MGP,MRGPRF,MYH10,NEXN,NGEF,OMD,OR1I1,P2RY10,PDE6A,PLN,PRELP,PRRX1,PRSS35,PTGIS,RSPO3,SFRP4,SHISA3,SIX4,SPOCD1,SPON1,SULF1,TAGLN,TIMP3,TPM2,UCHL1,VGLL3,WT1-AS,Z98257.1,ZFHX4,ZFPM2 |
| | | UNI2h | AC004865.2,AC073896.2,ACKR4,ACTA2,AD000090.1,AEBP1,AL391684.1,ANTXR1,AP000873.2,APOD,ARHGEF26,ARL17A,CCDC80,CCN1,CCN2,CDH3,CEMIP,CNKSR1,COL10A1,COL11A1,COL12A1,COL14A1,COL1A1,COL1A2,COL3A1,COL5A2,COL8A1,COMP,CRISPLD2,DCN,DKK2,DKK3,EFNB3,ELOVL4,FAP,FGF7,FNDC1,GGTLC2,GNAL,HDGFL3,HOXA5,IGF1,IGHG1,IGHG2,IGHG3,IGHG4,IGHGP,IGKC,IGKV3-15,IGLC2,IGLC3,INHBA,ISLR,ISM1,ITGA11,ITGBL1,KIF26B,LIMD1-AS1,LINC01048,LRRC17,LTBP2,LUM,MEDAG,MFAP4,MFAP5,MGP,MMP2,MRGPRF,MXRA8,MYH10,NEXN,NGEF,NLGN2,OMD,P2RY10,PDE6A,PLN,PNOC,POSTN,PRELP,PRKG1,PTGIS,RASL10A,RUNX2,SFRP2,SFRP4,SHISA3,SPON1,SULF1,TAGLN,THBS1,THBS2,TIMP3,TPM2,UCHL1,VGLL3,WT1-AS,Z98257.1,ZFHX4,ZFPM2 |
| | | Virchow | AC090204.1,AC093323.1,ACKR4,ACTA2,ADAMTSL5,ALDH1L2,ANXA4,AP000424.1,APTR,ARF4,ATP5MG,BNIP3,CALM1,CBX1,CCDC80,CCN1,CCN2,CDK14,COL11A1,COL12A1,COL14A1,COL1A2,COL3A1,COL8A1,DCN,DPY30,EEF1AKMT4,EIF2B1,FBXO17,FDFT1,FNDC1,FO704657.1,GAS5,GSTP1,HOXA5,HP,HPS3,HSCB,KIF26B,LRP6,LUM,MAP3K11,MCAT,MFAP5,MIR210HG,MIR4458HG,MLLT11,MPP6,MT1X,MT2A,MYO5B,NDRG1,NDUFA6,NIFK,OAS1,PABPC1,PALLD,PCBP4,PDAP1,POC1A,POLQ,PPP1R13L,PRDM4,PRKAB1,PRSS23,PTGIS,PYROXD1,RB1CC1,RCAN3,RER1,RNF41,ROCK2,RPL23,RPL23A,RPL36A,RPL38,RPL39,RPL6,RPL8,RPLP0,RPLP1,RPS4X,RPS8,SERF2,SERPINA3,SFRP4,SHISA3,SIX5,SLC17A3,SMIM30,SYNCRIP,TCEA1,TIMP3,TMEM64,TNFRSF19,TOX4,TPM1,TPT1,VPS18,ZNF805 |
| | | Virchow2 | ACTA2,AEBP1,AKAP12,AL691447.2,ANTXR1,ARL17A,BGN,CALD1,CCDC80,CCN1,CCN2,CCN4,CDC42EP1,CDH3,COL10A1,COL11A1,COL12A1,COL14A1,COL16A1,COL1A1,COL1A2,COL3A1,COL4A2,COL5A1,COL5A2,COL6A3,COL8A1,COMP,CRISPLD2,CRYBG2,CSRP1,CTSK,DCN,DKK3,EFEMP1,ELN,EMILIN1,F3,FAM155A,FAP,FBN1,FILIP1,FNDC1,FSTL1,IGF1,IGHG3,IGHG4,IGHGP,IGKC,IGLC2,IGLC3,INHBA,ISLR,ITGBL1,KCNK13,KIF26B,LIMS2,LTBP2,LUM,MFAP5,MGP,MMP2,MT1E,MXRA5,MXRA7,MXRA8,MYH10,MYL9,MYLK,NEXN,OXTR,PALLD,PLAT,PLN,POSTN,PRELP,PRKG1,PRRX1,PRSS23,PTGIS,RAB23,RFLNA,SFRP2,SFRP4,SHISA3,SPARC,SPOCD1,SPON1,SULF1,TAGLN,THBS1,THBS2,TIMP2,TIMP3,TMEM98,TPM2,UCHL1,VCAN,VCL,ZFPM2 |
| | | ViTB16 | ACTA2,AEBP1,ALCAM,ANTXR1,ARL17A,BGN,CALD1,CAPG,CCDC80,CCN1,CCN2,CD52,CDC42EP5,CDH11,CNN1,COL10A1,COL11A1,COL12A1,COL14A1,COL16A1,COL1A1,COL1A2,COL3A1,COL5A1,COL5A2,COL6A2,COL8A1,COMP,CRISPLD2,CSRP1,CTSB,DAAM1,DCN,DKK3,EFEMP1,EMILIN1,EMP1,FAP,FBN1,FNDC1,FOS,FSTL1,GAS6,GJA5,GLIPR1,GPNMB,HAMP,HIRA,HLA-DRA,HSPA8,IGF1,IGHG3,IGKC,IGLC3,INHBA,ITGBL1,KCTD9,LILRB4,LRP1,LTBP2,LUM,LYZ,MARCKS,MFAP5,MGP,MMP2,MRGPRF,MSR1,MT1E,MT2A,MXRA5,MYH10,MYL9,NPL,OLFML2B,PALLD,PDGFRA,POSTN,PRRX1,PRSS23,PTGIS,RAB23,S100A6,SERPINF1,SFRP2,SFRP4,SHISA3,SLC46A3,SPARC,SPON1,SRF,SULF1,TAGLN,THBS1,THBS2,TIMP1,TIMP2,TIMP3,TPM2,WDR74 |
| | Negatively Correlated Genes (early- enriched genes) | CONCH | ADGRG1,ANGPTL4,ATP1B1,BIRC3,C3,CA12,CALR,CANX,CD24,CKB,CLU,CNDP2,COX6A1,CP,CRYAB,CSTB,CYC1,DDIT4,DEGS1,EEF1G,ENO1,ERRFI1,FGB,FGG,FSTL3,FXYD2,GAPDH,GPX3,GRINA,GSDMD,GUK1,HBA2,HINT1,HLA-A,HP,HSP90B1,HSPB1,IGFBP3,IGFBP4,ITIH5,KRT18,KRT8,LAMB3,LBP,LDHA,LOX,MAP7D2,MGST1,MIF,MLEC,MT-ATP6,MT-ATP8,MT-CO1,MT-CO2,MT-CO3,MT-CYB,MT-ND2,MT-ND3,MT-ND4,MT-ND4L,MT-ND5,MYOCOS,NDRG1,NDUFA4L2,NEAT1,NNMT,NUDT4,NUPR1,OS9,P3H2,P4HB,PABPC1,PDK4,PDZK1IP1,PGK1,PKM,PLOD2,PRKCSH,RNASET2,RPL19,RPL8,RPLP0,RPS19,RPS20,RPS21,S100A10,SAA1,SERPINA1,SERPINF2,SERPING1,SLC16A3,SNHG25,SOD2,SPP1,TAGLN2,TGFBI,TMBIM6,TMSB10,TPI1,YBX3 |
| | | CONCH15 | AC090578.1,ADGRG1,AKR1C3,ANGPTL4,ATP1B1,BIRC3,C1R,C1S,C3,CA12,CALR,CANX,CD24,CERNA2,CERS2,CKB,CLU,CNDP2,COX6A1,COX6C,CP,CPD,CRYAB,CSRP2,CSTB,CXCL14,CYC1,DDIT4,DEGS1,FGA,FGB,FGG,FRZB,FXYD2,GAL3ST1,GAPDH,GPX3,GSDMD,HINT1,HLA-A,HP,HSP90B1,HSPB1,IGFBP3,IGFBP4,ITIH5,KRT18,KRT8,LBP,LDHA,LRP2,MAP7D2,MGST1,MIF,MLEC,MT-ATP6,MT-ATP8,MT-CO1,MT-CO2,MT-CO3,MT-CYB,MT-ND2,MT-ND3,MT-ND4,MT-ND4L,MT-ND5,MUC1,MYOCOS,NDRG1,NDUFA4L2,NEAT1,NNMT,NUPR1,P3H2,P4HB,PABPC1,PDK4,PDZK1IP1,PGK1,PKM,PLOD2,RNASET2,RPL23A,RPLP0,RPLP1,RPS20,S100A10,SAA1,SCARB1,SERPINA1,SERPINF2,SLC16A3,SNHG25,SOD2,SPP1,TIPARP,TMBIM6,TPI1,WFDC2,YBX3 |
| | | CTransPath | ADGRG1,AKR1C3,ALPL,ANKZF1,ATP1B1,BIRC3,BTG2,C1orf54,C3,CD24,CFB,CKB,CLU,CNDP2,COL23A1,CP,CRYAB,CXCL14,CYC1,DEGS1,DPP4,EEF2,EFNA1,EPS8L2,ERGIC1,FDPS,FGB,FXYD2,GAL3ST1,GAPDH,GIGYF1,GPX3,GSDMD,HBA2,HSD17B7,HSP90B1,ICA1L,IGFBP5,IGHM,IGLC1,ILF3,ISOC2,ITIH5,KCNJ16,LBP,LDHA,LINC01320,MALAT1,MAOA,MAP7D2,MEGF11,MGST1,MIF,MT-ATP6,MT-CO1,MT-CO2,MT-CO3,MT-CYB,MT-ND1,MT-ND2,MT-ND3,MT-ND4,MT-ND4L,MT-ND5,MUC1,NOL3,P4HB,PARP1,PDK4,PDZK1,PDZK1IP1,PGF,PHB2,PLA1A,PPP1R16A,PRDX2,PREX2,PSAT1,RERE,RGS5,RIDA,RPL19,RPL35A,RPL7,RPN2,SAA1,SEC61A1,SERPINF2,SFXN1,SGPP2,SNHG25,SNX10,SOD2,SPINK13,TAGLN2,TCF25,TP53INP1,TSPAN12,WFDC2,ZBED6CL |
| | | GigaPath | A2M,ACTB,ACTG1,ACTR2,APOC1,APOE,ARHGDIA,ATP6V0C,B2M,BTG2,C1QA,C1QB,CD24,CD74,CFL1,CRIP1,CST3,CTSZ,DDX3X,DUSP1,EEF1G,EEF2,ENO1,EPAS1,ERGIC1,FCER1G,FGL2,FTL,FUS,GLUL,HBA2,HLA-A,HLA-B,HLA-C,HLA-DPA1,HLA-DPB1,HLA-DQB1,HLA-DRA,HLA-E,HNRNPK,IGFBP7,IGHA1,IGHM,IL32,INPP5D,LCP2,LITAF,MCL1,MMP7,MT-ATP6,MT-CO1,MT-CO2,MT-CO3,MT-CYB,MT-ND2,MT-ND3,MT-ND4,MT-ND4L,NLRC5,PFN1,PSAP,PSMB9,RACK1,RGS1,RGS2,RGS5,RPL13,RPL18A,RPL19,RPL21,RPL27A,RPL28,RPL3,RPL32,RPL35,RPL35A,RPL41,RPL7,RPL7A,RPS11,RPS2,RPS27,RPS5,SAT1,SEC61A1,SERPING1,STAT2,TAGLN2,TAPBP,TMBIM6,TMEM176B,TMEM94,TMSB10,TNFSF13,TPI1,TYMP,UBA52,UBB,VWF,WARS |
| | | RN18 | ADGRG1,AGT,AKR1C3,ANGPTL4,ATP1B1,BIRC3,BSG,C1S,C3,CA12,CALR,CANX,CD24,CERS2,CKB,CLU,CNDP2,COX6A1,COX6C,CP,CRYAB,CXCL14,DDIT4,DEGS1,DPP4,ENO1,FGA,FGB,FGG,FRZB,FXYD2,GAL3ST1,GAPDH,GNS,GPX3,GRINA,GSDMD,HLA-A,HP,HSP90B1,HSPB1,HSPB8,IGFBP3,IGFBP4,ITIH5,ITM2B,KRT18,KRT8,LAMB3,LBP,LDHA,LRP2,MAP7D2,MIF,MLEC,MT-ATP6,MT-CO1,MT-CO2,MT-CO3,MT-CYB,MT-ND2,MT-ND3,MT-ND4,MT-ND4L,MT-ND5,MUC1,MYOCOS,NDRG1,NDUFA4L2,NEAT1,NFE2L1,NNMT,NUPR1,OS9,P4HB,PABPC1,PDK4,PDZK1IP1,PGK1,PKM,PLOD1,RNASET2,RPL23,RPL35A,RPLP0,RPLP1,RPS20,SAA1,SERPINA1,SERPINF2,SLC16A3,SNHG6,SOD2,SPP1,TGFBI,TMBIM6,TMED9,TPI1,TUBA1B,WFDC2 |
| | | UNI | ADGRG1,AKR1C3,ATP1B1,BIRC3,BTG2,C1S,C3,CA12,CALR,CANX,CD24,CD74,CFL1,CKB,CLU,CNDP2,COX6C,CP,CPD,CRYAB,CSTB,CTSD,CXCL14,CYC1,DDIT4,EEF1G,EEF2,ENO1,FGB,FXYD2,GAL3ST1,GAPDH,GPX3,GSDMD,HLA-A,HLA-B,HP,HSP90B1,HSPB1,IGFBP3,IGFBP4,ITIH5,ITM2B,LBP,LDHA,MALAT1,MAP7D2,MCL1,MGST1,MIF,MT-ATP6,MT-CO1,MT-CO2,MT-CO3,MT-CYB,MT-ND1,MT-ND2,MT-ND3,MT-ND4,MT-ND4L,MT-ND5,MUC1,MYOCOS,NDRG1,NDUFA4L2,NEAT1,NNMT,P4HB,PABPC1,PDK4,PDZK1IP1,PFN1,PGK1,PKM,PLOD2,RPL13,RPL19,RPL3,RPL35A,RPL41,RPL7,RPLP0,RPLP1,RPN2,RPS2,RPS20,S100A10,SERPINA1,SERPINF2,SERPING1,SLC16A3,SOD2,SPP1,TAPBP,TIPARP,TMBIM6,TMEM176B,TMSB10,TPI1,UBA52 |
| | | UNI2h | ADGRG1,AKR1C3,ATP1B1,BIRC3,C1S,C3,CA12,CALR,CANX,CD24,CERS2,CKB,CLU,CNDP2,COX6A1,COX6C,CP,CPD,CRYAB,CSTB,CXCL14,CYB5R3,CYC1,DDIT4,ENO1,FGB,FXYD2,GAL3ST1,GAPDH,GPX3,GSDMD,HINT1,HLA-A,HLA-B,HLA-E,HP,HSP90B1,HSPB1,HSPD1,IGFBP3,IGFBP4,ITIH5,KRT18,KRT8,LBP,LDHA,MALAT1,MAP7D2,MCL1,MGST1,MIF,MLEC,MT-ATP6,MT-CO1,MT-CO2,MT-CO3,MT-CYB,MT-ND1,MT-ND2,MT-ND3,MT-ND4,MT-ND4L,MT-ND5,MUC1,MYOCOS,NDRG1,NDUFA4L2,NEAT1,NNMT,NUPR1,P4HB,PABPC1,PDK4,PDZK1IP1,PGK1,PKM,PLOD2,PRDX2,RNASET2,RPL19,RPL7,RPLP0,RPLP1,RPN2,RPS19,RPS20,S100A10,SAA1,SERPINA1,SERPINF2,SERPING1,SLC16A3,SOD2,SPP1,TGFBI,TIPARP,TMBIM6,TMSB10,TPI1,TRAM1 |
| | | Virchow | AAAS,AASS,AC010331.1,AC073111.4,AC104971.1,AC107021.2,ACVRL1,AIM2,AL158835.1,AL391244.2,ALB,ALG8,ALKBH6,ANKRD17,ARL5A,ATP13A4,ATP2A3,ATP6AP1,BLOC1S2,BOC,C1QA,C1orf174,C5AR2,CAMSAP3,CCR7,CD19,CD79A,CDS2,CEP164,CHST2,CIITA,CIPC,CPSF2,DDX49,DENND6B,DIPK1B,DZIP3,ENAH,EPAS1,FAM174A,FBF1,FLVCR1-DT,GEMIN7-AS1,GIGYF1,GM2A,GORAB,H19,HBA2,IGHG4,IGHM,IGLC1,IL33,INPP5D,KIF14,KIF7,LFNG,LINC01871,LPAR6,MED14,NFS1,NIP7,NSUN6,OAS2,ORAI2,OTUD5,OXER1,P2RY1,POU2AF1,POU5F1B,PRDM11,RAB3GAP1,RASGEF1B,RASGRP2,RBM39,RGS2,RNF214,ROBO3,RSAD2,SEPTIN6,SHISA8,SP100,SPIB,SPOCK2,SRP54,STIM2,TIRAP,TMC6,TMEM30B,TRAF3IP3,TTC25,ULK1,VPS37C,WRN,XRRA1,ZBTB47,ZNF25,ZNF283,ZNF440,ZNF549,ZNF558 |
| | | Virchow2 | AC090578.1,ADGRG1,AKR1C3,ASPH,ATP1B1,BIRC3,BSG,C3,CA12,CALR,CANX,CD24,CDH6,CERNA2,CERS2,CKB,CLU,CNDP2,COX6A1,CP,CPD,CRYAB,CSTB,CXCL14,CYB5R3,CYC1,DDIT4,DEGS1,DPP7,EPS8L2,FGB,FGG,FXYD2,GAL3ST1,GAPDH,GATM,GPX3,GSDMD,GSTA1,HBA2,HP,HSP90B1,HSPB1,HSPD1,IGFBP3,IGFBP4,ITIH5,KRT18,KRT8,LAMB3,LBP,LDHA,LRP2,MAP7D2,MGST1,MIF,MLEC,MPV17L,MT-ATP6,MT-ATP8,MT-CO1,MT-CO2,MT-CO3,MT-CYB,MT-ND1,MT-ND2,MT-ND3,MT-ND4,MT-ND4L,MT-ND5,MUC1,MVK,MYOCOS,NDRG1,NDUFA4L2,NNMT,NUDT4,NUPR1,OSBPL11,P4HB,PABPC1,PDIA5,PDK4,PDZK1IP1,PGK1,PKM,POU5F1,PRDX2,RNASET2,RPS20,SAA1,SERPINA1,SERPINF2,SLC16A3,SPP1,TMBIM6,TNIP1,TPI1,TSKU,YBX3 |
| | | ViTB16 | AC090578.1,ADGRG1,AGT,AKR1C3,ANGPTL4,ATP1B1,BICC1,BIRC3,C3,CAPN6,CD24,CELSR1,CERS2,CKB,CLU,CNDP2,CP,CPNE3,CRYAB,CXCL14,CYC1,DDIT4,DEGS1,EFNA1,ERRFI1,FGA,FGB,FXYD2,GAPDH,GPX3,GSDMD,GSTA1,GSTA2,HBA2,HGD,HP,IFT57,IGFBP4,ISOC2,ITGB4,ITIH5,KRT18,LBP,LDHA,LINC00887,LINC01320,LINC02802,LYPLA1,MAP7D2,MGST1,MIF,MLEC,MT-ATP6,MT-ATP8,MT-CO1,MT-CO2,MT-CO3,MT-CYB,MT-ND1,MT-ND2,MT-ND3,MT-ND4,MT-ND4L,MT-ND5,MT3,MUC1,MVK,NDUFA4L2,P4HB,PABPC1,PDK4,PDZK1IP1,PGK1,PKM,PLOD1,POU5F1,RAB17,RNF19A,RPS20,SAA1,SERPINF2,SLC16A3,SLC3A1,SPINK13,SPON2,SPP1,SPSB1,STC2,TIPARP,TMEM9,TMEM94,TNFSF10,TP53INP1,WFDC2,XYLT2,YBX3,ZBED6CL,ZNF114,ZNF611 |

### Table.S8

| sample | tissue | link | Source and cancer subtypes | Transcription technology |
|---|---|---|---|---|
| MEND65 | brain | https://www.nature.com/articles/s41467 -023-36071-5 | Pediatric brain tumors: Medulloblastoma (WNT subtype); NOS subtype | Visium |
| MEND66 | brain | https://www.nature.com/articles/s41467 -023-36071-5 | Pediatric brain tumors: Medulloblastoma (WNT subtype); NOS subtype | Visium |
| NCBI629 | brain | https://academic.oup.com/neuro -oncology/article/24/Supplement_1/i45/6600572 | Spinal ependymoma (SP-EPN): subtype A/B; pediatric ependymoma: supratentorial ependymoma ZFTA fusion type (ST-EPN-ZFTA), posterior fossa group A ependymoma (PF-EPN-A) | Visium |
| NCBI631 | brain | https://academic.oup.com/neuro -oncology/article/24/Supplement_1/i45/6600572 | Spinal ependymoma (SP-EPN): subtype A/B; pediatric ependymoma: supratentorial ependymoma ZFTA fusion type (ST-EPN-ZFTA), posterior fossa group A ependymoma (PF-EPN-A) | Visium |
| NCBI632 | brain | https://academic.oup.com/neuro -oncology/article/24/Supplement_1/i45/6600572 | Spinal ependymoma (SP-EPN): subtype A/B; pediatric ependymoma: supratentorial ependymoma ZFTA fusion type (ST-EPN-ZFTA), posterior fossa group A ependymoma (PF-EPN-A) | Visium |
| NCBI635 | brain | https://academic.oup.com/neuro -oncology/article/24/Supplement_1/i45/6600572 | Spinal ependymoma (SP-EPN): subtype A/B; pediatric ependymoma: supratentorial ependymoma ZFTA fusion type (ST-EPN-ZFTA), posterior fossa group A ependymoma (PF-EPN-A) | Visium |
| NCBI638 | brain | https://academic.oup.com/neuro -oncology/article/24/Supplement_1/i45/6600572 | Spinal ependymoma (SP-EPN): subtype A/B; pediatric ependymoma: supratentorial ependymoma ZFTA fusion type (ST-EPN-ZFTA), posterior fossa group A ependymoma (PF-EPN-A) | Visium |
| NCBI640 | brain | https://academic.oup.com/neuro -oncology/article/24/Supplement_1/i45/6600572 | Spinal ependymoma (SP-EPN): subtype A/B; pediatric ependymoma: supratentorial ependymoma ZFTA fusion type (ST-EPN-ZFTA), posterior fossa group A ependymoma (PF-EPN-A) | Visium |
| TENX30 | brain | https://www.10xgenomics.com/datasets/human -glioblastoma -targeted -pan-cancer -panel -1-standard -1-2-0 | Pleomorphic glioblastoma | Visium |
| TENX31 | brain | https://www.10xgenomics.com/datasets/human -glioblastoma -whole -transcriptome -analysis -1-standard -1-2-0 | Pleomorphic glioblastoma | Visium |
| SPA100 | breast | https://www.science.org/doi/10.1126/science.aaf2403 | Invasive ductal carcinoma (INV); ductal carcinoma in situ (DCIS) | Spatial Transcriptomics |
| SPA101 | breast | https://www.science.org/doi/10.1126/science.aaf2403 | Invasive ductal carcinoma (INV); ductal carcinoma in situ (DCIS) | Spatial Transcriptomics |
| SPA102 | breast | https://www.science.org/doi/10.1126/science.aaf2403 | Invasive ductal carcinoma (INV); ductal carcinoma in situ (DCIS) | Spatial Transcriptomics |
| SPA103 | breast | https://www.science.org/doi/10.1126/science.aaf2403 | Invasive ductal carcinoma (INV); ductal carcinoma in situ (DCIS) | Spatial Transcriptomics |
| SPA104 | breast | https://www.science.org/doi/10.1126/science.aaf2403 | Invasive ductal carcinoma (INV); ductal carcinoma in situ (DCIS) | Spatial Transcriptomics |
| SPA105 | breast | https://www.science.org/doi/10.1126/science.aaf2403 | Invasive ductal carcinoma (INV); ductal carcinoma in situ (DCIS) | Spatial Transcriptomics |
| SPA106 | breast | https://www.science.org/doi/10.1126/science.aaf2403 | Invasive ductal carcinoma (INV); ductal carcinoma in situ (DCIS) | Spatial Transcriptomics |
| SPA107 | breast | https://www.science.org/doi/10.1126/science.aaf2403 | Invasive ductal carcinoma (INV); ductal carcinoma in situ (DCIS) | Spatial Transcriptomics |
| SPA108 | breast | https://www.science.org/doi/10.1126/science.aaf2403 | Invasive ductal carcinoma (INV); ductal carcinoma in situ (DCIS) | Spatial Transcriptomics |
| SPA109 | breast | https://www.science.org/doi/10.1126/science.aaf2403 | Invasive ductal carcinoma (INV); ductal carcinoma in situ (DCIS) | Spatial Transcriptomics |
| SPA111 | breast | https://www.science.org/doi/10.1126/science.aaf2403 | Invasive ductal carcinoma (INV); ductal carcinoma in situ (DCIS) | Spatial Transcriptomics |
| SPA112 | breast | https://www.science.org/doi/10.1126/science.aaf2403 | Invasive ductal carcinoma (INV); ductal carcinoma in situ (DCIS) | Spatial Transcriptomics |
| SPA114 | breast | https://www.science.org/doi/10.1126/science.aaf2403 | Invasive ductal carcinoma (INV); ductal carcinoma in situ (DCIS) | Spatial Transcriptomics |
| SPA115 | breast | https://www.science.org/doi/10.1126/science.aaf2403 | Invasive ductal carcinoma (INV); ductal carcinoma in situ (DCIS) | Spatial Transcriptomics |
| SPA116 | breast | https://www.science.org/doi/10.1126/science.aaf2403 | Invasive ductal carcinoma (INV); ductal carcinoma in situ (DCIS) | Spatial Transcriptomics |
| SPA117 | breast | https://www.science.org/doi/10.1126/science.aaf2403 | Invasive ductal carcinoma (INV); ductal carcinoma in situ (DCIS) | Spatial Transcriptomics |
| SPA118 | breast | https://www.science.org/doi/10.1126/science.aaf2403 | Invasive ductal carcinoma (INV); ductal carcinoma in situ (DCIS) | Spatial Transcriptomics |
| SPA55 | breast | https://www.science.org/doi/10.1126/science.aaf2403 | Invasive ductal carcinoma (INV); ductal carcinoma in situ (DCIS) | Spatial Transcriptomics |
| SPA57 | breast | https://www.science.org/doi/10.1126/science.aaf2403 | Invasive ductal carcinoma (INV); ductal carcinoma in situ (DCIS) | Spatial Transcriptomics |
| SPA60 | breast | https://www.science.org/doi/10.1126/science.aaf2403 | Invasive ductal carcinoma (INV); ductal carcinoma in situ (DCIS) | Spatial Transcriptomics |
| SPA61 | breast | https://www.science.org/doi/10.1126/science.aaf2403 | Invasive ductal carcinoma (INV); ductal carcinoma in situ (DCIS) | Spatial Transcriptomics |
| SPA64 | breast | https://www.science.org/doi/10.1126/science.aaf2403 | Invasive ductal carcinoma (INV); ductal carcinoma in situ (DCIS) | Spatial Transcriptomics |
| SPA66 | breast | https://www.science.org/doi/10.1126/science.aaf2403 | Invasive ductal carcinoma (INV); ductal carcinoma in situ (DCIS) | Spatial Transcriptomics |
| SPA67 | breast | https://www.science.org/doi/10.1126/science.aaf2403 | Invasive ductal carcinoma (INV); ductal carcinoma in situ (DCIS) | Spatial Transcriptomics |
| SPA68 | breast | https://www.science.org/doi/10.1126/science.aaf2403 | Invasive ductal carcinoma (INV); ductal carcinoma in situ (DCIS) | Spatial Transcriptomics |
| SPA69 | breast | https://www.science.org/doi/10.1126/science.aaf2403 | Invasive ductal carcinoma (INV); ductal carcinoma in situ (DCIS) | Spatial Transcriptomics |
| SPA73 | breast | https://www.science.org/doi/10.1126/science.aaf2403 | Invasive ductal carcinoma (INV); ductal carcinoma in situ (DCIS) | Spatial Transcriptomics |
| SPA80 | breast | https://www.science.org/doi/10.1126/science.aaf2403 | Invasive ductal carcinoma (INV); ductal carcinoma in situ (DCIS) | Spatial Transcriptomics |
| SPA82 | breast | https://www.science.org/doi/10.1126/science.aaf2403 | Invasive ductal carcinoma (INV); ductal carcinoma in situ (DCIS) | Spatial Transcriptomics |
| SPA83 | breast | https://www.science.org/doi/10.1126/science.aaf2403 | Invasive ductal carcinoma (INV); ductal carcinoma in situ (DCIS) | Spatial Transcriptomics |
| SPA84 | breast | https://www.science.org/doi/10.1126/science.aaf2403 | Invasive ductal carcinoma (INV); ductal carcinoma in situ (DCIS) | Spatial Transcriptomics |
| SPA90 | breast | https://www.science.org/doi/10.1126/science.aaf2403 | Invasive ductal carcinoma (INV); ductal carcinoma in situ (DCIS) | Spatial Transcriptomics |
| SPA91 | breast | https://www.science.org/doi/10.1126/science.aaf2403 | Invasive ductal carcinoma (INV); ductal carcinoma in situ (DCIS) | Spatial Transcriptomics |
| SPA92 | breast | https://www.science.org/doi/10.1126/science.aaf2403 | Invasive ductal carcinoma (INV); ductal carcinoma in situ (DCIS) | Spatial Transcriptomics |
| SPA93 | breast | https://www.science.org/doi/10.1126/science.aaf2403 | Invasive ductal carcinoma (INV); ductal carcinoma in situ (DCIS) | Spatial Transcriptomics |
| SPA94 | breast | https://www.science.org/doi/10.1126/science.aaf2403 | Invasive ductal carcinoma (INV); ductal carcinoma in situ (DCIS) | Spatial Transcriptomics |
| SPA95 | breast | https://www.science.org/doi/10.1126/science.aaf2403 | Invasive ductal carcinoma (INV); ductal carcinoma in situ (DCIS) | Spatial Transcriptomics |
| SPA96 | breast | https://www.science.org/doi/10.1126/science.aaf2403 | Invasive ductal carcinoma (INV); ductal carcinoma in situ (DCIS) | Spatial Transcriptomics |
| SPA97 | breast | https://www.science.org/doi/10.1126/science.aaf2403 | Invasive ductal carcinoma (INV); ductal carcinoma in situ (DCIS) | Spatial Transcriptomics |
| SPA98 | breast | https://www.science.org/doi/10.1126/science.aaf2403 | Invasive ductal carcinoma (INV); ductal carcinoma in situ (DCIS) | Spatial Transcriptomics |
| SPA99 | breast | https://www.science.org/doi/10.1126/science.aaf2403 | Invasive ductal carcinoma (INV); ductal carcinoma in situ (DCIS) | Spatial Transcriptomics |
| MISC33 | colon | https://www.cell.com/cell/fulltext/S0092 -8674(23)01222 -9?_returnURL=https%3A%2F%2Flinkinghub.elsevier.com%2Fretrieve%2Fpii%2FS0092867423012229%3Fshowall%3Dtrue#secsectitle0020 | Precancerous lesion subtypes (TA/TVA; SSL/HP); colorectal cancer subtypes (MSS CRC/microsatellite stable; MSI-H CRC/microsatellite instability -high); special/cross phenotypes (HM type TA/TVA/MSS; MSI-H/CIN cross subtype) | Visium |
| MISC35 | colon | https://www.cell.com/cell/fulltext/S0092 -8674(23)01222 -9?_returnURL=https%3A%2F%2Flinkinghub.elsevier.com%2Fretrieve%2Fpii%2FS0092867423012229%3Fshowall%3Dtrue#secsectitle0020 | Precancerous lesion subtypes (TA/TVA; SSL/HP); colorectal cancer subtypes (MSS CRC/microsatellite stable; MSI-H CRC/microsatellite instability -high); special/cross phenotypes (HM type TA/TVA/MSS; MSI-H/CIN cross subtype) | Visium |
| MISC36 | colon | https://www.cell.com/cell/fulltext/S0092 -8674(23)01222 -9?_returnURL=https%3A%2F%2Flinkinghub.elsevier.com%2Fretrieve%2Fpii%2FS0092867423012229%3Fshowall%3Dtrue#secsectitle0020 | Precancerous lesion subtypes (TA/TVA; SSL/HP); colorectal cancer subtypes (MSS CRC/microsatellite stable; MSI-H CRC/microsatellite instability -high); special/cross phenotypes (HM type TA/TVA/MSS; MSI-H/CIN cross subtype) | Visium |
| MISC40 | colon | https://www.cell.com/cell/fulltext/S0092 -8674(21)01381 -7?_returnURL=https%3A%2F%2Flinkinghub.elsevier.com%2Fretrieve%2Fpii%2FS0092867421013817%3Fshowall%3Dtrue | Precancerous lesion subtypes (conventional adenomas/ADs; serrated polyps/SERs); colorectal cancer subtypes (MSS CRC/microsatellite stable; MSI-H CRC/microsatellite instability -high) | Visium |
| MISC41 | colon | https://www.cell.com/cell/fulltext/S0092 -8674(23)01222 -9?_returnURL=https%3A%2F%2Flinkinghub.elsevier.com%2Fretrieve%2Fpii%2FS0092867423012229%3Fshowall%3Dtrue#secsectitle0020 | Precancerous lesion subtypes (TA/TVA; SSL/HP); colorectal cancer subtypes (MSS CRC/microsatellite stable; MSI-H CRC/microsatellite instability -high); special/cross phenotypes (HM type TA/TVA/MSS; MSI-H/CIN cross subtype) | Visium |
| MISC42 | colon | https://www.cell.com/cell/fulltext/S0092 -8674(23)01222 -9?_returnURL=https%3A%2F%2Flinkinghub.elsevier.com%2Fretrieve%2Fpii%2FS0092867423012229%3Fshowall%3Dtrue#secsectitle0020 | Precancerous lesion subtypes (TA/TVA; SSL/HP); colorectal cancer subtypes (MSS CRC/microsatellite stable; MSI-H CRC/microsatellite instability -high); special/cross phenotypes (HM type TA/TVA/MSS; MSI-H/CIN cross subtype) | Visium |
| MISC45 | colon | https://www.cell.com/cell/fulltext/S0092 -8674(23)01222 -9?_returnURL=https%3A%2F%2Flinkinghub.elsevier.com%2Fretrieve%2Fpii%2FS0092867423012229%3Fshowall%3Dtrue#secsectitle0020 | Precancerous lesion subtypes (TA/TVA; SSL/HP); colorectal cancer subtypes (MSS CRC/microsatellite stable; MSI-H CRC/microsatellite instability -high); special/cross phenotypes (HM type TA/TVA/MSS; MSI-H/CIN cross subtype) | Visium |
| MISC47 | colon | https://www.cell.com/cell/fulltext/S0092 -8674(23)01222 -9?_returnURL=https%3A%2F%2Flinkinghub.elsevier.com%2Fretrieve%2Fpii%2FS0092867423012229%3Fshowall%3Dtrue#secsectitle0020 | Precancerous lesion subtypes (TA/TVA; SSL/HP); colorectal cancer subtypes (MSS CRC/microsatellite stable; MSI-H CRC/microsatellite instability -high); special/cross phenotypes (HM type TA/TVA/MSS; MSI-H/CIN cross subtype) | Visium |
| MISC48 | colon | https://www.cell.com/cell/fulltext/S0092 -8674(23)01222 -9?_returnURL=https%3A%2F%2Flinkinghub.elsevier.com%2Fretrieve%2Fpii%2FS0092867423012229%3Fshowall%3Dtrue#secsectitle0020 | Precancerous lesion subtypes (TA/TVA; SSL/HP); colorectal cancer subtypes (MSS CRC/microsatellite stable; MSI-H CRC/microsatellite instability -high); special/cross phenotypes (HM type TA/TVA/MSS; MSI-H/CIN cross subtype) | Visium |
| MISC49 | colon | https://www.cell.com/cell/fulltext/S0092 -8674(23)01222 -9?_returnURL=https%3A%2F%2Flinkinghub.elsevier.com%2Fretrieve%2Fpii%2FS0092867423012229%3Fshowall%3Dtrue#secsectitle0020 | Precancerous lesion subtypes (TA/TVA; SSL/HP); colorectal cancer subtypes (MSS CRC/microsatellite stable; MSI-H CRC/microsatellite instability -high); special/cross phenotypes (HM type TA/TVA/MSS; MSI-H/CIN cross subtype) | Visium |
| MISC50 | colon | https://www.cell.com/cell/fulltext/S0092 -8674(23)01222 -9?_returnURL=https%3A%2F%2Flinkinghub.elsevier.com%2Fretrieve%2Fpii%2FS0092867423012229%3Fshowall%3Dtrue#secsectitle0020 | Precancerous lesion subtypes (TA/TVA; SSL/HP); colorectal cancer subtypes (MSS CRC/microsatellite stable; MSI-H CRC/microsatellite instability -high); special/cross phenotypes (HM type TA/TVA/MSS; MSI-H/CIN cross subtype) | Visium |
| MISC57 | colon | https://www.cell.com/cell/fulltext/S0092 -8674(23)01222 -9?_returnURL=https%3A%2F%2Flinkinghub.elsevier.com%2Fretrieve%2Fpii%2FS0092867423012229%3Fshowall%3Dtrue#secsectitle0020 | Precancerous lesion subtypes (TA/TVA; SSL/HP); colorectal cancer subtypes (MSS CRC/microsatellite stable; MSI-H CRC/microsatellite instability -high); special/cross phenotypes (HM type TA/TVA/MSS; MSI-H/CIN cross subtype) | Visium |
| MISC58 | colon | https://www.cell.com/cell/fulltext/S0092 -8674(23)01222 -9?_returnURL=https%3A%2F%2Flinkinghub.elsevier.com%2Fretrieve%2Fpii%2FS0092867423012229%3Fshowall%3Dtrue#secsectitle0020 | Precancerous lesion subtypes (TA/TVA; SSL/HP); colorectal cancer subtypes (MSS CRC/microsatellite stable; MSI-H CRC/microsatellite instability -high); special/cross phenotypes (HM type TA/TVA/MSS; MSI-H/CIN cross subtype) | Visium |
| MISC62 | colon | https://www.cell.com/cell/fulltext/S0092 -8674(23)01222 -9?_returnURL=https%3A%2F%2Flinkinghub.elsevier.com%2Fretrieve%2Fpii%2FS0092867423012229%3Fshowall%3Dtrue#secsectitle0020 | Precancerous lesion subtypes (TA/TVA; SSL/HP); colorectal cancer subtypes (MSS CRC/microsatellite stable; MSI-H CRC/microsatellite instability -high); special/cross phenotypes (HM type TA/TVA/MSS; MSI-H/CIN cross subtype) | Visium |
| MISC63 | colon | https://www.cell.com/cell/fulltext/S0092 -8674(23)01222 -9?_returnURL=https%3A%2F%2Flinkinghub.elsevier.com%2Fretrieve%2Fpii%2FS0092867423012229%3Fshowall%3Dtrue#secsectitle0020 | Precancerous lesion subtypes (TA/TVA; SSL/HP); colorectal cancer subtypes (MSS CRC/microsatellite stable; MSI-H CRC/microsatellite instability -high); special/cross phenotypes (HM type TA/TVA/MSS; MSI-H/CIN cross subtype) | Visium |
| MISC66 | colon | https://www.cell.com/cell/fulltext/S0092 -8674(23)01222 -9?_returnURL=https%3A%2F%2Flinkinghub.elsevier.com%2Fretrieve%2Fpii%2FS0092867423012229%3Fshowall%3Dtrue#secsectitle0020 | Precancerous lesion subtypes (TA/TVA; SSL/HP); colorectal cancer subtypes (MSS CRC/microsatellite stable; MSI-H CRC/microsatellite instability -high); special/cross phenotypes (HM type TA/TVA/MSS; MSI-H/CIN cross subtype) | Visium |
| MISC67 | colon | https://www.cell.com/cell/fulltext/S0092 -8674(21)01381 -7?_returnURL=https%3A%2F%2Flinkinghub.elsevier.com%2Fretrieve%2Fpii%2FS0092867421013817%3Fshowall%3Dtrue | Precancerous lesion subtypes (conventional adenomas/ADs; serrated polyps/SERs); colorectal cancer subtypes (MSS CRC/microsatellite stable; MSI-H CRC/microsatellite instability -high) | Visium |
| MISC72 | colon | https://www.cell.com/cell/fulltext/S0092 -8674(23)01222 -9?_returnURL=https%3A%2F%2Flinkinghub.elsevier.com%2Fretrieve%2Fpii%2FS0092867423012229%3Fshowall%3Dtrue#secsectitle0020 | Precancerous lesion subtypes (TA/TVA; SSL/HP); colorectal cancer subtypes (MSS CRC/microsatellite stable; MSI-H CRC/microsatellite instability -high); special/cross phenotypes (HM type TA/TVA/MSS; MSI-H/CIN cross subtype) | Visium |
| MISC73 | colon | https://www.cell.com/cell/fulltext/S0092 -8674(23)01222 -9?_returnURL=https%3A%2F%2Flinkinghub.elsevier.com%2Fretrieve%2Fpii%2FS0092867423012229%3Fshowall%3Dtrue#secsectitle0020 | Precancerous lesion subtypes (TA/TVA; SSL/HP); colorectal cancer subtypes (MSS CRC/microsatellite stable; MSI-H CRC/microsatellite instability -high); special/cross phenotypes (HM type TA/TVA/MSS; MSI-H/CIN cross subtype) | Visium |
| TENX128 | colon | https://www.10xgenomics.com/datasets/visium -hd-cytassist -gene-expression -libraries -of-human -crc | Sigmoid colon and rectal cancer | Visium HD |
| TENX147 | colon | https://www.biorxiv.org/content/10.1101/2024.06.04.597233v1.full | Adjacent normal tissue; colorectal adenocarcinoma | Xenium |
| TENX148 | colon | https://www.biorxiv.org/content/10.1101/2024.06.04.597233v1.full | Adjacent normal tissue; colorectal adenocarcinoma | Xenium |
| TENX149 | colon | https://www.biorxiv.org/content/10.1101/2024.06.04.597233v1.full | Adjacent normal tissue; colorectal adenocarcinoma | Xenium |
| TENX152 | colon | https://www.biorxiv.org/content/10.1101/2024.06.04.597233v1.full | Adjacent normal tissue; colorectal adenocarcinoma | Visium |
| TENX70 | colon | https://www.10xgenomics.com/datasets/human -colorectal -cancer -11-mm-capture -area -ffpe -2-standard | Colorectal adenocarcinoma (T3, stage IIA) | Visium |
| TENX90 | colon | https://www.10xgenomics.com/datasets/visium -cytassist -gene-expression -libraries -of-post -xenium -human -colon -cancer -ffpe -using -the-human -whole -transcriptome -probe -set-2-standard | Colonic adenocarcinoma (Stage IIA) | Visium |
| TENX91 | colon | https://www.10xgenomics.com/datasets/visium -cytassist -gene-expression -libraries -of-post -xenium -human -colon -cancer -ffpe -using -the-human -whole -transcriptome -probe -set-2-standard | Colonic adenocarcinoma (Stage IIA) | Visium |
| TENX92 | colon | https://www.10xgenomics.com/datasets/visium -cytassist -gene-expression -libraries -of-post -xenium -human -colon -cancer -ffpe -using -the-human -whole -transcriptome -probe -set-2-standard | Colonic adenocarcinoma (Stage IIA) | Visium |
| ZEN38 | colon | https://www.biorxiv.org/content/10.1101/2023.01.23.525135v2 | Colorectal adenocarcinoma; molecular subtype classification: including CMS1, CMS2, CMS3, CMS4, and mixed subtype CMS1-CMS2 | Visium |
| ZEN42 | colon | https://www.biorxiv.org/content/10.1101/2023.01.23.525135v2 | Colorectal adenocarcinoma; molecular subtype classification: including CMS1, CMS2, CMS3, CMS4, and mixed subtype CMS1-CMS3 | Visium |
| ZEN44 | colon | https://www.biorxiv.org/content/10.1101/2023.01.23.525135v2 | Colorectal adenocarcinoma; molecular subtype classification: including CMS1, CMS2, CMS3, CMS4, and mixed subtype CMS1-CMS4 | Visium |
| ZEN45 | colon | https://www.biorxiv.org/content/10.1101/2023.01.23.525135v2 | Colorectal adenocarcinoma; molecular subtype classification: including CMS1, CMS2, CMS3, CMS4, and mixed subtype CMS1-CMS5 | Visium |
| INT12 | lymph node | https://pubmed.ncbi.nlm.nih.gov/35231421/ | Clear cell renal cell carcinoma (ccRCC) | Visium |
| INT13 | lymph node | https://pubmed.ncbi.nlm.nih.gov/35231421/ | Clear cell renal cell carcinoma (ccRCC) | Visium |
| INT14 | lymph node | https://pubmed.ncbi.nlm.nih.gov/35231421/ | Clear cell renal cell carcinoma (ccRCC) | Visium |
| INT15 | lymph node | https://pubmed.ncbi.nlm.nih.gov/35231421/ | Clear cell renal cell carcinoma (ccRCC) | Visium |
| INT16 | lymph node | https://pubmed.ncbi.nlm.nih.gov/35231421/ | Clear cell renal cell carcinoma (ccRCC) | Visium |
| INT19 | lymph node | https://pubmed.ncbi.nlm.nih.gov/35231421/ | Clear cell renal cell carcinoma (ccRCC) | Visium |
| INT21 | lymph node | https://pubmed.ncbi.nlm.nih.gov/35231421/ | Clear cell renal cell carcinoma (ccRCC) | Visium |
| INT22 | lymph node | https://pubmed.ncbi.nlm.nih.gov/35231421/ | Clear cell renal cell carcinoma (ccRCC) | Visium |
| INT24 | lymph node | https://pubmed.ncbi.nlm.nih.gov/35231421/ | Clear cell renal cell carcinoma (ccRCC) | Visium |
| INT4 | lymph node | https://pubmed.ncbi.nlm.nih.gov/35231421/ | Clear cell renal cell carcinoma (ccRCC) | Visium |
| INT5 | lymph node | https://pubmed.ncbi.nlm.nih.gov/35231421/ | Clear cell renal cell carcinoma (ccRCC) | Visium |
| INT7 | lymph node | https://pubmed.ncbi.nlm.nih.gov/35231421/ | Clear cell renal cell carcinoma (ccRCC) | Visium |
| INT8 | lymph node | https://pubmed.ncbi.nlm.nih.gov/35231421/ | Clear cell renal cell carcinoma (ccRCC) | Visium |
| NCBI681 | lymph node | https://www.nature.com/articles/s41467 -022-34581-2 | Invasive ductal carcinoma of the breast with lymph node metastasis; molecular subtype: luminal (A/B type), human epidermal growth factor receptor 2 overexpressed (HER2) | Visium |
| NCBI682 | lymph node | https://www.nature.com/articles/s41467 -022-34581-2 | Invasive ductal carcinoma of the breast with lymph node metastasis; molecular subtype: luminal (A/B type), human epidermal growth factor receptor 2 overexpressed (HER3) | Visium |
| NCBI683 | lymph node | https://www.nature.com/articles/s41467 -022-34581-2 | Invasive ductal carcinoma of the breast with lymph node metastasis; molecular subtype: luminal (A/B type), human epidermal growth factor receptor 2 overexpressed (HER4) | Visium |
| INT25 | prostate | None (internal) | | Visium |
| INT26 | prostate | None (internal) | | Visium |
| INT27 | prostate | None (internal) | | Visium |
| INT28 | prostate | None (internal) | | Visium |
| INT35 | prostate | None (internal) | | Visium |
| MEND140 | prostate | https://www.nature.com/articles/s41586 -022-05023-2 | Prostate cancer (including Gleason grade I and IV) and prostatic intraepithelial neoplasia, benign/pericancerous epithelial tissue | Visium |
| MEND141 | prostate | https://www.nature.com/articles/s41586 -022-05023-2 | Prostate cancer (including Gleason grade I and IV) and prostatic intraepithelial neoplasia, benign/pericancerous epithelial tissue | Visium |
| MEND142 | prostate | https://www.nature.com/articles/s41586 -022-05023-2 | Prostate cancer (including Gleason grade I and IV) and prostatic intraepithelial neoplasia, benign/pericancerous epithelial tissue | Visium |
| MEND143 | prostate | https://www.nature.com/articles/s41586 -022-05023-2 | Prostate cancer (including Gleason grade I and IV) and prostatic intraepithelial neoplasia, benign/pericancerous epithelial tissue | Visium |
| MEND145 | prostate | https://www.nature.com/articles/s41586 -022-05023-2 | Prostate cancer (including Gleason grade I and IV) and prostatic intraepithelial neoplasia, benign/pericancerous epithelial tissue | Visium |
| MEND146 | prostate | https://www.nature.com/articles/s41586 -022-05023-2 | Prostate cancer (including Gleason grade I and IV) and prostatic intraepithelial neoplasia, benign/pericancerous epithelial tissue | Visium |
| MEND147 | prostate | https://www.nature.com/articles/s41586 -022-05023-2 | Prostate cancer (including Gleason grade I and IV) and prostatic intraepithelial neoplasia, benign/pericancerous epithelial tissue | Visium |
| MEND148 | prostate | https://www.nature.com/articles/s41586 -022-05023-2 | Prostate cancer (including Gleason grade I and IV) and prostatic intraepithelial neoplasia, benign/pericancerous epithelial tissue | Visium |
| MEND150 | prostate | https://www.nature.com/articles/s41586 -022-05023-2 | Prostate cancer (including Gleason grade I and IV) and prostatic intraepithelial neoplasia, benign/pericancerous epithelial tissue | Visium |
| MEND151 | prostate | https://www.nature.com/articles/s41586 -022-05023-2 | Prostate cancer (including Gleason grade I and IV) and prostatic intraepithelial neoplasia, benign/pericancerous epithelial tissue | Visium |
| MEND153 | prostate | https://www.nature.com/articles/s41586 -022-05023-2 | Prostate cancer (including Gleason grade I and IV) and prostatic intraepithelial neoplasia, benign/pericancerous epithelial tissue | Visium |
| MEND154 | prostate | https://www.nature.com/articles/s41586 -022-05023-2 | Prostate cancer (including Gleason grade I and IV) and prostatic intraepithelial neoplasia, benign/pericancerous epithelial tissue | Visium |
| MEND156 | prostate | https://www.nature.com/articles/s41586 -022-05023-2 | Prostate cancer (including Gleason grade I and IV) and prostatic intraepithelial neoplasia, benign/pericancerous epithelial tissue | Visium |
| MEND157 | prostate | https://www.nature.com/articles/s41586 -022-05023-2 | Prostate cancer (including Gleason grade I and IV) and prostatic intraepithelial neoplasia, benign/pericancerous epithelial tissue | Visium |
| MEND158 | prostate | https://www.nature.com/articles/s41586 -022-05023-2 | Prostate cancer (including Gleason grade I and IV) and prostatic intraepithelial neoplasia, benign/pericancerous epithelial tissue | Visium |
| MEND160 | prostate | https://www.nature.com/articles/s41586 -022-05023-2 | Prostate cancer (including Gleason grade I and IV) and prostatic intraepithelial neoplasia, benign/pericancerous epithelial tissue | Visium |
| MEND162 | prostate | https://www.nature.com/articles/s41586 -022-05023-2 | Prostate cancer (including Gleason grade I and IV) and prostatic intraepithelial neoplasia, benign/pericancerous epithelial tissue | Visium |
| MEND18 | prostate | https://www.ncbi.nlm.nih.gov/pmc/articles/PMC9482614/ | Castration -resistant prostate cancer / Prostate cancer receiving androgen deprivation therapy (ADT) | Spatial Transcriptomics |
| MEND19 | prostate | https://www.ncbi.nlm.nih.gov/pmc/articles/PMC9482614/ | Castration -resistant prostate cancer / Prostate cancer receiving androgen deprivation therapy (ADT) | Spatial Transcriptomics |
| MEND24 | prostate | https://www.ncbi.nlm.nih.gov/pmc/articles/PMC9482614/ | Castration -resistant prostate cancer / Prostate cancer receiving androgen deprivation therapy (ADT) | Spatial Transcriptomics |
| MEND59 | prostate | https://www.nature.com/articles/s41467 -023-36071-5 | Prostate tumor | Visium |
| MEND60 | prostate | https://www.nature.com/articles/s41467 -023-36071-5 | Prostate tumor | Visium |
| MEND61 | prostate | https://www.nature.com/articles/s41467 -023-36071-5 | Prostate tumor | Visium |
| MEND62 | prostate | https://www.nature.com/articles/s41467 -023-36071-5 | Prostate tumor | Visium |
| TENX157 | prostate | https://www.10xgenomics.com/datasets/xenium -prime -ffpe -human -prostate | Prostate tumor | Xenium |
| TENX40 | prostate | https://www.10xgenomics.com/datasets/human -prostate -cancer -adenocarcinoma -with -invasive -carcinoma -ffpe -1-standard -1-3-0 | Invasive adenocarcinoma (Stage III, Gleason score: 7) | Visium |
| TENX46 | prostate | https://www.10xgenomics.com/datasets/human -prostate -cancer -acinar -cell-carcinoma -ffpe -1-standard | Invasive acinar cell carcinoma (Stage IV, Gleason score: 7) | Visium |
| MEND37 | skin | https://www.ncbi.nlm.nih.gov/pmc/articles/PMC7391009/ | Cutaneous squamous cell carcinoma (cSCC) | Visium |
| MEND38 | skin | https://www.ncbi.nlm.nih.gov/pmc/articles/PMC7391009/ | Cutaneous squamous cell carcinoma (cSCC) | Visium |
| MEND39 | skin | https://www.ncbi.nlm.nih.gov/pmc/articles/PMC7391009/ | Cutaneous squamous cell carcinoma (cSCC) | Visium |
| MEND40 | skin | https://www.ncbi.nlm.nih.gov/pmc/articles/PMC7391009/ | Cutaneous squamous cell carcinoma (cSCC) | Visium |
| NCBI760 | skin | https://www.ncbi.nlm.nih.gov/pmc/articles/PMC7391009/ | Cutaneous squamous cell carcinoma (cSCC) | Spatial Transcriptomics |
| NCBI764 | skin | https://www.ncbi.nlm.nih.gov/pmc/articles/PMC7391009/ | Cutaneous squamous cell carcinoma (cSCC) | Spatial Transcriptomics |
| NCBI765 | skin | https://www.ncbi.nlm.nih.gov/pmc/articles/PMC7391009/ | Cutaneous squamous cell carcinoma (cSCC) | Spatial Transcriptomics |
| NCBI766 | skin | https://www.ncbi.nlm.nih.gov/pmc/articles/PMC7391009/ | Cutaneous squamous cell carcinoma (cSCC) | Spatial Transcriptomics |
| NCBI768 | skin | https://www.ncbi.nlm.nih.gov/pmc/articles/PMC7391009/ | Cutaneous squamous cell carcinoma (cSCC) | Spatial Transcriptomics |
| NCBI769 | skin | https://www.ncbi.nlm.nih.gov/pmc/articles/PMC7391009/ | Cutaneous squamous cell carcinoma (cSCC) | Spatial Transcriptomics |
| NCBI770 | skin | https://www.ncbi.nlm.nih.gov/pmc/articles/PMC7391009/ | Cutaneous squamous cell carcinoma (cSCC) | Spatial Transcriptomics |
| ZEN78 | skin | https://www.biorxiv.org/content/10.1101/2023.07.27.550727v1 | BRAF V600E mutant metastatic melanoma | Visium |
| ZEN79 | skin | https://www.biorxiv.org/content/10.1101/2023.07.27.550727v1 | BRAF V601E mutant metastatic melanoma | Visium |
| ZEN80 | skin | https://www.biorxiv.org/content/10.1101/2023.07.27.550727v1 | BRAF V602E mutant metastatic melanoma | Visium |
| ZEN81 | skin | https://www.biorxiv.org/content/10.1101/2023.07.27.550727v1 | BRAF V603E mutant metastatic melanoma | Visium |
